# Plasmodesmata-located proteins regulate plasmodesmal function at specific cell interfaces in Arabidopsis

**DOI:** 10.1101/2022.08.05.502996

**Authors:** Zhongpeng Li, Su-Ling Liu, Christian Montes-Serey, Justin W. Walley, Kyaw Aung

## Abstract

Plasmodesmata (PD) are membrane-lined channels connecting adjoining plant cells. PD control symplasmic intercellular communication by allowing molecules to move between cells. Plant polysaccharide callose (ß-1,3-glucan) is deposited at PD, affecting plasmodesmal function; however, the regulation of PD at different cell interfaces is largely unknown. This study discovered that two PD-located proteins, PDLP5 and PDLP6, are expressed in non-overlapping cell types. The constitutive expression of PDLP5 and PDLP6 results in the overaccumulation of PD callose at different cell interfaces and starch hyperaccumulation in different cell types within mature leaves. Using a proximity labeling approach, we identified sucrose synthase 6 (SUS6) as a functional partner of PDLP6. We further demonstrated that PDLP6 physically and genetically interacts with SUS6. In addition, callose synthase 7 (CalS7) interacts with both SUS6 and PDLP6 and is required for PDLP6’s function. We propose that PDLP6-SUS6-CalS7 forms a callose synthase complex in the vasculature to regulate the plasmodesmal function.

## Introduction

Different cell types have evolved specialized functions to effectively support a multicellular organism to operate as a unit. Cells need to maintain their own cellular identity while effectively communicating with surrounding cells and tissues. Communication between cells can achieve through two major pathways: the apoplastic and symplastic routes. In an apoplastic pathway, molecules released from a cell in the apoplast are perceived by or enter other cells through various mechanisms. A symplastic pathway allows the movement of molecules between cells through physical channels connecting adjoining cells (Lucas et al., 2009; Ariazi et al., 2017).

Plasmodesmata (PD) are membrane-lined channels connecting adjoining plant cells, allowing the exchange of signals and resources among cells (Lucas et al., 2009). The PD-dependent communication among cells is fundamental for developmental regulation and stress responses in plants (Sager and Lee, 2018). PD also allow translocating of photosynthetic products from mature leaves (source tissues) to non-photosynthetic organs (sink tissues) in plants (Comtet et al., 2017; Liesche et al., 2017). In Arabidopsis, chloroplasts within mesophyll cells (MCs) are mainly responsible for converting solar energy into chemical energy, producing sugars. The carbohydrate can be stored temporarily in the chloroplasts as starch and translocated to different parts of the plants. Sugars move from MCs to sieve elements (SEs) through different pathways to translocate photosynthetic products. It’s generally believed that photosynthetic products move symplasmically through PD between photosynthetic cells, including MCs and bundle sheath cells (BSs), and are loaded into phloem for long-distance transport (Lucas et al., 2013; Braun, 2022). In Arabidopsis, three major cell types compose phloem: phloem parenchyma cells (PPs), companion cells (CCs), and SEs. Single-cell transcriptomes of Arabidopsis vascular tissues revealed that there might be at least two subpopulations of BS, PP, and CC (Kim et al., 2021; Braun, 2022).

Within phloem, symplasmic and apoplasmic phloem loading pathways have been reported for different plant species. Symplasmic phloem loading moves sugars through PD connecting different phloem cell types. Arabidopsis utilizes apoplasmic phloem loading, in which sugars are exported from PPs into the extracellular space by sucrose-proton cotransporters (SWEETs) (Chen et al., 2012) and imported from the apoplast into CCs by sucrose-proton symporter 2 (SUC2) (Srivastava et al., 2008). Arabidopsis *sweet11;12* and *suc2* mutants are compromised in sugar translocation from source to sink, resulting in hyperaccumulation of starch in mature leaves and stunted plant growth (Srivastava et al., 2008; Chen et al., 2012). In addition to the sugar transporter, a choline transporter-like 1 (CHER1) also affects sugar translocation. Arabidopsis *cher1-4* mutant has a reduced number of PD between different cell types in leaves (Kraner et al., 2017). The mutant is compromised in sugar movement and loading in mature leaves, resulting in starch hyperaccumulation and stunted plant growth (Kraner et al., 2017). Sugars from CC moves into SE through PD (Lucas et al., 2013; Liesche et al., 2017), completing the sugar-loading process. The enucleated SE transports sugars to other parts of the plants (Oparka and Turgeon, 1999; Chen et al., 2015; Zhang et al., 2018).

It is well documented that callose (ß-1,3-glucan) is deposited at PD within cell walls to restrict the plasmodesmal aperture (Wu et al., 2018), regulating the PD function. Callose synthases (CalSs) catalyze callose biosynthesis. It’s been proposed that CalSs form multisubunit enzyme complexes with sucrose synthases (SUSs) (Amor et al., 1995), which generate uridine diphosphate glucose (UDPG), and others. CalSs utilize UDPG as a glucose donor to synthesize callose (Li et al., 2003; Brownfield et al., 2007; De Storme and Geelen. 2014). In addition, recent findings highlighted the roles of PD-located proteins (PDLPs) in modulating the plasmodesmal functions. PDLPs are type I membrane proteins containing a signal peptide, two DOMAINS OF UNKNOWN FUNCTION 26 (DUF26), and a transmembrane domain followed by a short stretch of cytoplasmic tail (Thomas et al., 2008). The extracellular domain of PDLP5 exhibits high structural similarity to fungal lectins, which bind mannose; however, it is yet to demonstrate the plant polysaccharide binding ability of PDLP5 (Vaattovaara et al., 2019).

Despite their important roles in regulating callose biosynthesis at PD, molecular functions of PDLPs are unknown. Here, we characterized the functions of PDLPs using a gain-of-function approach in Arabidopsis. The overexpression of PDLP5 and PDLP6 hinders the PD-dependent movement of sugar in different cell types, resulting in starch hyperaccumulation in mature leaves. We further demonstrated that PDLP5 and PDLP6 are expressed in and function at different cell interfaces. In addition, we identified tissue-specific functional partners of PDLP6, SUS6 and CalS7. Our findings suggest that PDLP6-SUS6-CalS7 functions specifically in the vasculature, likely in SE, to regulate the plasmodesmal function.

## Results

### The overexpression of PDLP6 affects plant growth and PD-mediated long-distance symplastic transportation

To better understand the function of PDLPs, we individually overexpressed all eight members of PDLP (PDLP1-8) in Arabidopsis wild-type Col-0 using a 35S promoter. We chose to overexpress the PDLPs to avoid a gene redundancy issue within the protein family. The PDLPs were fused with His and Flag (HF) tag. Here, we report the characterization of *Pro35S:PDLP5-HF* and *Pro35S:PDLP6-HF* (hereafter *PDLP5-HF* and *PDLP6-HF*). Consistent with a previous report (Lee et al., 2011), a stunted growth phenotype was observed for *PDLP5-HF* lines compared to a vector control *Pro35S:HF-YFP* (hereafter *HF-YFP*; Supplemental Figure 1, A and E). Immunoblot analyses assessed the levels of protein expression in the transgenic plants. Similar to a previous report (Lee et al., 2011), we observed a negative correlation between the expression level of PDLP5 and the growth phenotype (Supplemental Figure 1, A and B). In addition, *PDLP6-HF* transgenic lines exhibit a stunted plant growth phenotype (Figure 1A). Like PDLP5, transgenic plants with a higher expression level of PDLP6 exhibit more severe dwarfism in plant growth (Figure 1, A and B). The findings suggest that the overexpression of PDLP5 and PDLP6 affects plant growth. As PDLP5 negatively regulates the plasmodesmal function (De Storme and Geelen. 2014), we examined the PD permeability in *PDLP6-HF* using a CFDA feeding assay. 5(6)-carboxyfluorescein diacetate (CFDA) is a symplastic tracer commonly used to determine the PD-dependent movement of the dye among plant tissues (Oparka et al., 1994). Plant cells can rapidly uptake CFDA and cleave it into CF using intracellular esterases. Since CF is a fluorescent molecule and membrane-impermeable, CF’s movement in plant cells depends on PD. CFDA was applied to a cotyledon of two-week-old Arabidopsis seedlings. The presence of CF in root tips of CFDA-fed seedlings was detected and quantified to determine the PD-mediated long-distance symplamic transportation in the transgenic plants. Compared to wild-type Col-0, we observed a significant decrease in CF signals in the root tips of *PDLP6-HF* lines (Figure 1, D and E). The findings suggest that PDLP6 also negatively regulates the plasmodesmal function.

**Figure 1.**
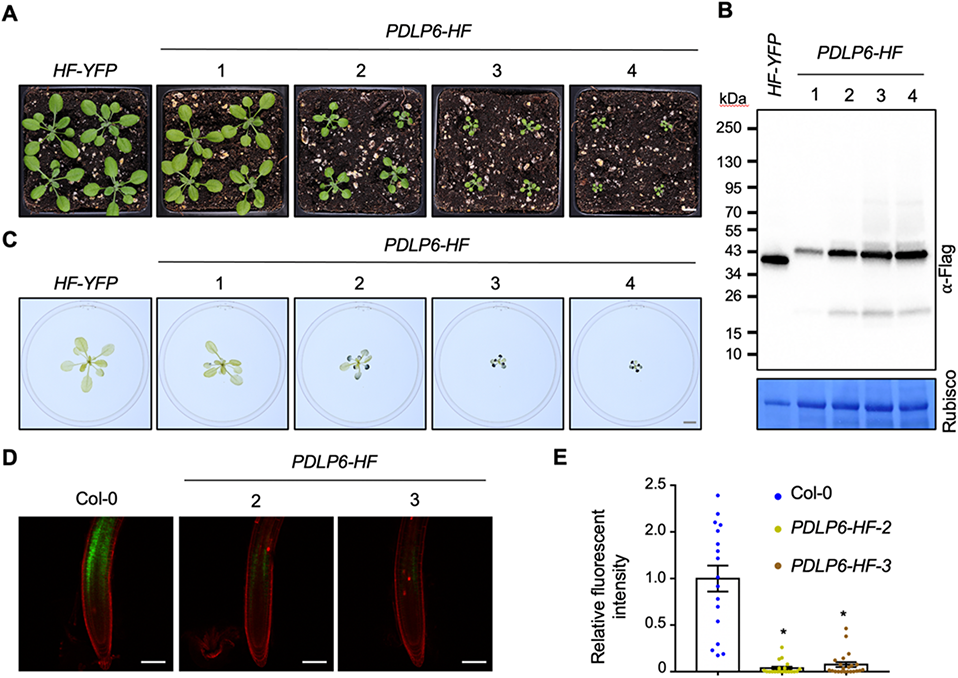
The overexpression of PDLP6 leads to growth defects, starch overaccumulation, and reduced PD permeability. (**A**) 3-week-old Arabidopsis plants were grown under a light intensity used for standard Arabidopsis growth (110 µmol m^-2^ s^-1^). Images were taken using the same magnification. Scale bar = 1 cm. Four independent transgenic lines (1-4) were shown. *HF-YFP* and *PDLP6-HF* refer to *Pro35S:HF-YFP* and *Pro35S:PDLP6-HF,* respectively. (**B**) Immunoblot analysis detects the expression of PDLP6-HF in four independent transgenic plants. An anti-Flag antibody was used to detect the expression of Flag-fusion proteins. Rubisco serves as a loading control. (**C**) Starch accumulation of plants shown in a. Tissues were harvested at the end of the night and stained using Lugol’s iodine solution. Images were taken using the same magnification. Scale bars = 1 cm. (**D**) PD permeability was determined by a CFDA loading assay. CF signals are shown in green. Propidium Iodide (PI) was used as a root cell wall stain. PI signals are shown in red. Scale bars = 100 µm. (**E**) CF fluorescent signal intensity in roots was quantified using Fiji. The plot shows the mean with SEM. Col-0, n = 17; *PDLP6-HF-2*, n = 20; and *PDLP6-HF-3*, n = 22. Asterisks indicate statistically significant differences (*t*-Test; two-paired; *P* < 0.01).

### The overexpression of PDLP5 or PDLP6 leads to starch hyperaccumulation in mature leaves

In addition to a stunted growth phenotype, *PDLP6-HF* also exhibits anthocyanin overaccumulation in mature leaves and late-flowering phenotypes (Supplemental Figure 1, C-G). These growth phenotypes are reminiscent of Arabidopsis mutants compromised in sugar translocation, *i.e., sweet11;12* and *suc2* (Srivastava et al., 2008; Chen et al., 2012). We thus hypothesized that the overexpression of PDLP5 and PDLP6 affects the PD-dependent sugar translocation in mature leaves. Transitory starch is the major product of photosynthesis and accumulates inside chloroplasts of source leaves during the day. It is hydrolyzed to maltose and glucose to synthesize sucrose, which is then transported via phloem to sink tissues during the following night (Smith et al., 2005; Stitt and Zeeman, 2012; Noronha et al., 2018; Geiger, 2020). When sugar loading is compromised, starch hyperaccumulates in mature leaves. Using Lugol’s iodine solution, we determined starch content in the transgenic plants (Tran et al., 2019). Tissues were harvested at the end of the night for the starch staining since overall starch accumulation in mature leaves is at its lowest (Graf et al., 2010; Chen et al., 2012). We included *cher1-4* and *sweet11;12* mutants as the mutants hyperaccumulate starch in mature leaves (Chen et al., 2012; Kraner et al., 2017). Wild-type Col-0 and *HF-YFP* were used as negative controls. Under a light intensity used for a standard Arabidopsis growth (110 µmol m^-2^ s^-1^), we reproduced the starch hyperaccumulation phenotype in *cher1-4* (Supplemental Figure 2A). We also found that *PDLP6-HF* hyperaccumulates starch in mature rosette leaves, exhibiting a dark blue color (Figure 1C and Supplemental Figure 2A). Similar to a plant growth phenotype, transgenic plants with a higher expression level of PDLP6 contain more starch in their mature leaves (Figure 1, A-C).

We didn’t observe starch hyperaccumulation phenotype in *PDLP5-HF* and *sweet11;12* mutant when the plants were grown under a standard Arabidopsis growth condition (Supplemental Figure 2A). As *sweet11;12* was reported to exhibit starch hyperaccumulation only when the plants were grown under high light conditions (Chen et al., 2012), we irradiated the mutants and transgenic plants with high light intensity (200 µmol m^-2^ s^-1^) for a week before the starch staining. The starch hyperaccumulation phenotype became apparent in *sweet11;12* and *PDLP5-HF* (Figure 2A). We also quantified the starch and sucrose content in mature leaves (Leach and Braun, 2016). Compared with *HF-YFP,* a higher level of both sucrose and starch was detected in *PDLP5-HF* and *PDLP6-HF* (Supplemental Figure 2B).

**Figure 2.**
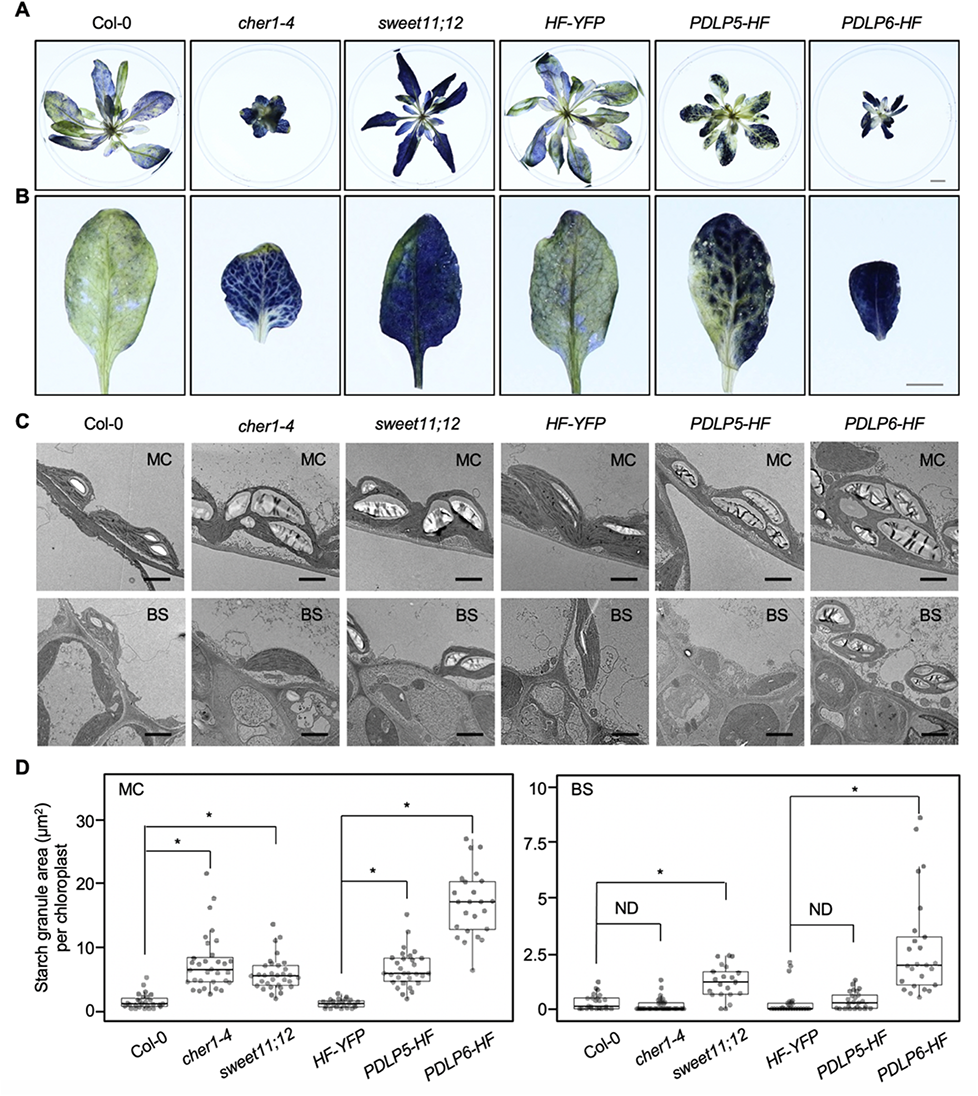
The overexpression of PDLP5 and PDLP6 leads to starch overaccumulation in different leaf cell types. (**A**) Starch staining of 5-week-old Arabidopsis plants. 4-week-old Arabidopsis plants were irradiated with high light intensity (200 µmol m^-2^ s^-1^) for a week. Samples were collected at the end of the night for starch staining using Lugol’s iodine solution. *HF-YFP, PDLP5-HF* and *PDLP6-HF* refer to *Pro35S:HF-YFP, Pro35S:PDLP5-HF,* and *Pro35S:PDLP6-HF,* respectively. Images were taken using the same magnification. Scale bar = 1 cm. (**B**) Starch accumulation patterns in different genotypes. Images were captured with the same magnification. Scale bar = 0.5 cm. (**C**) TEM images show the starch granules in chloroplasts in mesophyll and bundle sheath cells of mature leaves. MC: mesophyll cell; BS: Bundle sheath cell. Scale bars = 2 µm. (**D**) Quantification of starch granule areas. Each dot represents the total starch granule area in one chloroplast. Col-0, n = 27; *cher1-4*, n = 32; *sweet11;12*, n = 32; *HF-YFP*, n = 24; *PDLP5-HF*, n = 30; and *PDLP6-HF*, n = 24 for MC. Col-0, n = 23; *cher1-4*, n = 38; *sweet11;12*, n = 21; *HF-YFP*, n = 26; *PDLP5-HF*, n = 25; and *PDLP6-HF*, n = 25 for BS. Asterisks indicate statistically significant differences (Mann-Whitney *U* Test; two-paired; *P* < 0.01). ND, no statistical difference.

To further characterize the function of PDLP6 in regulating the symplamtic movement of sugars, we generated *pdlp6-1* and *pdlp6-2* mutants using CRISPR/Cas9 technology (Tsutsui and Higashiyama, 2017). *Pdlp6-1* and *pdlp6-2* carry 26 bp deletion and a G to T mutation around a gRNA target site, respectively (Supplemental Figure 3A). The deletion and mutation lead to a frameshift and premature termination of PDLP6 translation. The development and growth of *pdlp6* mutants are indistinguishable from that of wild-type Col-0 (Supplemental Figure 3B). Using Lugol’s iodine solution, *pdlp6* mutants exhibit a similar level of starch in mature leaves compared to that of wild-type Col-0 (Supplemental Figure 3C). To directly visualize starch granules in chloroplasts within mature leaves, we performed transmission electron microscopy (TEM). We observed significantly fewer starch grains in *pdlp6-1* mutant compared to that of wild-type Col-0 and *pdlp5* mutant (Supplemental Figure 3, D and E). The findings suggest that the absence of functional PDLP6 allows the more efficient movement or loading of sugars in mature leaves.

**Figure 3.**
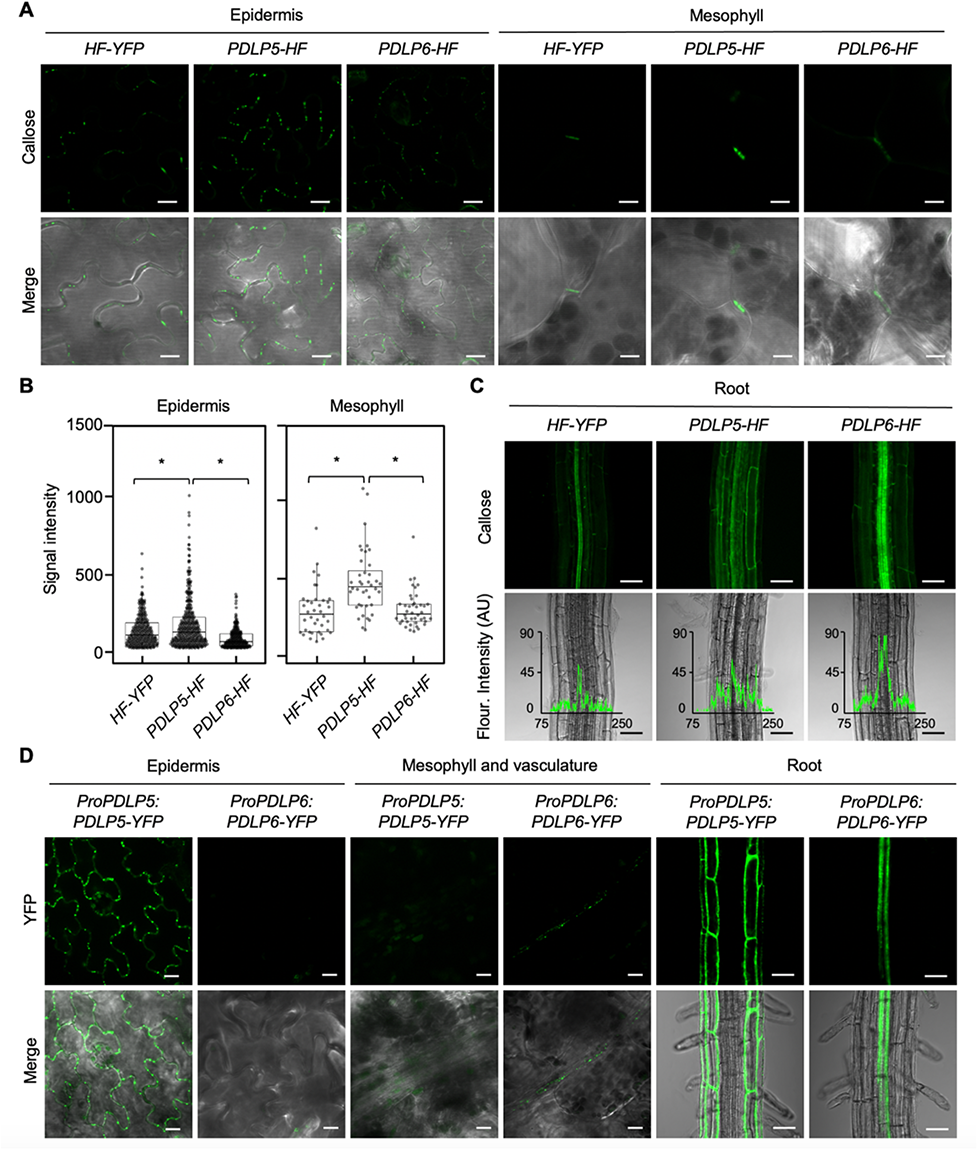
PDLP5 and PDLP6 function at different cell interfaces and express in different tissues. (**A**) Callose accumulation between leaf epidermal or mesophyll cells. Green signals represent aniline blue-stained callose at PD (top panel). Merged images show the signals from callose and bright-field (lower panel). Scale bars = 10 µm. (**B**) Quantitative data show callose accumulation between epidermal cells and mesophyll cells. Asterisks indicate statistically significant differences (Mann-Whitney *U* Test; *P* < 0.05). *HF-YFP*, n = 283; *PDLP5-HF*, n = 359; and *PDLP6-HF*, n = 233 for epidermis. *HF-YFP*, n = 38; *PDLP5-HF*, n = 40; and *PDLP6-HF*, n = 42 for mesophyll cell. (**C**) Callose accumulation in roots of the transgenic plants. Green signals represent callose accumulation in roots (top panel). Merged images show the bright field and fluorescence intensity profile (arbitrary unit: AU) of callose accumulation along the horizontal lines (lower panel). Numbers on the X-axis indicate the distance (µm) across the region analyzed. Scale bars = 50 µm. (**D**) The tissue-specific expression of PDLP5-YFP and PDLP6-YFP. The fusion proteins were driven by their own native promoter. Confocal images were captured from 2-week-old Arabidopsis seedlings. Green signals represent the expression of the YFP fusion proteins in different cell types (top panel). Merged images show the signals from YFP and bright-field (lower panel). Scale bars for epidermis and vasculature = 10 µm. Scale bars for root = 50 µm.

### The overexpression of PDLP5 or PDLP6 leads to distinct starch hyperaccumulation patterns

In addition, we observed distinct starch accumulation patterns in mature leaves of *PDLP5-HF*, *PDLP6-HF*, *sweet11;12*, and *cher1-4*. *PDLP6-HF* and *sweet11;12* accumulate starch evenly, whereas *PDLP5-HF* and *cher1-4* lack starch accumulation in and around vascular tissues in mature leaves (Figure 2B). As SWEET11 and SWEET12 specifically express at PPs within phloem (Chen et al., 2012), sugars can likely move from MCs to PPs in the *sweet11;12* mutant. *cher1-4*, on the other hand, has a reduced number of PD between MCs to allow the efficient symplastic movement of sugars (Kraner et al., 2017); consequently, most sugars might be trapped within MCs. To determine cell type-specific starch accumulation in the mutants and transgenic plants, leaf discs from high light-treated plants were subjected to histological sectioning and starch staining. Periodic acid/Schiff (PAS) reagent labels polysaccharides in the cell wall and starch grains in chloroplasts. Consistent with the whole tissue staining (Figure 2B), *cher1-4*, *sweet11;12*, *PDLP5-HF*, and *PDLP6-HF* exhibit darker PAS-stained starch grains in chloroplasts of MCs compared to that of Col-0 and *HF-YFP* (Supplemental Figure 2C). In line with our prediction, *cher1-4* and *PDLP5-HF* contain fewer starch in BSs, whereas *sweet11;12* and *PDLP6-HF* contain many PAS-stained starch grains in their BSs (Supplemental Figure 2C). To visualize starch granules in different cell types within mature leaves, we performed a TEM analysis. *cher1-4*, *sweet11;12*, *PDLP5-HF*, and *PDLP6-HF* contain larger starch granules in chloroplasts within MCs compared to their controls, whereas enlarged starch granules were only observed in BSs of *sweet11;12* and *PDLP6-HF* (Figure 2, C and D). Together, our findings suggest that the overexpression of PDLP5 and PDLP6 blocks the movement of sugars at different cell types during sugar translocation in mature leaves.

To test the important roles of symplamic sugar movement from MCs to phloem, we performed genetic crosses between *cher1-4* and *PDLP6-HF* and *cher1-4* and *sweet11;12*. F_3_ populations of the crosses were subjected to a PCR genotyping assay to identify homozygous lines for *cher1-4* and *sweet11;12* (Supplemental Figure 4, A and C). As the expression level of PDLP6-HF determines the plant growth and starch accumulation phenotypes of the transgenic plants, we examined the expression level of the fusion proteins using immunoblot analysis (Supplemental Figure 4B). As shown in Figure 4D, the starch accumulation pattern in *cher1-4 PDLP6-HF* and *cher1-4 sweet11;12* resembles that in *cher1-4*, which lacks starch hyperaccumulation in and around vascular tissues. In addition, *cher1-4 PDLP6-HF* and *cher1-4 sweet11;12* show a more stunted growth phenotype compared to cher1-4 mutant (Supplemental Figure 4D). The findings highlight the important roles of symplasmic sugar movement from mesophyll cells to phloem.

**Figure 4.**
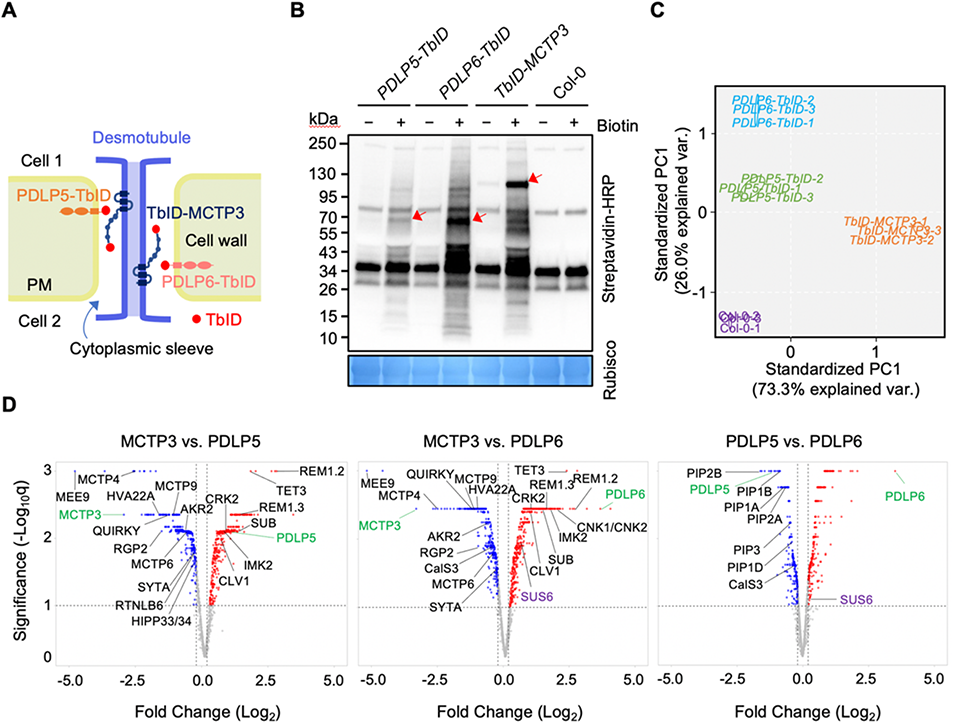
A proximity labeling assay to identify functional partners of plasmodesmal proteins. (**A**) A diagram shows the localization of PDLP5-TbID, PDLP6-TbID, and TbID-MCTP3 fusion proteins to different membranes of PD. *PDLP5-TbID*, *PDLP6-TbID,* and *TbID-MCTP3* refer to *ProUBQ10:PDLP5-TurboID-3xFlag*, *ProUBQ10:PDLP6-TbID-3xFlag,* and *ProUBQ10:TbID-3xFlag-MCTP3,* respectively. (**B**) 2-week-old seedlings of wild-type Col-0 and Arabidopsis transgenic plants were subjected to biotin treatment and immunoblot analysis using a streptavidin-HRP antibody to determine the activity of the TbID fusion proteins. Rubisco serves as a loading control. Red arrows indicate the biotinylated TbID fusion proteins. (**C**) Principal component analysis (PCA) was performed and visualized using the ggbiplot R package as part of the TMT-NEAT analysis pipeline. Three replicates (1-3) were analyzed for each genotype. (**D**) Volcano plots show significantly enriched proteins in PDLP5, PDLP6, and MCTP3 samples. Candidates were filtered using cutoffs log_2_FC > 0.2 or < −0.2 and q < 0.1. Plots were generated using VolcaNoseR. A few of the known and putative PD-associated proteins are labeled.

Together, these results suggest that sugar movement is likely blocked at different steps in different genotypes: (1) sugar movement between MC-MC is blocked in *cher1-4* as PD numbers between MCs are greatly reduced in the mutant, (2) sugar movement between MC-MC and/or MC-BS is blocked in *PDLP5-HF* since no significant starch accumulation was observed in BSs, (3) sugar movement between PP-CC is blocked in *sweet11;12* as sucrose efflux from PPs is compromised, and (4) sugar movement in vascular tissues (between BS-PP or CC-SE or SE-SE) is blocked in *PDLP6-HF* since starch hyperaccumulation in BSs was observed.

### PDLP5 and PDLP6 promote PD callose deposition at different cell interfaces

While the molecular mechanism underlying PDLP function is unclear, the expression level of PDLP5 is highly correlated with callose accumulation at PD in Arabidopsis and *N. benthamiana* (Lee et al., 2011; Li et al., 2021). To further investigate the cell type-specific functions of PDLP5 and PDLP6, we examined whether the overexpression of PDLP5 or PDLP6 leads to the overaccumulation of PD callose at specific cell interfaces. *PDLP5-HF*, *PDLP6-HF*, and *HF-YFP* leaves were stained with aniline blue to detect callose accumulation at PD. Consistent with the previous report (Lee et al., 2011), we observed a higher callose accumulation between epidermal cells (ECs) in *PDLP5-HF* compared to *HF-YFP*. We also detected a higher callose level at PD connecting MCs in *PDLP5-HF* (Figure 3, A and B). As PDLP6 is expected to function specifically in the vasculature, we examined the callose accumulation between vascular cells. The roots of the transgenic plants were stained with aniline blue and imaged with confocal microscopy due to their tissue transparency. *PDLP6-HF* exhibits the highest callose accumulation in vascular tissues (Figure 3C). In contrast, a much higher callose accumulation was observed in the epidermis and cortex of *PDLP5-HF* than in *HF-YFP* and *PDLP6-HF* (Figure 3C). Fluorescence intensity profiles were consistent among data collected from 25 individual transgenic plants for each genotype (Supplemental Figure 5). These findings support that the overexpression of PDLP5 and PDLP6 leads to a higher PD callose accumulation at PD at specific cell interfaces.

### PDLP5 and PDLP6 express in different tissues/cell types

The distinct patterns of starch accumulation and corresponding PD callose accumulation at specific cell interfaces in *PDLP5-HF* and *PDLP6-HF* prompted us to hypothesize that endogenous PDLP5 and PDLP6 express in different cell types. To test the hypothesis, we generated native promoter-driven *ProPDLP5:PDLP5-YFP* and *ProPDLP6:PDLP6-YFP* transgenic Arabidopsis plants. Using confocal microscopy, the expression of PDLP5-YFP was mainly detected between ECs, whereas no PDLP5-YFP signals were observed in MCs and leaf vasculature (Figure 3D). PDLP6-YFP, on the other hand, was detected specifically in leaf vasculature (Figure 3D). Similarly, PDLP6-YFP expresses specifically in root vasculature, whereas PDLP5-YFP was detected in the epidermis and cortex (Figure 3D). We also used native promoters of *PDLP5* or *PDLP6* (*ProPDLP5* or *ProPDLP6*) to drive the expression of GUS. Histochemical staining results showed that the PDLP5 promoter is active in ECs and MCs in the leaf and epidermis and cortex in the root, while the PDLP6 promoter is active specifically in vasculature tissues in both leaf and root (Supplemental Figure 6).

Using a constitutive promoter *Pro35S*, PDLP5-HF and PDLP6-HF are expected to express in most cell types. The assumption was supported by the findings that PDLP5-YFP and PDLP6-YFP were detected in most cell types in leaves and roots of Arabidopsis transgenic plants, *Pro35S:PDLP5-YFP* and *Pro35S:PDLP6-YFP* (Supplemental Figure 7). Our findings raise an intriguing question of how the expression of the PDLPs in most cell types gives rise to the cell type-specific functions of the proteins. As PDLPs are not predicted to catalyze callose biosynthesis, the enzymes or proteins involved in synthesizing callose functioning together with the PDLPs might express only in a cell type-specific manner.

### Identification of putative PDLP5 and PDLP6 functional partners using a proximity labeling (PL) approach

To identify functional partners of PDLP5 and PDLP6, we conducted a PL assay. Recent studies have successfully used the method to identify functional or interacting partners of plant proteins, including nuclear membrane proteins (Mair et al., 2019; Zhang et al., 2019; Huang et al., 2020). We generated the following two transgenic lines: *ProUBQ10:PDLP5-TurboID-3*×*Flag* and *ProUBQ10:PDLP6-TbID-3*×*Flag* (hereafter *PDLP5-TbID* and *PDLP6-TbID*). PDLPs are localized to the plasma membrane within plasmodesmata (Thomas et al., 2008) (PD-PM; Figure 4A). Since multiple C2 domains and transmembrane protein 3 (MCTP3) is targeted to the ER membrane of PD (PD-ER) (Brault et al., 2019), we generated and included *ProUBQ10:TbID-3*×*Flag-MCTP3* as a control (hereafter *TbID-MCTP3*; Figure 4A). We also included wild-type Col-0 as another control to detect the baseline of biotinylated proteins in Arabidopsis. The expression and enzymatic activity of the TbID fusion proteins in the transgenic plants were detected using immunoblot analysis (Figure 4B and Supplemental Figure 8A). Total biotinylated proteins were enriched and subjected to LC/MS-MS for protein identification. Principal component analysis (PCA) showed a clear separation of the samples by genotype (Figure 4C). MCTP3 is distinctly separated from PDLP5 and PDLP6, as well as Col-0. PDLP5, PDLP6, and MCTP3 are specifically enriched in the transgenic plants expressing the corresponding fusion proteins (Figure 4D and Supplemental Data set 1). Many known PD-localized proteins were enriched by PDLP5, PDLP6, and/or MCTP3 (Figure 4D and Supplemental Figure 8, B-D). Several proteins with potential functions at PD-ER (e.g., MCTP4, RTNLB8, RTNLB6, HVA22a, and HVA22c) were significantly enriched by MCTP3. PDLP5 and PDLP6, on the other hand, enrich many confirmed PD proteins targeted to PD-PM (e.g., CLV1, REM1.2, and REM1.3) (Figure 4D). Our findings suggest that the PL assay is a powerful tool to identify functional partners of PD proteins. Given the juxtaposition of the two membrane systems, the PM and the ER, the PL assay shows promise in resolving the functional protein complexes at a nanometer resolution within PD.

### PDLP6 functions together with SUS6

To maximize the identification of PDLP functional partners, we chose to express the PDLP5-TbID and PDLP6-TbID fusion proteins with a constitutive promoter *UBQ10*. Consequently, the two baits enrich a large set of similar proteins (Figure 4D and Supplemental Data set 1). Despite the similarity, PDLP5 and PDLP6 also enrich specific putative functional partners. PDLP5 significantly enriches several members of plasma membrane intrinsic proteins (PIPs/aquaporins), while PDLP6 significantly enriches SUS6 (Figure 4D). Given the potential roles of SUS6 in callose biosynthesis, we further characterized the relationship between PDLP6 and SUS6. Arabidopsis genome encodes six members of SUSs, sequentially named SUS1-6 (Baud et al., 2004). Among them, SUS6 is expressed specifically in SE (Yao et al., 2020). The findings suggest that PDLP6 might function together with SUS6 to regulate callose biosynthesis in vascular tissues, likely in SE. We first determined the PD association of SUS6 by transiently expressing SUS6-superfolder GFP (sfGFP) in *N. benthamiana*. Most of the SUS6-sfGFP fusion proteins were detected in the cytosol as aggregates, while some signals co-localized with aniline-blue-stained callose (Figure 5A), suggesting that SUS6 can localize to PD. Moreover, the transient overexpression of PDLP6-HF increased the PD association of SUS6-sfGFP, showing significantly more sfGFP signals overlapped with aniline blue-stained callose (Supplemental Figure 9).

**Figure 5.**
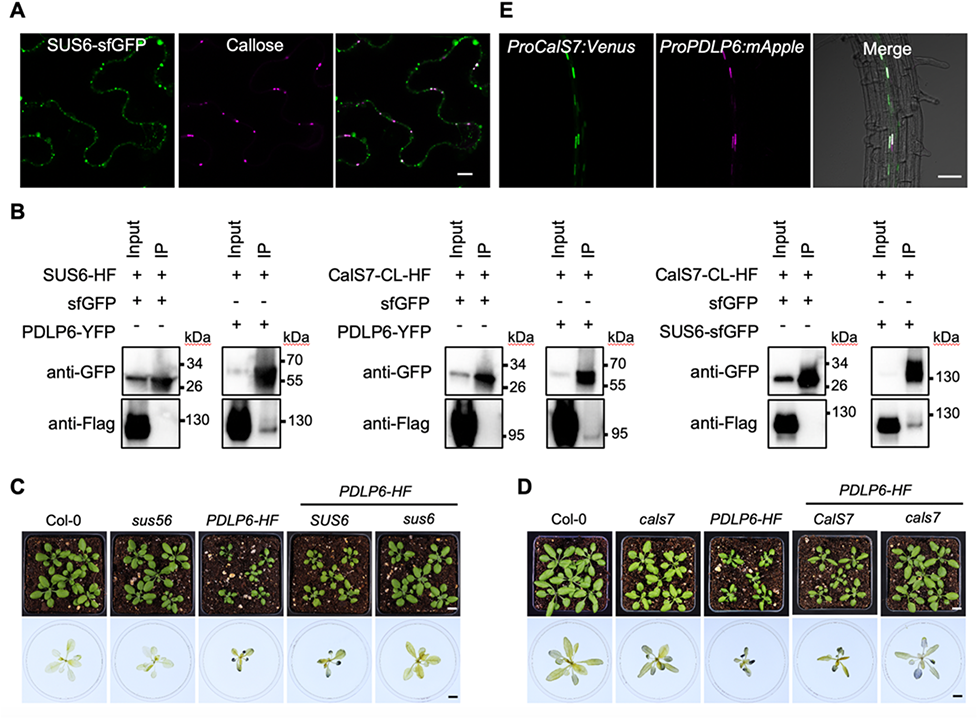
PDLP6 functions together with SUS6 and CalS7. (**A**) Confocal images show the PD localization of SUS6-sfGFP. Agrobacteria harboring *SUS6-sfGFP* were infiltrated into *N. benthamiana*. Callose was stained using aniline blue. Scale bars = 10 µm. (**B**) Co-IP assay indicates the interaction between PDLP6 and SUS6, PDLP6 and CalS7, and SUS6 and CalS7. Agrobacteria harboring different plasmids were co-infiltrated into *N. benthamiana*. Samples were collected 2 days after infiltration and subjected to Co-IP. (**C**) Genetic interactions between *PDLP6* and *SUS6*. (**D**) Genetic interactions between *PDLP6* and *CalS7*. Homozygous lines were isolated from F_3_ progenies of the genetic crosses of *sus56* with *Pro35S:PDLP6-HF (PDLP6-HF)* and *cals7* with *PDLP6-HF*. Images in the lower panel show the starch accumulation phenotype. Plants were collected at the end of the night for starch staining using Lugol’s iodine solution. Images were taken using the same magnification. Scale bar = 1 cm. (**E**) *CalS7* and *PDLP6* promoters are active in the same root cell type. Confocal images show the expression of Venus and mApple in Arabidopsis transgenic plants carrying *ProCalS7:NLS-3×Venus* (*ProCalS7:Venus*) and *ProPDLP6:NLS-3×mApple* (*ProPDLP6:mApple*). Scale bar = 50 µm.

We next determined the physical interaction between PDLP6 and SUS6 using a co-immunoprecipitation (co-IP) assay. PDLP6-YFP and SUS6-HF were transiently expressed in *N. benthamiana*. As shown in Figure 5B, PDLP6-YFP pulls down SUS6-HF, while a free sfGFP does not. The findings showed that PDLP6 physically interacts with SUS6 *in planta*. To elucidate the role of SUS6 in regulating the plasmodesmal function, we introduced *sus6* mutation into *PDLP6-HF* via a genetic cross. We first generated a cross between the *sus56* double mutant (Bieniawska et al., 2007; Baroja-Fernández et al., 2012) and *PDLP6-HF*. F_2_ segregating progenies were screened for PDLP6 overexpressor in SUS5 and SUS6 wild-type background (hereafter *PDLP6-HF SUS6*) as well as *sus6* mutant background (hereafter *PDLP6-HF sus6*). A PCR genotyping assay was applied to identify the genotypes of SUS5 and SUS6 (Supplemental Figure 10A). The expression level of PDLP6-HF was determined using immunoblot analysis. The selected lines expressed a comparable level of PDLP6-HF (Supplemental Figure 10B). *PDLP6-HF SUS6* exhibits a similar stunted growth phenotype as *PDLP6-HF*, while *sus6* largely suppresses the plant growth phenotype of *PDLP6-HF* (Figure 5C). In addition, *sus6* also suppresses starch hyperaccumulation in mature leaves of *PDLP6-HF* (Figure 5C). Similar phenotypes were observed for two independent progenies of the crosses. Independent progenies were shown in fig. S10C. The findings suggest that SUS6 is required for PDLP6’s function in the vasculature.

### PDLP6 functions together with CalS7

Callose biosynthesis is catalyzed by callose synthases CalSs (De Storme and Geelen, 2014). The Arabidopsis genome encodes twelve members of CalSs, sequentially named CalS1-12 (Verma and Hong, 2001). *CalS7* was previously reported to be expressed in phloem (Barratt et al., 2011; Xie et al., 2011). Recent findings reported that CalS7 is specifically expressed in SEs (Kalmbach et al., 2022). CalS7 is required for callose deposition in sieve plates and sieve pores formation (Barratt et al., 2011; Xie et al., 2011). We thus hypothesized that PDLP6-SUS6-CalS7 might form a callose synthase complex in phloem, likely in SE. We tested the physical interactions among the three proteins. As the cytoplasmic loop (CL) of CalSs is predicted to interact with SUS6 (De Storme and Geelen, 2014; Verma and Hong, 2001), we cloned the CL of CalS7-HF (hereafter CalS7_CL_-HF). Using a coIP assay, we observed the physical interaction between SUS6 and CalS7_CL_ as well as PDLP6 and CalS7_CL_ (Figure 5B). To further characterize the functions of CalS7, we examined the genetic interaction between *PDLP6* and *CalS7*. Using the same strategy mentioned above, we isolated *PDLP6-HF cals7* and *PDLP6-HF CalS7* (Supplemental Figure 10). *cals7* mutant largely suppresses plant growth and starch accumulation phenotypes of *PDLP6-HF*, suggesting the genetic interaction between the two genes (Figure 5D).

According to single-cell transcriptomes of Arabidopsis vascular tissues, *PDLP6* is expressed specifically in PP (Kim et al., 2021). To determine vascular cell type-specific expression of PDLP6-YFP, we expressed the fusion proteins in transgenic plants expressing different phloem cell markers: phloem parenchyma (PP) marker (*ProSWEET13:SWEET13-mCherry*) (Kim et al., 2021), companion cell (CC) marker (*ProSUC2:PP2A1-mCherry*) and sieve elements marker (*ProSEOR2:SEOR2-mCherry*) (Cayla et al., 2015). T_3_ populations of transgenic lines expressing both PDLP6-YFP and the markers were isolated and subjected to confocal microscopy analysis. As shown in Supplemental Figure 11, PDLP6-YFP does not express in the same cell type as the PP, CC, or SE marker. It is noted that the SEOR2 promoter is active in immature SEs in leaves (Cayla et al., 2015). Also, SEOR1-YFP can move from protophloem sieve element (PSE) into phloem pole pericycle cells (PPP) in roots (Ross-Elliott et al., 2017). It is highly possible that the signal of SEOR2-mCherry driven by *ProSEOR2* was also mainly present in PPP in roots. As an alternative approach, we examined the promoter activity of *PDLP6* (*ProPDLP6*) and *CalS7* (*ProCalS7*) in Arabidopsis. A recent report demonstrated that CalS7 is specifically expressed in SEs (Verma and Hong, 2001). We thus generated Arabidopsis transgenic plants coexpressing *ProPDLP6:NLS-3*×*mApple* and *ProCalS7:NLS-3*×*Venus*. T_2_ populations of the transgenic plants were screened for the presence of mApple and Venus using confocal microscopy. As shown in Figure 5E, mApple and Venus signals were detected in the same root cell type, suggesting that *PDLP6* and *CalS7* promoters are active in the same cell type, likely in SEs. Altogether, our findings reveal the function of cell type-specific callose synthase complexes, which regulate the plasmodesmal function at specific cell interfaces.

## Discussion

### Gain of function study leads to the identification of cell type-specific functions of PDLPs

To avoid a gene redundancy issue, we employed a gain of function approach to characterize the function of PDLP5 and PDLP6. Despite the proteins being ubiquitously expressed in most cell types in the transgenic plants overexpressing PDLP-YFP fusion proteins (likely the same for PDLP-HF fusion proteins) (Supplemental Figure 7), the expression of PDLP5-HF and PDLP6-HF blocks the movement of sugar at specific cell interfaces (Figure 2). The findings suggest that functional partners required for the PDLPs are expressed in specific cell types. The assumption was supported by the finding that PDLP6 significantly enriches SUS6, which is expressed specifically in SE (Yao et al., 2020), in our PL assay (Figure 4D). We further demonstrated the physical and genetic interactions between PDLP6 and SUS6 (Figure 5, B and C), suggesting that PDLP6 functions together with SUS6 in regulating callose accumulation at PD in the vasculature. From our PL assay, we didn’t identify CalS7 as a partner of PDLP6. Given the SE-specific expression of CalS7 (Kalmbach et al., 2022), we investigated the roles of CalS7 in PDLP6-dependent PD regulation. Our data showed that CalS7 genetically interacts with PDLP6 and physically interacts with SUS6 as well as PDLP6 (Figure 5, B and D). Together, PDLP6-SUS6-CalS7 might form a callose biosynthesis complex in SE. We hypothesize that PDLP6 functions as a scaffold protein, recruiting SUS6 and CalS7 to PD for depositing callose at PD. The hypothesis is supported by the findings that co-transient expression of SUS6 and PDLP6 significantly increases the PD association of SUS6-YFP (Supplemental Figure 9). Arabidopsis genome encodes at least 8 PDLPs, 6 SUSs, and 12 CalSs, providing many possible combinations of callose biosynthesis complexes to function at different cell interfaces. As we demonstrated here, the identification of cell type-specific expression of the proteins involved in callose biosynthesis could provide a road map to define cell type-specific callose synthase complexes in plants.

In Arabidopsis, two different metabolic pathways have been proposed for generating UDPG, which is used as a glucose donor for callose and cellulose biosynthesis (Barnes and Anderson, 2018). SUS can directly convert sucrose to UDPG and fructose. The pathway is energetically more economical. Although SUSs might not be crucial for UDPG production during cellulose biosynthesis, *sus5;6* double mutant exhibits a lower level of callose in SEs (Barratt et al., 2009). Together with the SE-specific expression of SUS5 and SUS6, the proteins might be crucial for UDPG production in SE for callose biosynthesis. Cell type-specific expression of SUS showed that none of the SUSs is ubiquitously expressed in Arabidopsis (Yao et al., 2020). Consequently, a SUS-dependent UDPG production pathway might not be a dominant metabolic pathway in most cell types.

Alternatively, UDPG can be generated through a cytosolic convertase (CINV)-dependent pathway (Barratt et al., 2009; Barnes and Anderson, 2018). In addition to CINV, hexokinase (HXK), phosphoglucoisomerase (PGM), and UDP-glucose pyrophosphorylase (UGP) are required to generate UDPG from sucrose (Kleczkowski et al., 2004). During cellulose biosynthesis, a CINV-dependent pathway is considered to be the major metabolic pathway in generating UDPG (Barnes and Anderson, 2018). In this pathway, UGP converts Glucose-1-Phosphate (Glc-1-P) into UDPG. Further research is needed to understand whether the CINV pathway is crucial for a PDLP5-containing callose synthase complex in ECs and MCs.

### Cell type-specific expression and function of PDLPs

The single-cell transcriptome of Arabidopsis leaf phloem cells detects different members of PDLP transcripts in different cell types: *PDLP3* is detected in MCs, *PDLP6* is detected in PPs, *PDLP7* is detected in guard cells and PPs, and *PDLP8* is detected in guard cells and CCs (Kim et al., 2021). The findings imply the cell type-specific functions of PDLPs. In this study, we demonstrated the cell type-specific expression and function of PDLP5 and PDLP6. We detected PDLP5-YFP mainly in ECs in leaves and epidermis and cortex in roots (Figure 3D). From starch and callose hyperaccumulation phenotypes (Figure 2 and Figure 3, A-C), it is evident that PDLP5 functions in MCs when the protein is overexpressed. The absence of detectable PDLP5-YFP in MCs might be due to the lower expression level of PDLP5-YFP. As GUS was detected in MCs of *pPDLP5-GUS* transgenic plants (Supplemental Figure 6), the promoter of PDLP5 is likely active in MCs. It is also possible that PDLP5 is mostly expressed in leaf ECs, whereas the functional partners of PDLP5 are present both in the ECs and MCs. Thus, the overexpression of PDLP5 in MCs results in the overaccumulation of callose at PD between the MCs (Figure 3, A and B). A recent report showed that PDLP5-GFP expresses in different root cell types during different stages of lateral root primordium development (Barnes and Anderson, 2018). PDLP5-GFP was specifically detected in endodermis during the early stage of lateral root primordium (LRP) development. In the later stage, the fusion protein was also detected in the cortex and epidermis (Sager et al., 2020). The findings suggest the cell type-specific expression and function of PDLP5 during different developmental stages in regulating plant growth and development.

*PDLP6* transcripts were specifically detected in PP (Kim et al., 2021). In this study, we did not observe the expression of PDLP6-YFP in Arabidopsis transgenic plants expressing the *ProSWEET13:SWEET13-mCherry* PP marker (Supplemental Figure 11). As PDLP6 functions together with SUS6 and CalS7, which are specifically expressed in SE, we postulated that PDLP6 protein is expressed in SEs. The assumption was supported by the findings that the promoters of *PDLP6* and *CalS7* are active in the same cells in the root elongation zone (Figure 5E). It’s also noted that the expression of PDLP6-YFP is difficult to detect in leaves when the fusion protein was expressed under a *PDLP6* native promoter. It is possible that PDLP6 expresses in one of the subpopulations of PP or transiently in precursor SEs to regulate the function of PD.

It has been well-established that morphologies of plasmodesmata connecting different cell types vary depending on the functions of the cell (Faulkner, 2018; Lee and Frank, 2018). For example, simple PD, pore PD, or funnel PD can be found in protophloem sieve elements interfacing with different CC or phloem pole pericycle cells (Ross-Elliott et al., 2017). Although it is unknown whether PDLPs are involved in the formation or maintenance of different types of PD, it is tantalizing to hypothesize that different members of PDLPs play specific roles in recruiting cell interface-specific callose synthase complexes to regulate the PD structure and function. Investigation of PD ultrastructure in PDLP overexpressors and *pdlp* knock-out mutants (including *pdlp* higher-order mutants) using electron tomography as recently described (Nicolas et al., 2022) will uncover the functional roles of PDLPs in regulating the PD structure.

### Functional roles of PD in sugar movement between cells in mature leaves

In Arabidopsis, chloroplast-rich mesophyll cells are mainly responsible for photosynthesis in mature leaves. Sugars produced via photosynthesis need to load into phloem for long-distance sugar transport. PD have been implicated to play important roles in allowing sugars to move symplasmically from MCs to phloem during sugar loading (Zhang and Turgeon, 2018; Braun, 2022). Alternatively, it is plausible that sugars are exported out of MCs and move to phloem through the apoplasm (Giaquinta et al., 1983; Ayre, 2011). Here, our findings further support that the symplasmic movement of sugars from MCs to PPs is an important cellular route during sugar movement and loading in mature leaves. The overexpression of PDLP5 results in the overaccumulation of callose between MCs, blocking the movement of sugars out of MCs and resulting in starch hyperaccumulation (Figure 2). A different line of evidence was provided from the characterization of *cher1-4*. The mutant has much less PD between mesophyll cells and other cell types in leaves, resulting in starch hyperaccumulation mostly in MCs. In addition, *CHER1* is epistatic to *PDLP6*, *SWEET11*, and *SWEET12* (Supplemental Figure 4). Together, these findings support that the symplasmic movement of sugars along the cellular route from MCs to PPs in mature leaves is the major contributor to sugar translocation in mature leaves.

### A PL assay is a promising methodology for uncovering novel molecular players functioning at PD

We utilized a recently developed PL approach (Branon et al., 2018) to identify functional partners of PDLP5 and PDLP6, which are targeted to PD-PM. We included MCTP3 as a control for the PDLPs as the protein is targeted to PD-ER (Brault et al., 2019). As shown in Figure 4D, we were able to identify many proteins enriched specifically by PDLP5, PDLP6, or MCTP3. The identification of candidate proteins with potential functions at PD greatly boosts confidence in confirming and determining novel functions of other candidate proteins with no clear functions at PD (Supplemental Data set 1). We identified several putative functional partners of PDLP5 and PDLP6, including leucine-rich repeats receptor-like kinases and cysteine-rich receptor-like kinases (Supplemental Data set 1). It’s tantalizing to hypothesize that the PD-localized receptor-like kinases regulate the activity of callose synthase complexes through phosphorylation. We also identified a small set of proteins specifically enriched by PDLP5 or PDLP6 (Figure 4D). Here, we report SUS6 as a cell/tissue-specific functional partner of PDLP6. Despite using a *UBQ10* promoter, which expresses the PDLPs ubiquitously, we identify cell type-specific functional partners of PDLP6. We also observed PDLP5 significantly enriches several members of aquaporins (PIPs). Recently findings showed that PIP2;1 and PIP2;4 can transport reactive oxygen species, including hydrogen peroxide (H_2_O_2_) (Rodrigues et al., 2017; Wang et al., 2020). Given the functional roles of PDLP5 and H_2_O_2_ in plant immunity, functional characterization of the aquaporins has the potential in revealing the novel functions of PIPs in regulating PD during microbial defense. Together, our findings suggest that we were able to identify cell/tissue type-specific functional partners of PDLP5 and PDLP6 despite expressing the fusion proteins with a constitutive promoter. It’s highly plausible that the use of native promoter in expressing TbID fusion protein might greatly enrich cell/tissue type-specific functional partners; however, the strategy also likely misses the detection of candidate proteins due to the lower expression level of TbID fusion proteins. A combination of a constitutive promoter and a native promoter would likely facilitate the identification of functional partners of proteins of interest. Our findings demonstrated that a PL assay is a powerful tool to study the plasmodesmal biology.

## Materials and Methods

### Plant Material, Growth Conditions, Transformation, and Plant Selection

*Arabidopsis thaliana* (Arabidopsis) and *Nicotiana benthamiana* plants were grown at 22°C with 50% humidity and irradiated with 110 µmol m^-2^ s^-1^ white light for 16 h per day. To grow plants under high light intensity, plants were irradiated with 200 µmol m^-2^ s^-1^ white light. Arabidopsis T-DNA insertion mutants, *cals7* (SALK_048921), *cher1-4* (SALK_065853), *pdlp5* (SAIL_46_E06.v1), and *sweet11;12* (CS68845) were obtained from ABRC (Columbus, OH). *sus56* is a gift from Dr. Alison M Smith’s lab. *pdlp6-1* and *pdlp6-2* were generated using CRISPR/Cas9 technology in this study. Transgenic Arabidopsis plants were generated using the simplified transformation method (https://plantpath.wisc.edu/simplified-arabidopsis-transformation-protocol/). To select transgenic plants harboring transgenes containing the hygromycin-resistance gene, T_1_ seeds were germinated on a ½ × LS medium containing 25 µg/mL hygromycin. For those harboring transgenes containing the glufosinate-resistance gene, T_1_ seeds were germinated on soil and sprayed with 0.1% (v/v) Finale Herbicide (Bayer) and 0.05% (v/v) Silwet L-77 (PhytoTech) around 10 days after germination. The T_2_ or T_3_ plants were selected on a ½ × LS medium containing 10 µg/mL glufosinate-ammonium (MilliporeSigma).

### Gene Cloning and Plasmid Construction

If not specified otherwise, constructs generated in this work used a standard Gateway cloning system (Invitrogen). Coding sequences, promoters, or genomic DNA fragments were amplified with Gateway-compatible primers from cDNA synthesized from total RNA extracted from wild-type Col-0 seedlings or total genomic DNA extracted from Col-0 using Phusion High-Fidelity DNA polymerase (ThermoFisher). PCR fragments were first cloned into the pDONR 207 entry vector and subsequently cloned into different destination vectors. CRISPR-P 2.0 (Liu et al., 2017) was used to design the guide sequence for PDLP6 (http://crispr.hzau.edu.cn/CRISPR2/). All primers and vectors used for cloning are listed in Table S1 and S2. In this study, we reported the following constructs: *Pro35S:PDLP5-HF*, *Pro35S:PDLP6-HF*, *Pro35S:HF-YFP*, *Pro35S:PDLP5-YFP*, *Pro35S:PDLP6-YFP*, *ProPDLP5:PDLP5-YFP*, *ProPDLP6:PDLP6-YFP*, *pKIR-CRISPR-PDLP6*, *ProSWEET13:SWEET13-mCherry*, *ProSUC2:PP2A1-mCherry*, *ProSEOR2:SEOR2-mCherry*, *ProUBQ10:PDLP5-TbID-3xFlag*, *ProUBQ10:PDLP6-TbID-3xFlag*, *ProUBQ10: TbID-3xFlag-MCTP3*, *ProCalS7:NLS3-3xVenus*, *ProPDLP6:NLS-3xmApple*, *ProPDLP5:GUS*, and *ProPDLP6:GUS*.

### Whole Tissue Starch Staining

The aerial portion of 3-4-week-old Arabidopsis plants grown under regular light were harvested at the end of the night. For high light-treated plants, 4-week-old Arabidopsis plants grown under a regular light intensity (110 µmol m^-2^ s^-1^) were treated with higher light intensity (200 µmol m^-2^ s^-1^) for 7-10 days. The inflorescence was removed and the rosette leaves were decolored with 95% ethanol. The samples were then washed with ddH_2_O and stained with Lugol’s iodine solution (Sigma-Aldrich) for 5 minutes and rinsed with ddH_2_O. The images were captured with a Canon camera 1-2 hours after destaining.

### Tissue Sectioning and Starch Staining

Leaf punch samples were collected and fixed in FAA fixative (5% formaldehyde, 5% glacial acetic acid, 50% ethyl alcohol). Samples were dehydrated through graded ethanol series (70, 85, 95, and 100%) for 3-6 hours for each concentration. Samples were infiltrated into LR White hard grade resin (Electron Microscopy Sciences) and polymerized at 55°C for 48 hours. Sections were made using a Leica UC6 ultramicrotome at 1.5 µm thickness. Sections were stained for non-soluble polysaccharides as the following: slides with sections were immersed in periodic acid for 5 minutes, rinsed in distilled water for 5 minutes, stained in Schiff’s reagent (Electron Microscopy Sciences) for 10 minutes, rinsed in running tap water for 5 minutes, and air-dried. Dry slides were coverslipped using Permount mounting media (Fisher Scientific). Images were captured using AxioImager A2.

### Transmission electron microscopy (TEM)

Leaves were dissected using a leaf punch and samples were fixed with 3% glutaraldehyde (w/v) and 1% paraformaldehyde (w/v) in 0.1 M cacodylate at 4°C. Samples were rinsed 3 times in 0.1 M cacodylate buffer at room temperature. The samples were post-fixed in 1% osmium tetroxide in 0.1 M cacodylate for 1 hour, followed by a 5-minute wash in dH_2_O and en-bloc staining with 2% uranyl acetate for 1 hour. The samples were then dehydrated in a graded ethanol series, cleared with ultra-pure acetone, and infiltrated/embedded using SPURR’s epoxy resin (Electron Microscopy Sciences). Resin blocks were polymerized for 48 hours at 70°C. Thick and ultrathin sections were made using a Leica UC6 ultramicrotome. Ultrathin sections (60-70 nm) were collected onto copper grids and images were captured using a JEOL JSM2100 scanning and transmission electron microscope.

### Anthocyanin extraction and quantification

Fresh tissues were frozen with liquid nitrogen and homogenized with 1600 miniG (SPEX). 1 ml of extraction buffer (45% methanol and 5% acetic acid) was added to the homogenized tissues. The samples were incubated at 4°C for 30 min and centrifuged at 14,000 g for 5 min at room temperature. The supernatant was transferred into a new 1.5 ml tube. 200 μl of the supernatant was pipetted to 96-well plates and the absorbances at 530 nm and 657 nm were determined using plate reader (SpectraMax iD3). Anthocyanin content was calculated by [Abs_530−_ (0.25 × Abs_657_)/g fresh weight.

### Starch and soluble sugars extraction and quantification

The whole rosettes from 5-week-old Arabidopsis plants treated with high light for a week were collected at the end of the night and immediately frozen in liquid nitrogen before being stored at −80°C until quantification. Soluble sugar and starch were extracted according to Leach & Braun (2016). A methanol:chloroform:water (MCW)-based extraction was performed to isolate soluble sugars. Samples were grinded in liquid nitrogen and 100 mg of powder was transferred into 1.5 ml EP tube. 1 ml of MCW extraction buffer was added. The samples were vortexed briefly, incubated in 50°C water bath for 30 min, and centrifuged at 14,000 *g* for 5 min at room temperature. The supernatant was transferred into 15 ml conical tube and stored on ice. Repeat the extraction process twice and the supernatant was combined in the 15 ml conical tube. The pellet was saved for starch determination. 0.6 volumes of water were added to the 15 ml conical tube. The samples were vortexed and centrifuged at 4,650 *g* for 5 min at room temperature. Aqueous (top) phase containing the soluble sugars was pipetted into a 1.5 ml EP tube and stored at −20°C until measurement. Sucrose concentration was determined using Megazyme kit (catalog No. K-SUFRG) following manufacturer instructions. Starch was solubilized from the MCW-extracted tissue pellet and enzymatically degraded into glucose for quantification. The pellet was resuspended in 330 μl of DMSO and heated at 100°C for 5 min to solubilize and gelatinize the starch. 50 μl of the slurry was diluted by 950 μl of 100 mM sodium acetate (pH 5.0). α-amylase (30 U) was added and the samples were vortexed briefly, heated at 100°C for 15 min, and the cooled at 50°C for 3 min. Amyloglucosidase (66 U) was added and the samples were vortexed briefly, incubated at 50°C for 1 hour, and centrifuged at 14,000 *g* for 5 min at room temperature. The supernatant containing starch-derived glucose was transferred into a new 1.5 ml EP tube. Starch concentration was determined using Megazyme kit (catalog No. K-TSTA-50A) following manufacturer instructions.

### Aniline Blue Staining

Mature leaves from 4-week-old Arabidopsis plants were infiltrated with 0.1 mg/ml aniline blue in 1x PBS buffer (pH 7.4). Ten-day-old seedlings were vacuum-infiltrated with 0.1 mg/ml of aniline blue in 1x PBS buffer (pH 7.4) to detect callose accumulation in roots. The tissues were stained for ∼ 5 min prior to imaging. Stained leaf and root tissues were imaged using Zeiss LSM 700 Laser Scanning Confocal System. Callose in mature leaves was quantified using FIJI. For epidermis, images were converted from lsm to tiff. 8-bit images were used for analysis. Black and white images highlighting callose were created by Auto Threshold which was set by RenyiEntropy white method. A Particle Analysis tool was used to outline callose with sizes from 0.10 to 20 µm2 and circularity from 0.15 to 1.00. Quantitative numerical values in µm2 were then exported. For mesophyll cells, an aniline blue-stained callose area was manually selected and the signal intensity was determined by measuring integrated density. Semi-quantitative evaluation of the relative level of aniline blue-stained callose in the root tissues of the transgenic plants was performed using FIJI. Images were converted from lsm to tiff. 16-bit images were used for analysis. Horizontal lines were drawn near the bottom of the images crossing all root cell types as shown in Fig. 2d. A plot profile was used to generate a two-dimensional graph. Value on the y-axis represents the relative signal intensity as an arbitrary unit (AU).

### GUS Activity Staining

GUS staining was performed as previously described (Li et al., 2016) with minor modifications. Mature leaves and small seedlings were immersed in GUS solution (100 mM sodium phosphate buffer [pH 7.0], 10 mM Na_2_EDTA, 1 mM K_3_ [Fe(CN)_6_], 1 mM K_4_ [Fe(CN)_6_], 0.1% Triton X-100, and 1 mM X-Gluc). Samples were vacuumed for 10-40 min, followed by incubation in darkness at 37°C for 2-16 h. After staining, samples were de-stained in 75% ethanol. For imaging transverse leaf sections, GUS-stained leaves were embedded in 3% agarose and sectioned with a vibrating blade microtome (Leica VT1000 S). Images were taken using ZEISS Axio Observer.

### Confocal Imaging

All confocal images were captured with a confocal laser-scanning microscope (Zeiss LSM 700). A small piece of tissue was mounted with water on a glass slide. For leaf tissues, the abaxial side was imaged. YFP, sfGFP, and Venus were excited at 488 nm and emission was collected over 510–550 nm using SP555. Aniline blue-stained callose was excited at 405 nm and emission was collected over 420–480 nm using SP 490. mApple was excited at 555 nm and emission was collected over 590-630 nm using SP 640. CF was excited at 488 nm and emission was collected over 505-545 nm using SP 555. PI was excited at 555 nm and emission was collected using SP 640.

### Immunoblot Analyses

Arabidopsis leaves were frozen with liquid nitrogen and homogenized with 1600 miniG (SPEX). Protein extraction buffer (60 mM Tris-HCl [pH 8.8], 2% [v/v] glycerol, 0.13 mM EDTA [pH 8.0], and 1× protease inhibitor cocktail complete from Roche) was added to the homogenized tissues (100 µl/10 mg). The samples were vortexed for 30 s, heated at 70°C for 10 min, and centrifuged at 13,000 *g* for 5 min at room temperature. The supernatants were then transferred to new tubes. For SDS-PAGE analysis, 10 µl of the extract in 1x Laemmli sample buffer (Bio-Rad) was separated on 4–15% Mini-PROTEAN TGX precast protein gel (Bio-Rad). The separated proteins were transferred to a polyvinylidene fluoride membrane (Bio-Rad) using a Trans-Blot Turbo Transfer System RTA transfer kit following the manufacturer’s instructions (Bio-Rad). The membrane was incubated in a blocking buffer (3% [v/v] BSA, 50 mM Tris base, 150 mM NaCl, 0.05% [v/v] Tween 20 [pH 8.0]) at room temperature for 1 h, then incubated overnight with a 1:10,000 dilution of an ⍺-GFP (abcam catalog No. ab290), ⍺-Flag-HRP (Sigma-Aldrich catalog No. A8592) or ⍺-Streptavidin-HRP antibody (abcam catalog No. ab191338) at 4°C. The membrane was washed four times with 1× TBST (50 mM Tris base, 150 mM NaCl, 0.05% [v/v] Tween 20 [pH 8.0]) for 10 min. For an ⍺-GFP, 1:20,000 goat anti-rabbit IgG (Thermo Fisher Scientific catalog No. 31,460) was used as a secondary antibody. The membrane was washed four times with 1× TBST (50 mM Tris base, 150 mM NaCl, 0.05% [v/v] Tween 20 [pH 8.0]) for 10 min. The signals were visualized with SuperSignal West Dura Extended Duration Substrate (Pierce Biotechnology).

### Transient overexpression for subcellular localization and Co-Immunoprecipitation (CoIP)

*Agrobacterium tumefaciens* GV3101 harboring the plasmid of interest was adjusted to an optical density of A_600_ 0.1 using sterilized ddH_2_O and infiltrated into 6-week-old *Nicotiana benthamiana* leaves. Infiltrated tissues were subjected to live-cell imaging or CoIP assay 2 d after infiltration. CoIP assay was performed as previously described (Aung et al., 2020). One gram fresh weight of tissues were ground in liquid nitrogen and lysed with 3 ml of RIPA buffer (50 mM Tris-HCl pH 7.5, 150 mM NaCl, 1% NP-40, 1% Sodium deoxycholate, and 0.1% SDS with 1x complete protease inhibitor cocktail (Roche) on a rotator at 4°C for 1 h. The samples were centrifuged at 13,000 *g* for 10 mins, filtered with two layers of Miracloth and centrifuged again to remove cell debris. 60 μl of the supernatants were served as input controls. 20 μl of 4 x SDS loading buffer (Bio-rad) was added into input samples and heated at 95°C for 5 min. 15 μl of GFP-Trap®_A (ChromoTek) was washed three times with RIPA buffer and added into the supernatant. The samples were mixed on a rotator at 4 °C for 1 h. The agarose beads were spun down at 100 g for 1 min and washed four times with RIPA buffer. Proteins co-immunoprecipitated with the YFP-fusion protein were eluted by adding 60 μl of 1x SDS loading buffer and heating at 95°C for 5 min. The protein samples were analyzed by immunoblot assay.

### Statistical Analysis

Column plots were created using GraphPad Prism. Box plots were created with an online software (https://huygens.science.uva.nl/PlotsOfData/). Mann-Whitney U Test (www.socscistatistics.com/tests/mannwhitney/default2.aspx) or Student’s *T*-Test (https://www.socscistatistics.com/tests/studentttest/default2.aspx) were performed for testing statistical significance of differences.

### Proximity labeling (PL)

The PL assay was performed according to Mair et al., 2019 with minor modifications. Three independent transgenic events from each transgenic line were subjected to biotin labeling and streptavidin enrichment. 14-day-old seedlings were carefully removed from ½ LS medium, transferred into 40 ml of a 50 μM biotin solution and incubated at room temperature for 3 hr. The biotin solution was then removed and seedlings were quickly rinsed with ice-cold water for three times. The samples were homogenized using a pestle and mortar in liquid nitrogen. Around 1.5 ml of leaf powder was resuspended with 2 ml of RIPA lysis buffer (50 mM Tris-HCl pH 7.5, 150 mM NaCl, 1% NP-40, 1% Sodium deoxycholate, and 0.1% SDS with 1x complete protease inhibitor cocktail (Roche)). The samples were vortexed to mix and incubate at 4°C on a rotator for 10 min. To digest cell walls and DNA/RNA, the samples were incubated with 0.5 μl of Lysonase (Millipore) on a rotator at 4°C for 15 min. The samples were centrifuged at 15,000 *g* for 10 mins at 4°C. The clear supernatant was applied to a PD-10 desalting column to remove excess free biotin using gravity following the manufacturer’s instructions. 2.5 ml of the protein extract was loaded onto the desalting column. Proteins were then eluted with 3.5 ml of equilibration buffer (RIPA buffer without 1x complete protease inhibitor cocktail). The desalted protein extracts were quantified by Bradford assay and a complete protease inhibitor cocktail was added to each sample to reach final concentrations of 1x complete.

### Affinity purification of biotinylated proteins

To enrich biotinylated proteins, 150 µl streptavidin-coated beads (Dynabeads MyOne Streptavidin C1 beads) pre-washed with extraction buffer were mixed with protein extracts and incubated on a rotator at 4°C overnight. The Streptavidin beads were then washed with 1 ml of the following solutions for 2 min each: 2 × cold extraction buffer, 1 × cold 1 M KCl, 1 × cold 100 mM Na_2_CO_3_, 1 × 2M Urea in 10 mM Tris pH 7.5, 6 × 50 mM Tris 7.5. The beads were resuspended in 200 μl of 50 mM Tris 7.5 and stored at −80 °C for subsequent analysis until the next step.

### MS sample preparation

Biotinylated proteins on streptavidin beads eluted from beads by incubation at 95°C for 10 minutes in 1x S-Trap lysis buffer (5% SDS, 50mM TEAB, pH 8.5) supplemented with 12.5 mM biotin. Eluted samples were subjected to S-Trap sample processing technology (Catalog number C02-micro-80, ProtiFi, USA), following manufacturer protocol. Samples were reduced in 2 mM TCEP, alkylated in 50 mM iodoacetamide, and digested into peptides at 37°C in one round of overnight incubation with 1 µg of Trypsin and a second incubation of 4 hours with 0.1 µg trypsin plus 0.1 µg Lys-C. Peptides were further desalted using SepPack C18 columns (Waters). Tandem Mass Tag (TMT, Thermo Scientific) labeling was performed on purified peptides from each sample as previously reported (Song et al., 2020). TMT labeling reaction was stopped using 5% hydroxylamine and the quenched samples were then pooled. Pooled samples were subjected to high pH fractionation using Pierce High pH Reversed-Phase Peptide Fractionation Kit (Thermo Scientific) following manufacturer instructions. The obtained 8 fractions were further concatenated (pooled) as follow: fraction 1 with fraction 5, fraction 2 with fraction 6, fraction 3 with fraction 7, and fraction 4 with 8. Samples were dried in a SpeedVac and resuspended in 0.1% TFA in Optima™ grade H_2_O (Fisher).

### LC-MS/MS

Chromatography was performed on a Thermo UltiMate 3000 UHPLC RSLCnano. Peptides were desalted and concentrated on a PepMap100 trap column (300 µM i.d. x 5 mm, 5 µm C18, 100 Å µ-Precolumn, Thermo Scientific) at a flow rate of 10 µL min-1. Sample separation was performed on a 200 cm Micro-Pillar Array Column (µ-PAC, Pharmafluidics) with a flow rate of ∼300 nL min-1 over a 150 min reverse phase gradient (80% ACN in 0.1% FA from 1% to 15% over 5 min, 15% to 20.8% over 20 min, from 20.8% to 43.8% over 80 min, from 43.8% to 99.0% in 11 min and kept at 99.0% for 5 min). Eluted peptides were analyzed using a Thermo Scientific Q-Exactive Plus high-resolution quadrupole Orbitrap mass spectrometer, which was directly coupled to the UHPLC. Data dependent acquisition was obtained using Xcalibur 4.0 software in positive ion mode with a spray voltage of 2.3 kV and a capillary temperature of 275 °C and an RF of 60. MS1 spectra were measured at a resolution of 70,000, an automatic gain control (AGC) of 3e6 with a maximum ion time of 100 ms and a mass range of 400-2000 m/z. Up to 15 MS2 were triggered at a resolution of 35,000. A fixed first mass of 115 m/z. An AGC of 1e5 with a maximum ion time of 50 ms, a normalized collision energy of 33, and an isolation window of 1.3 m/z were used. Charge exclusion was set to unassigned, 1, 5–8, and >8. MS1 that triggered MS2 scans were dynamically excluded for 25 s.

### Proteomics data analysis

Raw data were analyzed using MaxQuant version 2.1.0.0 (Cox and Mann, 2008). Spectra were searched, using the Andromeda search engine (Cox et al., 2011), against *Arabidopsis thaliana* TAIR10 annotation (www.arabidopsis.org). The proteome files were complemented with reverse decoy sequences and common contaminants by MaxQuant. Carbamidomethyl cysteine was set as a fixed modification while methionine oxidation and protein N-terminal acetylation were set as variable modifications. The sample type was set to “Reporter Ion MS2” with “TMT18plex” selected for both lysine and N-termini. TMT batch-specific correction factors were configured in the MaxQuant modifications tab (TMT Lot No. XA338617). Digestion parameters were set to “specific” and “Trypsin/P;LysC”. Up to two missed cleavages were allowed. A false discovery rate, calculated in MaxQuant using a target-decoy strategy (Elias and Gygi, 2007), less than 0.01 at both the peptide spectral match and protein identification level was required. The match between runs feature of MaxQuant was not utilized. Statistical analysis on the MaxQuant output was performed using the TMT-NEAT Analysis Pipeline (Clark et al., 2021). Proteins were called as significantly enriched interactors if they were under a false discovery rate cutoff of q-value < 0.1 and a log_2_FC > 0.2 or < −0.2. Principal component analysis (PCA) was performed and visualized using the ggbiplot R package as part of the TMT-NEAT analysis pipeline. Volcano plots were generated using VolcaNoseR.

## Acknowledgments

We thank Tracey Stewart from Roy J. Carver High Resolution Microscopy Facility at Iowa State University (ISU) for helping with histological sectioning, PAS staining, and TEM material preparation and imaging. We thank the Aung lab members Dr. Yani Chen, Haris Variz, Madelyn M Clements, Saiful Islam, Panchali Chakraborty, and a former member Dr. Kemesh Regmi (currently at Indiana University Bloomington) and Tyler Weide. We thank Dr. Yanhai Yin from ISU, Dr. Michelle Guo from ISU, Dr. Anne Rea from Michigan State University and Dr. Yu-Ti Cheng from Duke university for discussion and proofreading. We thank Dr. Alison Smith (John Innes Centre) for generously providing *sus56* seeds and Dr. Tetsuya Higashiyama (Nagoya University) for pKIR1.1 vector. We also thank the ABRC for providing the T-DNA insertion mutants and pMDC163 vector and Addgene for providing plasmid # 127366.

## Funding

This work was supported by the National Institute of General Medical Science (Grant R00GM115766 to K.A.), the ISU Crop Bioengineering Center seed grant (K.A.), the ISU Plant Science Institute (J.W.W.), and Hatch Act funds (Project No. IOW04308 to J.W.W.).

## Conflict of interest statement

The authors declare no conflict of interest regarding this study.

## Author Contributions

Z.L. designed and performed most of the experiments, including data analysis. S.L. designed and generated the vectors and plasmids. C.M., and J.W.W. designed, performed LC/MS-MS. C.M., Z.L., and K.A. analyzed the MS data. K.A. designed the experiments and analyzed the data. K.A., Z.L., and J.W.W. wrote the manuscript with inputs from all authors.

The author responsible for distribution of materials integral to the findings presented in this article in accordance with the policy described in the Instructions for Authors (https://academic.oup.com/plcell) is: Kyaw Aung

---

**Supplemental Figure 1.**
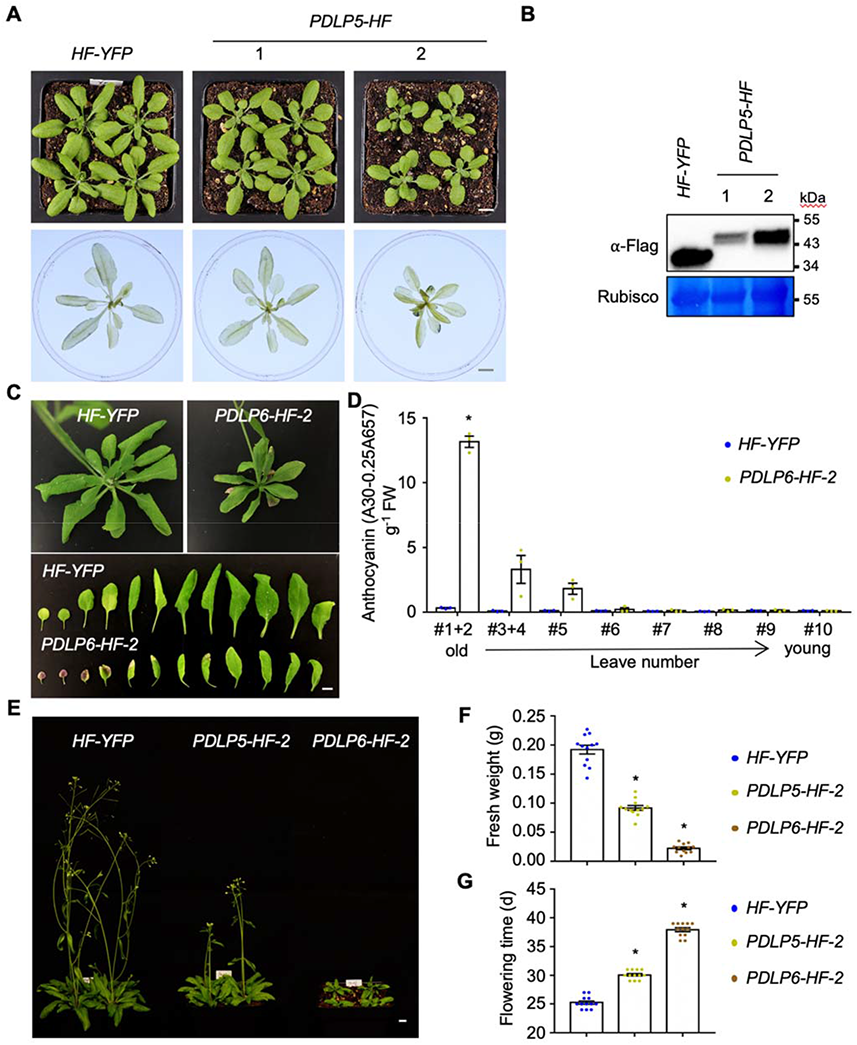
Growth phenotypes of *PDLP5-HF* and *PDLP6-HF*. (**A**) The overexpression of PDLP5 leads to stunted plant growth (top panel). *HF-YFP* serves as a control. Images were taken from 25-day-old plants growing under a standard light condition (110 µmol m^-2^ s^-1^) using the same magnification. Tissues were harvested at the end of the night and stained using Lugol’s iodine solution (lower panel). Scale bar = 1 cm. (**B**) Immunoblot analysis detects the expression of PDLP5-HF in the two transgenic lines. An anti-Flag antibody was used to detect the expression of Flag-fusion proteins. Rubisco serves as a loading control. (**C**) Older leaves of *PDLP6-HF* exhibit purple color. Scale bar = 1 cm. (**D**) Anthocyanin accumulation in leaves shown in c. Anthocyanins were extracted from different leaves (old to young) and measured at absorbance 530 nm and 657 nm using a microplate reader. The plot shows the mean with SEM (n = 3). Asterisks indicate differences that are statistically significant (*t*-Test; two-paired; *P* < 0.01). (**E**) Late-flowering phenotypes of *PDLP5-HF* and *PDLP6-HF*. *HF-YFP* serves as a control. Images were taken from 35-day-old plants using the same magnification. Scale bar = 1 cm. (**F**) Quantification of plant growth by fresh weight. The above-ground tissues of 25-day-old plants were quantified. The plot shows the mean with SEM (n = 12). Asterisks indicate differences that are statistically significant (*t*-Test; two-paired; *P* < 0.01). (**G**) Quantification of flowering time. Flowering time of *PDLP5-HF*, *PDLP6-HF*, and *HF-YFP* was measured as the number of days at bolting. The plot shows the mean with SEM (n = 12). Asterisks indicate differences that are statistically significant (*t*-Test; two-paired; *P* < 0.01).

**Supplemental Figure 2.**
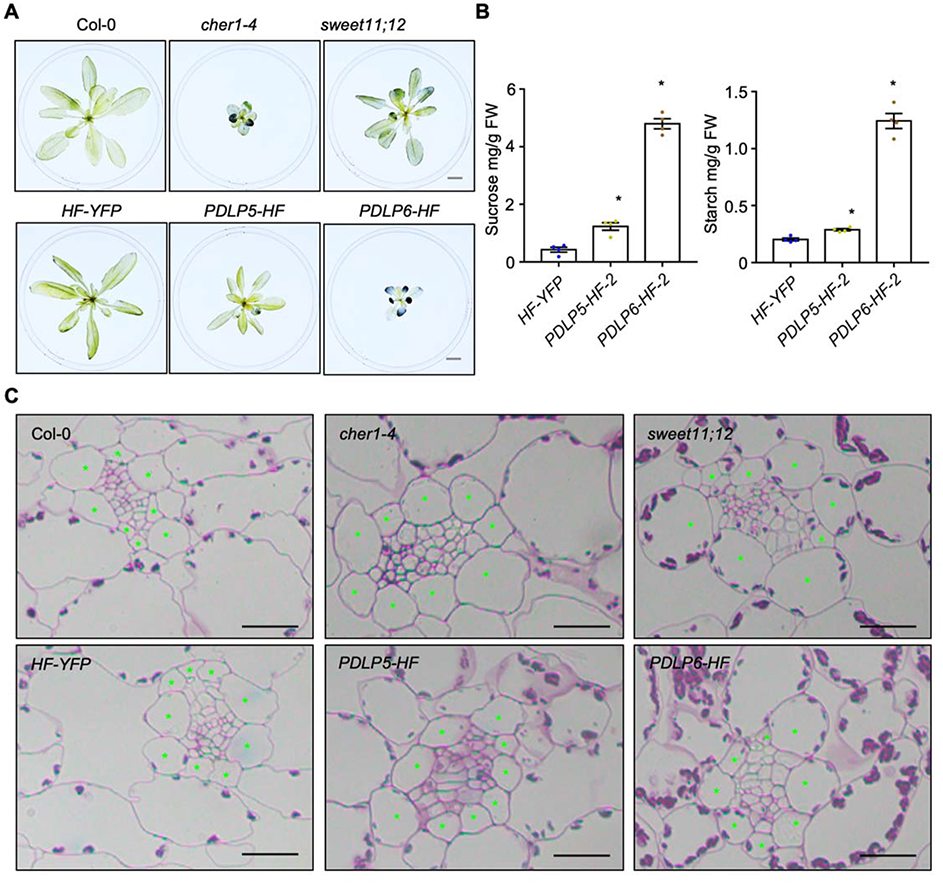
Starch accumulation in mature leaves of different Arabidopsis genotypes. (**A**) Starch staining of plants grown under a light intensity used for a standard Arabidopsis growth condition (110 µmol m^-2^ s^-1^) for four weeks. Samples were collected at the end of the night for starch staining using Lugol’s iodine solution. Images were taken using the same magnification. Scale bars = 1 cm. (**B**) Quantification of starch and sucrose contents in transgenic plants expressing *PDLP5-HF* or *PDLP6-HF*. *HF-YFP* serves as a control. The plots show the mean with SEM (n = 4). Asterisks indicate differences that are statistically significant (*t*-Test; two-paired; *P* < 0.01). (**C**) Histological sections of Arabidopsis leaves. Mature leaves of 5-week-old high light-treated plants were subjected to sectioning and staining with periodic acid/Schiff reagent, which stains polysaccharide in cell wall and starch grains in chloroplasts. Asterisks mark bundle sheath cells. Scale bars = 20 µm.

**Supplemental Figure 3.**
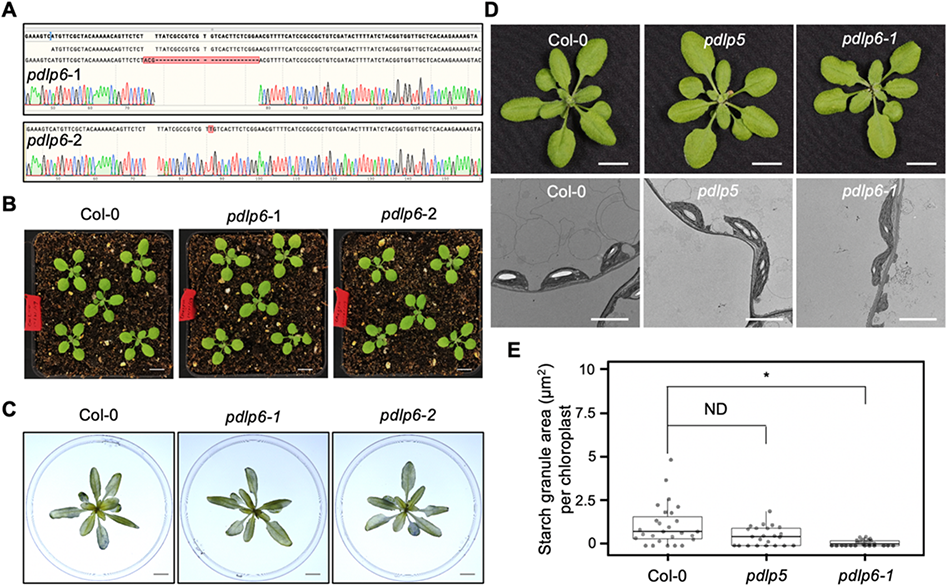
Characterization of *pdlp6* mutants. (**A**) Genotyping of *pdlp6-1* and *pdlp6-2* mutants generated by CRISPR/Cas9 technology. *pdlp6-1* and *pdlp6-2* carry 26 bp deletion and a G to T mutation around the gRNA target sites, respectively. (**B**) 2-week-old Arabidopsis plants were grown under a standard Arabidopsis growth condition. Images were taken using the same magnification. Scale bars = 1 cm. (**C**) Starch staining of *pdlp6* mutants. Tissues were harvested at the end of the night and stained using Lugol’s iodine solution. Images were taken using the same magnification. Scale bars = 1 cm. (**D**) Plant morphologies (top panel) and TEM images (lower panel) of chloroplasts from mesophyll cells of Col-0, *pdlp5,* and *pdlp6-1* mutants. Samples for TEM were collected from 5-week-old high light treated plants. Scale bars = 1 cm (top panel) or 5 μm (lower panel). (**E**) Quantification of starch granule areas per chloroplast. Each dot represents the total starch granule area in one chloroplast. Col-0, n = 27; *pdlp5*, n = 23; and *pdlp6-1*, n = 25. An asterisk indicates differences that are statistically significant (Mann-Whitney *U* Test; two-paired; *P* < 0.01). ND, no statistical difference.

**Supplemental Figure 4.**
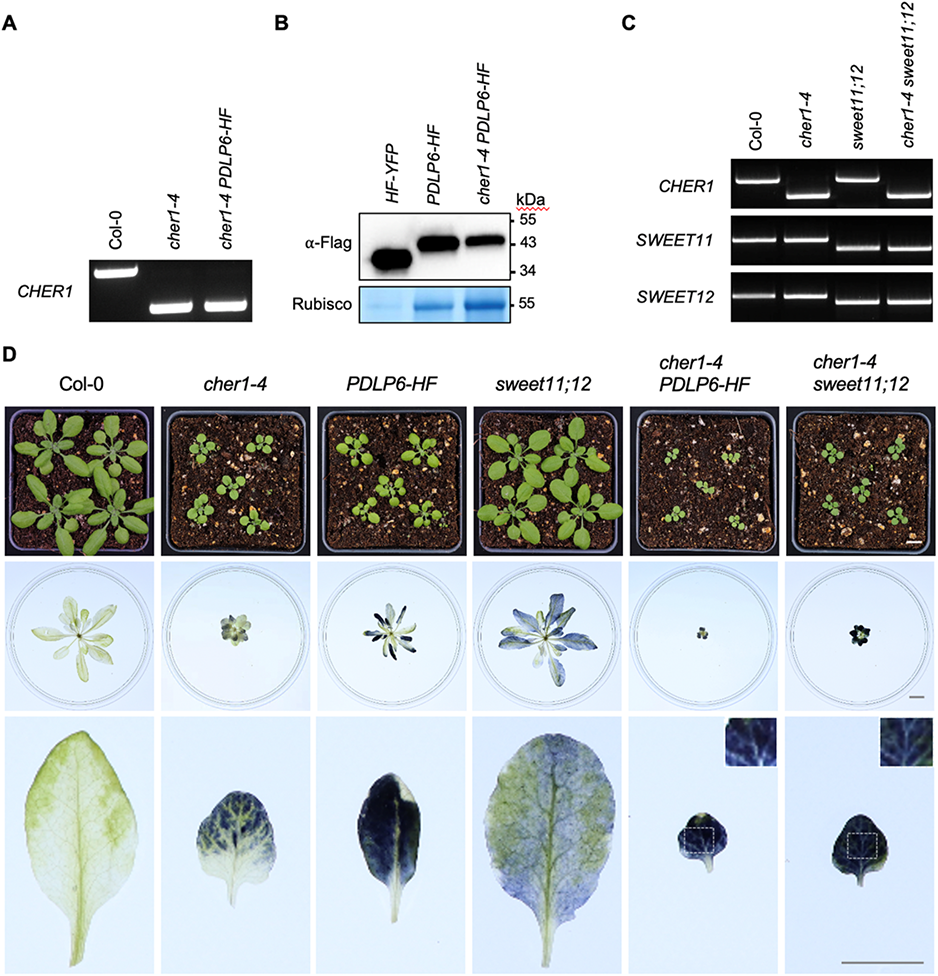
Genetic interactions between *CHER1*, *SWEET11;12*, and *PDLP6*. (**A**) A PCR Genotyping assay to isolate cher1-4. A pair of primers (LP and RP) was used to amplify genomic DNA and another pair of primers (SALK LBb1.3 and RP) was used to amplify the presence of t-DNA for each genotype. Upper bands indicate the PCR products of genomic amplifications. The lower bands indicate the presence of T-DNA insertions in both chromosomes (homozygous lines). (**B**) Immunoblot analysis detects the expression of PDLP6-HF. An anti-Flag antibody was used to detect the expression of Flag-fusion proteins. Rubisco serves as a loading control. (**C**) A PCR Genotyping assay to isolate sweet11, and sweet12 homozygous lines as mentioned above. (**D**) 3.5-week-old plants were grown under a light intensity used for a standard Arabidopsis growth condition (110 µmol m^-2^ s^-1^) (top panel). Tissues from 6-week-old plants were collected at the end of the night for starch staining using Lugol’s iodine solution (middle panel). Enlarged images of leaves highlight the different patterns of starch accumulation (lower panel). Images were taken using the same magnification. Scale bars = 1 cm. Inserts show enlarged parts of leaves to indicate the veins.

**Supplemental Figure 5.**
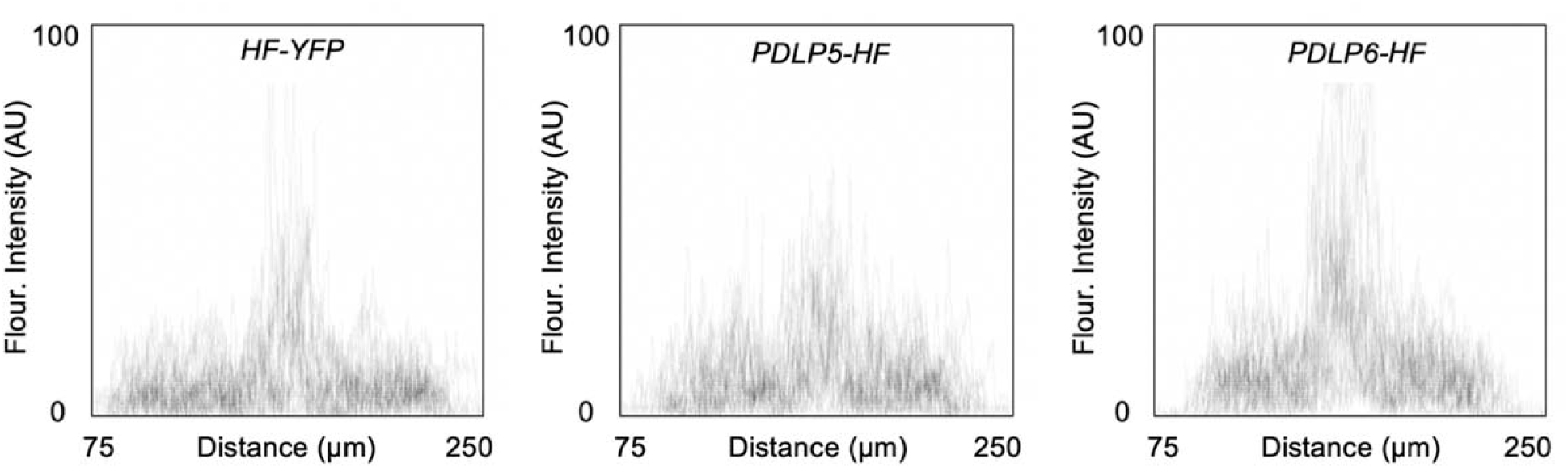
Fluorescence intensity profiles of callose signals in roots. Semi-quantitative evaluation of the relative level of aniline blue-stained callose in root cells (Fig. 3C) was performed by analyzing the signal intensity across different root cell types. Confocal images were captured from the maturation zone of 10-day-old seedlings. 25 individual transgenic plants were analyzed for each genotype and fluorescence intensity profiles (arbitrary unit: AU) of callose accumulation were combined within the genotype. Numbers on the X-axis indicate the distance (µm) across the region analyzed.

**Supplemental Figure 6.**
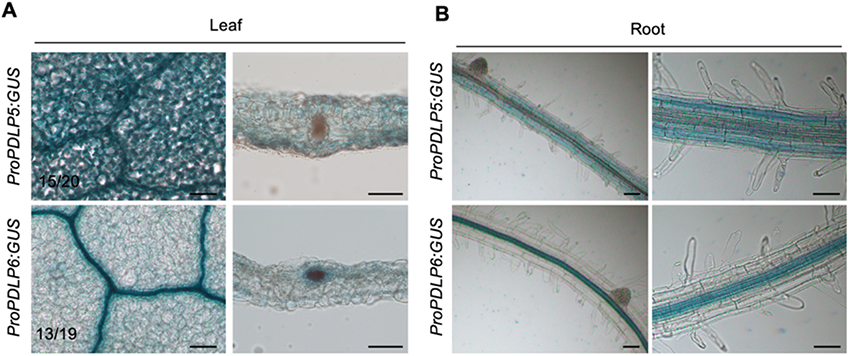
Histochemical staining of GUS activity in Arabidopsis *ProPDLP5:GUS* and *ProPDLP6::GUS* transgenic plants. (**A**) Leaves of 4-week-old Arabidopsis T_1_ transgenic plants were subjected to GUS staining. Images on the left column were captured from the leaf surface. Images on the right column were taken from GUS-stained mature leaf transverse sections. The tissues were embedded in 3% agarose and sectioned with a vibrating blade microtome. The numbers indicate the transgenic plants exhibit the shown GUS activity pattern out of the total independent transgenic plants analyzed. Scale bars = 100 µm. (**B**) Roots of 2-week-old Arabidopsis T_2_ transgenic plants were subjected to GUS staining. Scale bars = 50 µm.

**Supplemental Figure 7.**
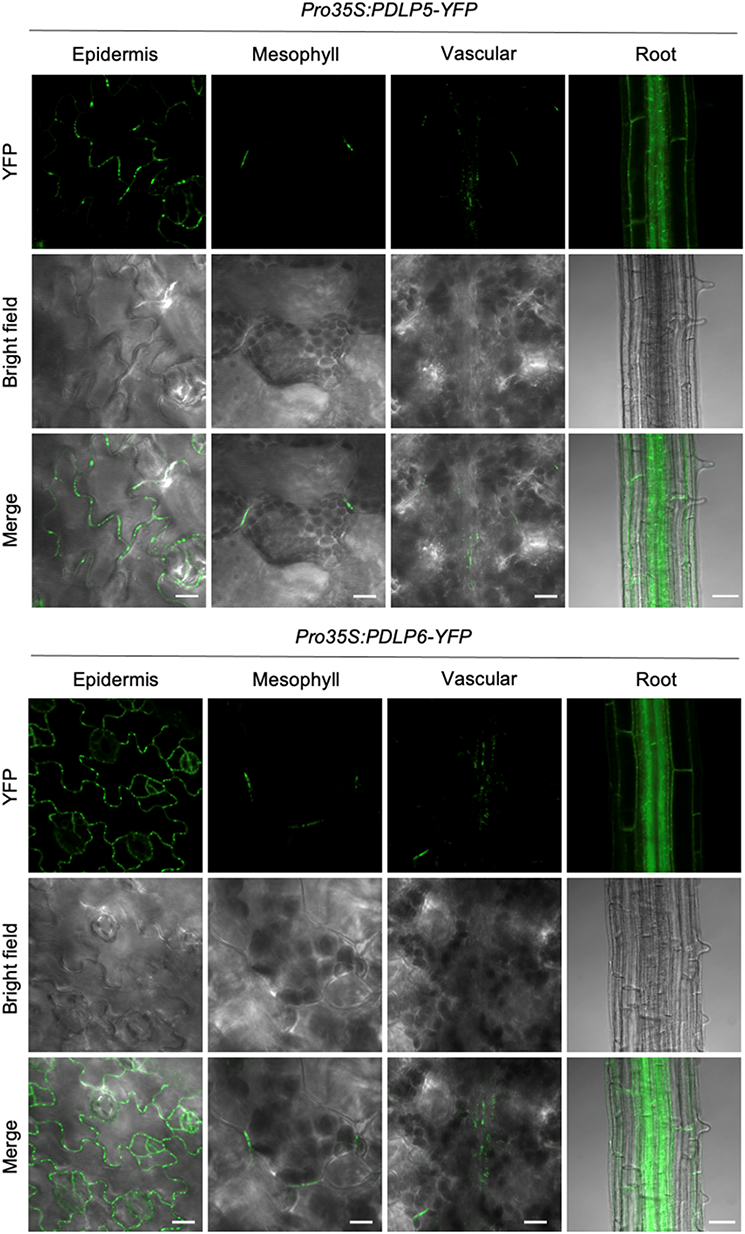
Ubiquitous expression of PDLP5-YFP and PDLP6-YFP fusion proteins in *Pro35S:PDLP5-YFP* and *Pro35S:PDLP6-YFP* transgenic plants. The expression of the fusion proteins was detected in epidermal cells, mesophyll cells, vascular cells in leaves, and most cell types in roots. Confocal images were captured from 2-week-old Arabidopsis seedlings. Green signals represent the expression of YFP fusion proteins. Scale bars for epidermis, mesophyll, and vasculature = 10 µm. Scale bars for root = 50 µm.

**Supplemental Figure 8.**
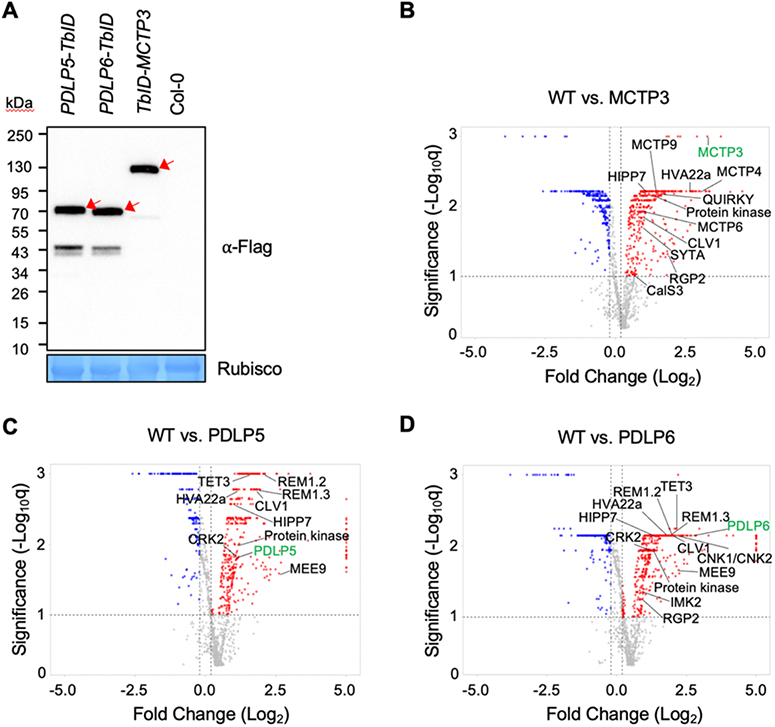
The expression of PDLP5-TbID, PDLP6-TbID, and TbID-MCTP3 and their enriched proteins. (**A**) 2-week-old seedlings of wild-type Col-0 and Arabidopsis transgenic plants were subjected to immunoblot analysis using an anti-Flag antibody. Rubisco serves as a loading control. Red arrows indicate the TbID fusion proteins. (**B-D**), Volcano plots show significantly enriched proteins in MCTP3, PDLP5, and PDLP6 samples compared with WT. Candidates were filtered using cutoffs log_2_FC > 0.2 or < −0.2 and q < 0.1. Plots were generated using VolcaNoseR. A few of the known and putative PD-associated proteins are labeled.

**Supplemental Figure 9.**
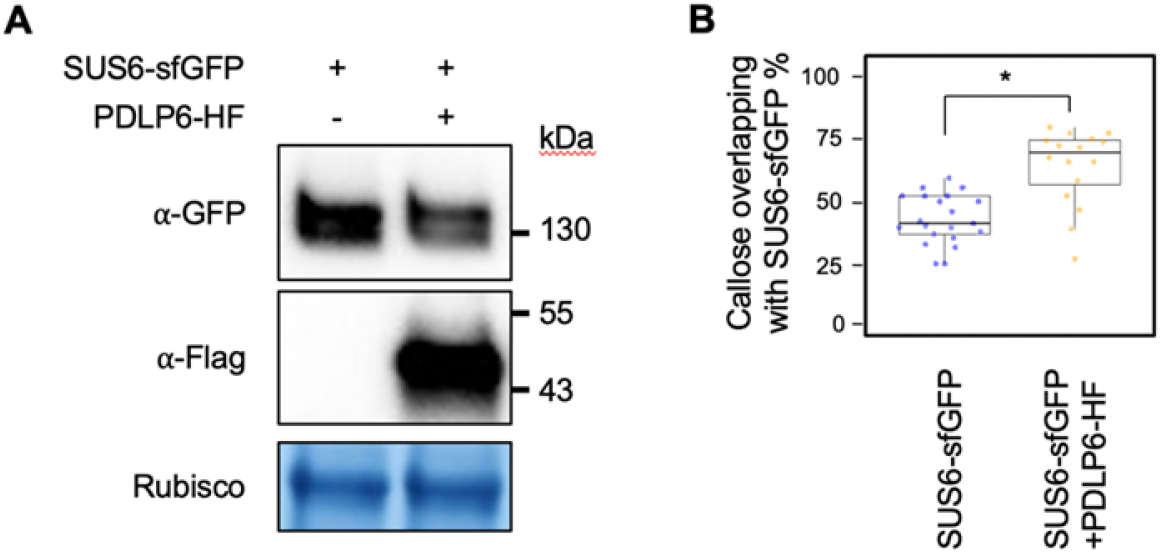
The expression of PDLP6-HF enhances the PD association of SUS6-sfGFP. (**A**) Immunoblot analysis detects the expression of SUS6-GFP and PDLP6-HF transiently expressed in *N. benthamiana*. Anti-Flag and anti-GFP antibodies were used to detect the expression of Flag-and GFP-fusion proteins, respectively. Rubisco serves as a loading control. (**B**) Quantitative data show that the expression of PDLP6-HF enhances the PD association of SUS6-sfGFP. Agrobacteria harboring *SUS6-sfGFP* and *PDLP6-HF* were co-infiltrated into *N*. *benthamiana*. Aniline blue stained-callose signals overlapping with sfGFP signals were quantified. SUS6-sfGFP, n = 21; and SUS6-sfGFP + PDLP6-HF, n = 16. An asterisk indicates differences that are statistically significant (Mann-Whitney *U* Test; two-paired; *P* <0.01).

**Supplemental Figure 10.**
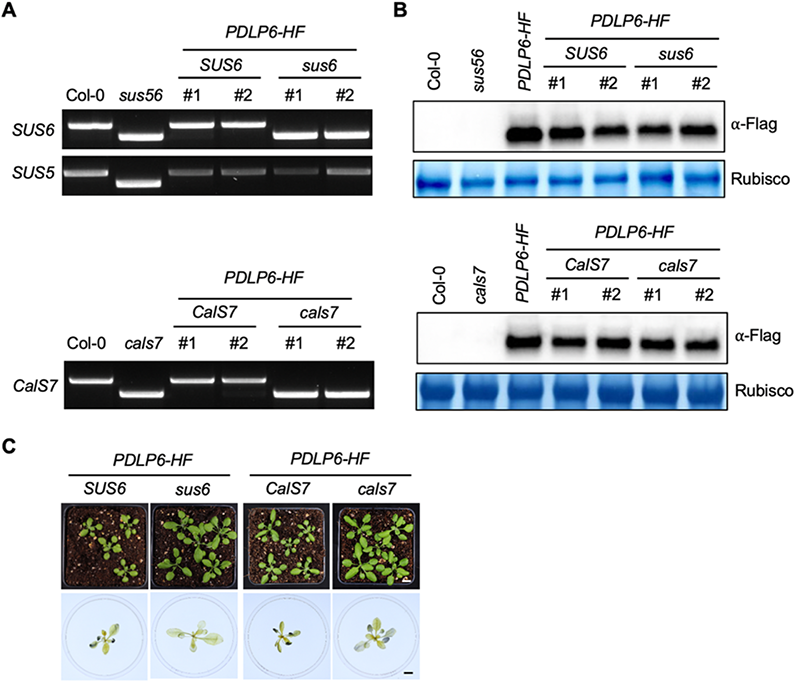
Characterization of progenies of *sus56* × *PDLP6-HF* and *cals7* × *PDLP6-HF*. (**A**) PCR Genotyping analysis. A pair of primers (LP and RP) was used to amplify genomic DNA and another pair of primers (SALK LBb1.3 and RP) was used to amplify the presence of t-DNA for each genotype. Upper bands indicate the PCR products of genomic amplifications. The lower bands indicate the presence of T-DNA insertions in both chromosomes (homozygous lines). (**B**) Immunoblot analysis detects the expression of PDLP6-HF. An anti-Flag antibody was used to detect the expression of Flag-fusion proteins. Rubisco serves as a loading control. (**C**) Independent F_3_ progenies of *sus56* crossing *Pro35S::PDLP6-HF (PDLP6-HF)* and *cals7* crossing *Pro35S::PDLP6-HF (PDLP6-HF)* shown in Fig. 5, C **and** D. Images were taken using the same magnification. Scale bars = 1 cm. Plants were collected at the end of the night and subjected to starch staining.

**Supplemental Figure 11.**
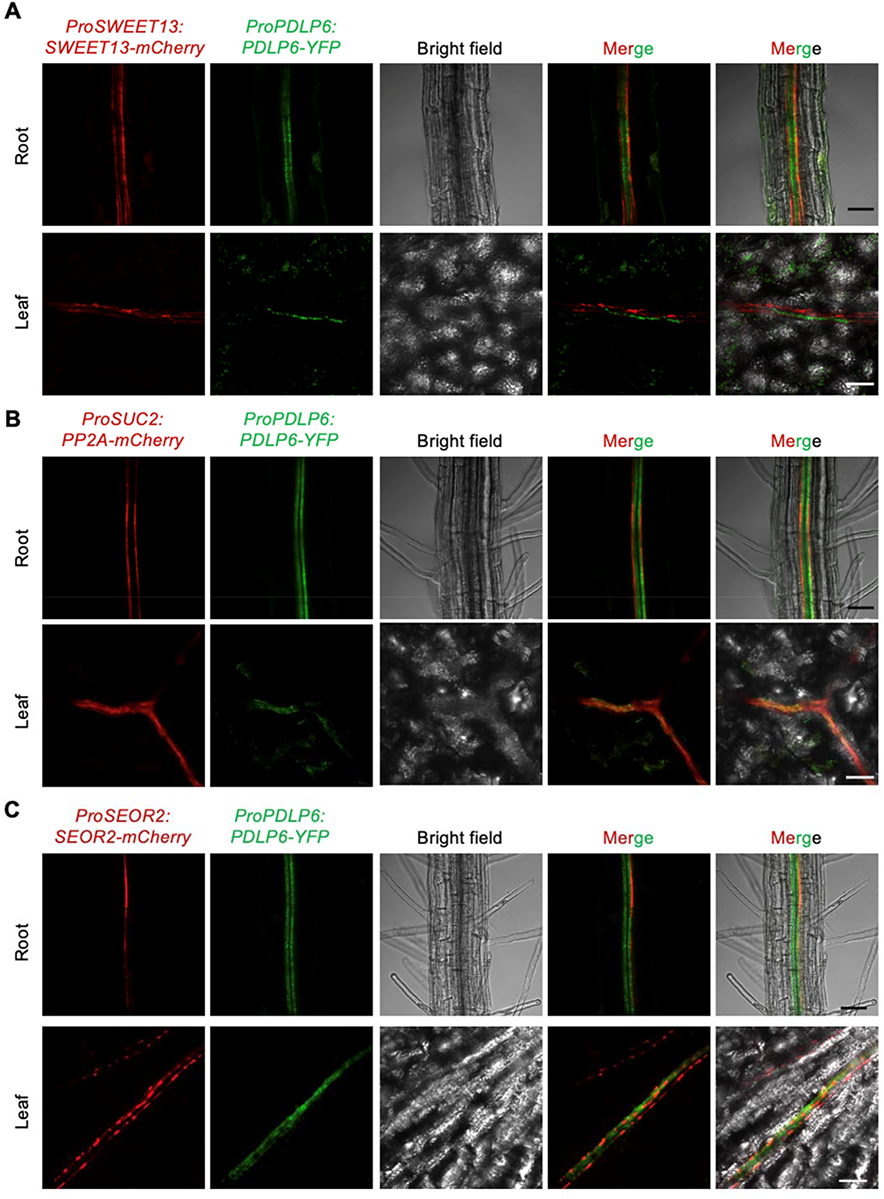
Cell type-specific expression of PDLP6. Confocal images show the expression of PDLP6-YFP in Arabidopsis transgenic plants expressing different cell type markers. (**A**) *ProSWEET13:SWEET13-mCherry* is a PP marker. (**B**) *ProSUC2:PP2A-mCherry* is a CC marker. (**C**) *ProSEOR2:SEOR2-mCherry* is a SE marker. *ProPDLP6:PDLP6-YFP* was introduced into the marker lines using genetic crosses. 2-week-old seedlings were subjected for confocal imaging. Images were captured from roots (top panel) and leaves (lower panel) of F_3_ populations of the transgenic lines expressing both mCherry and YFP. Scale bars = 50 µm.

**Supplemental Table 1.**
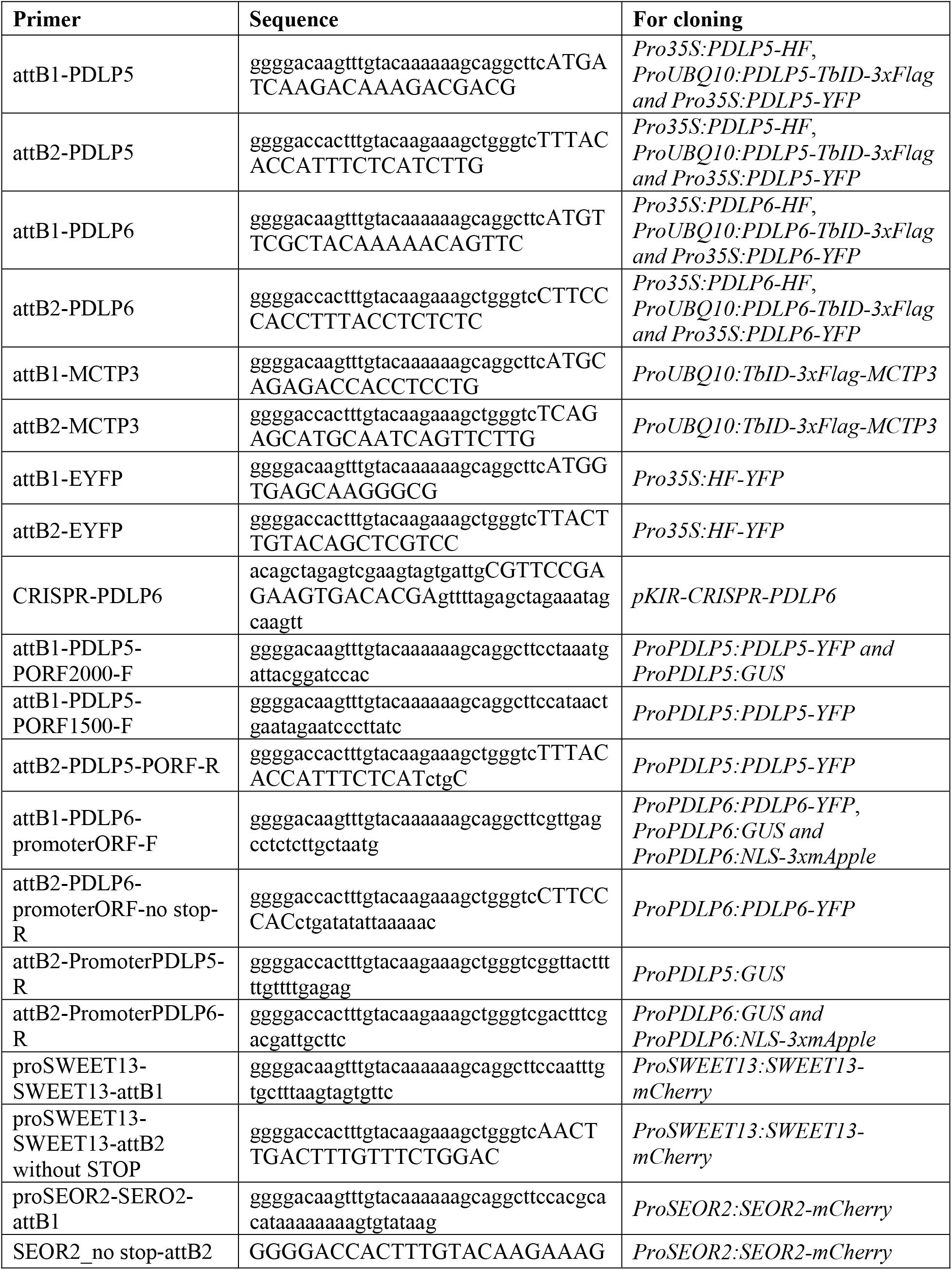
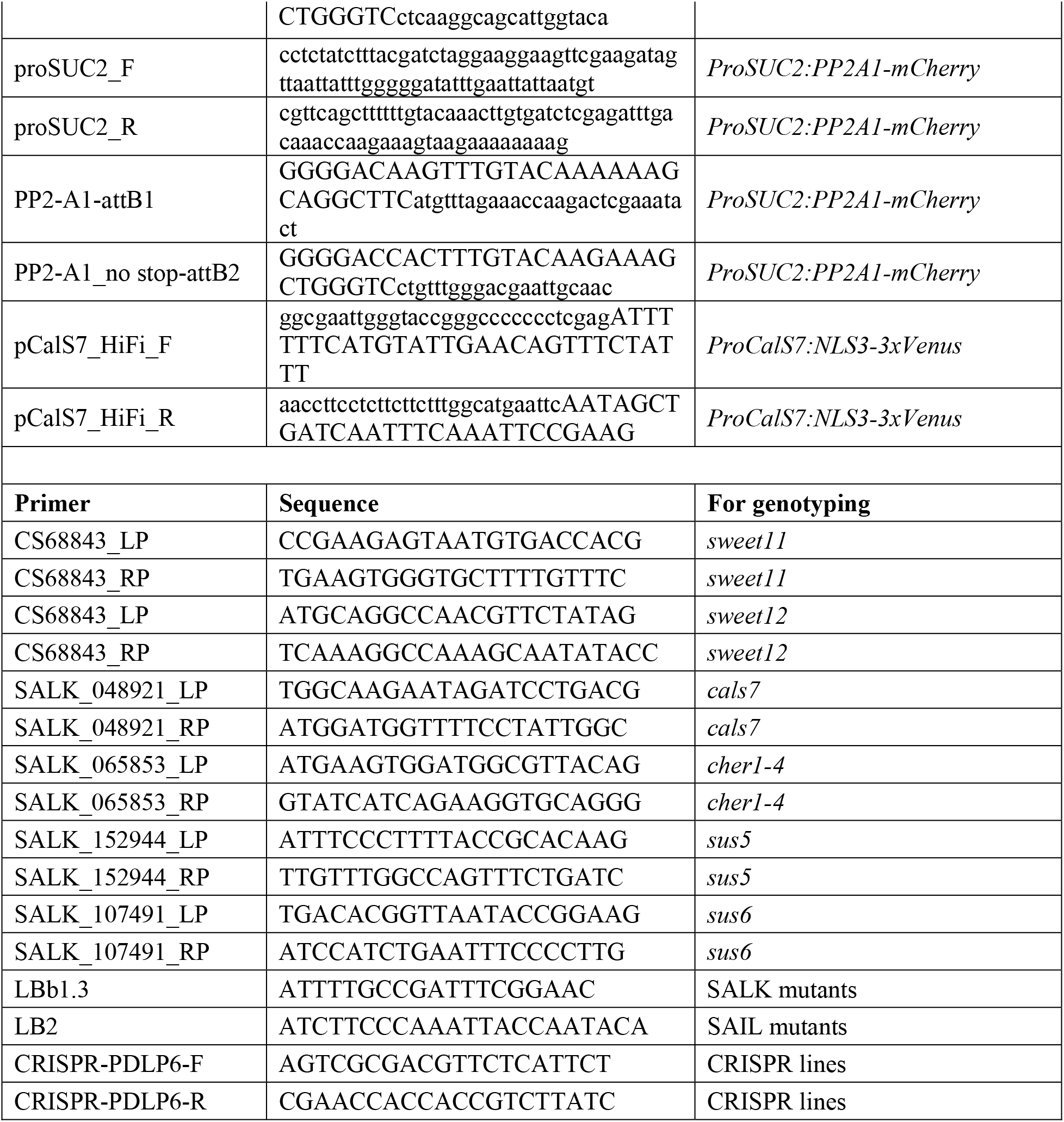
Primers used in this study.

**Supplemental Table 2.**
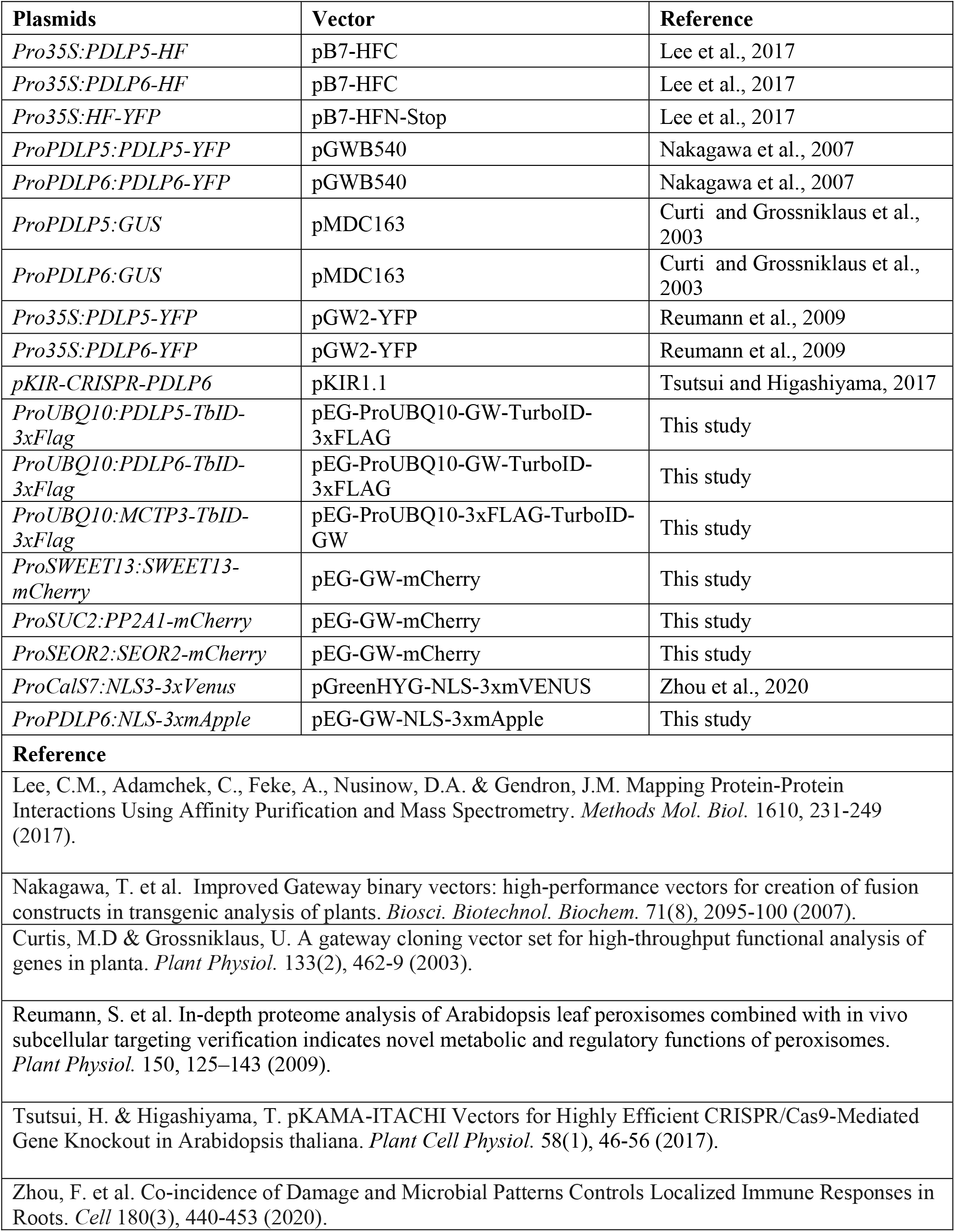
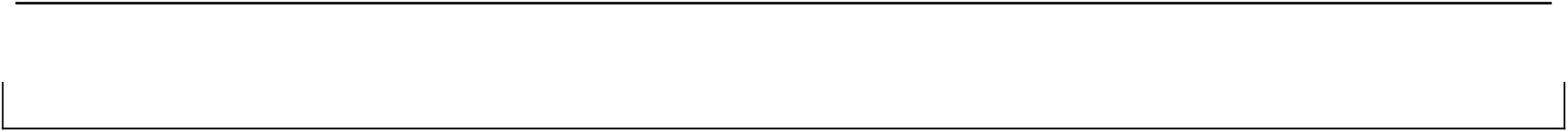
Plasmids used in this study.

**Supplemental Data Set 1. TbID data sheets (provided as a separate file .xlsx)**

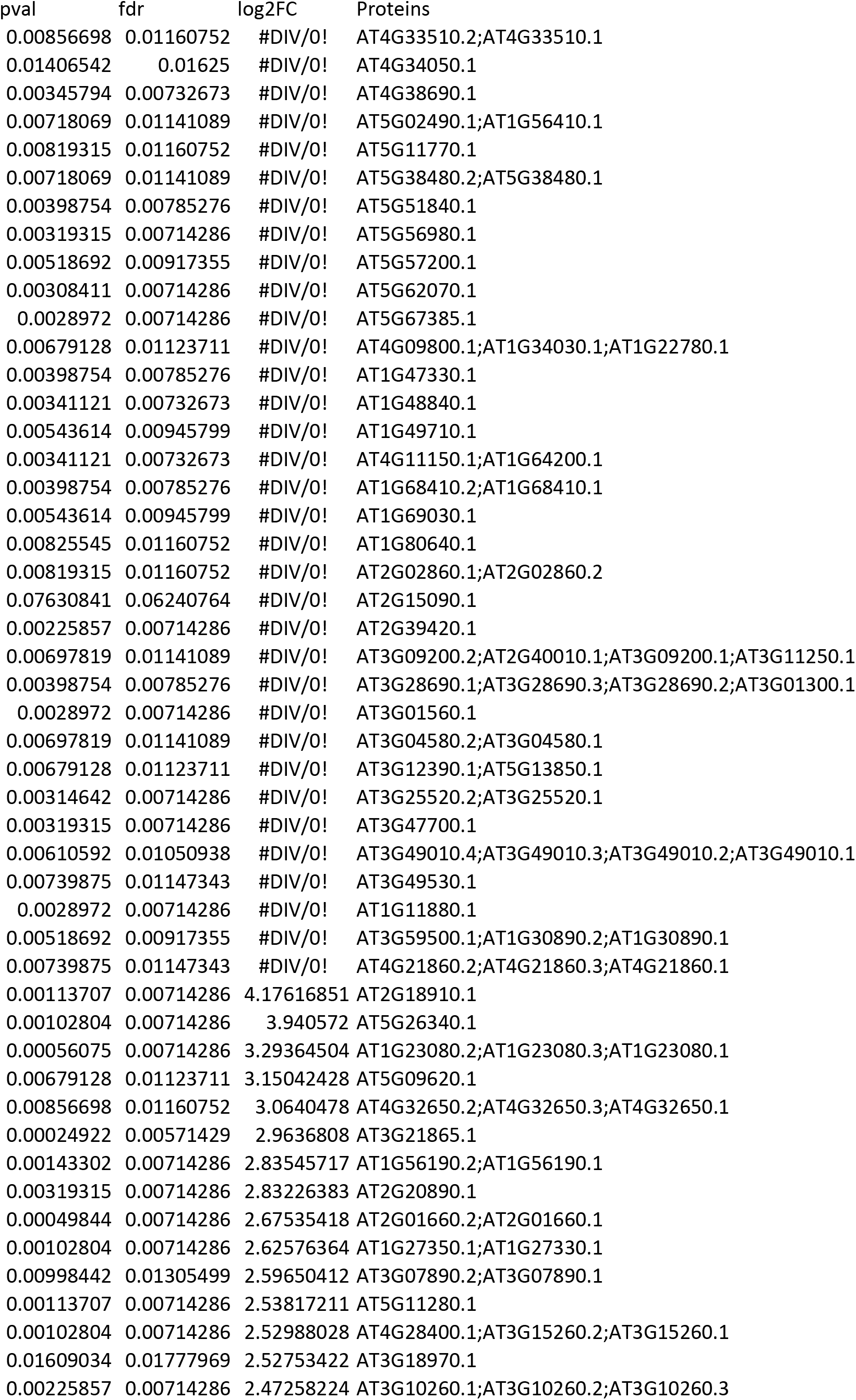
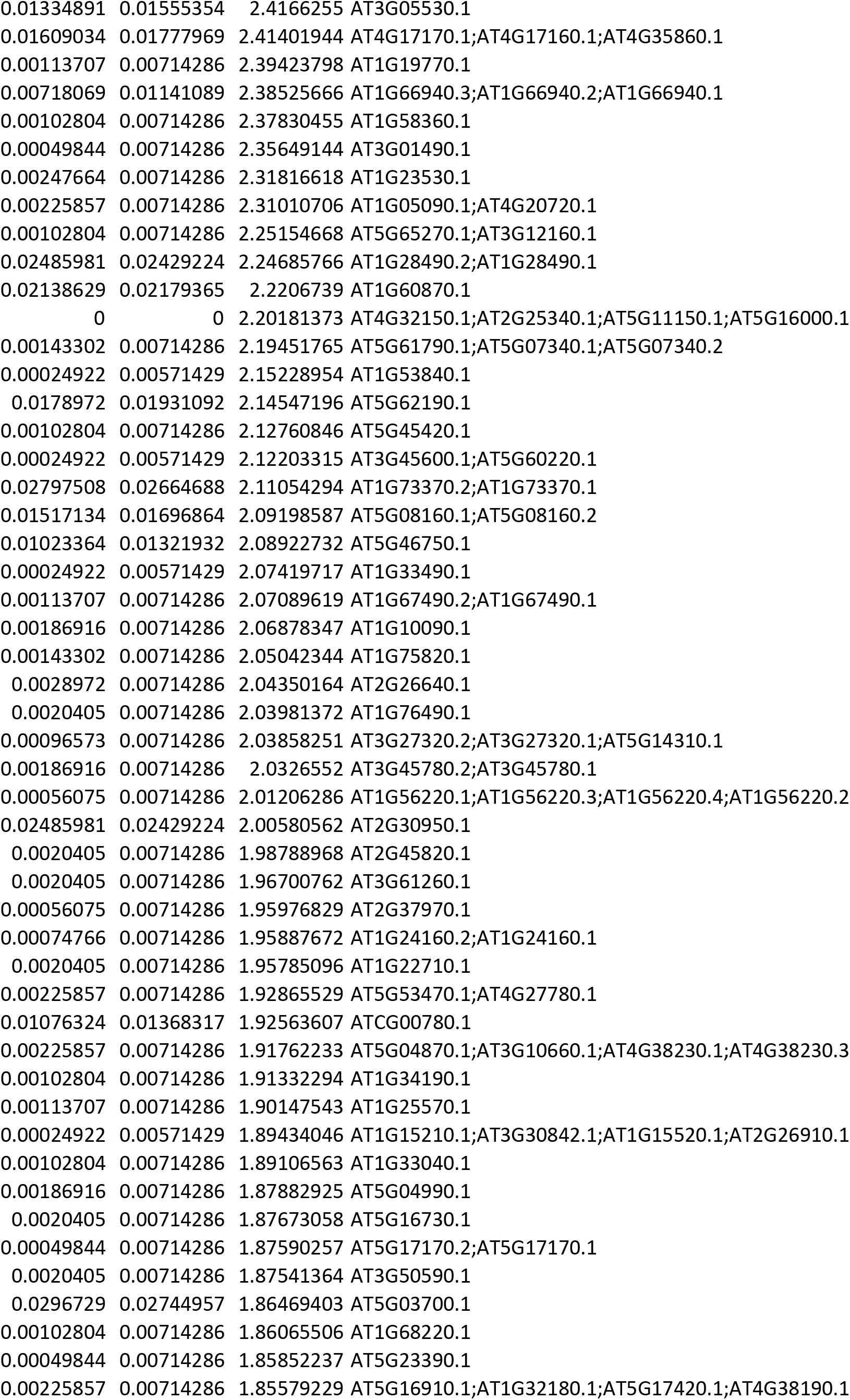
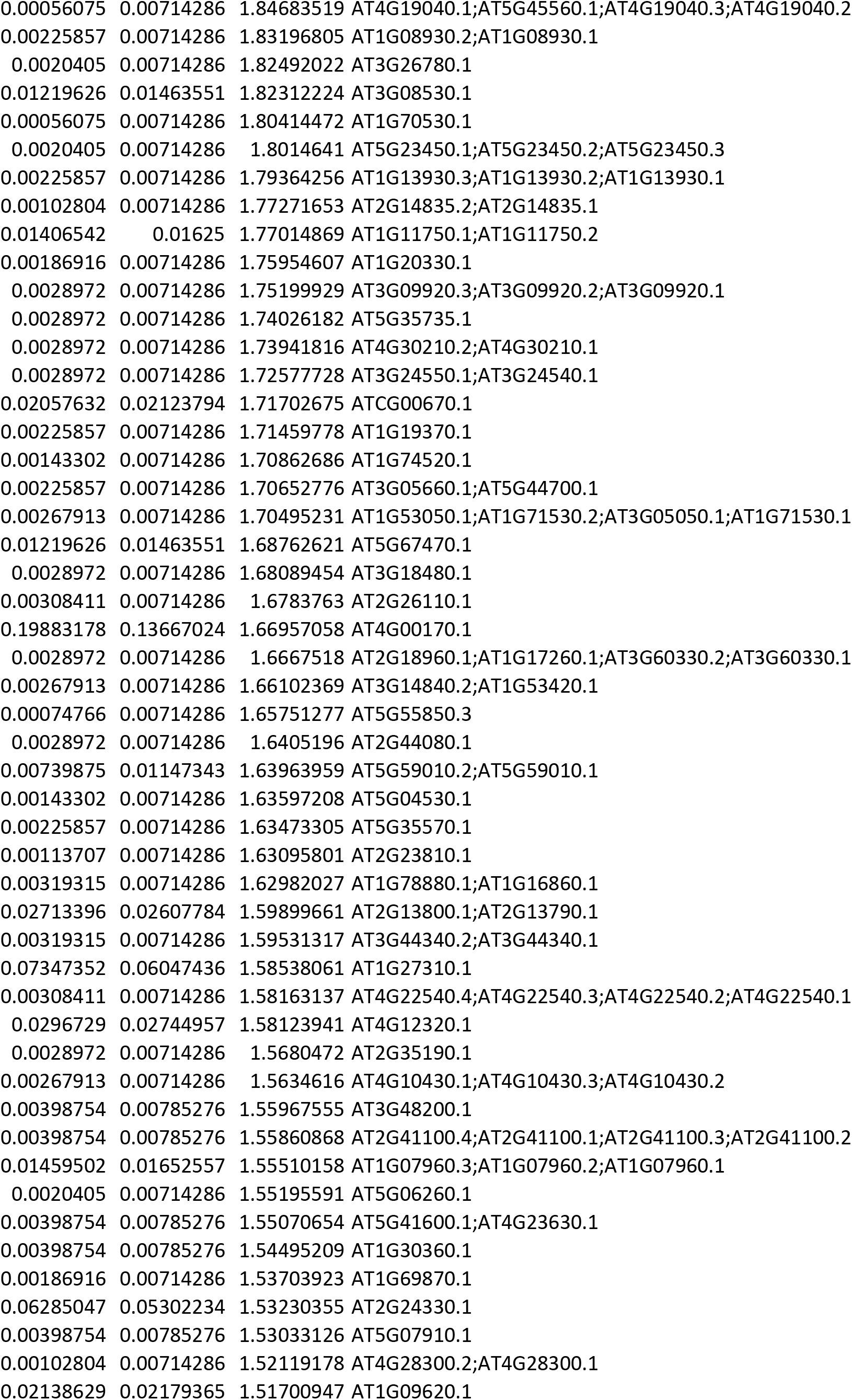
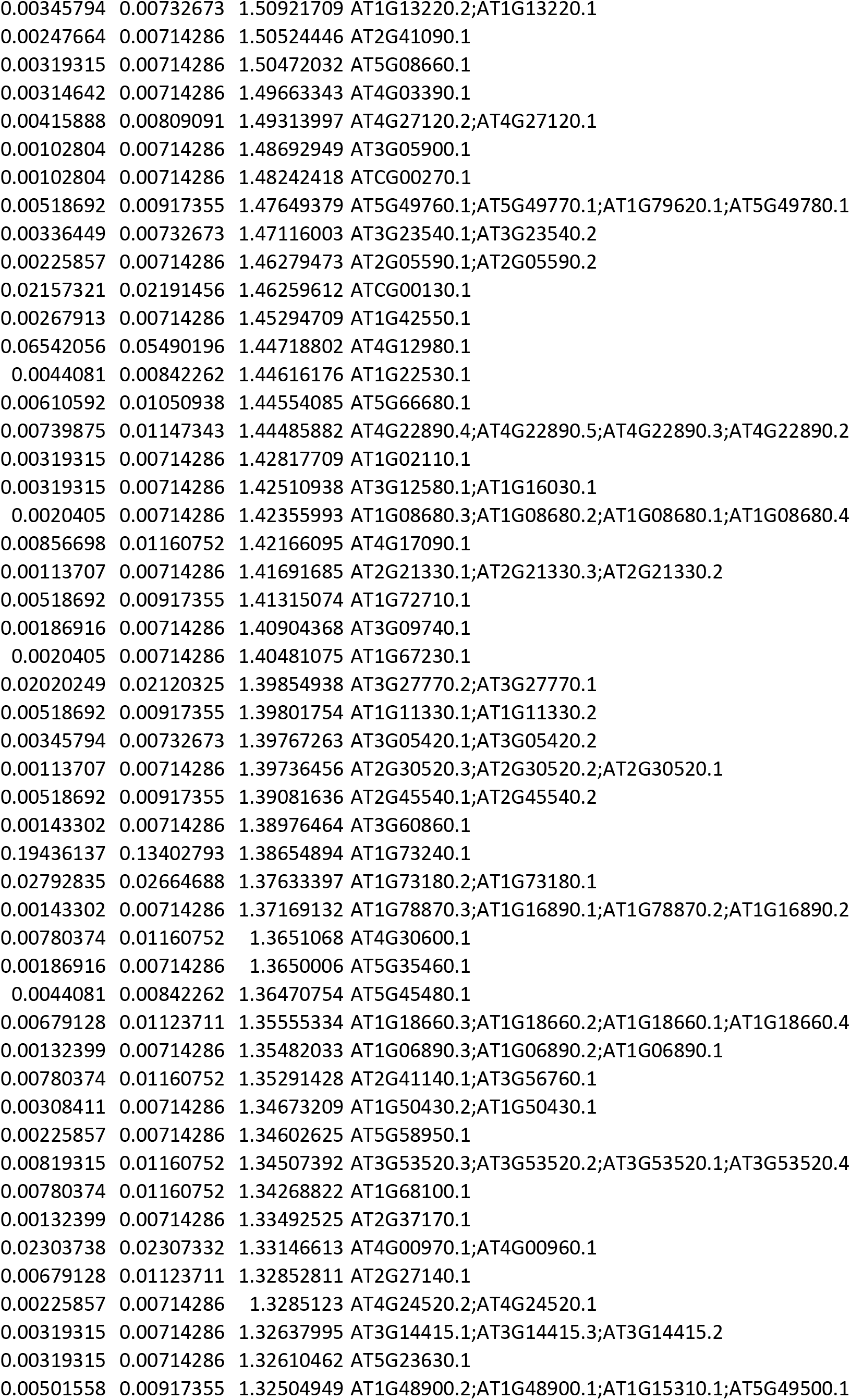
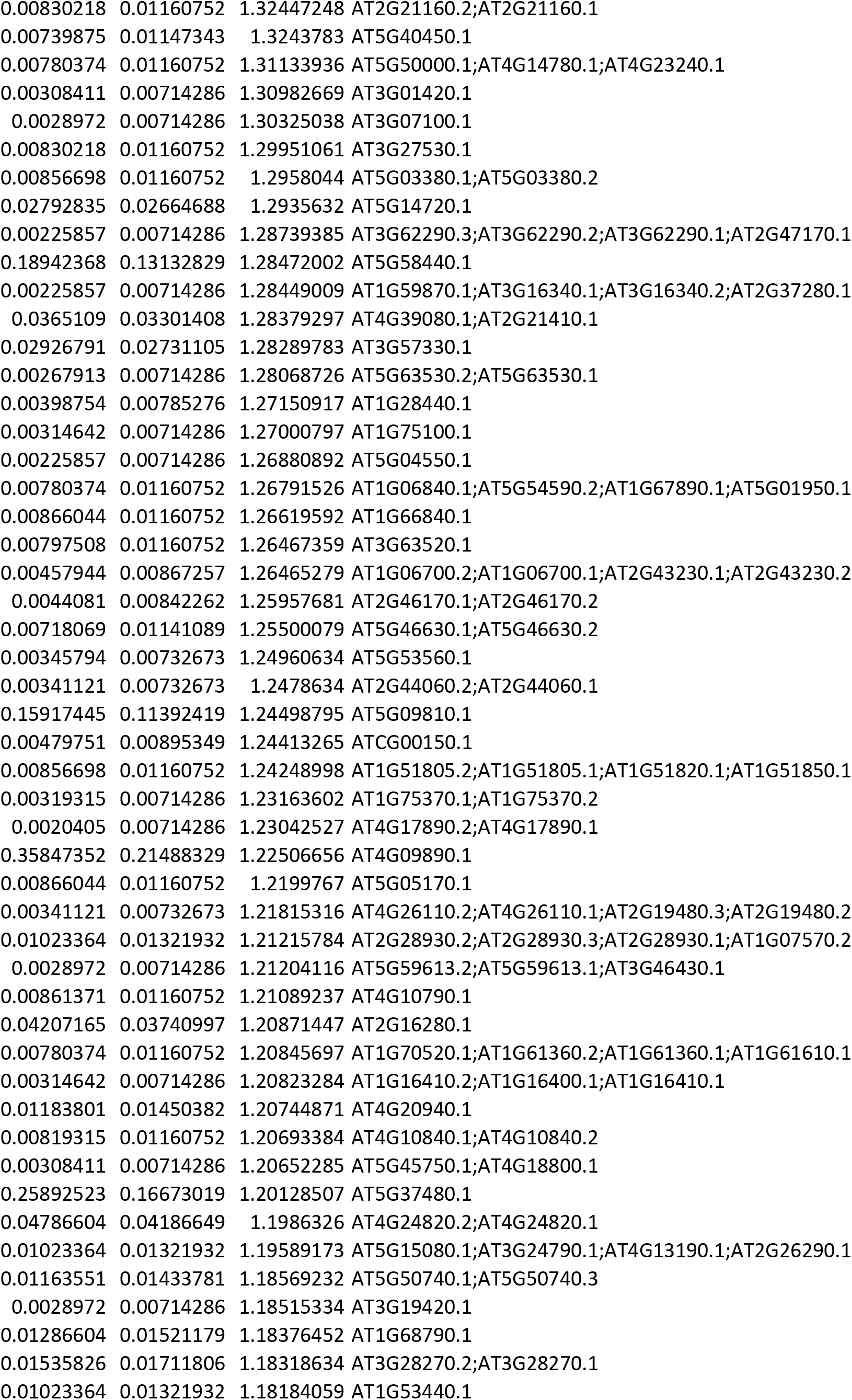
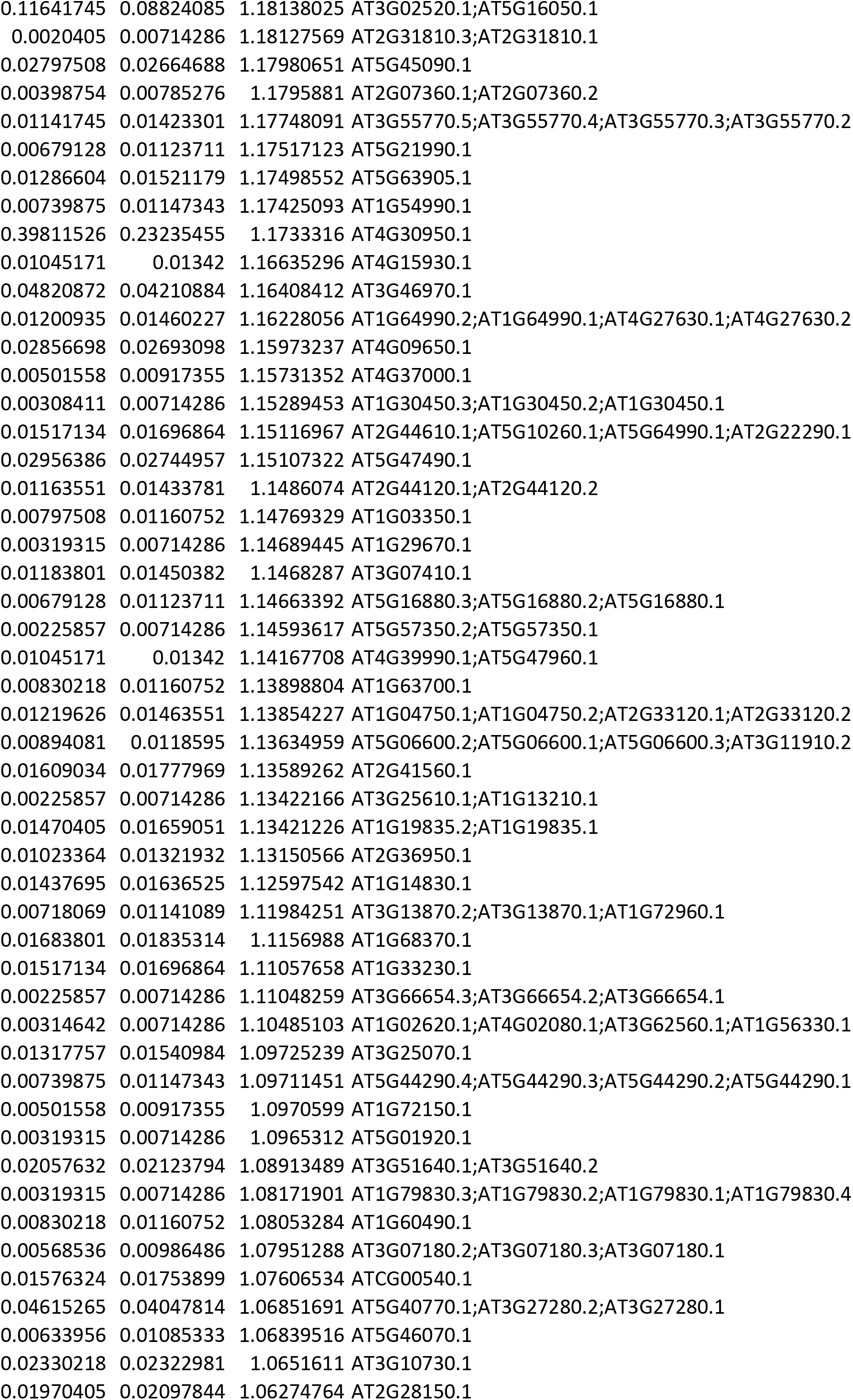
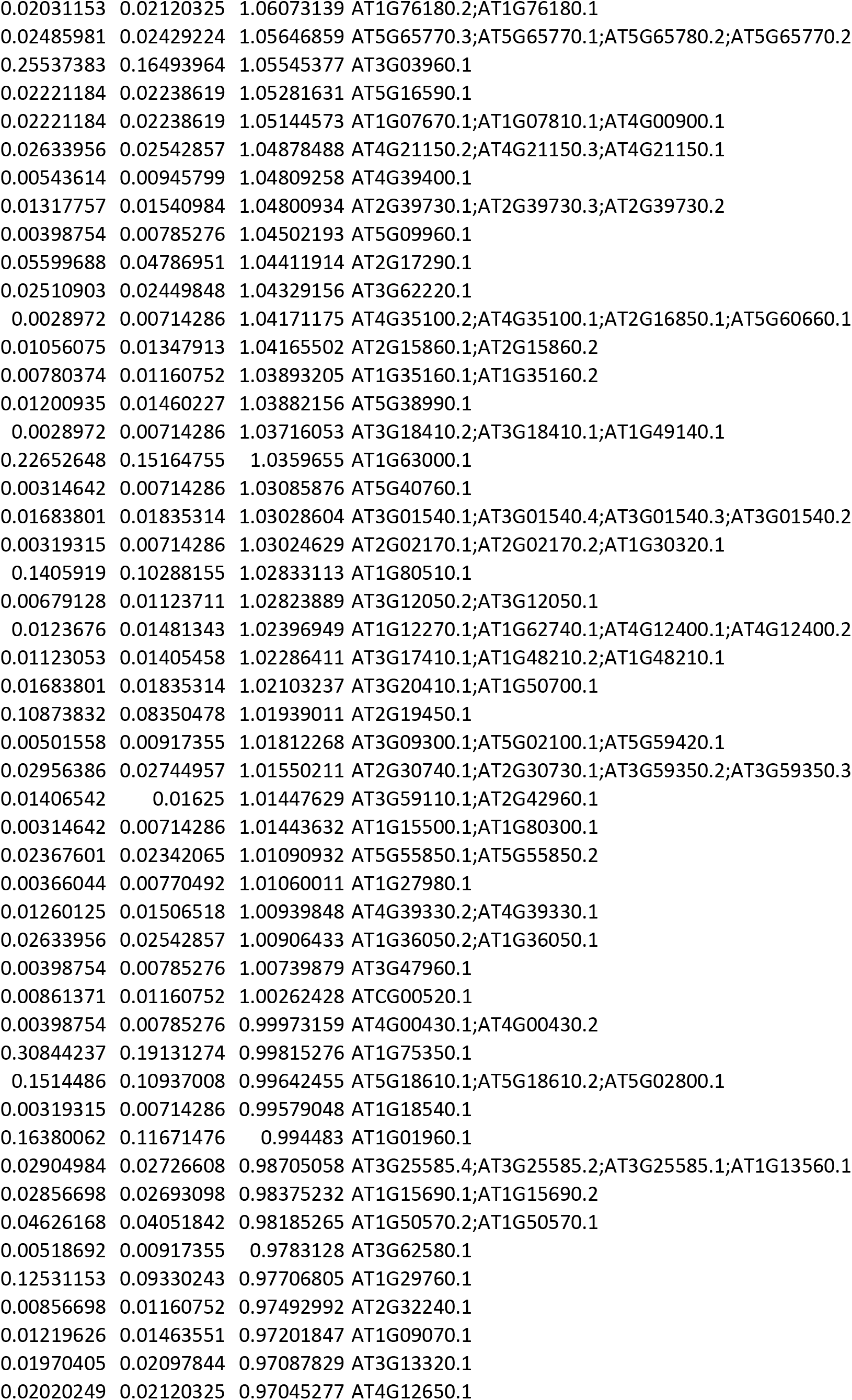
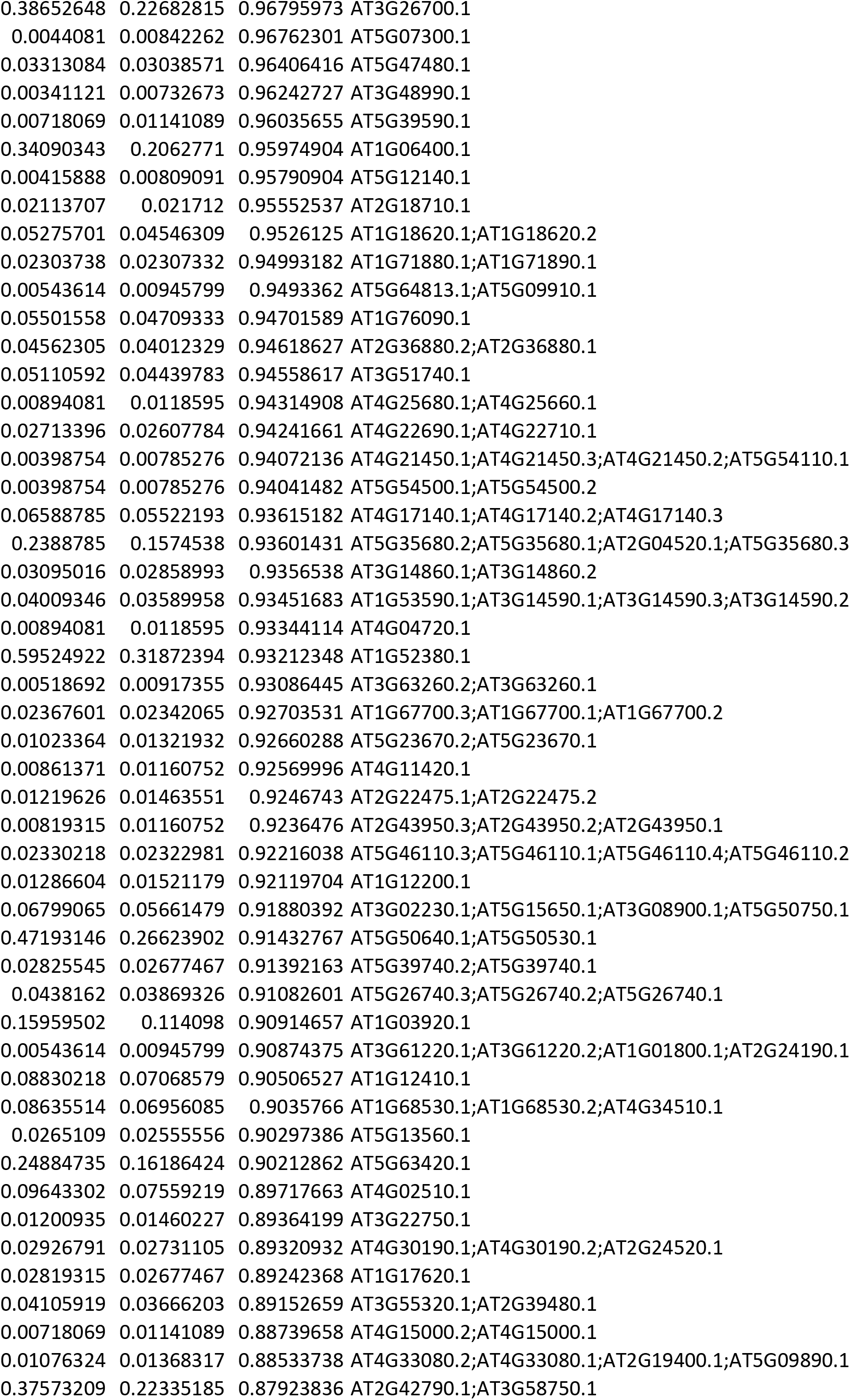
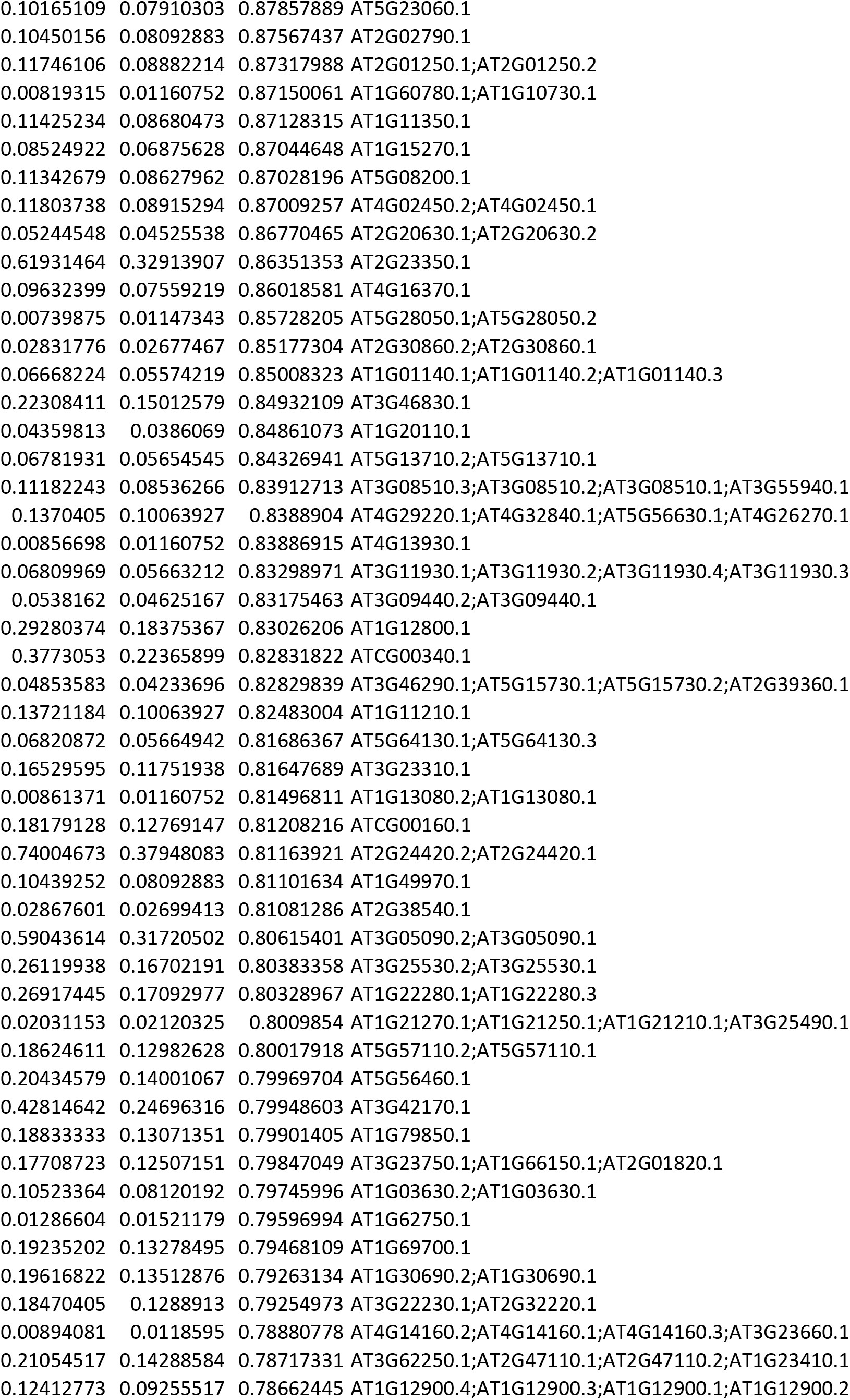
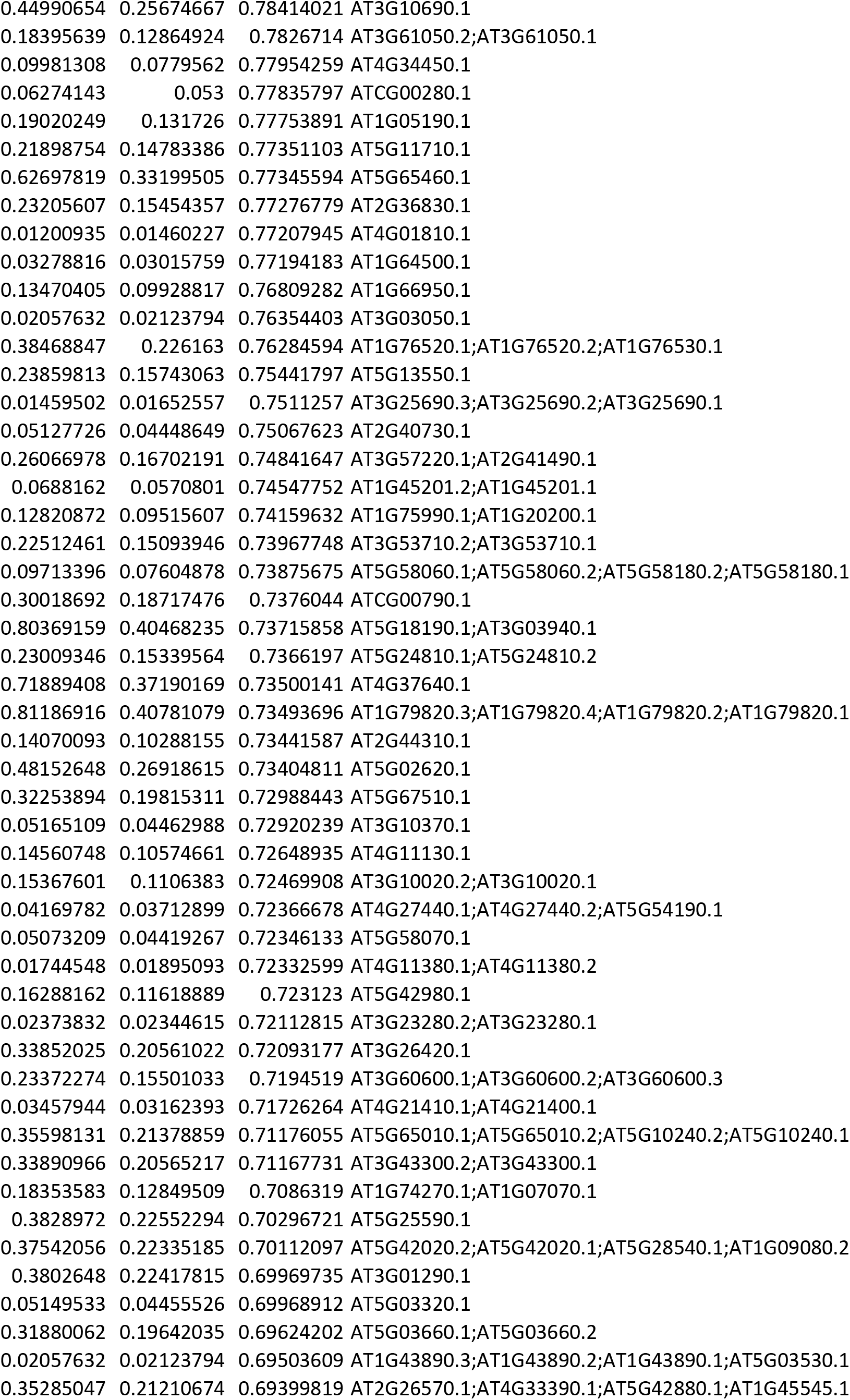
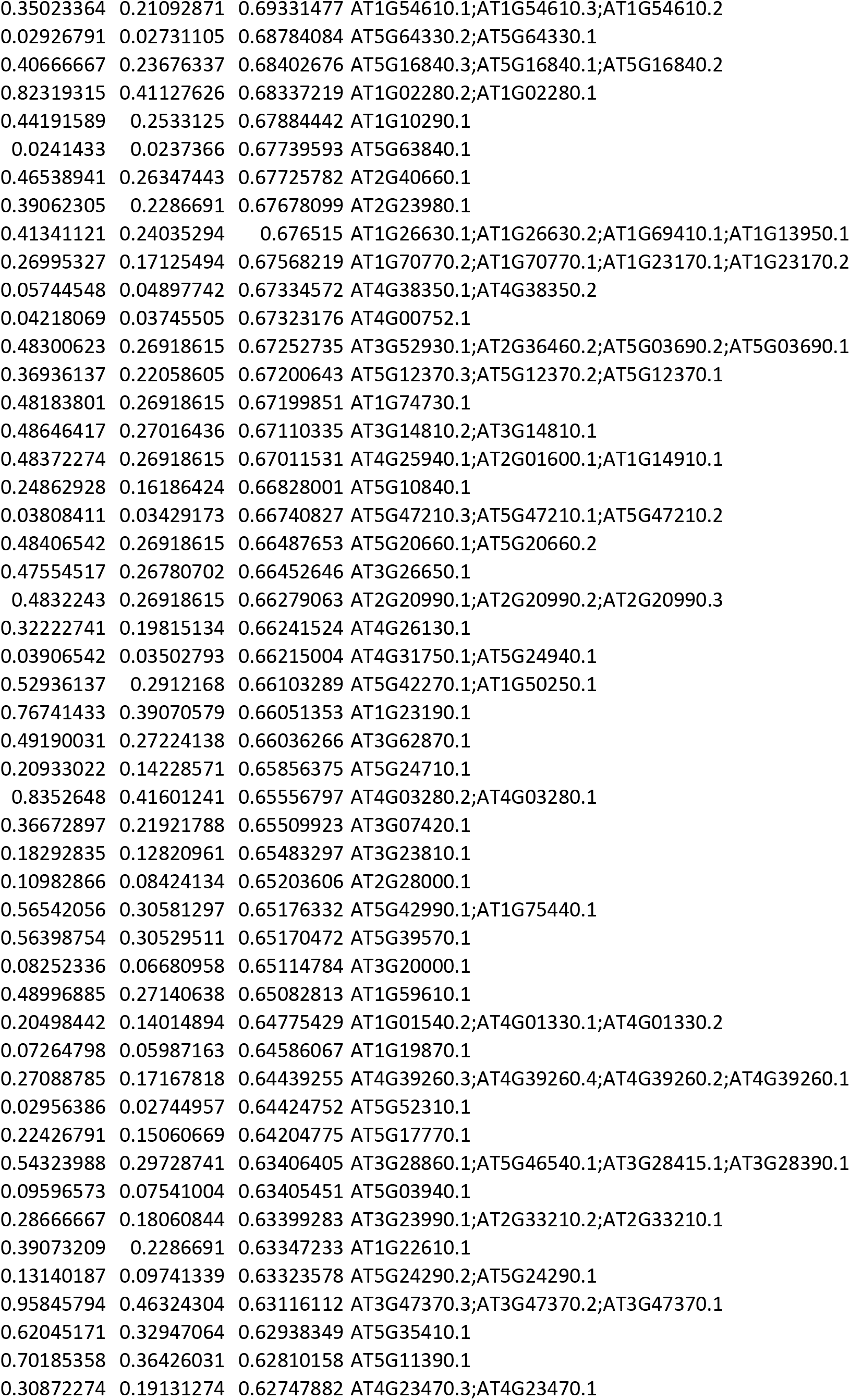
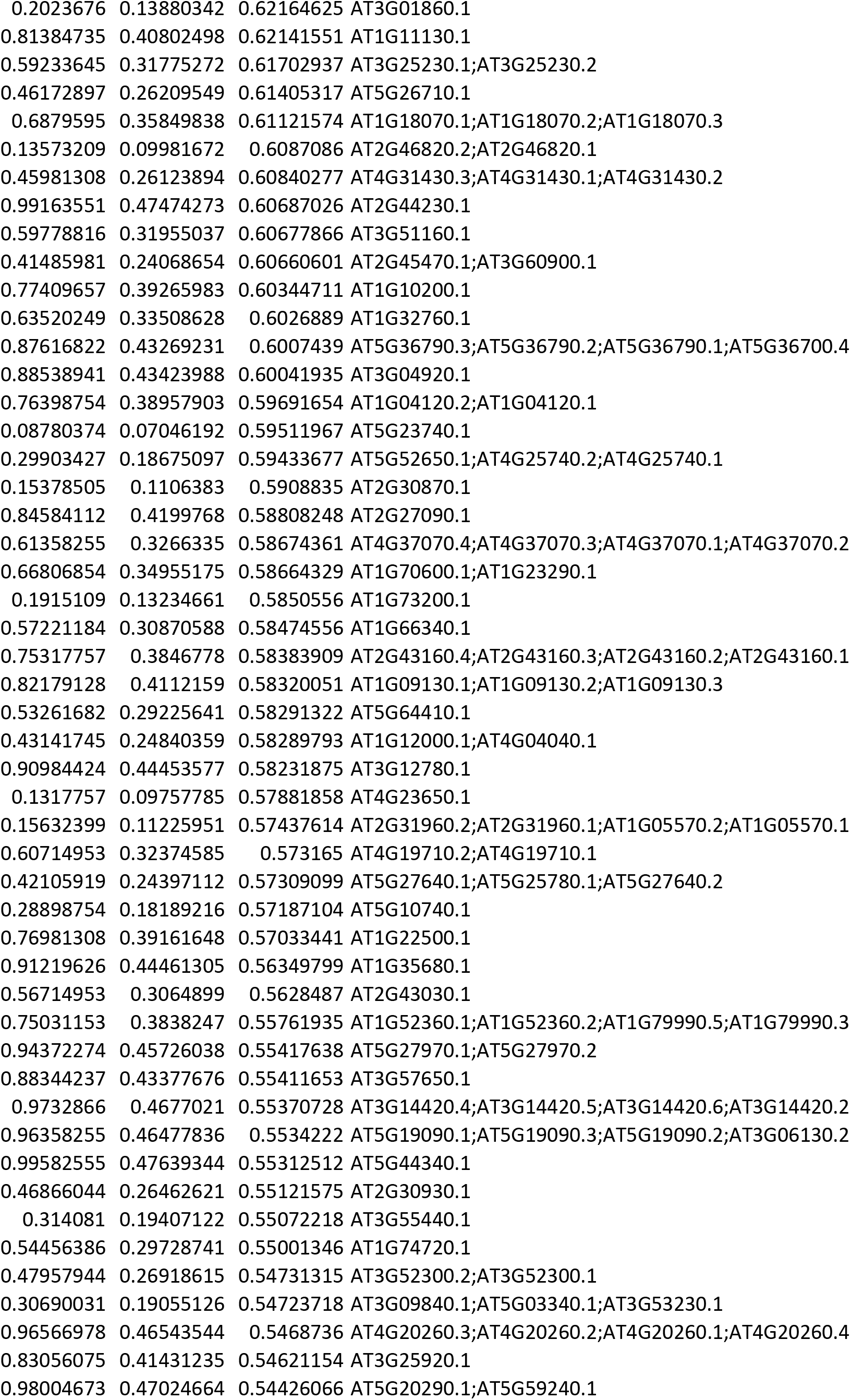
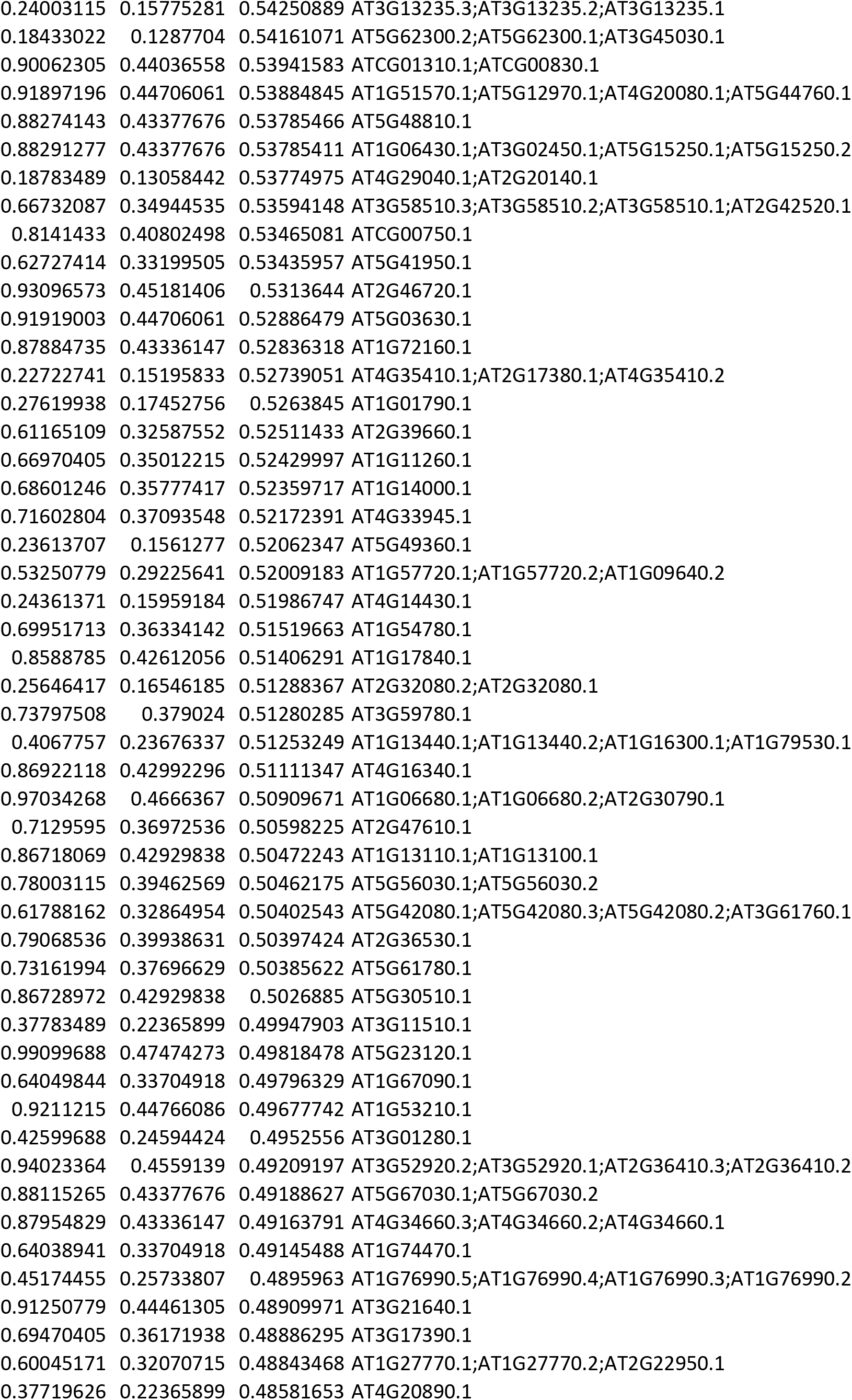
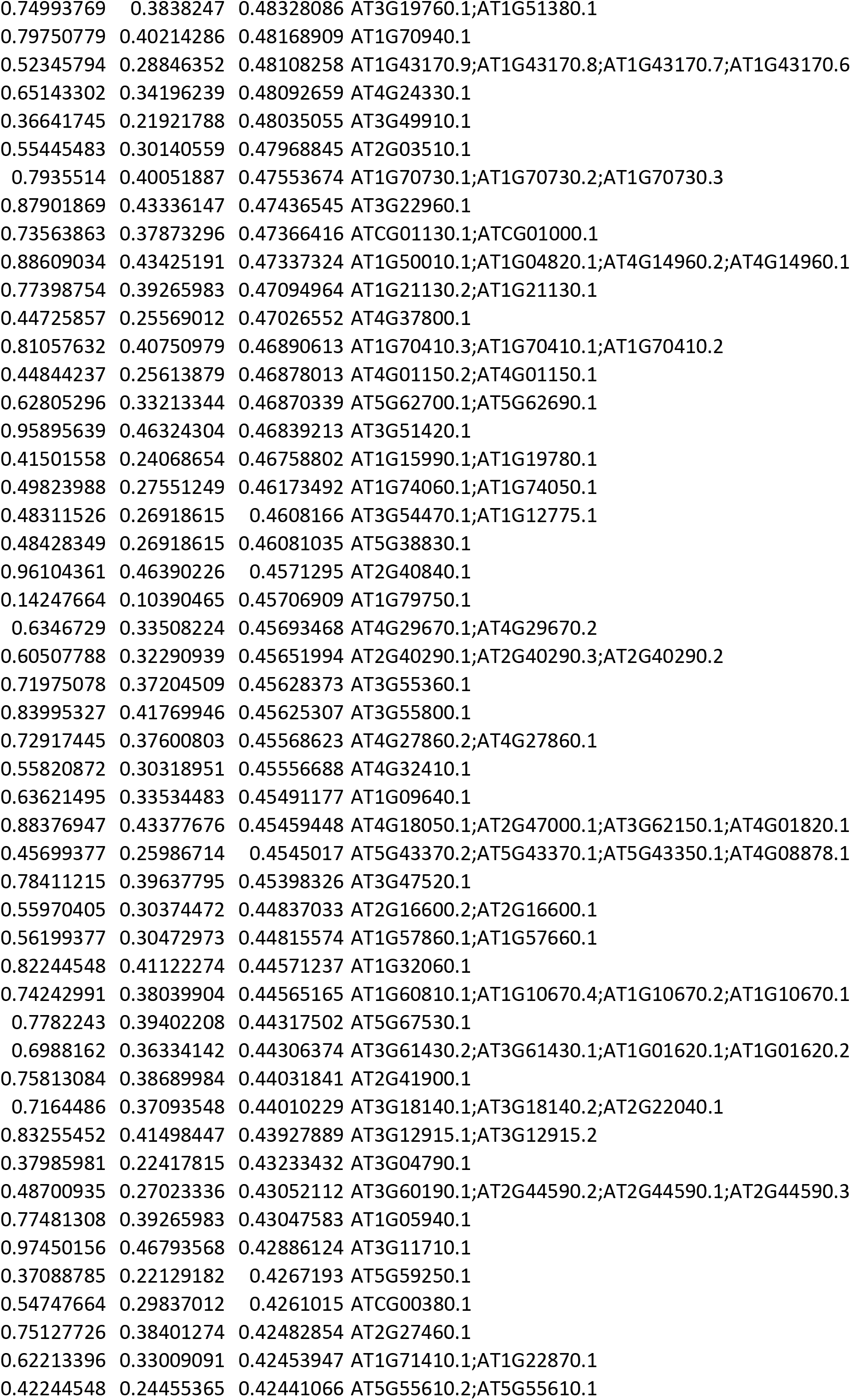
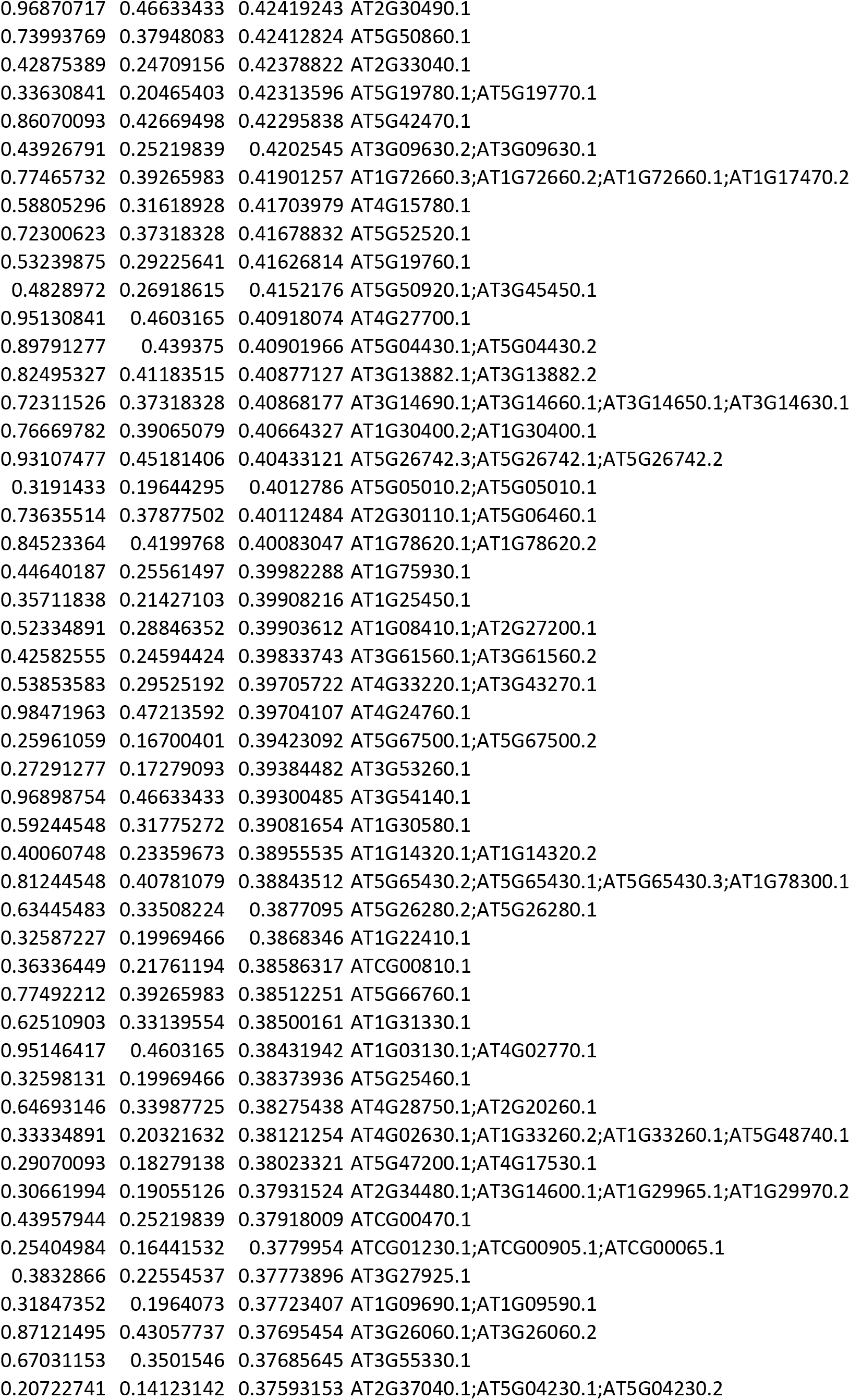
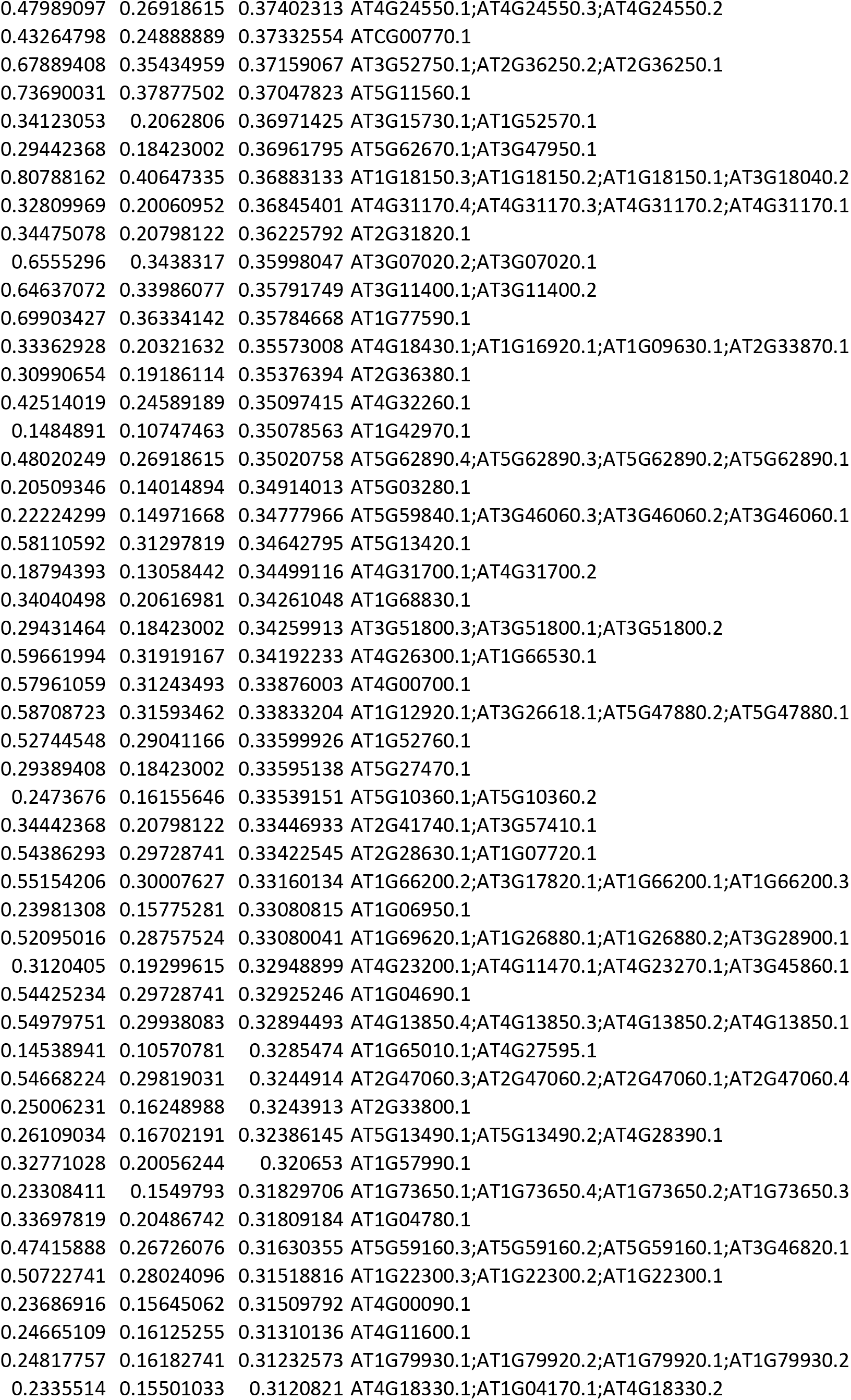
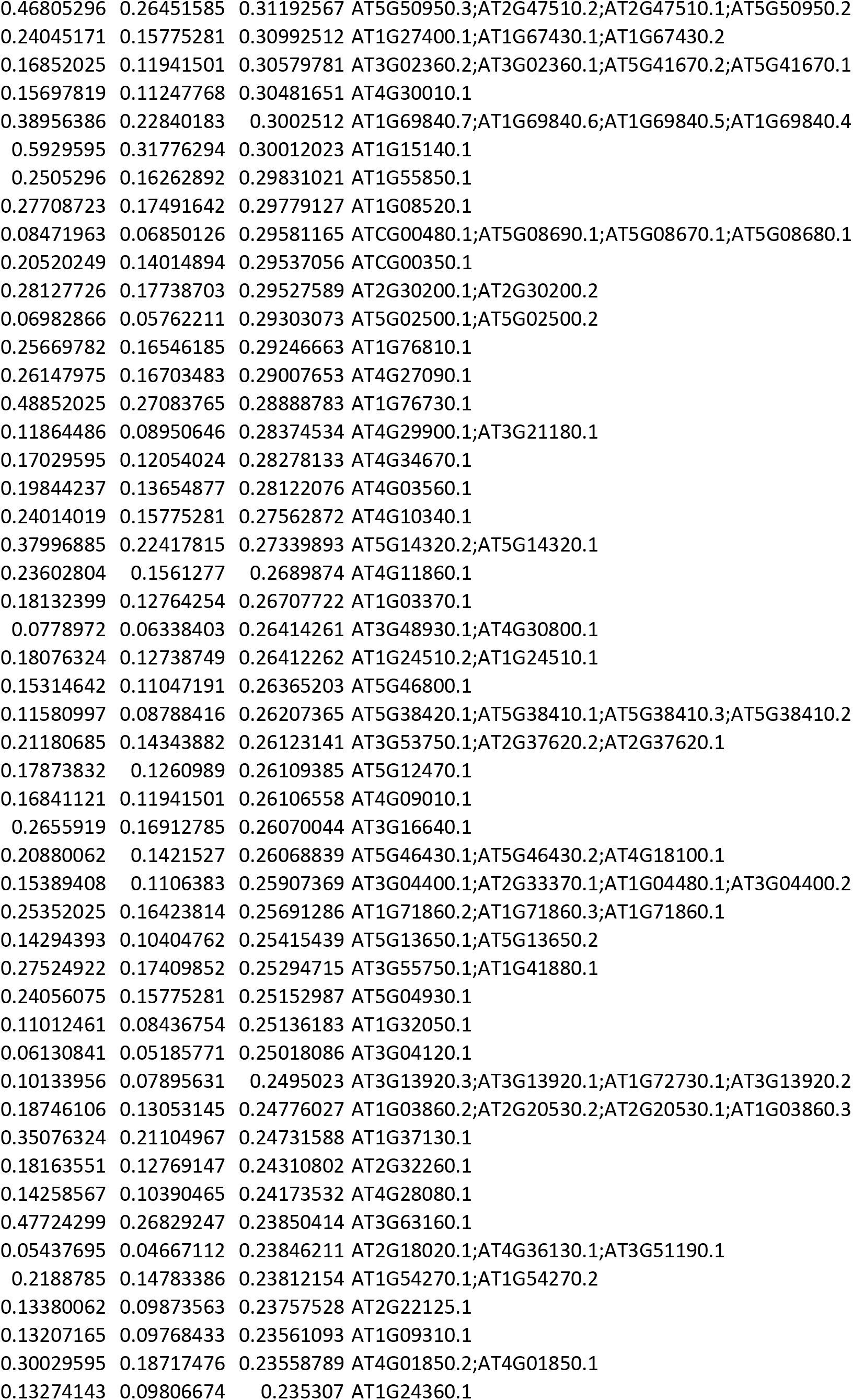
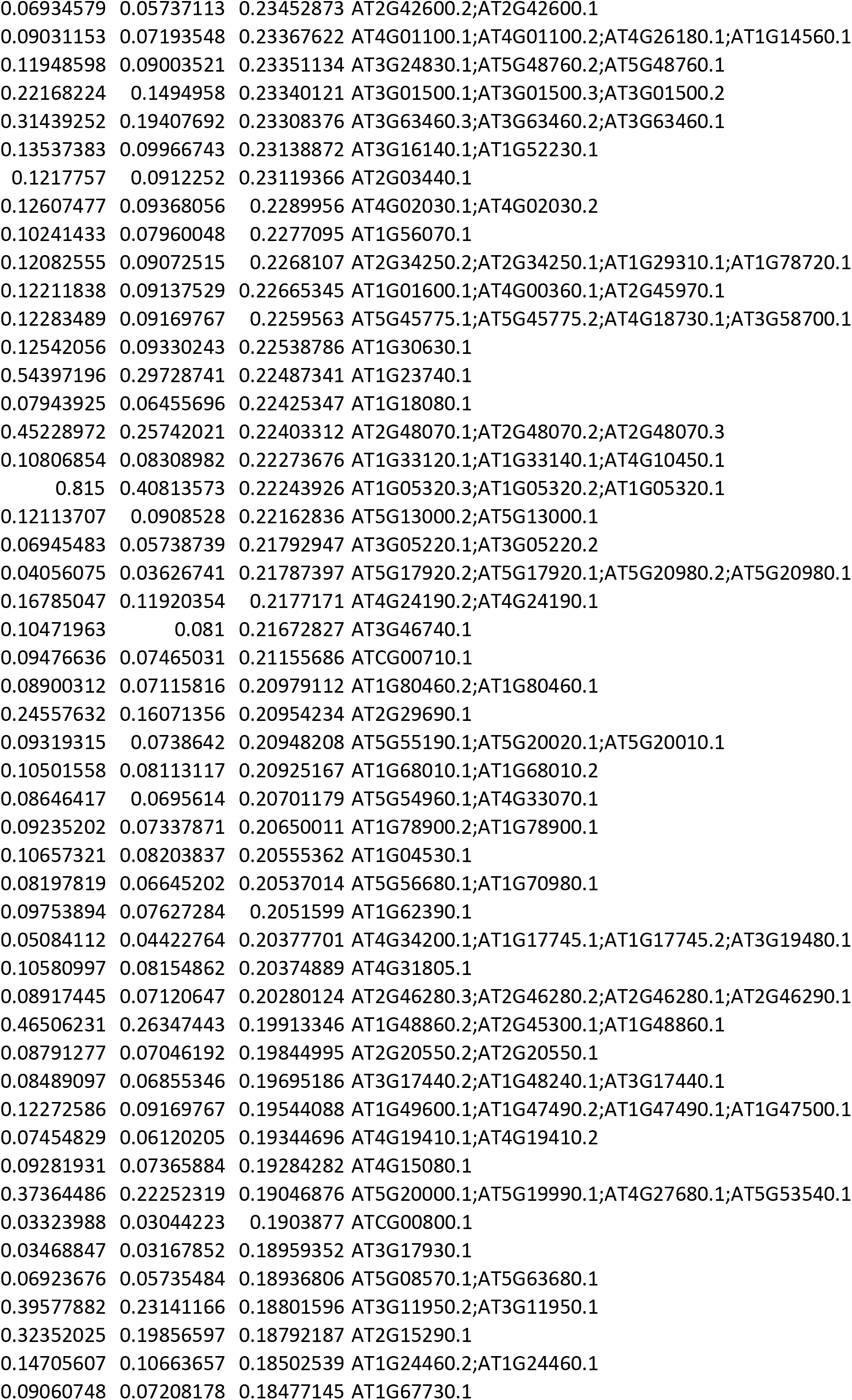
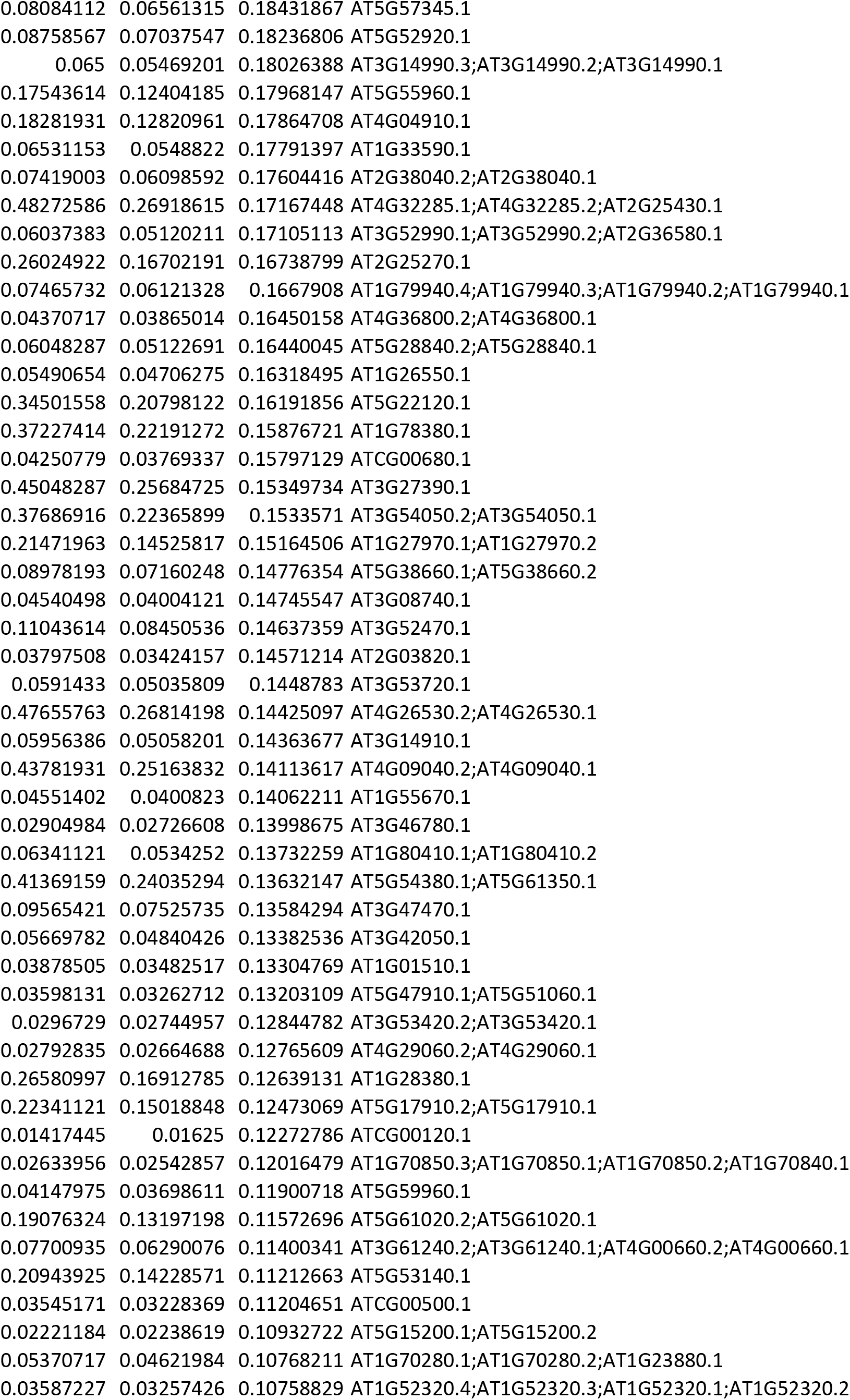
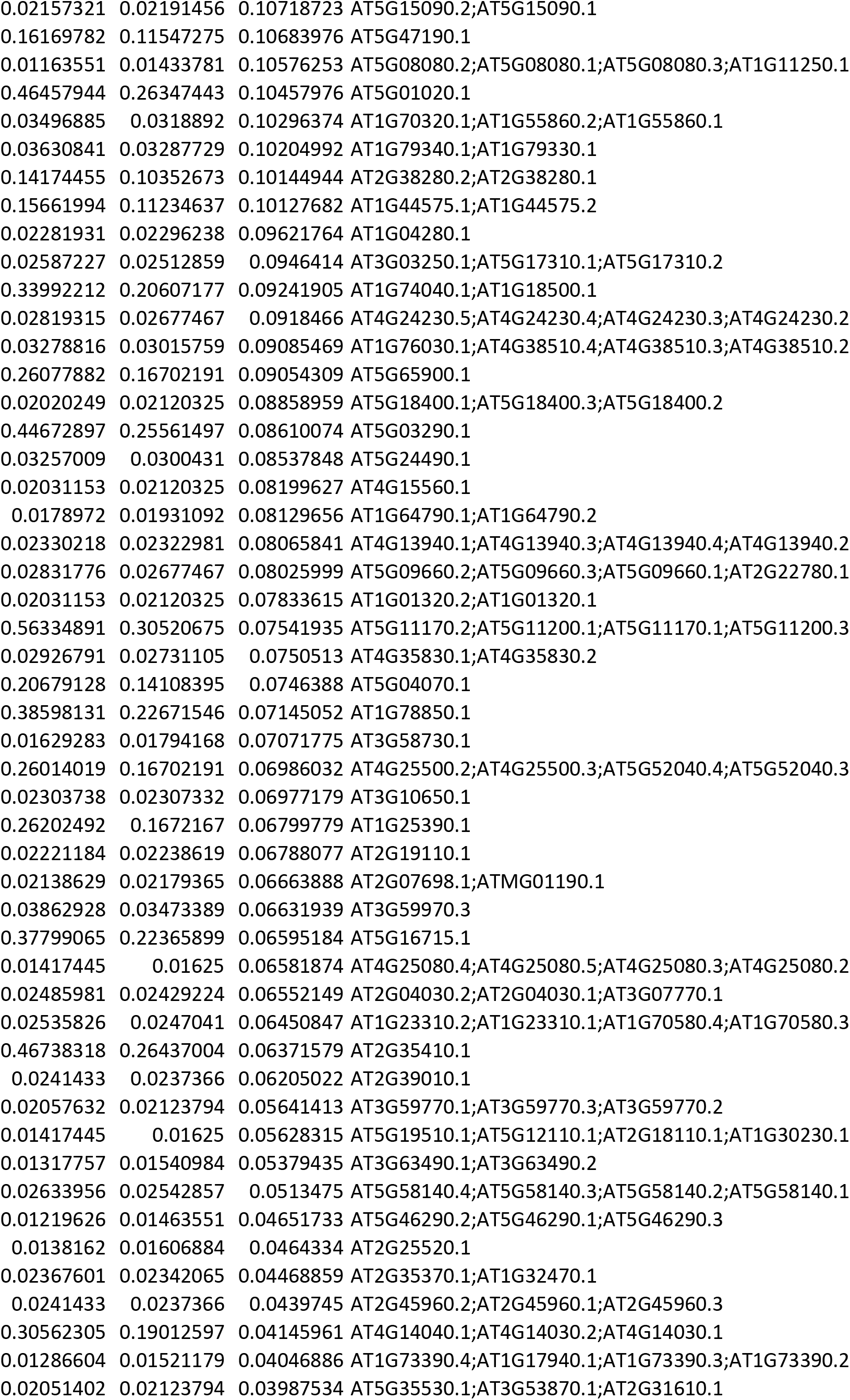
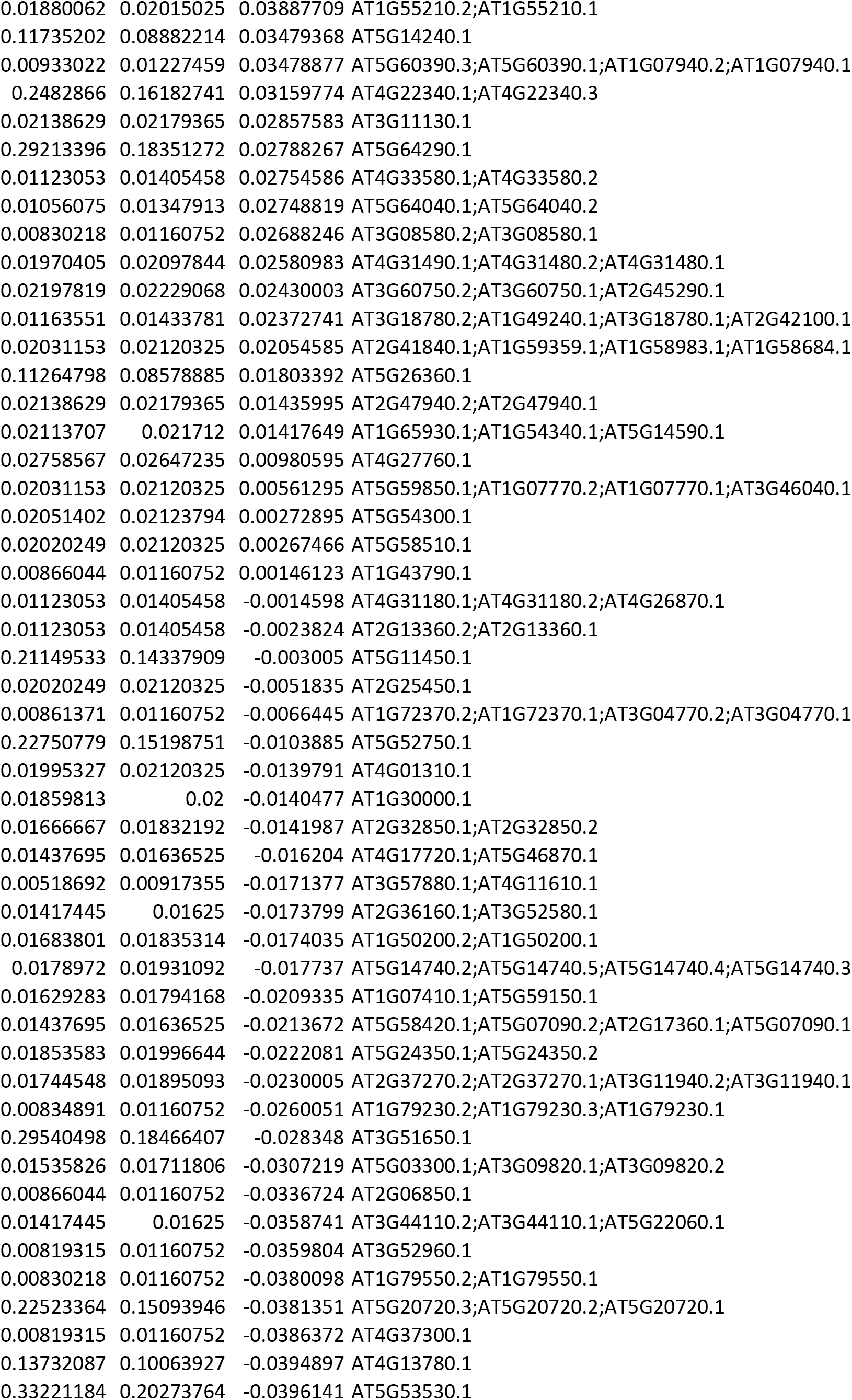
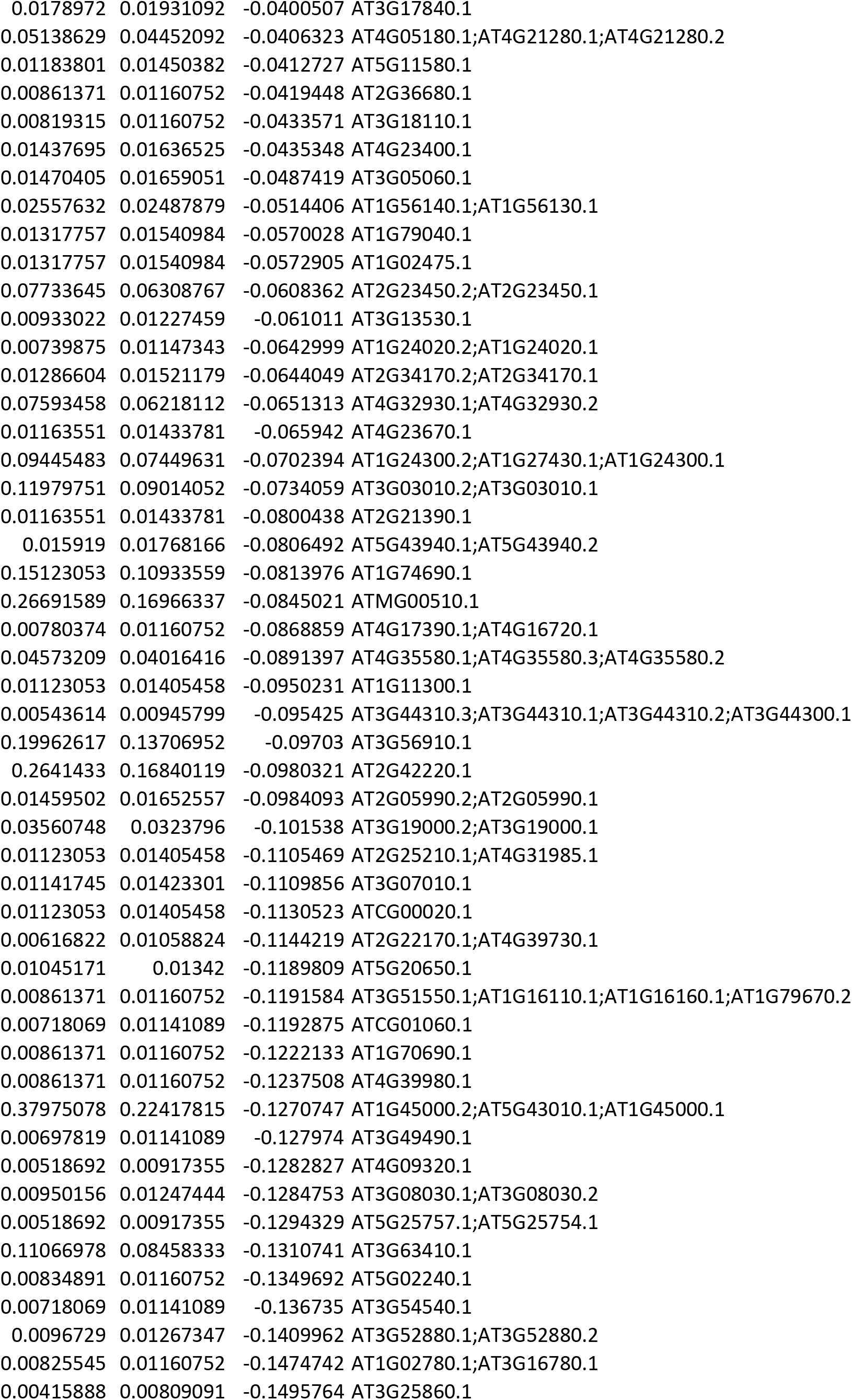
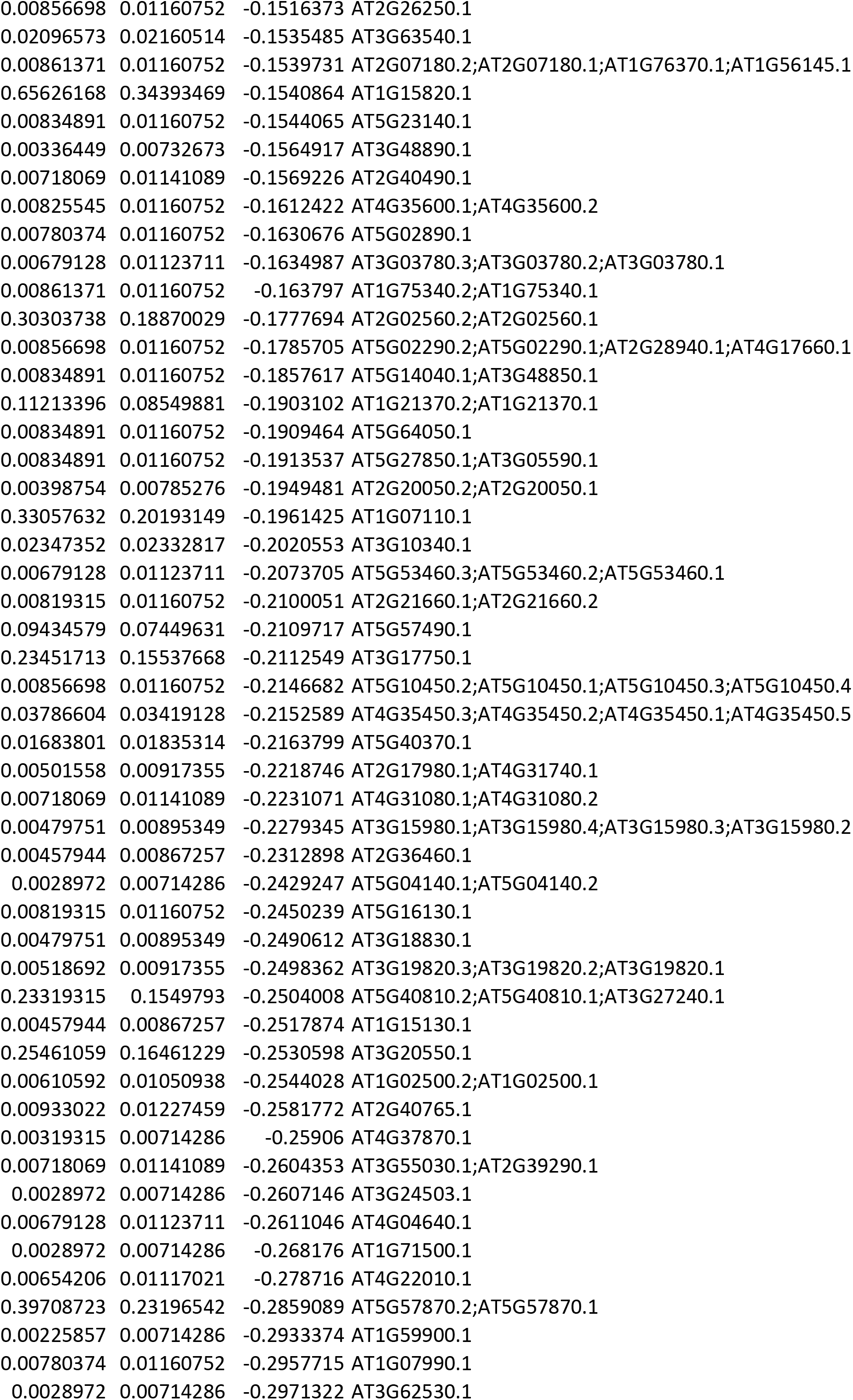
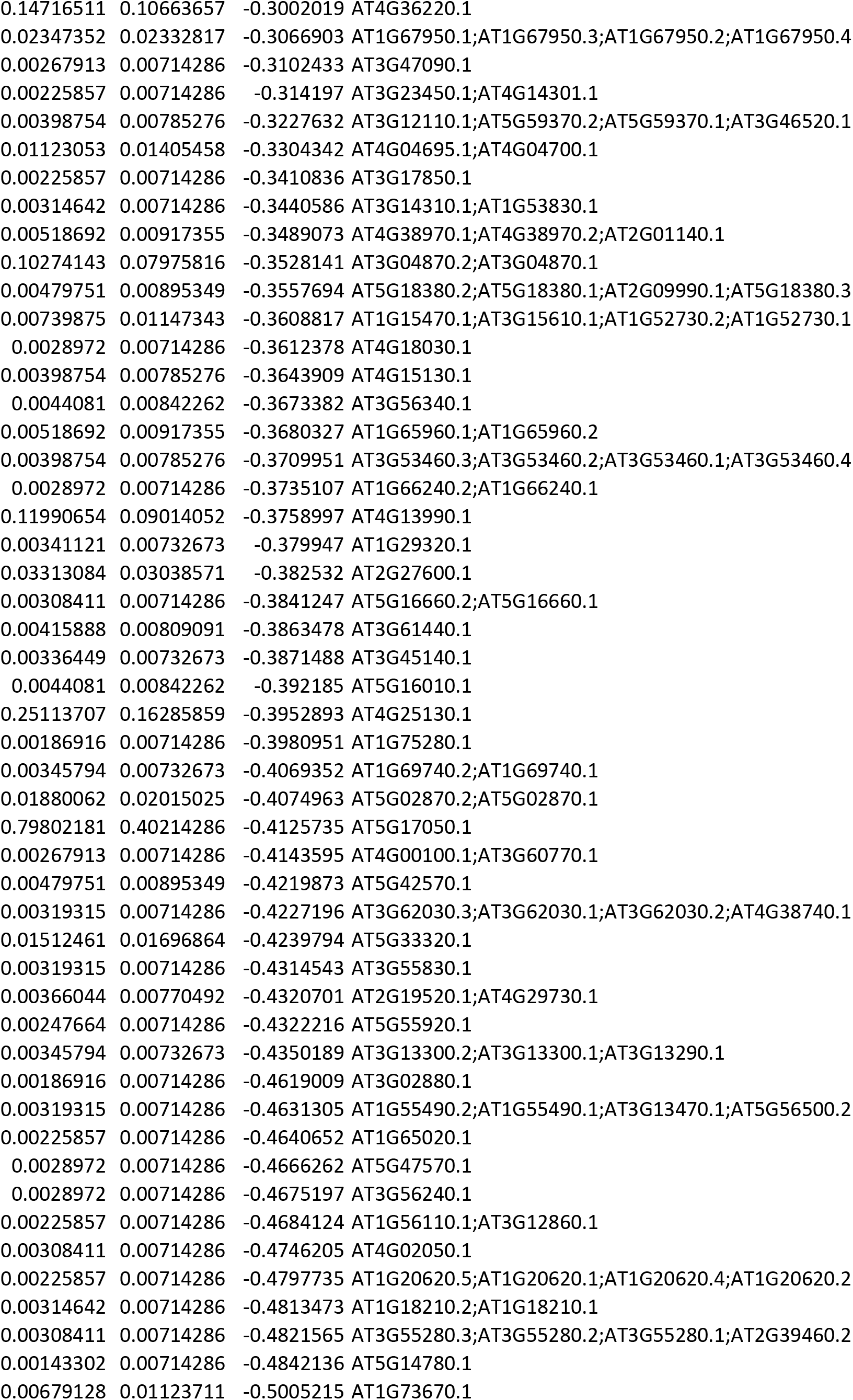
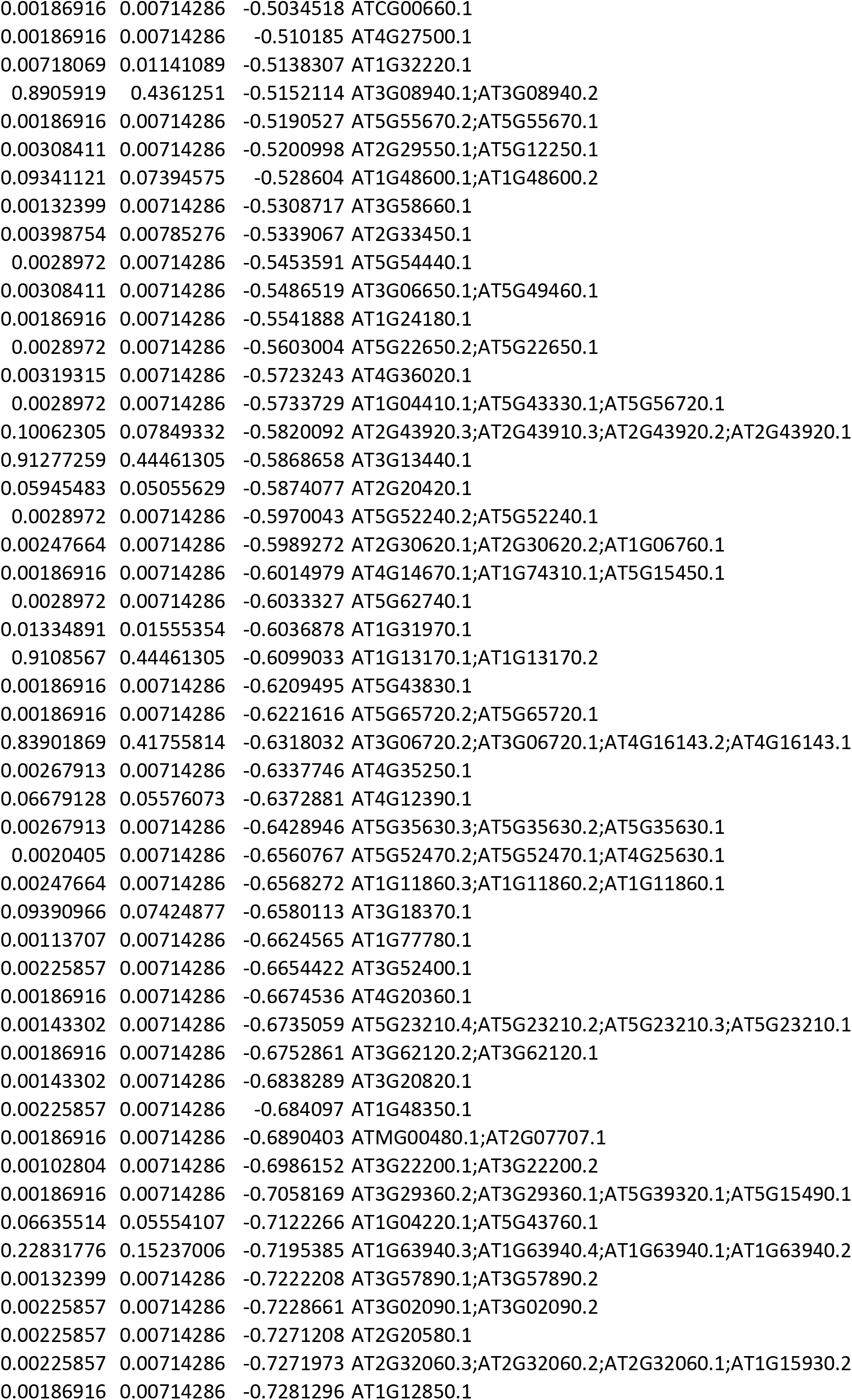
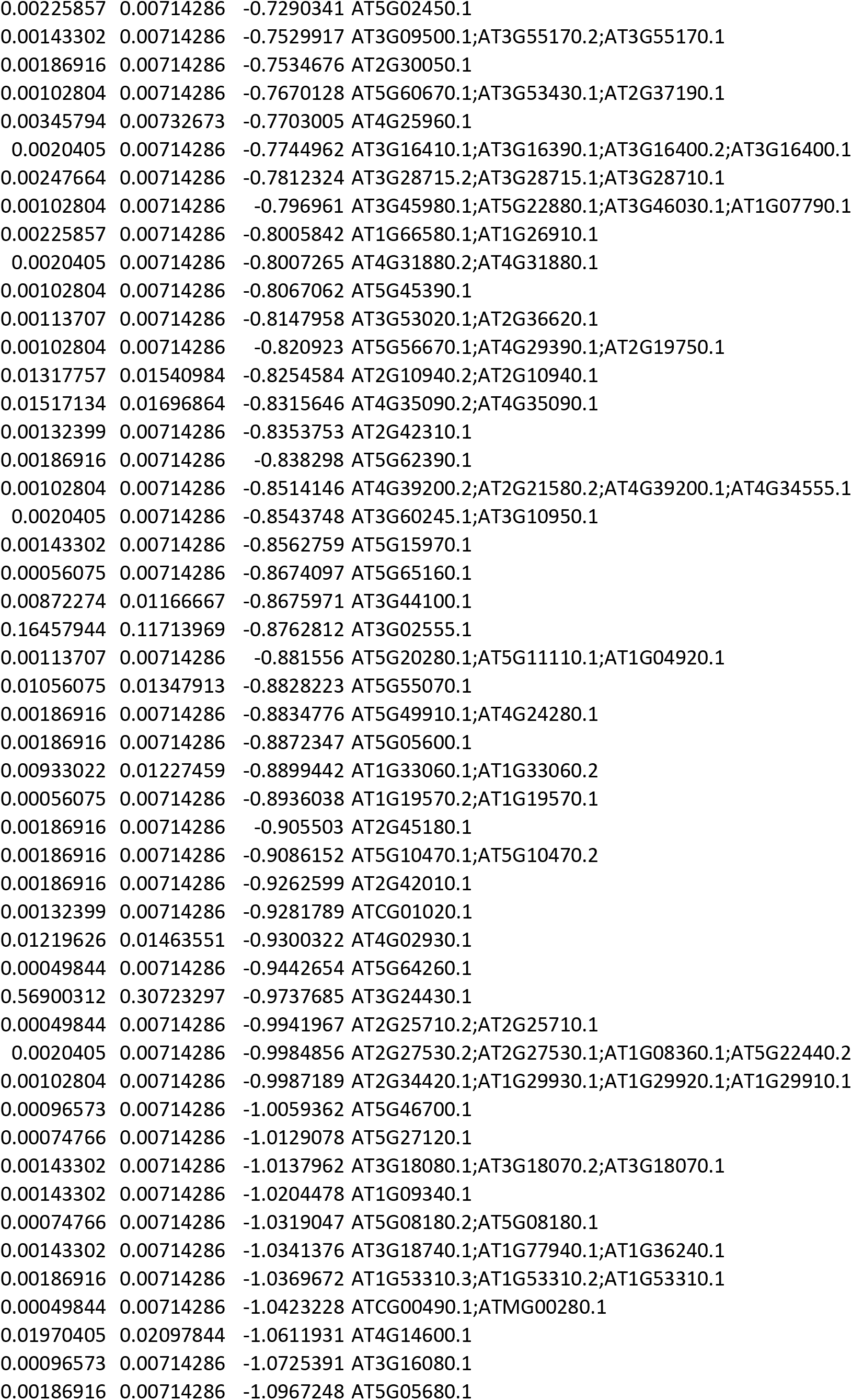
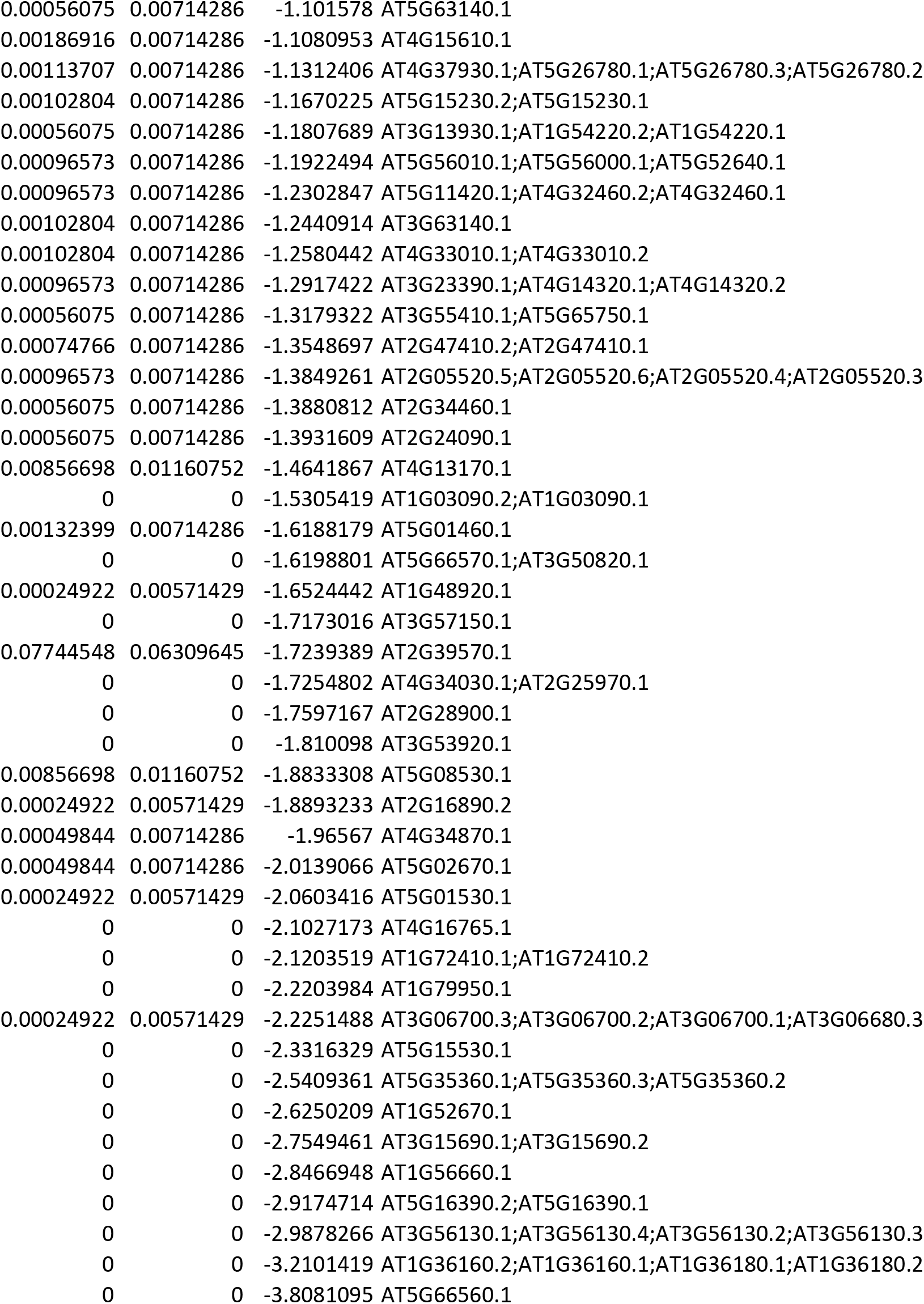
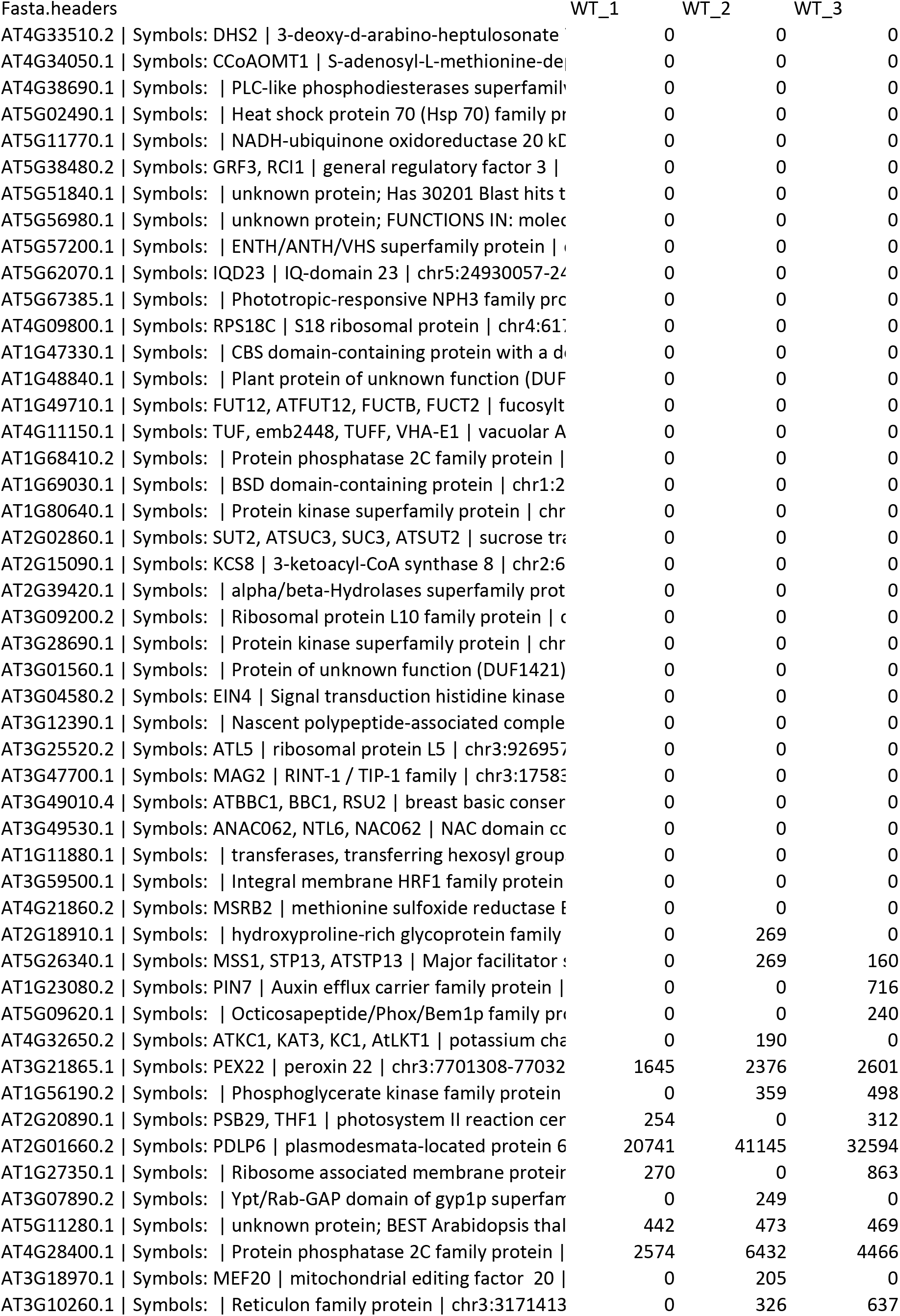
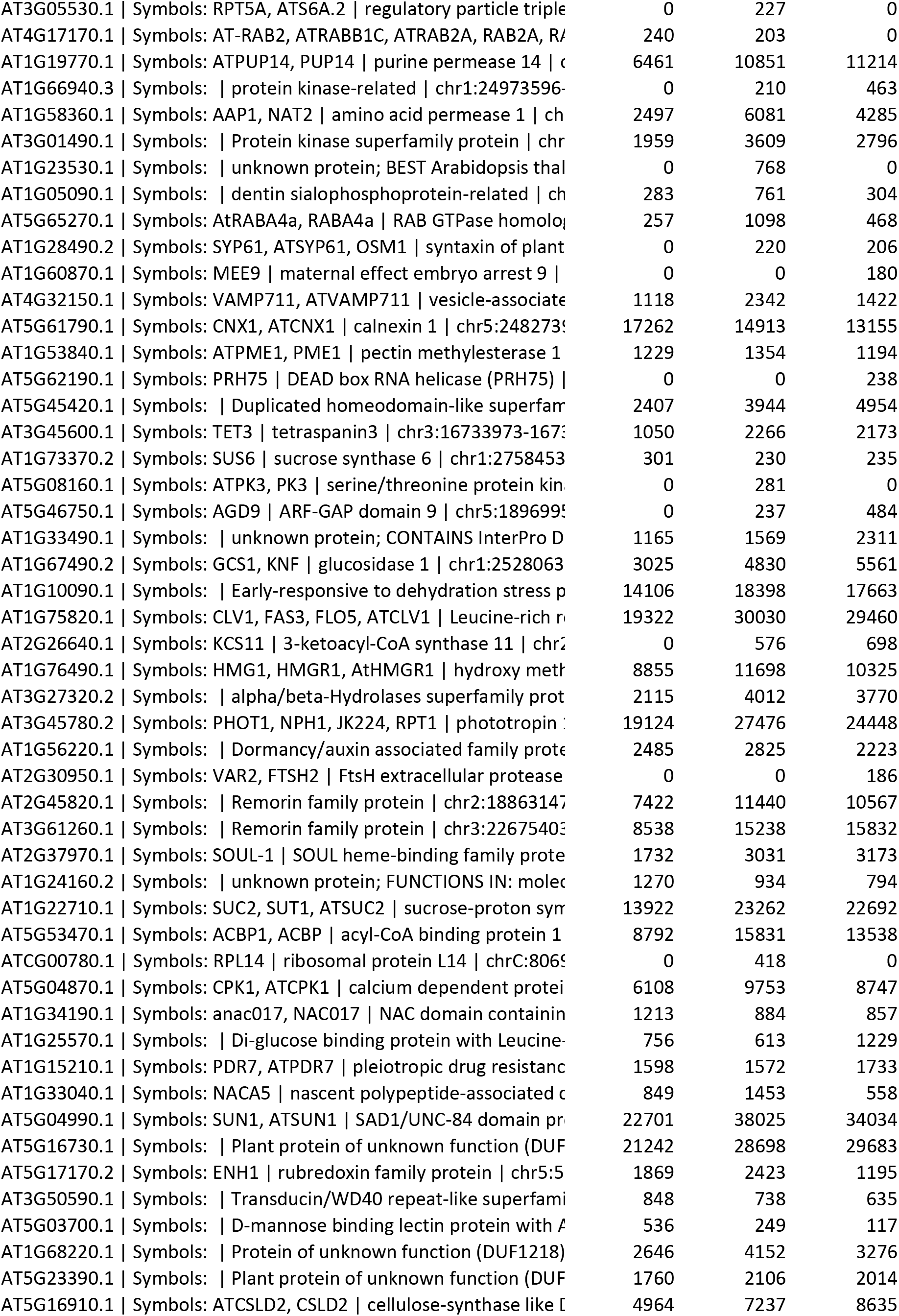
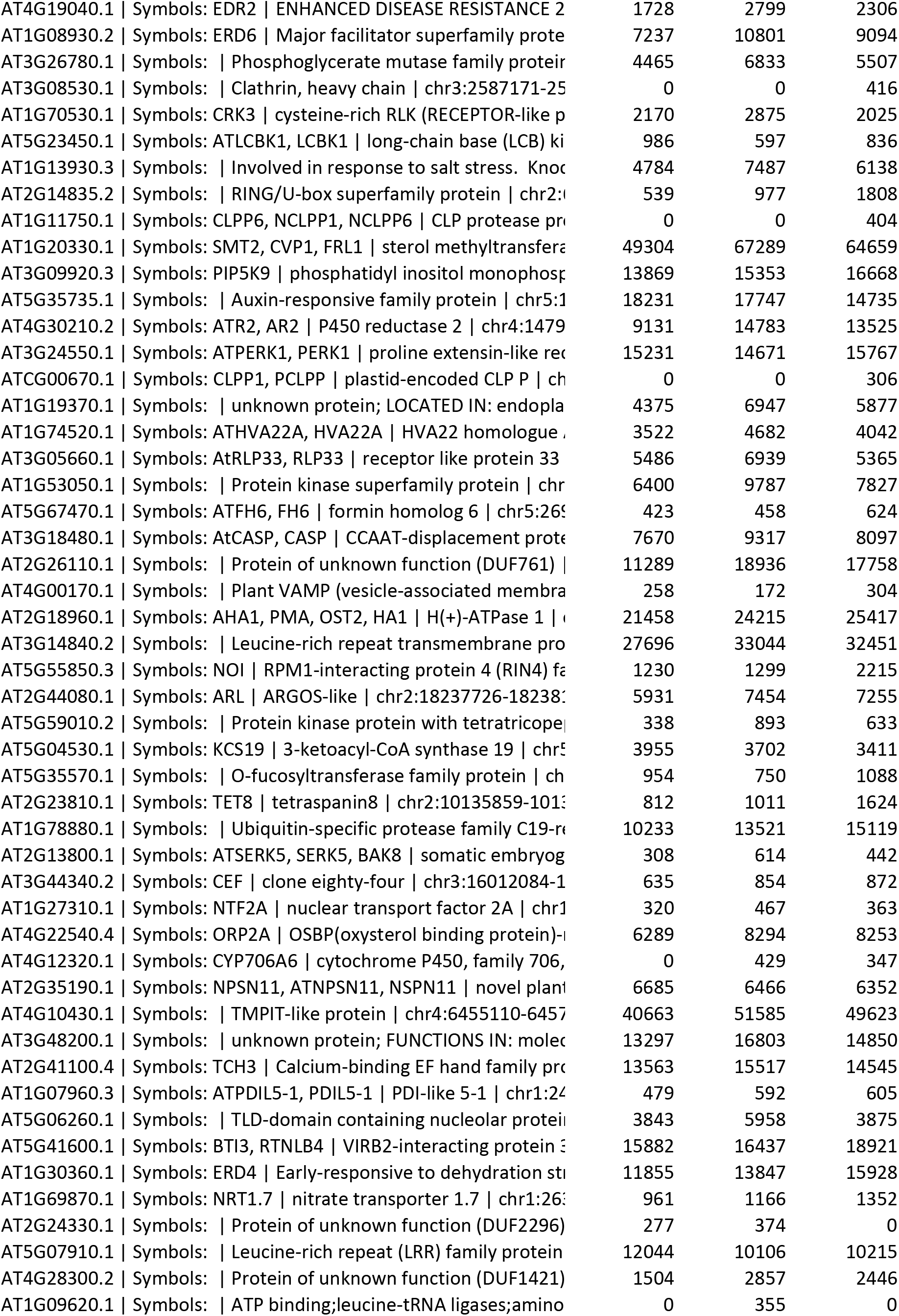
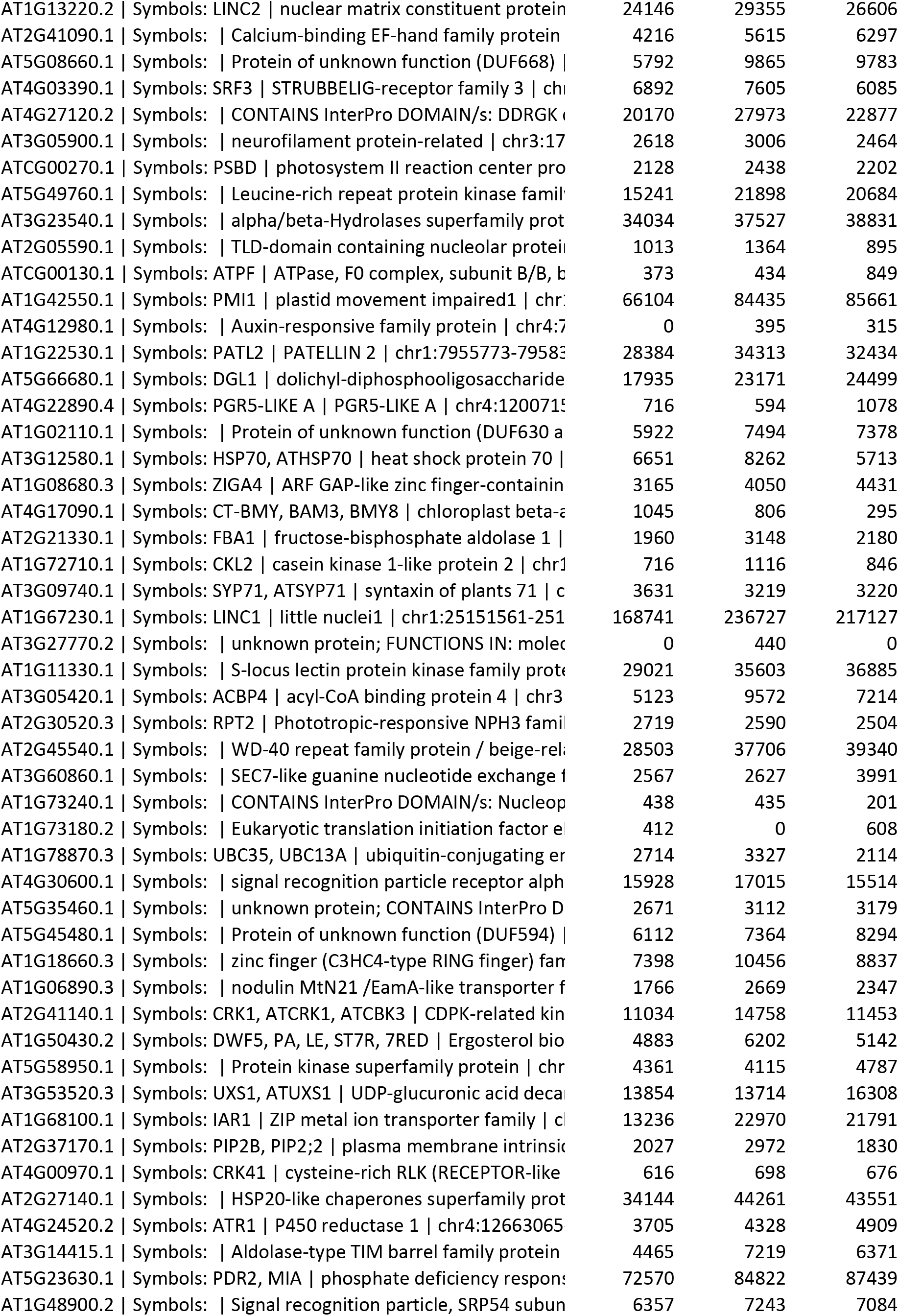
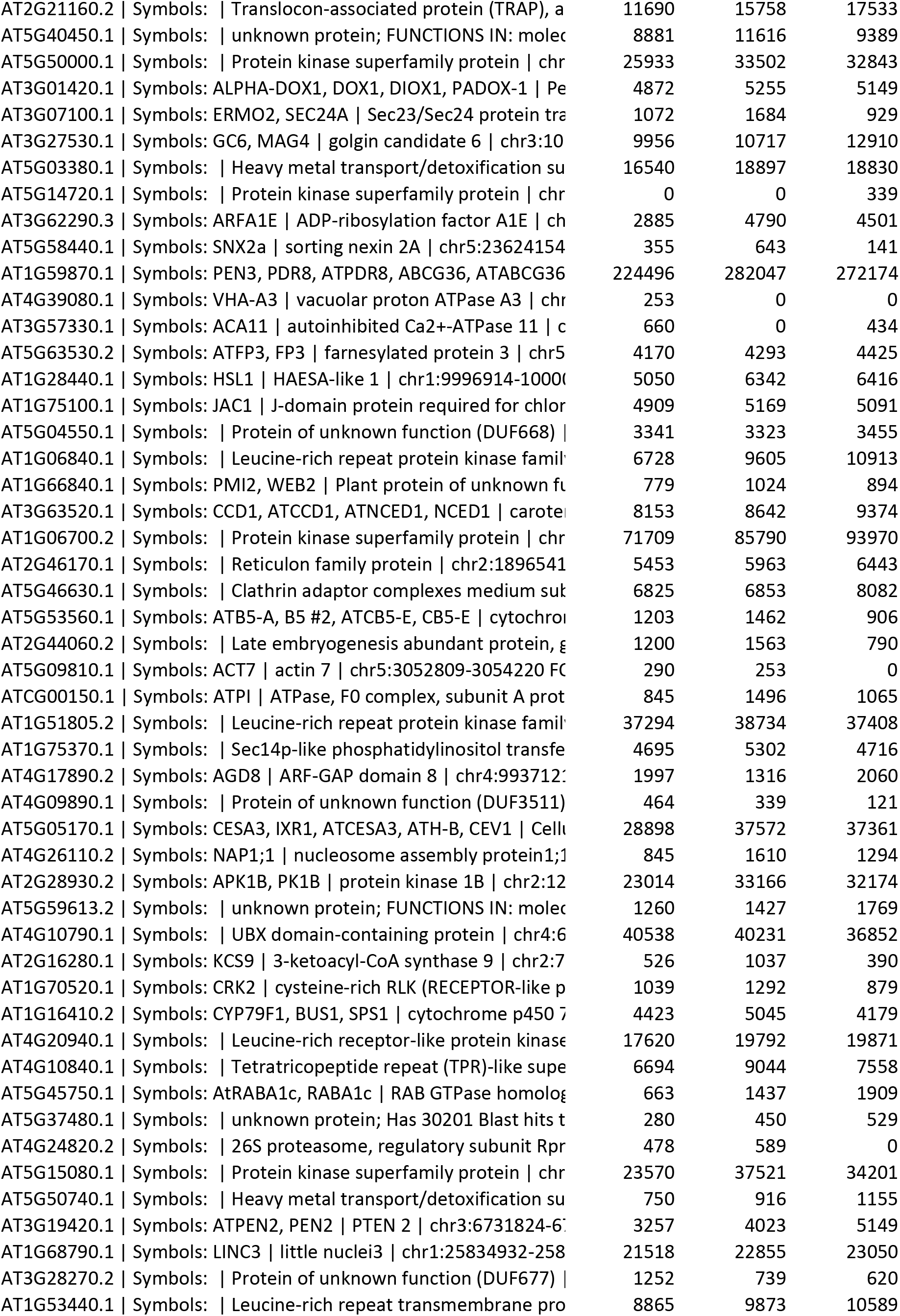
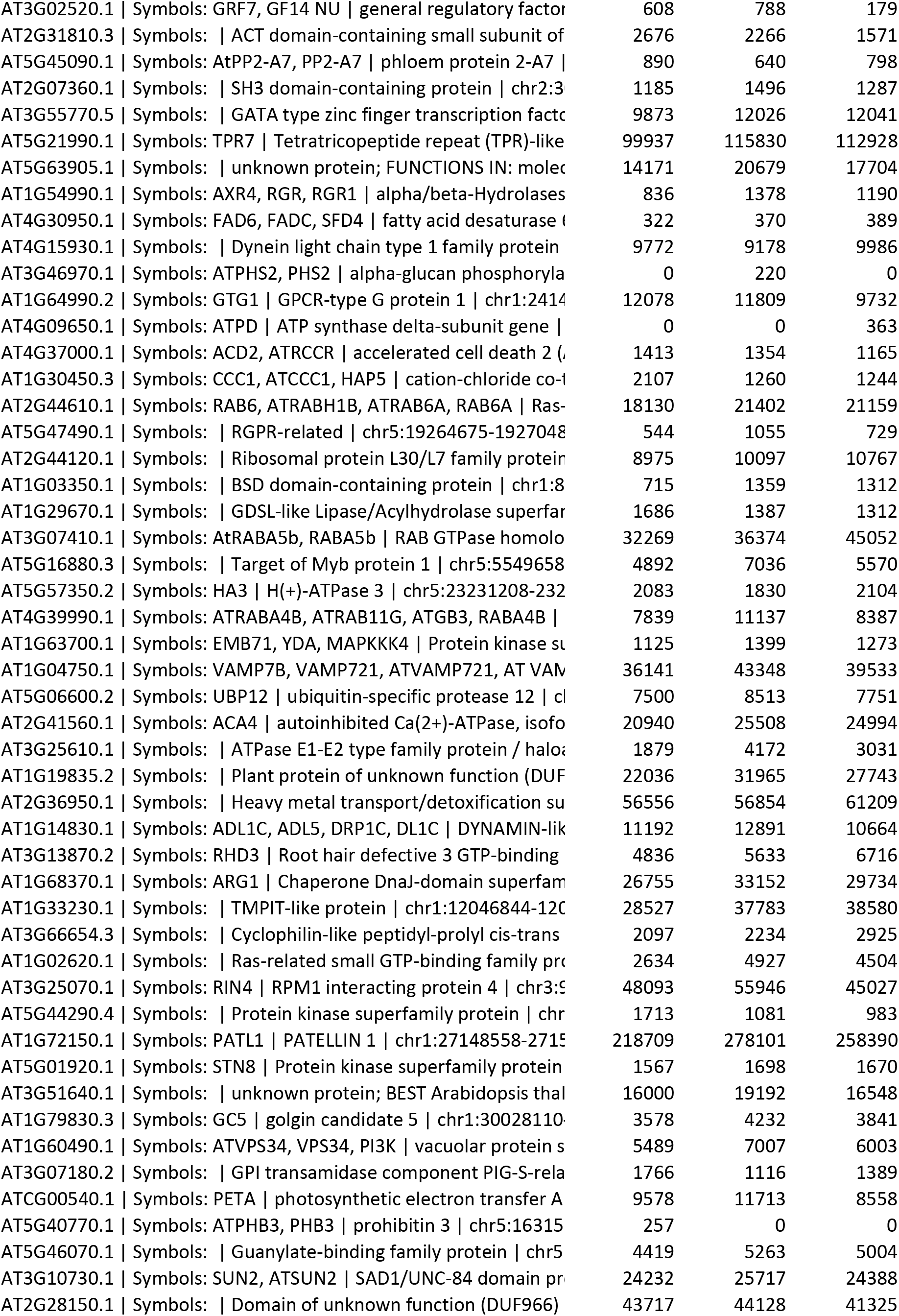
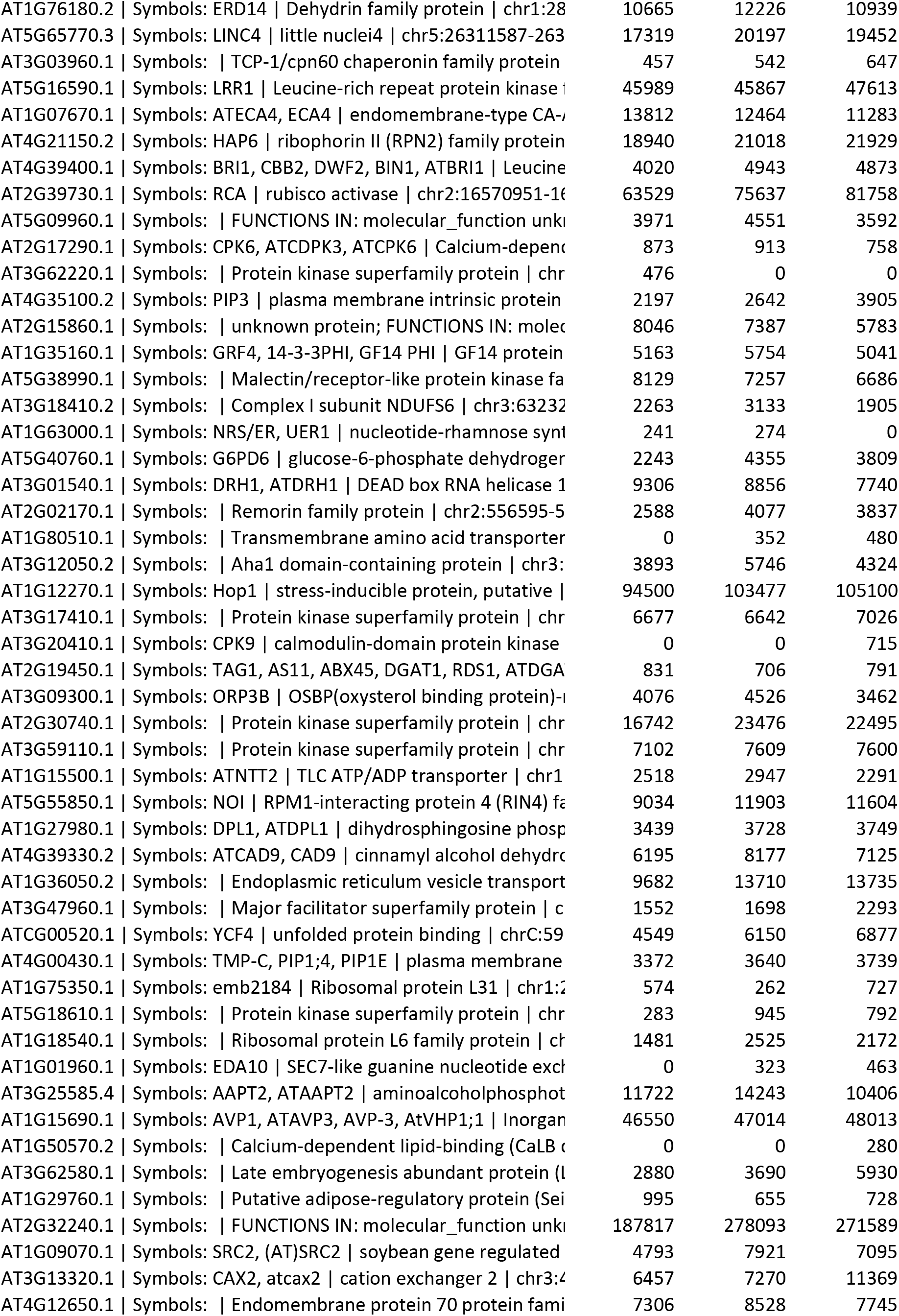
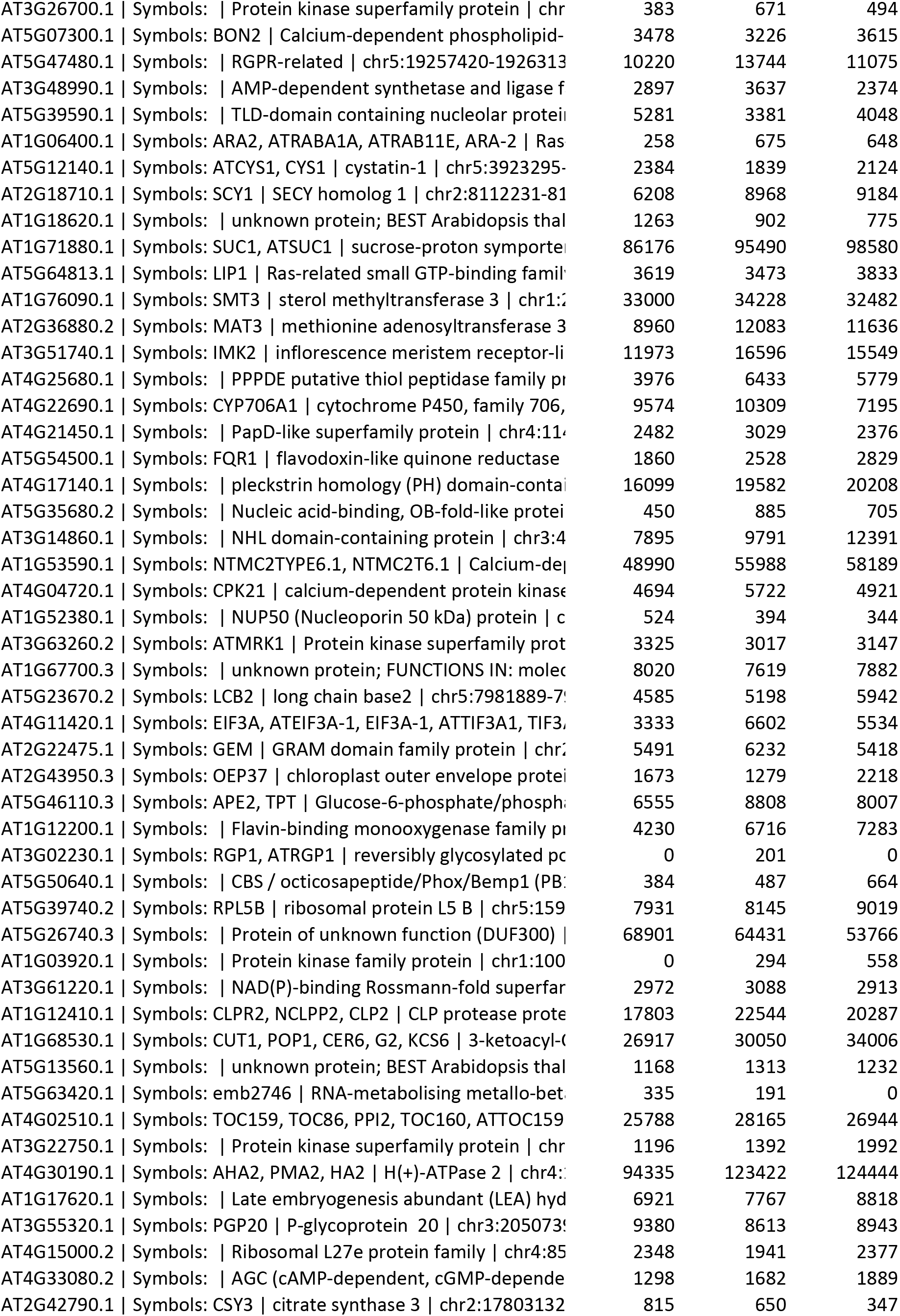
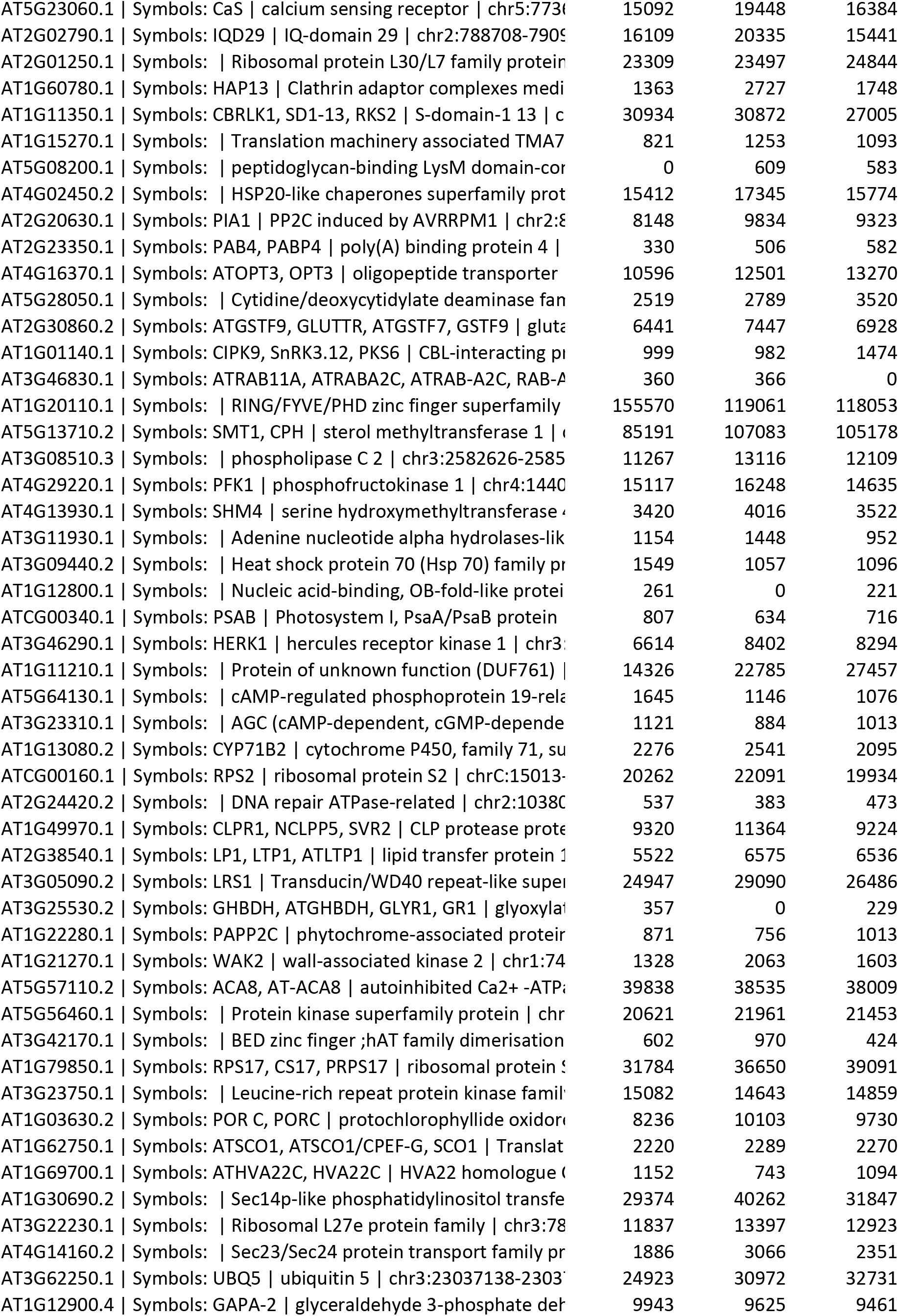
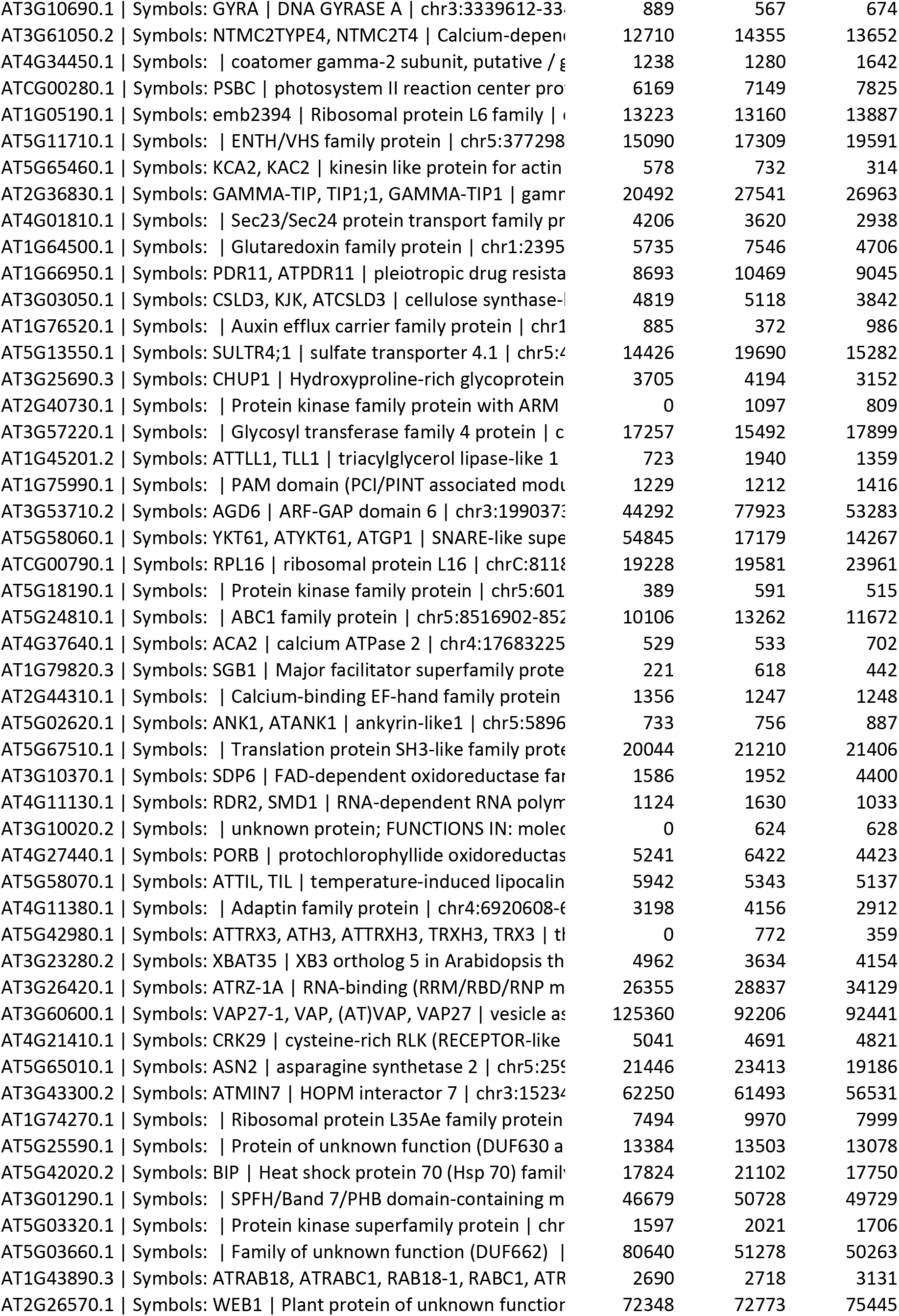
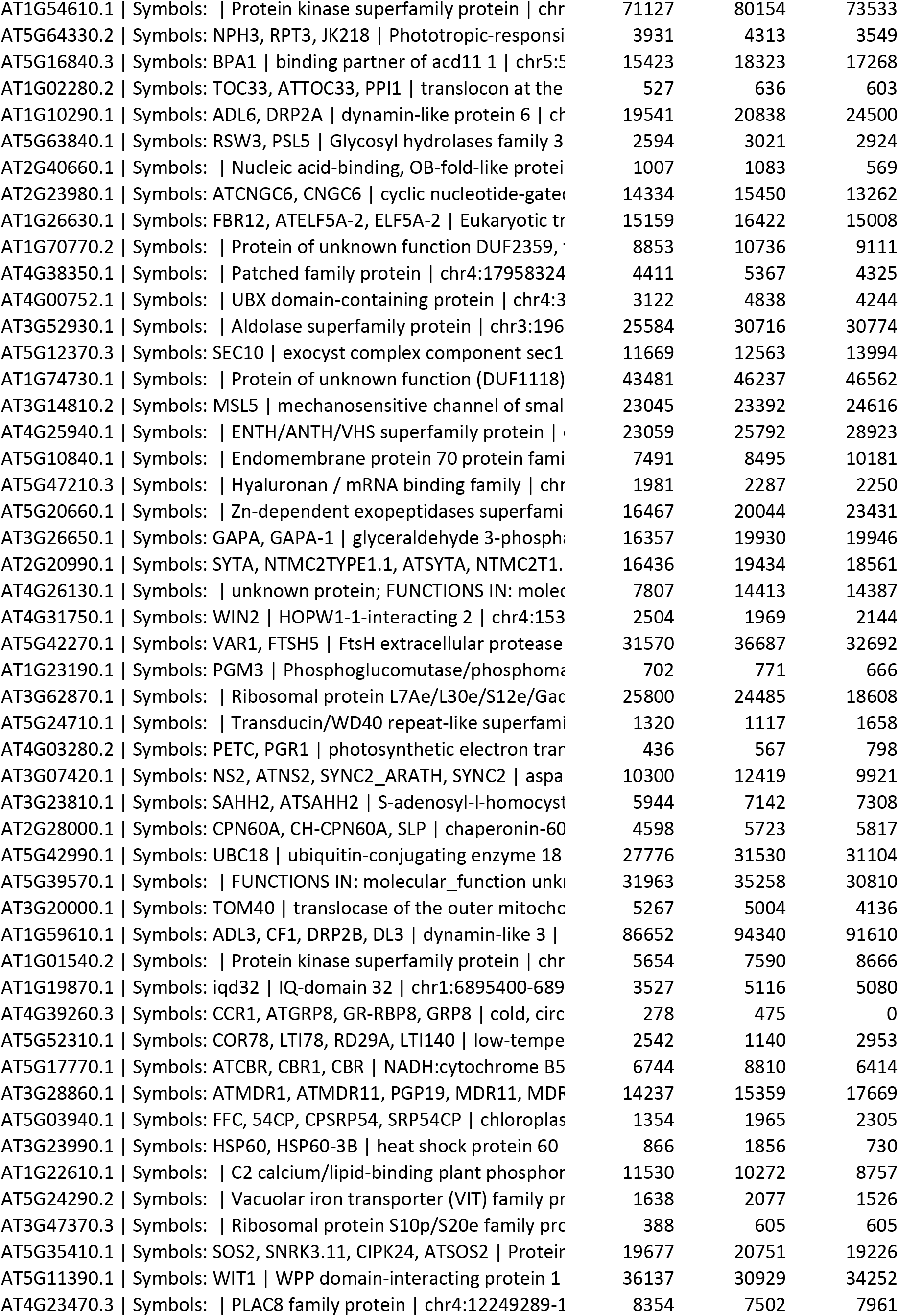
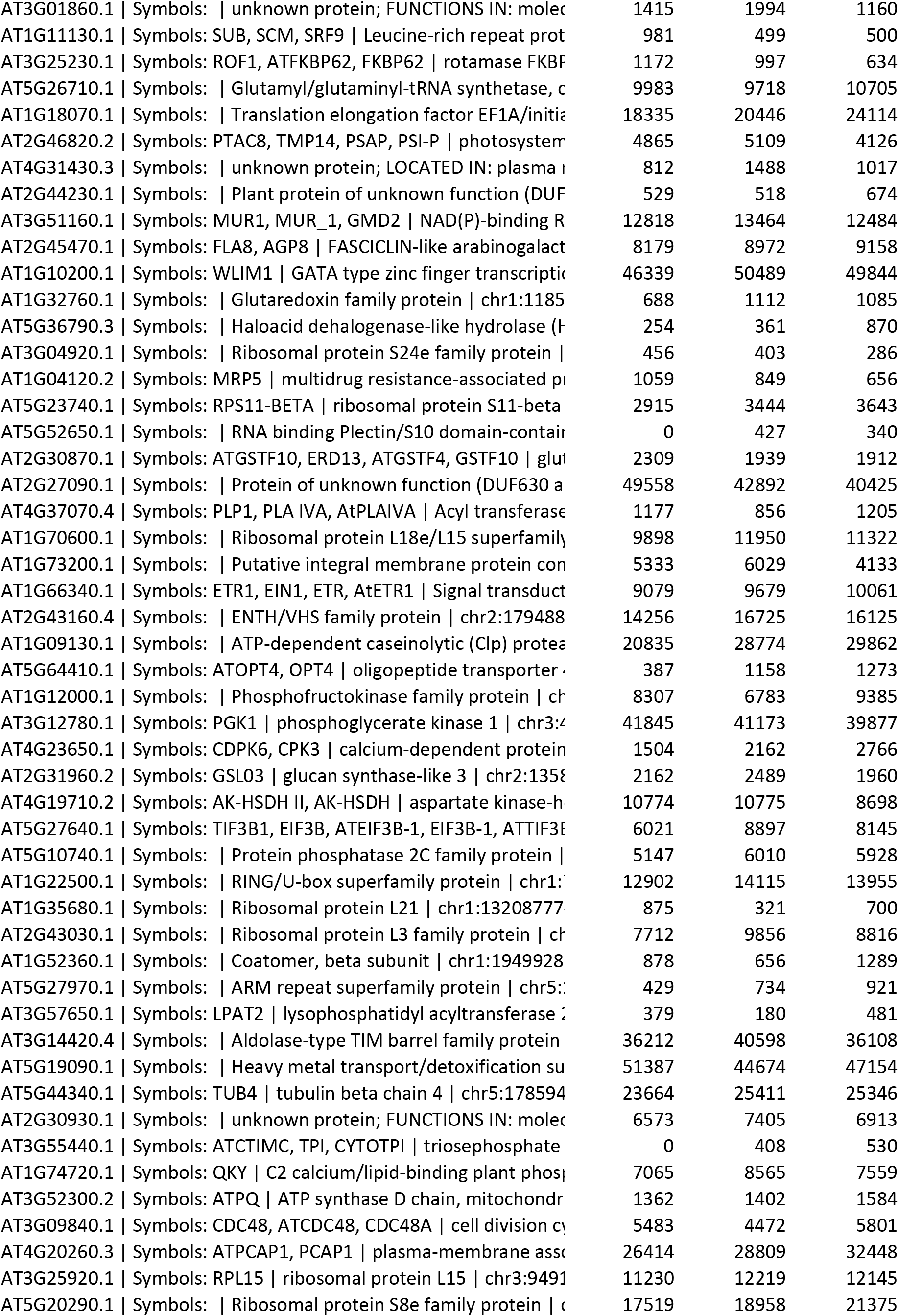
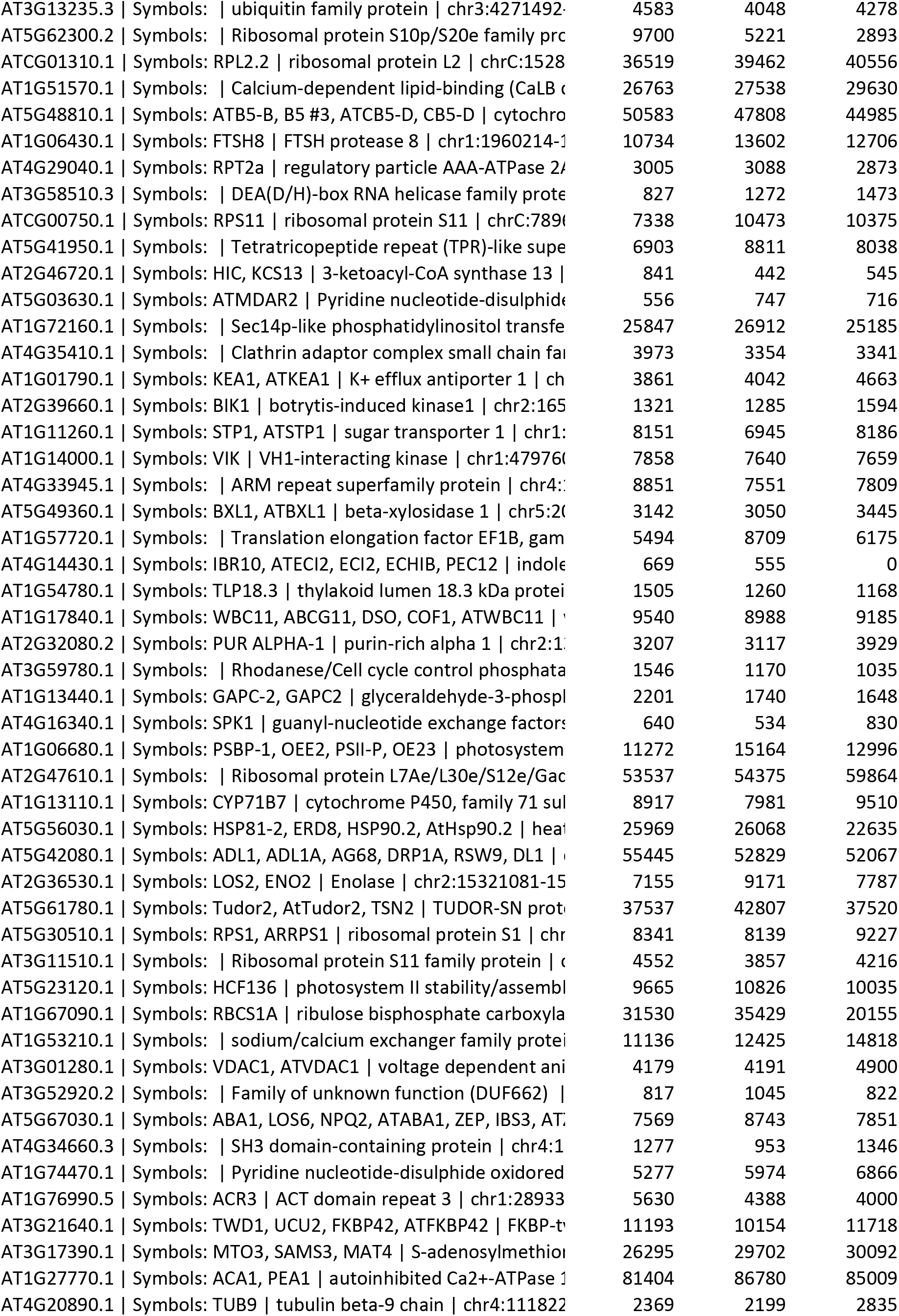
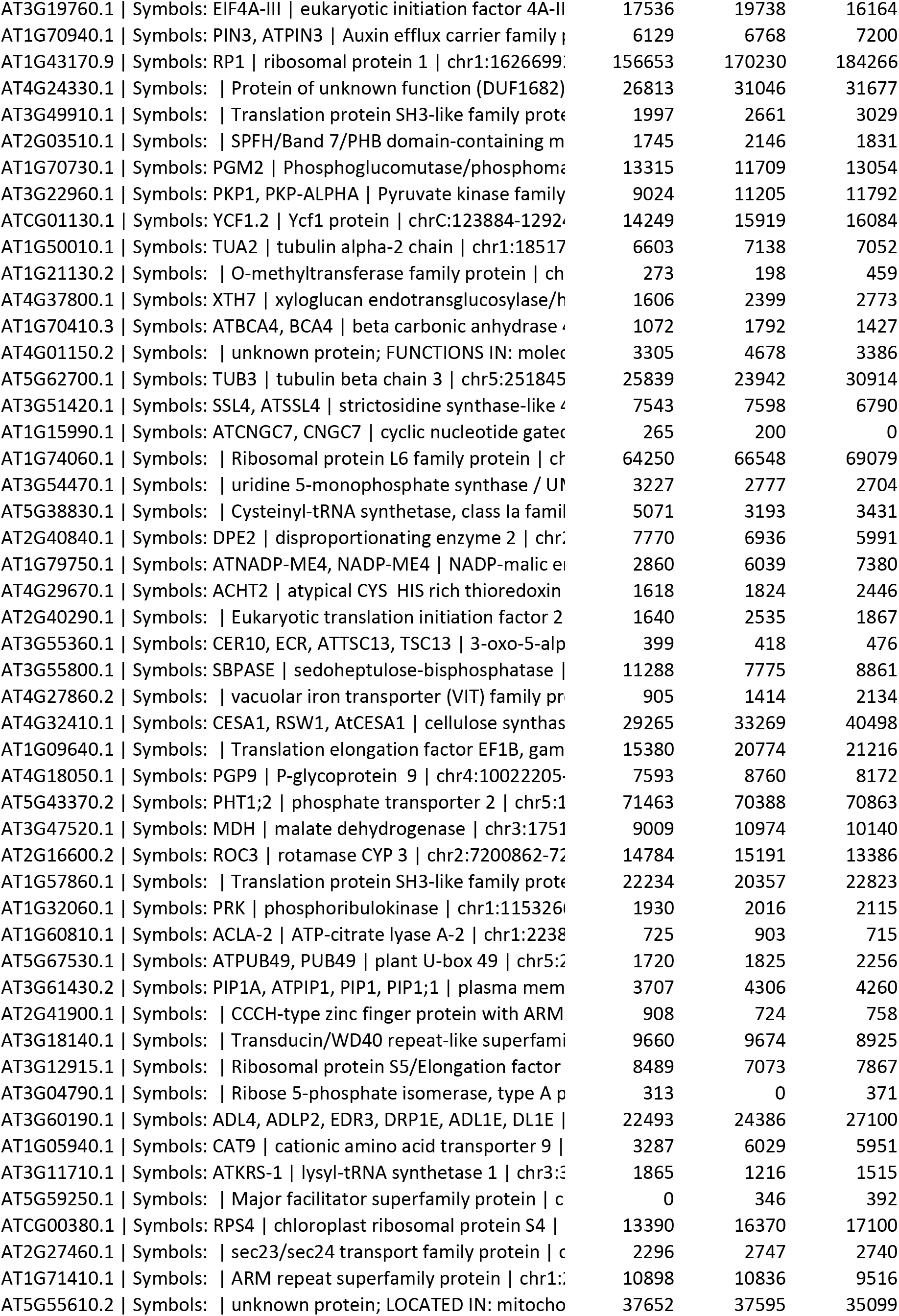
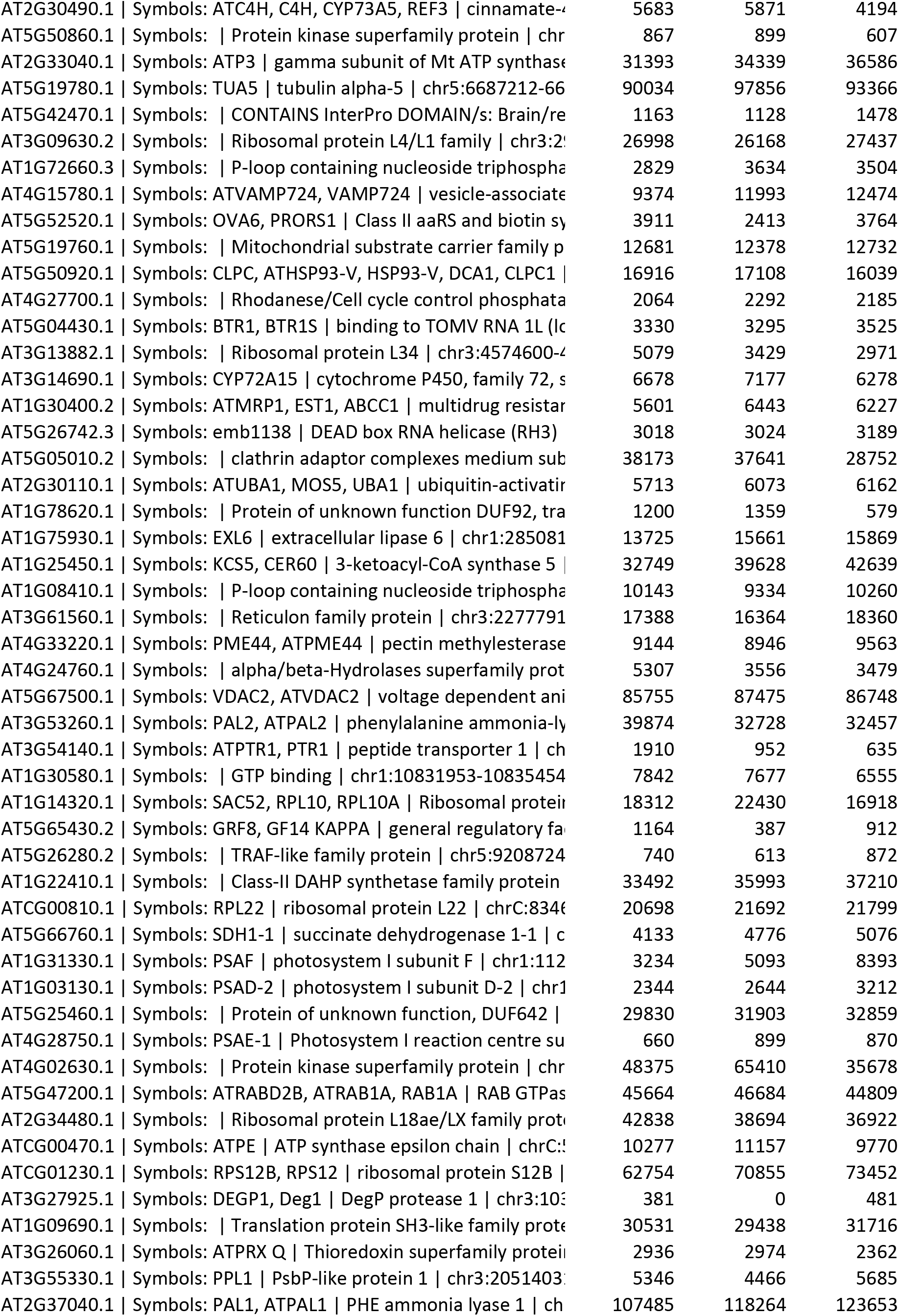
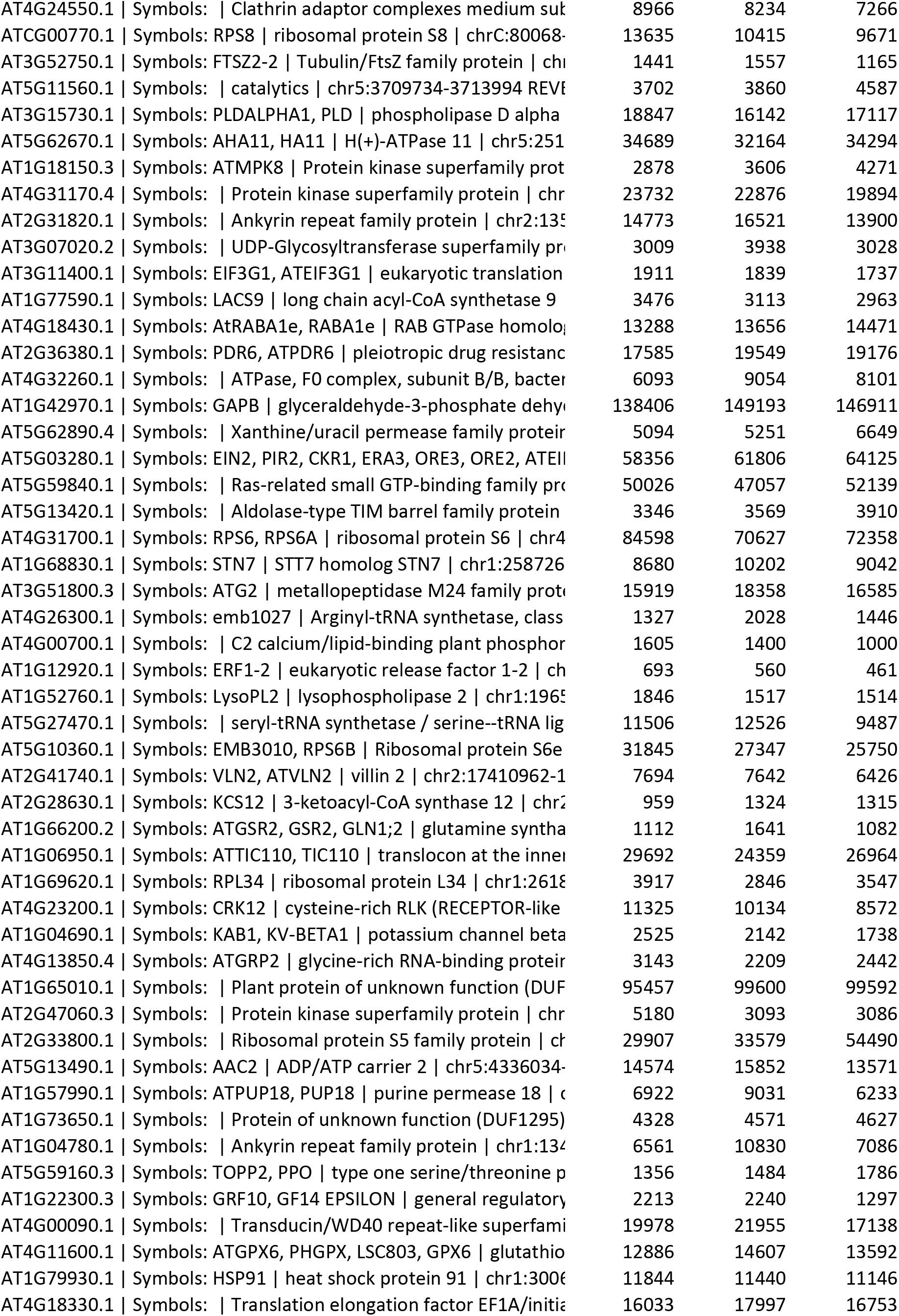
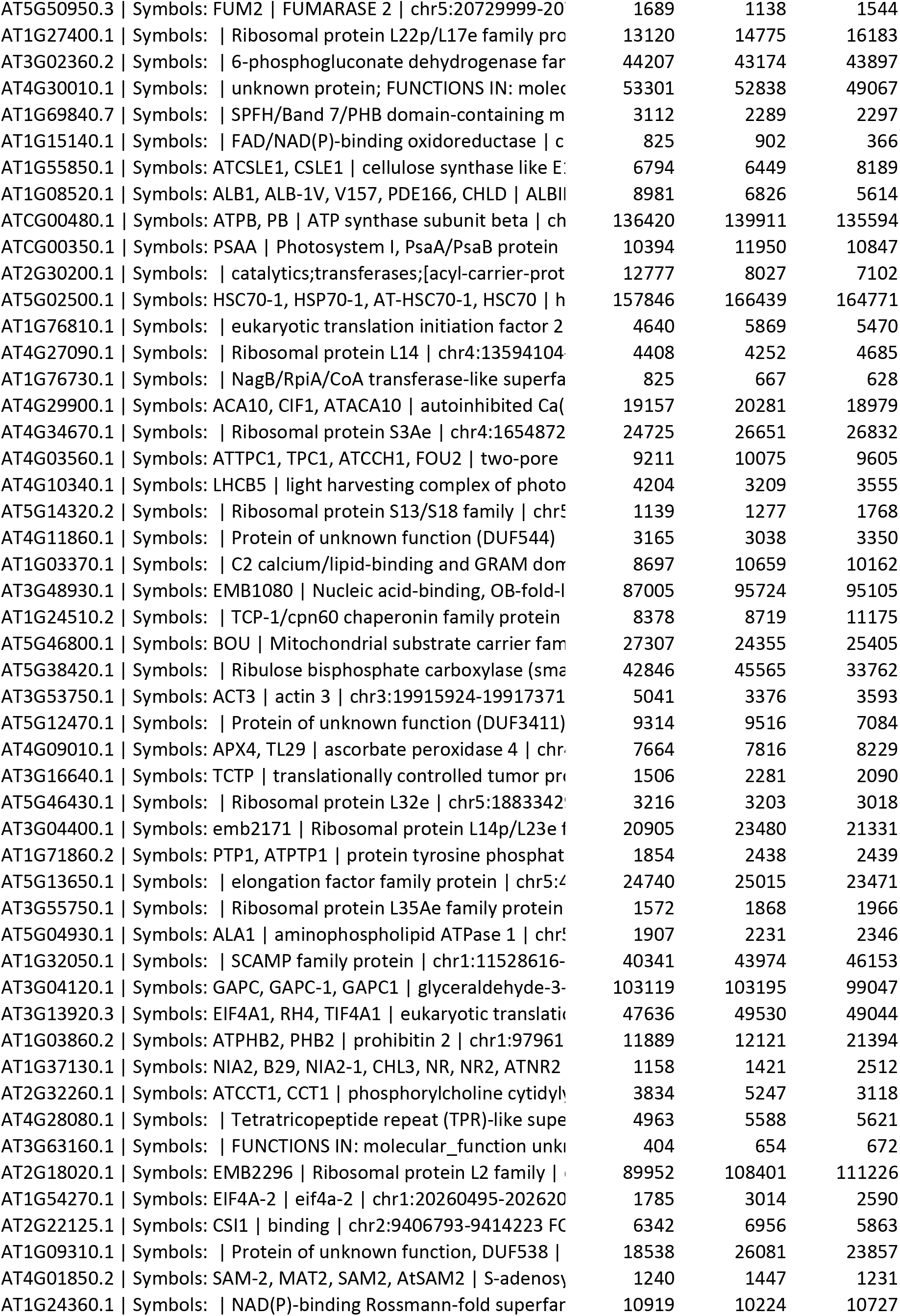
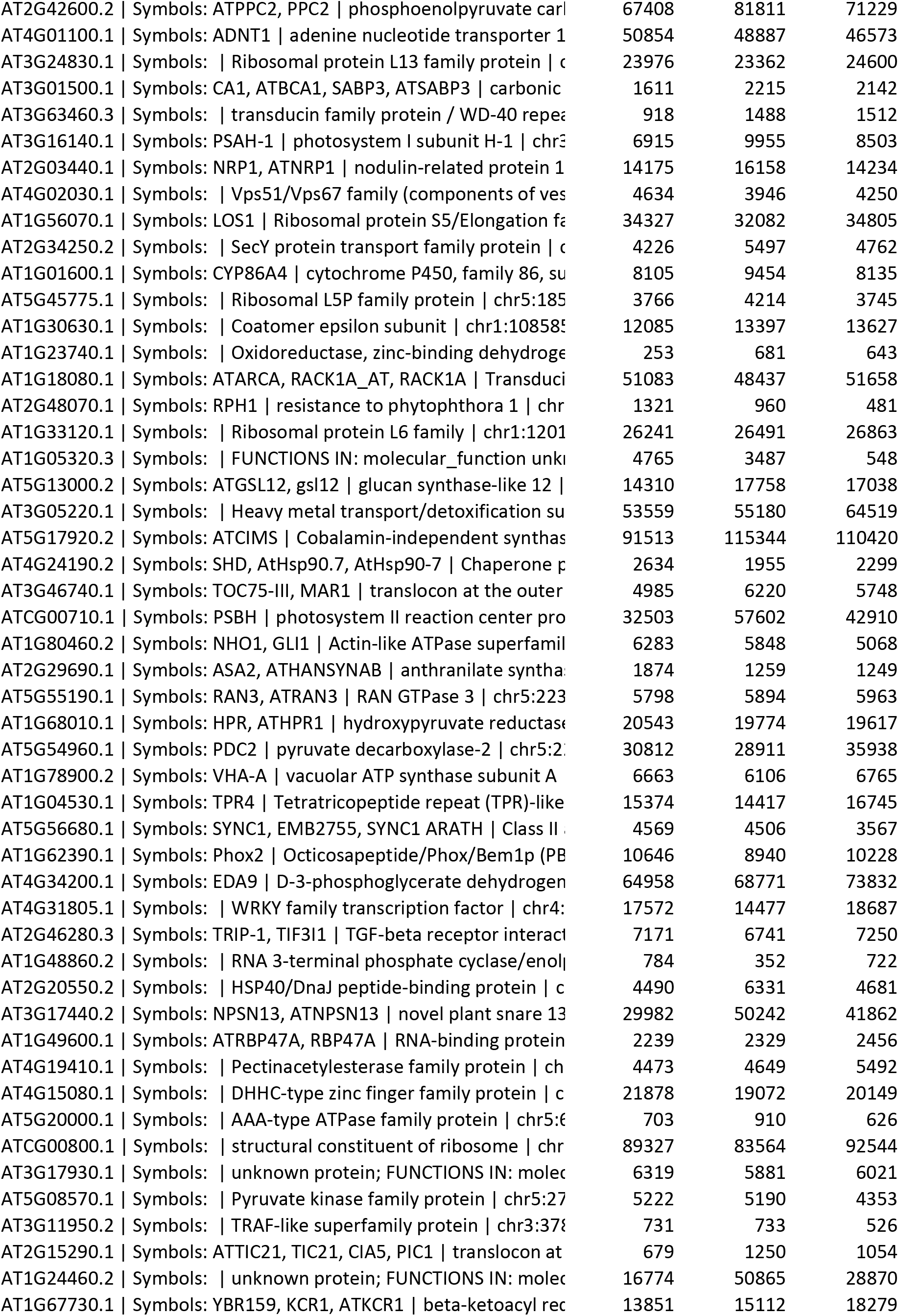
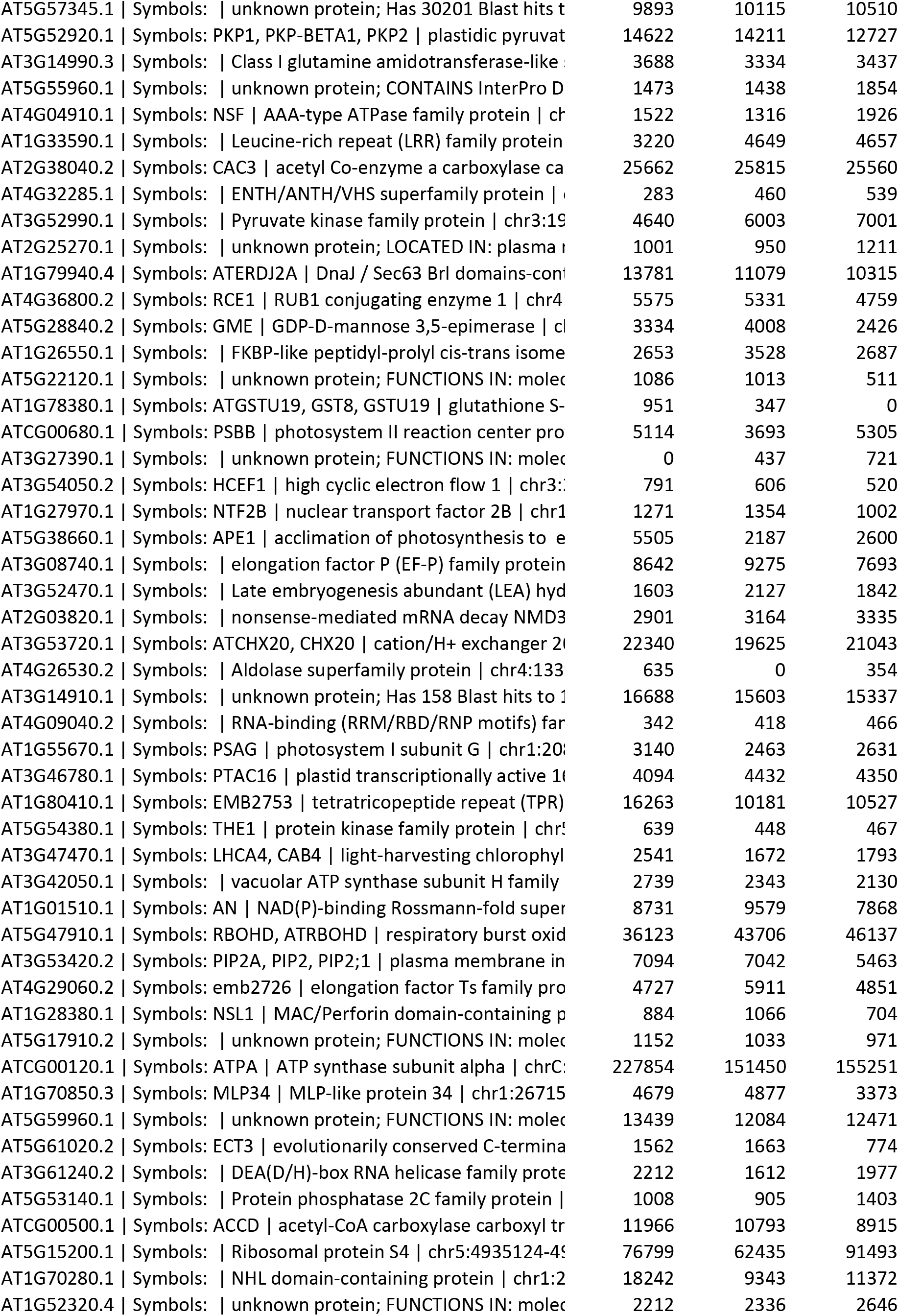
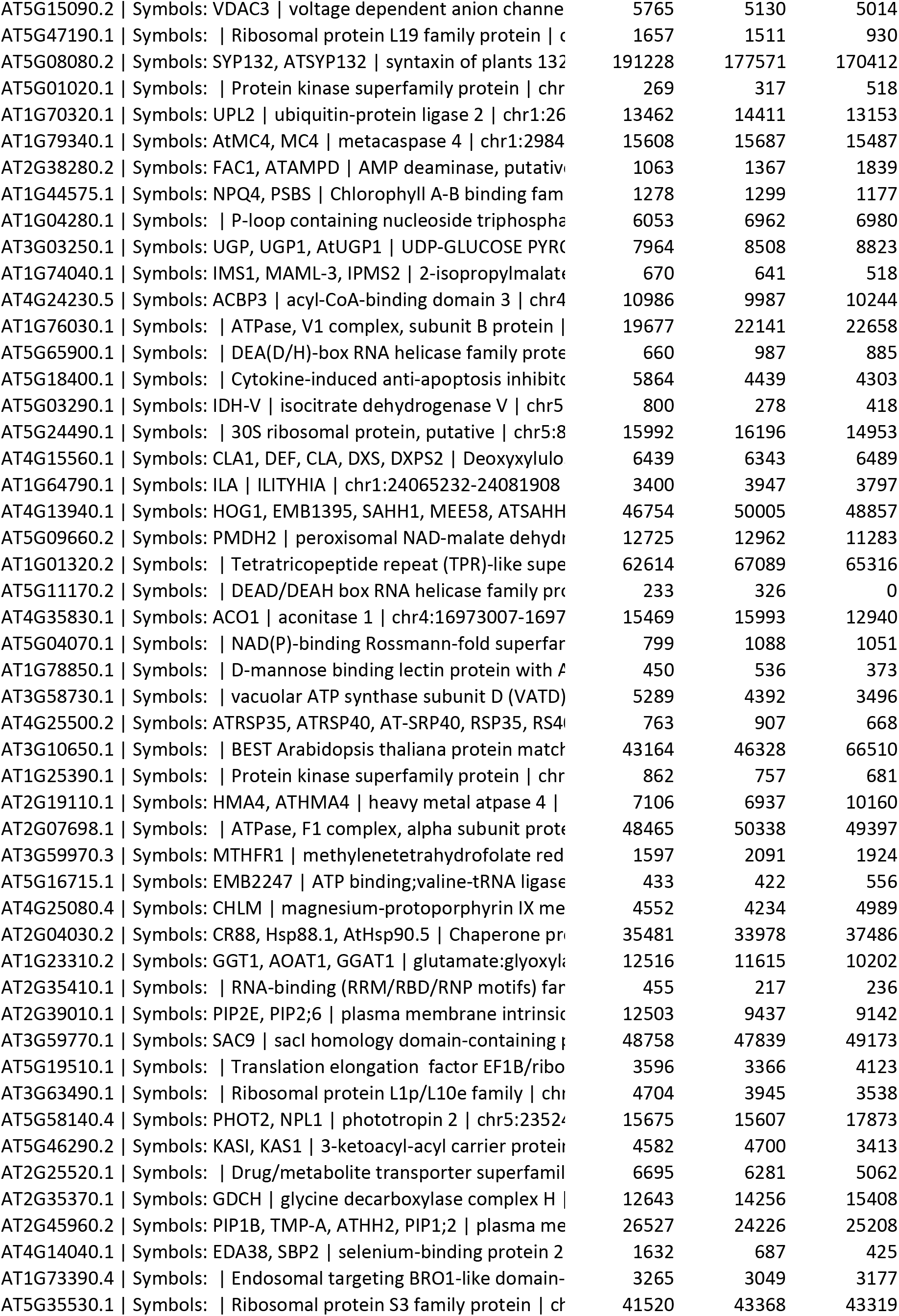
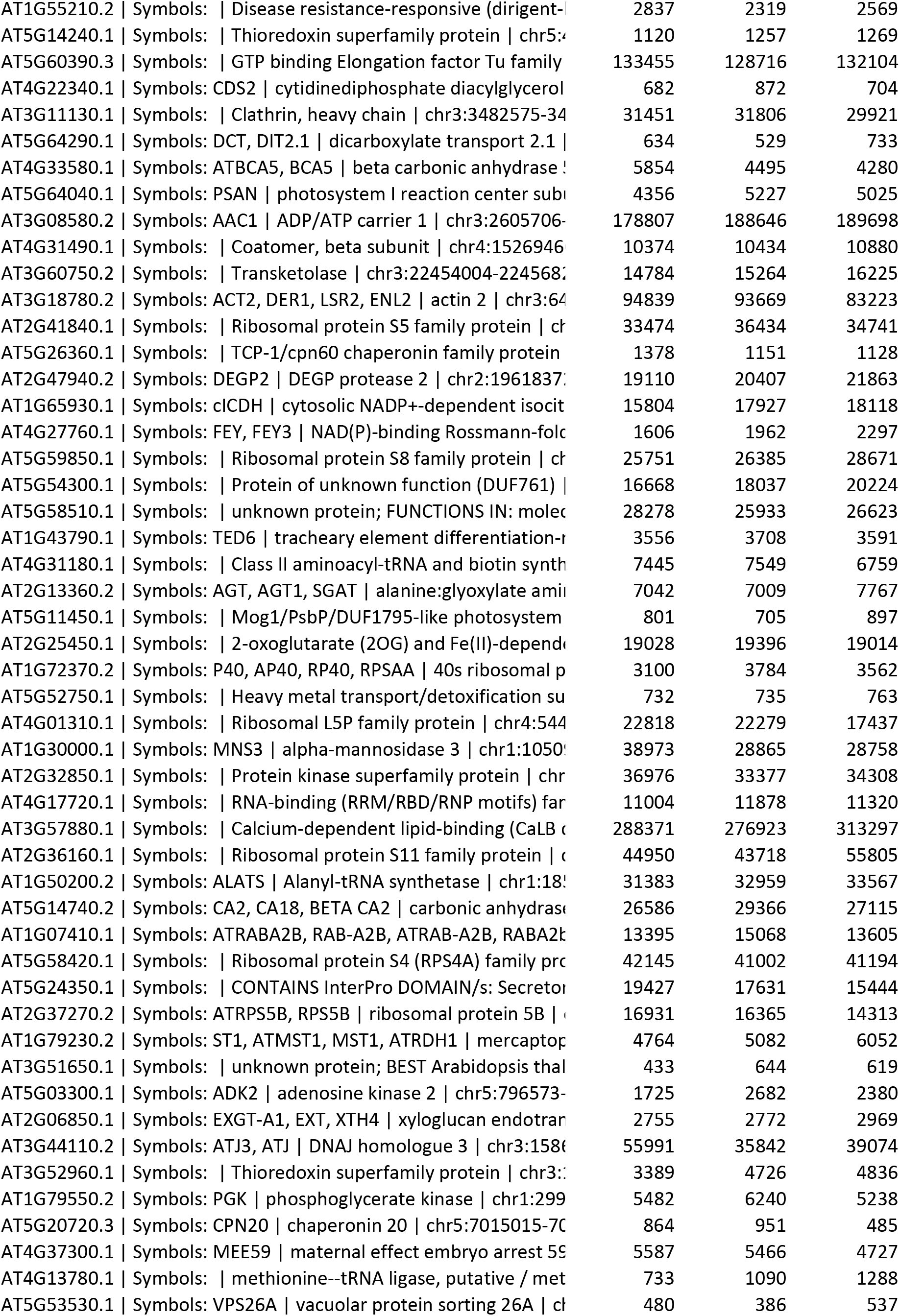
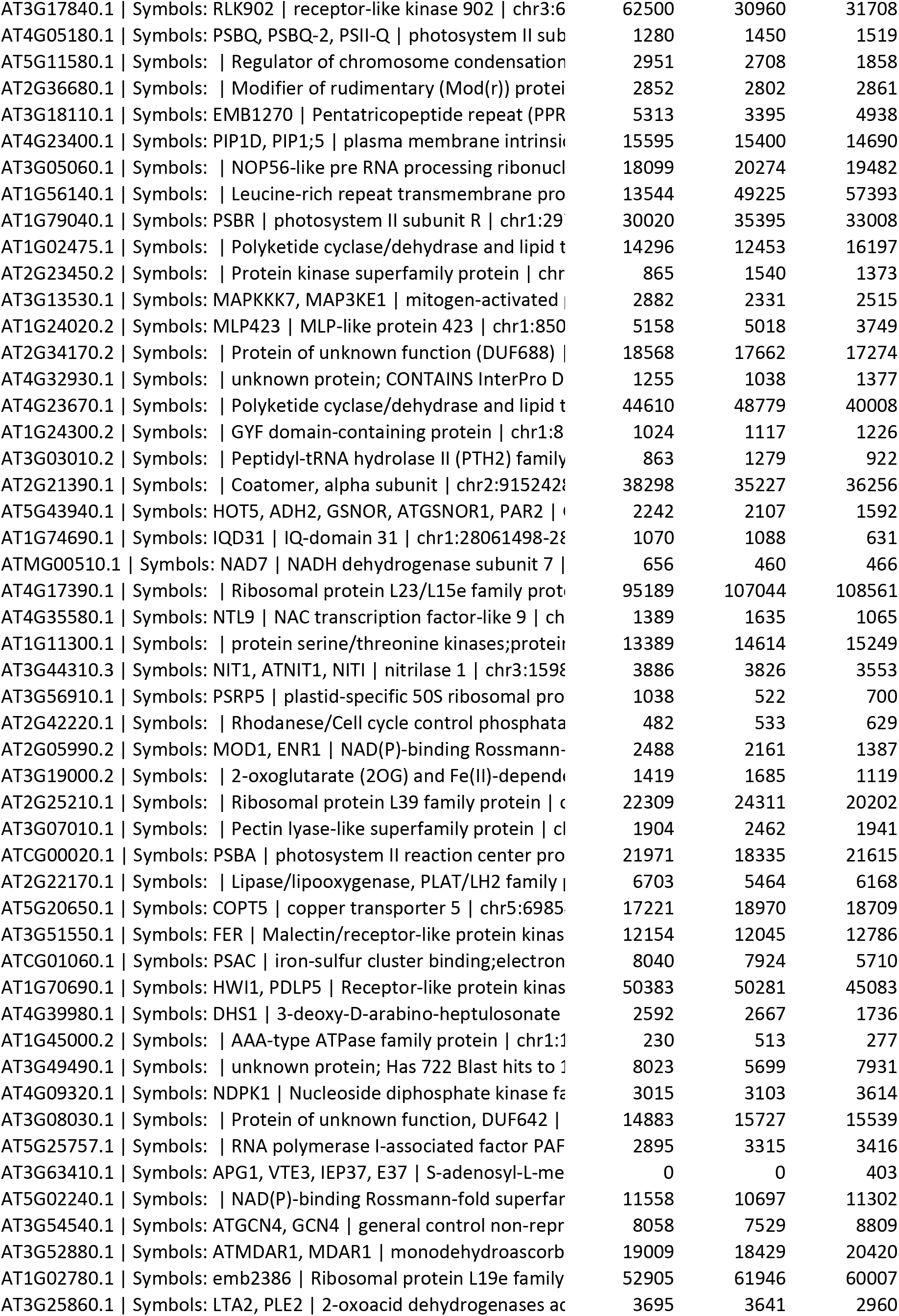
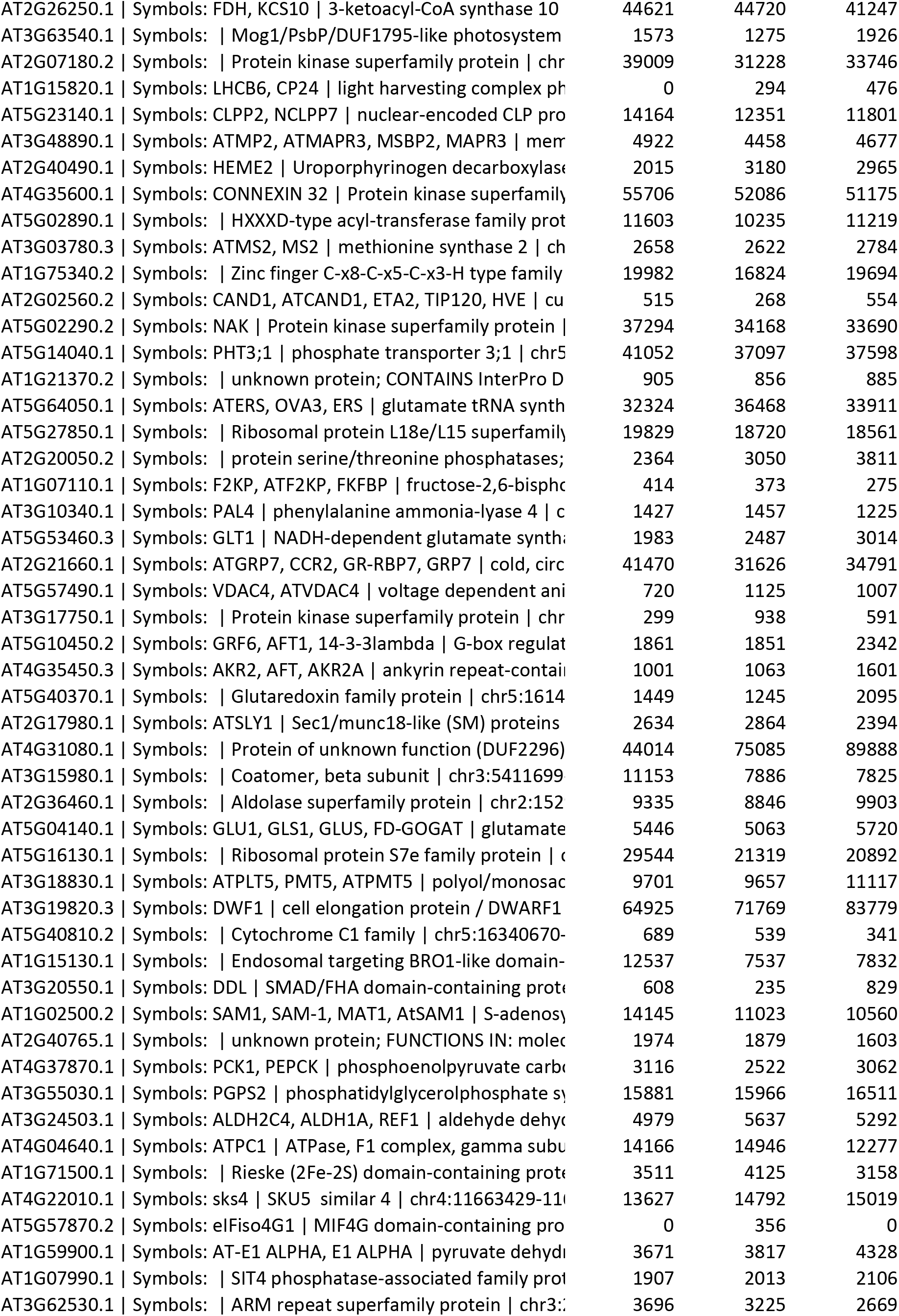
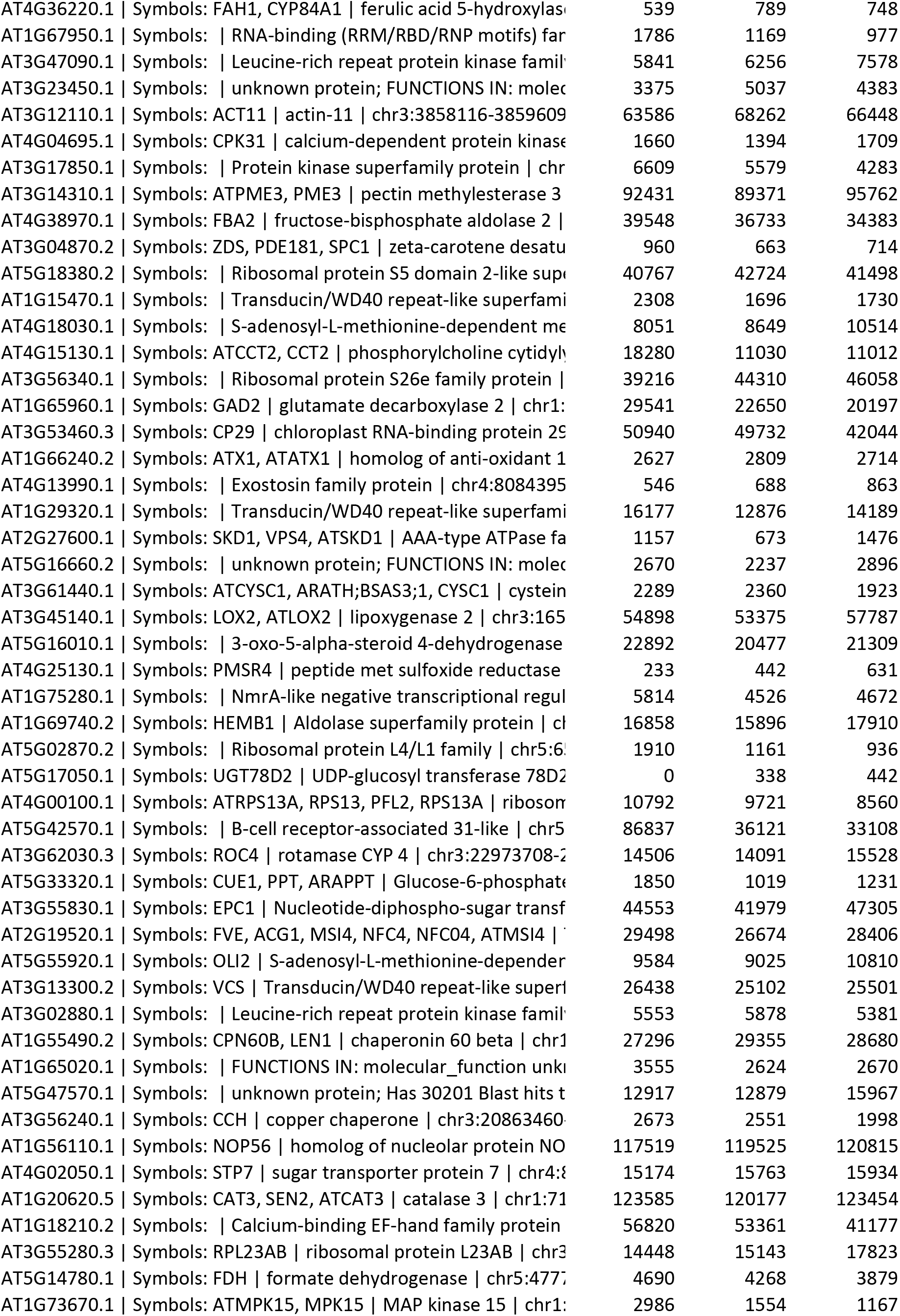
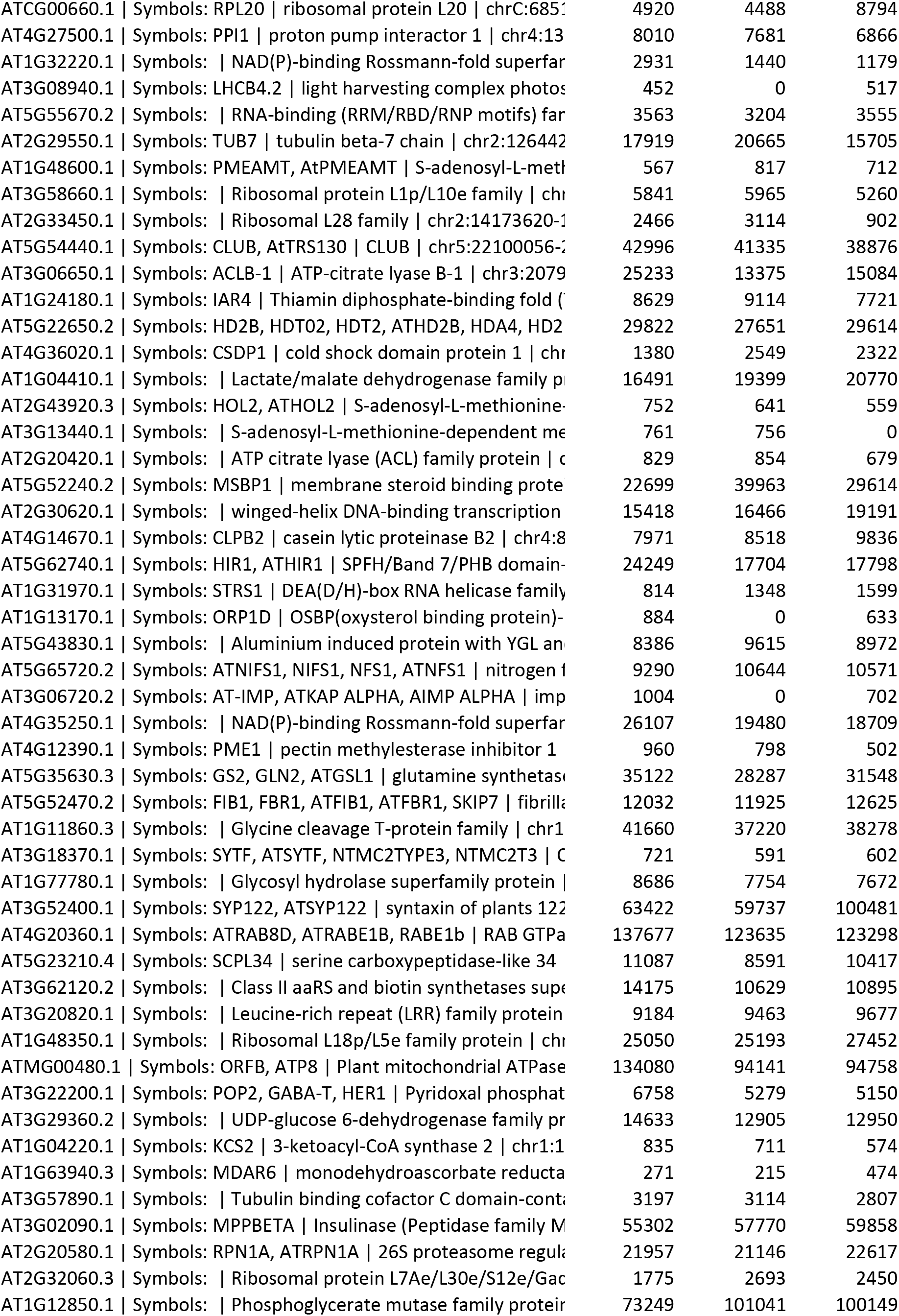
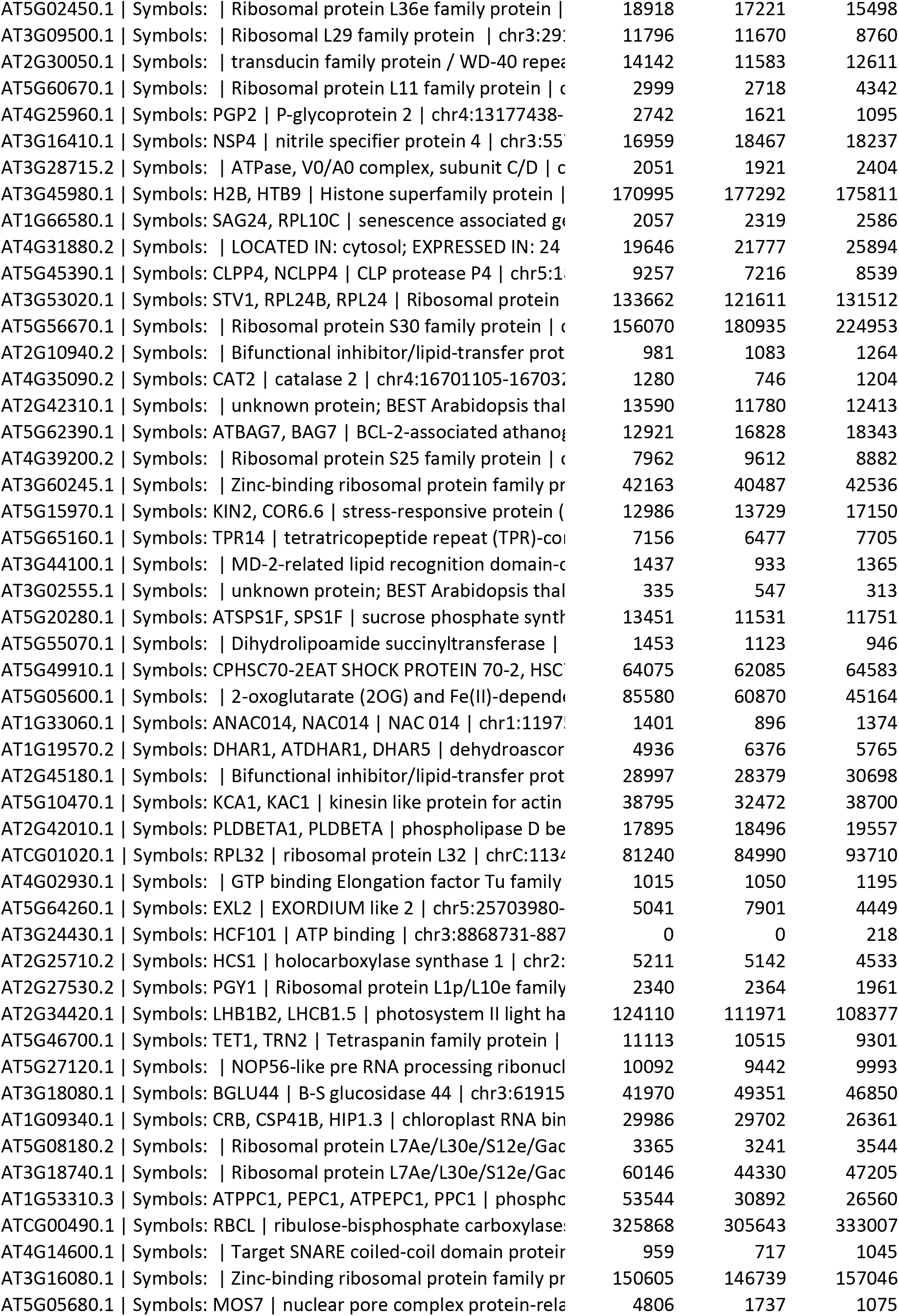
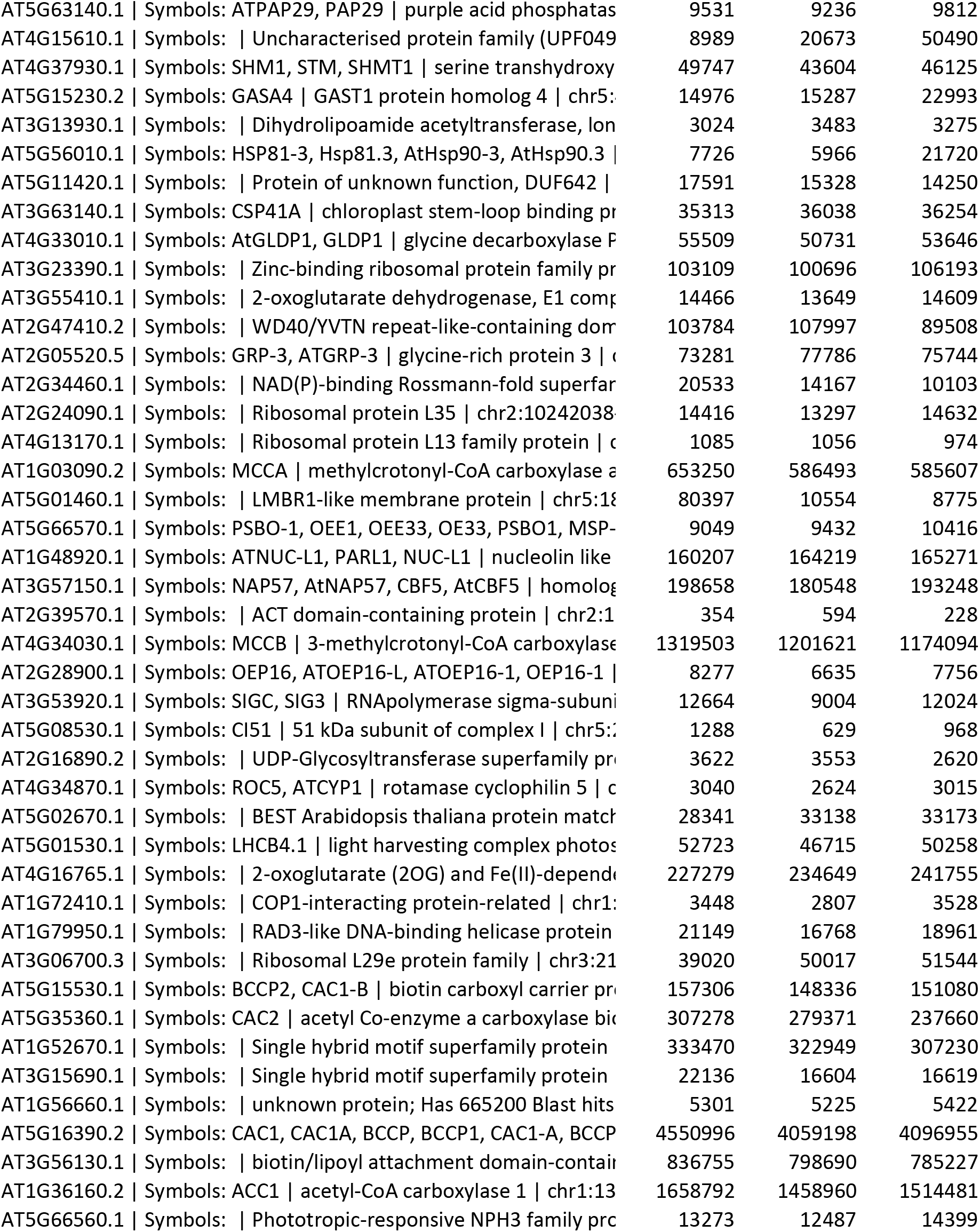
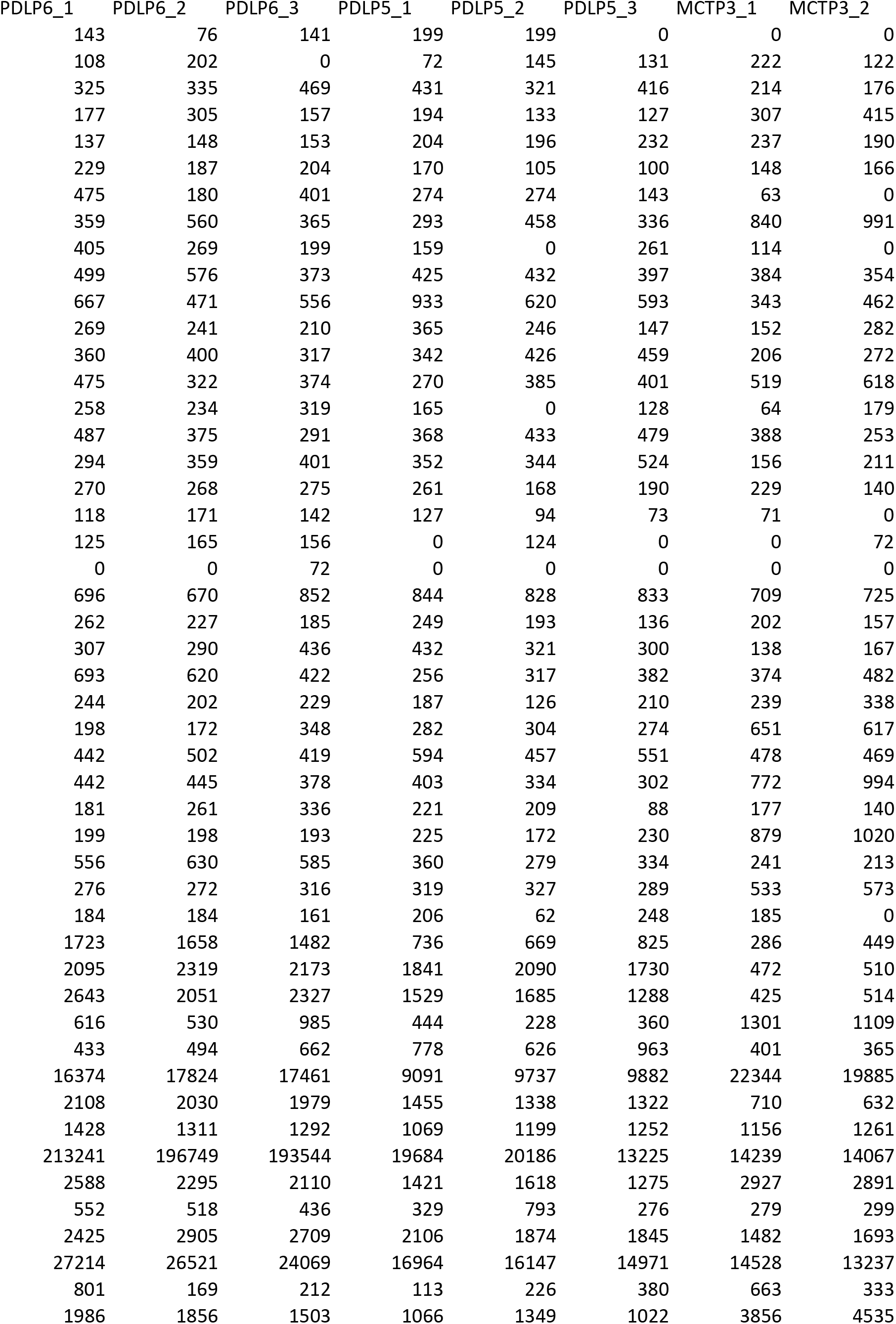
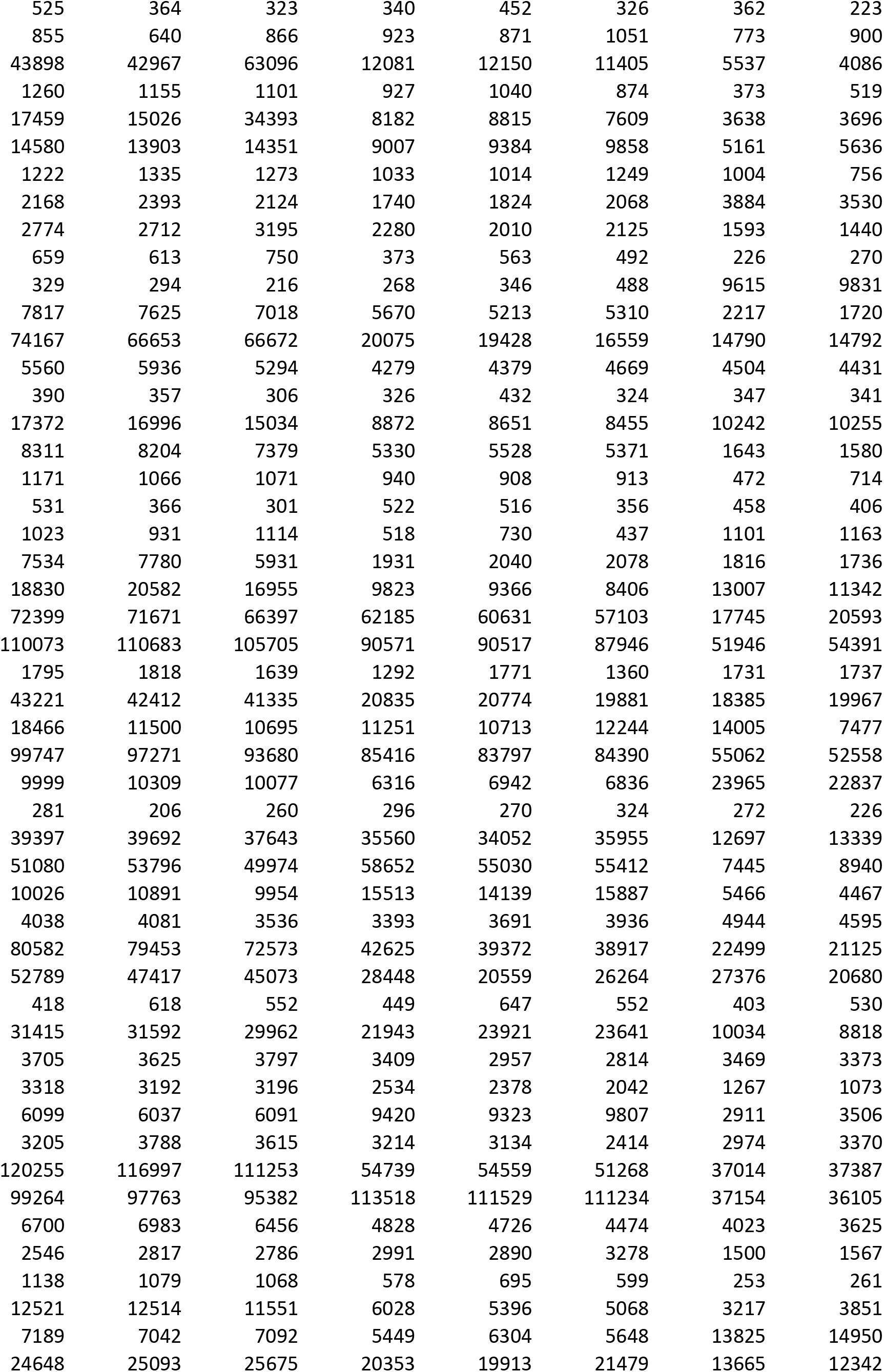
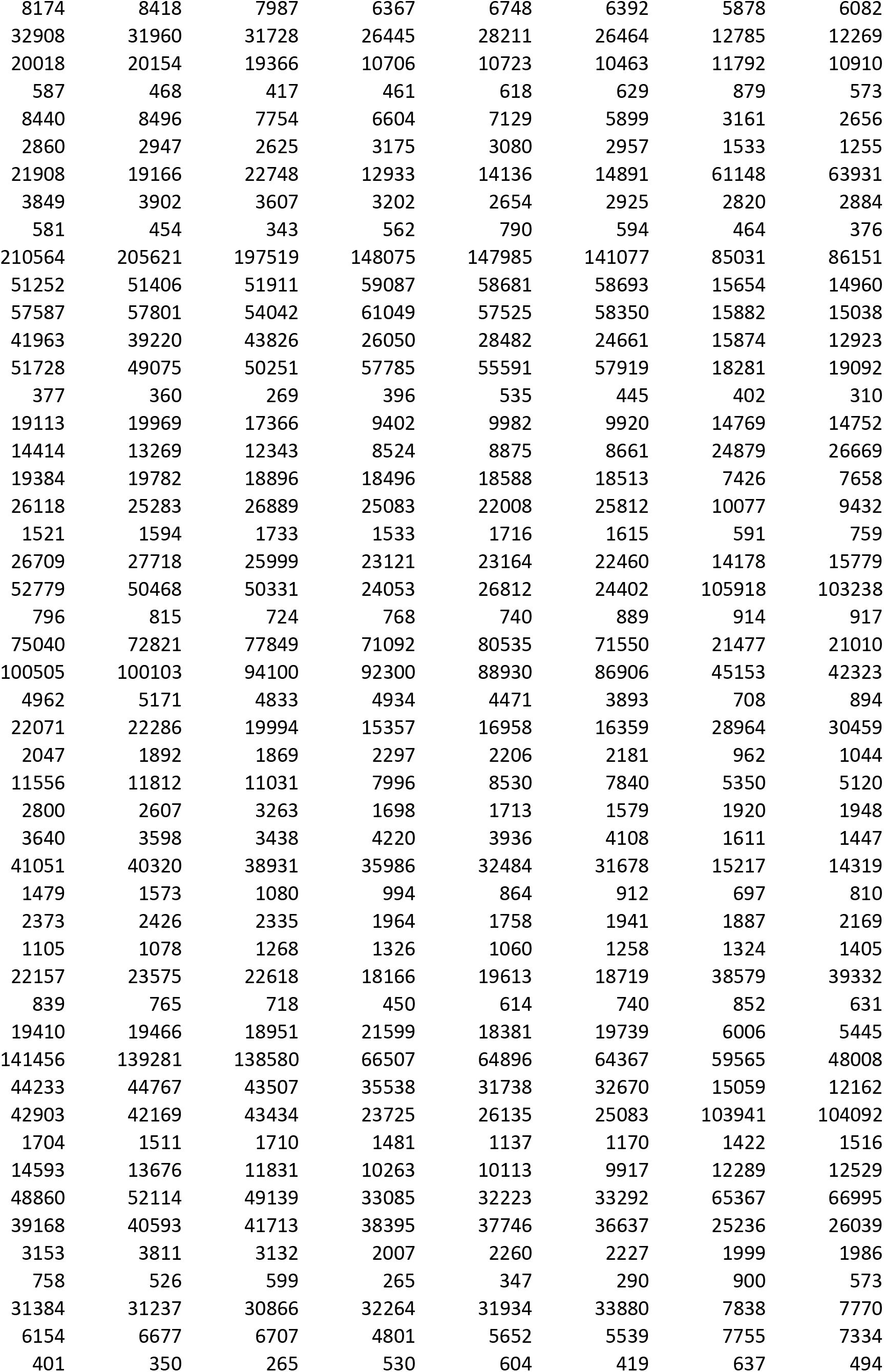
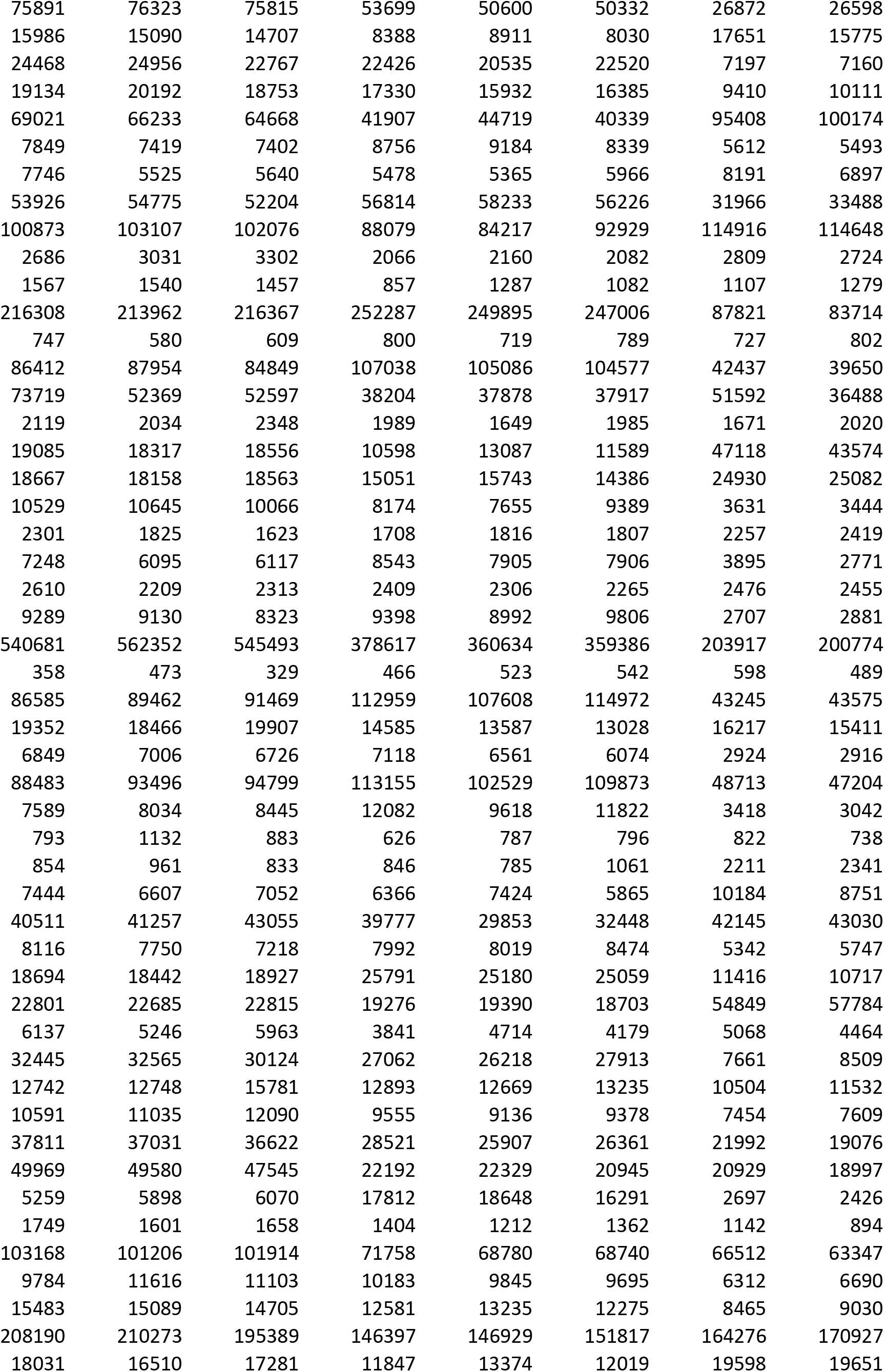
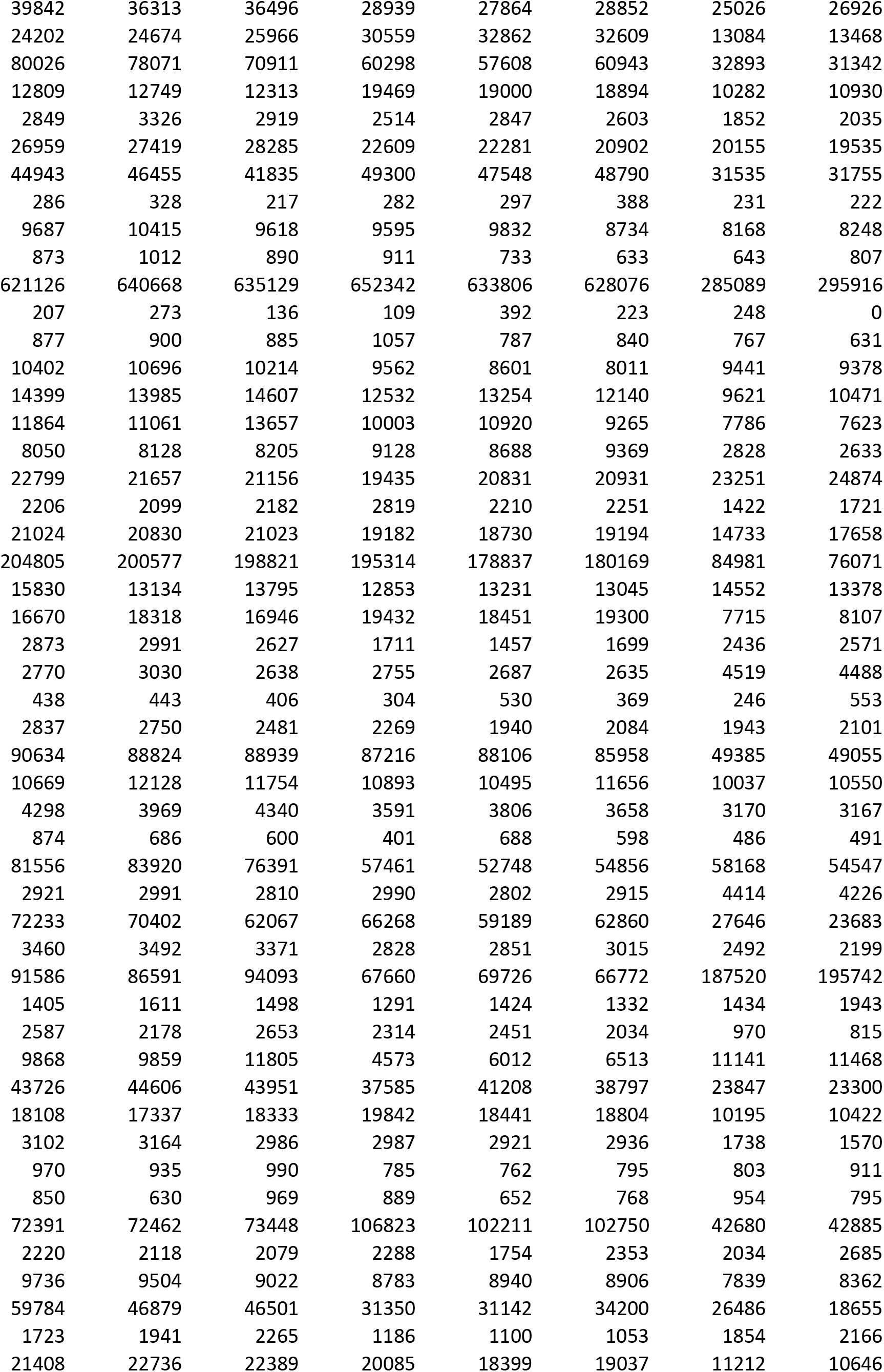
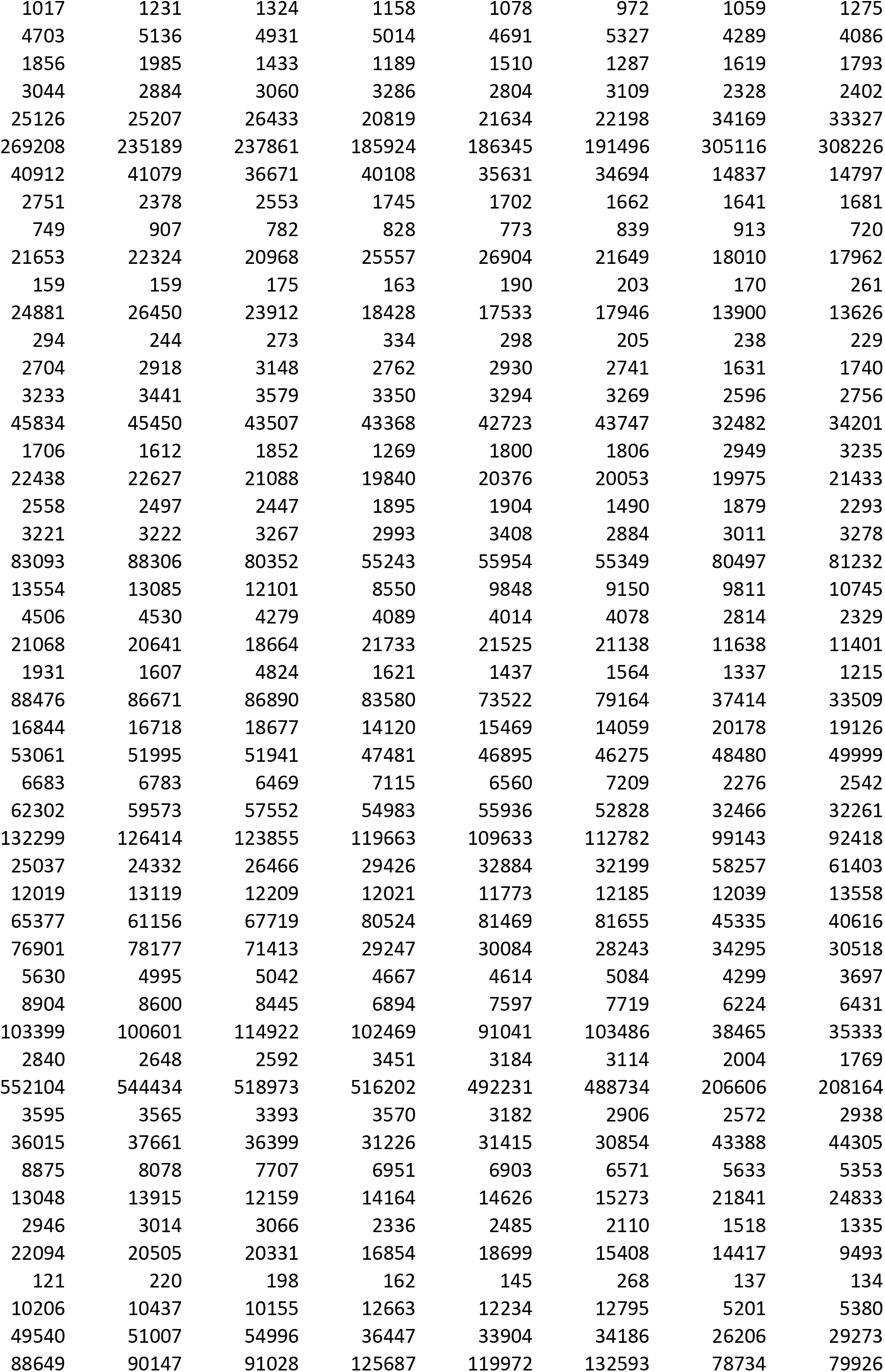
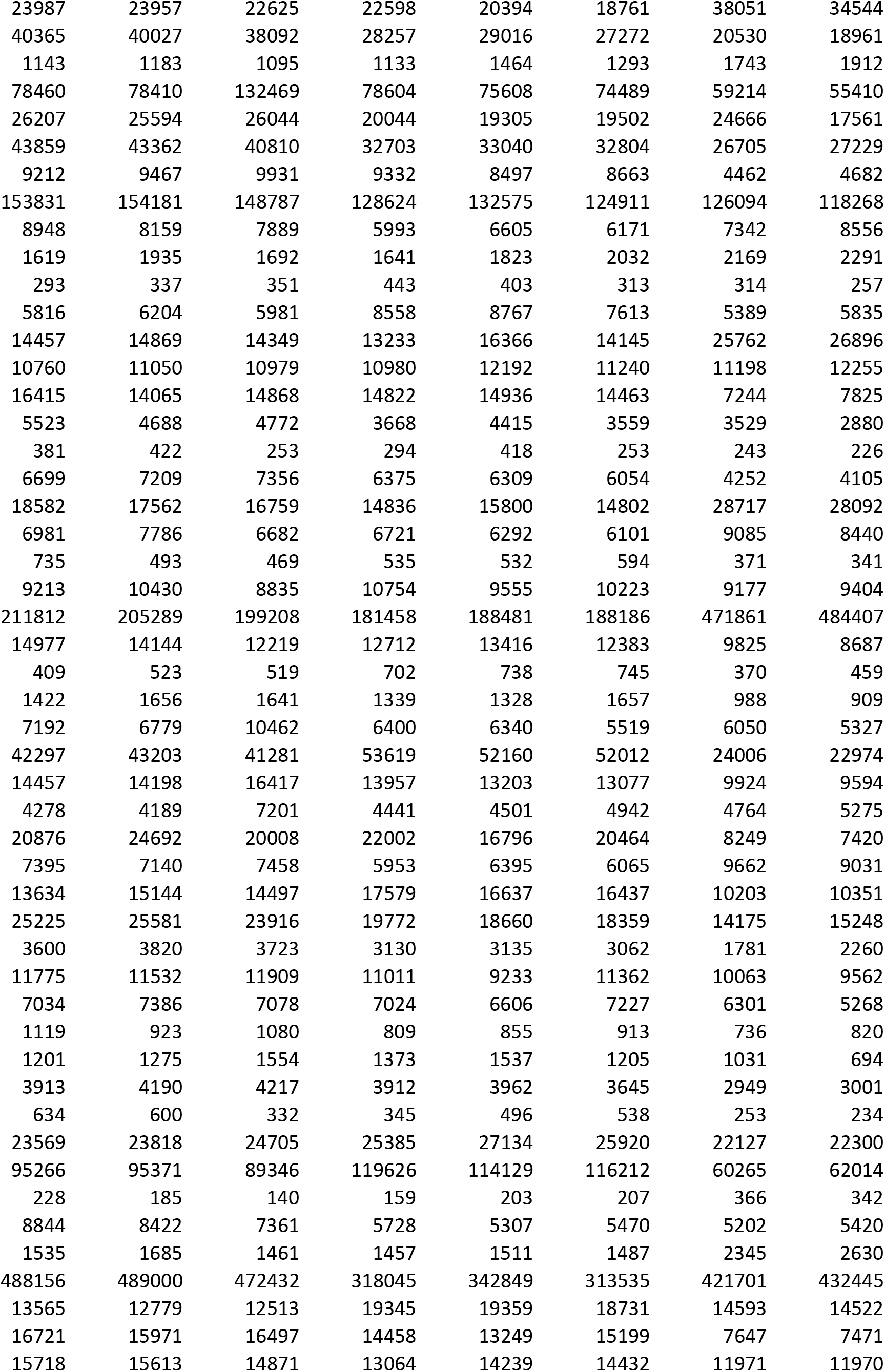
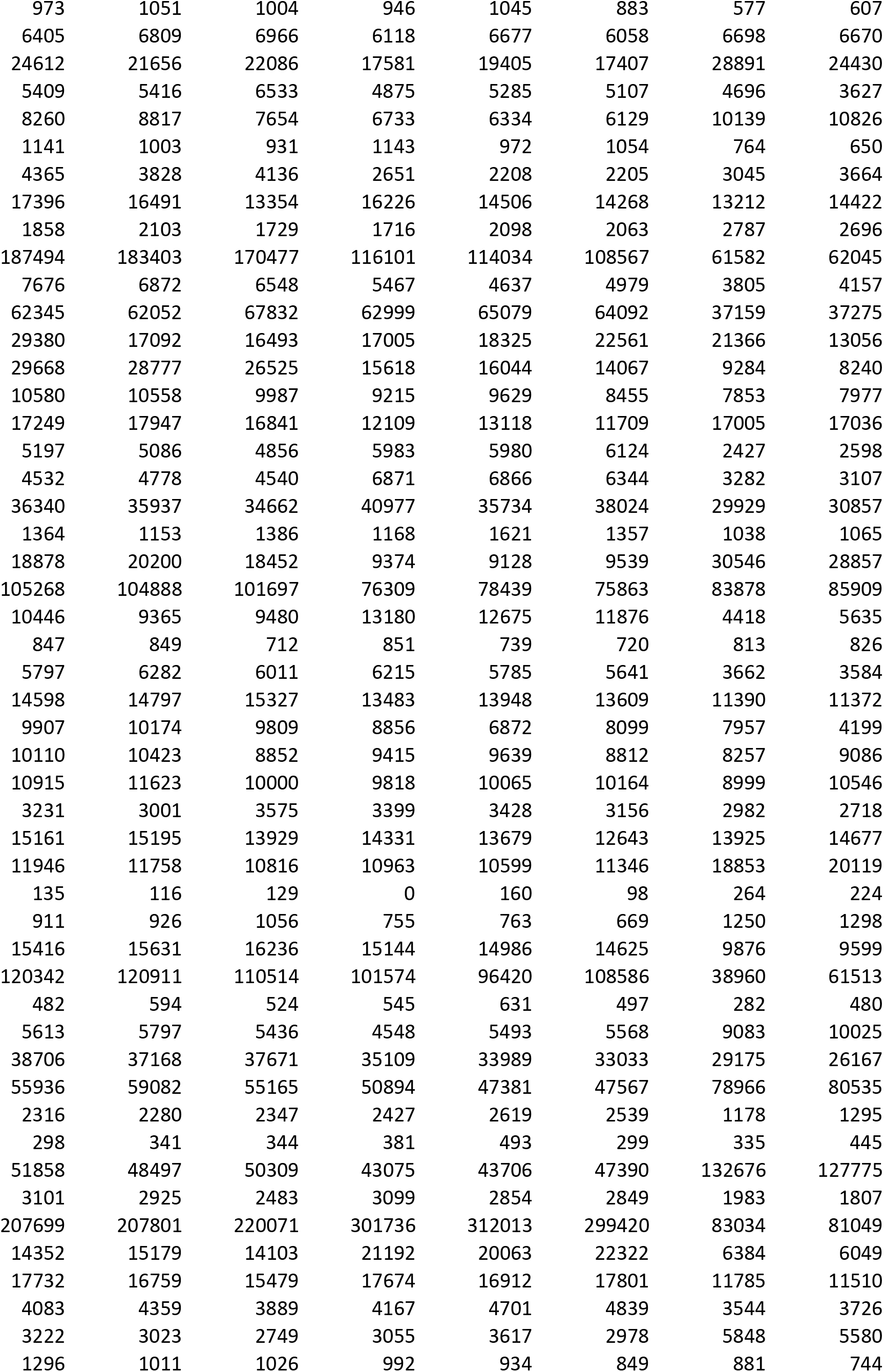
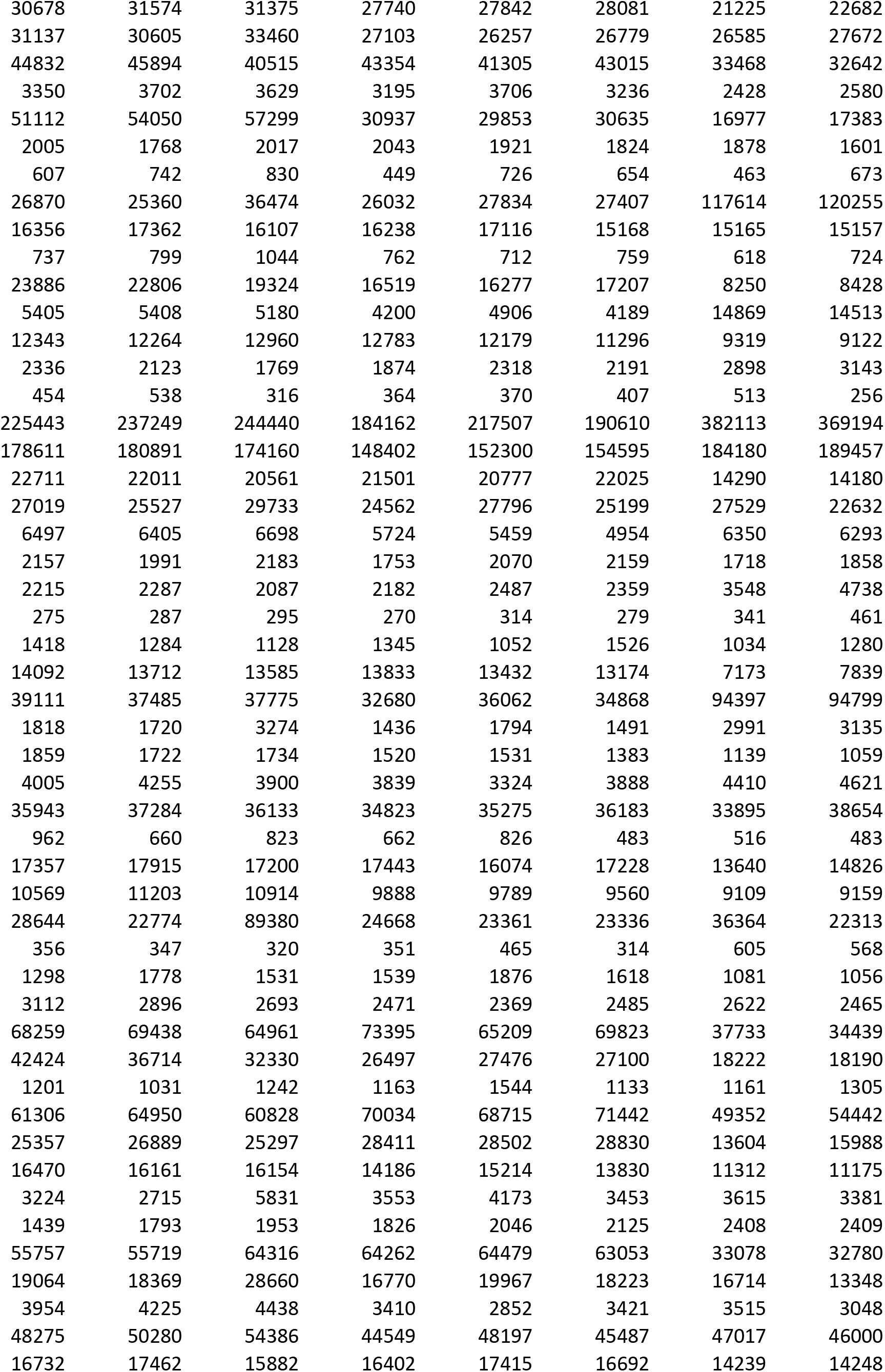
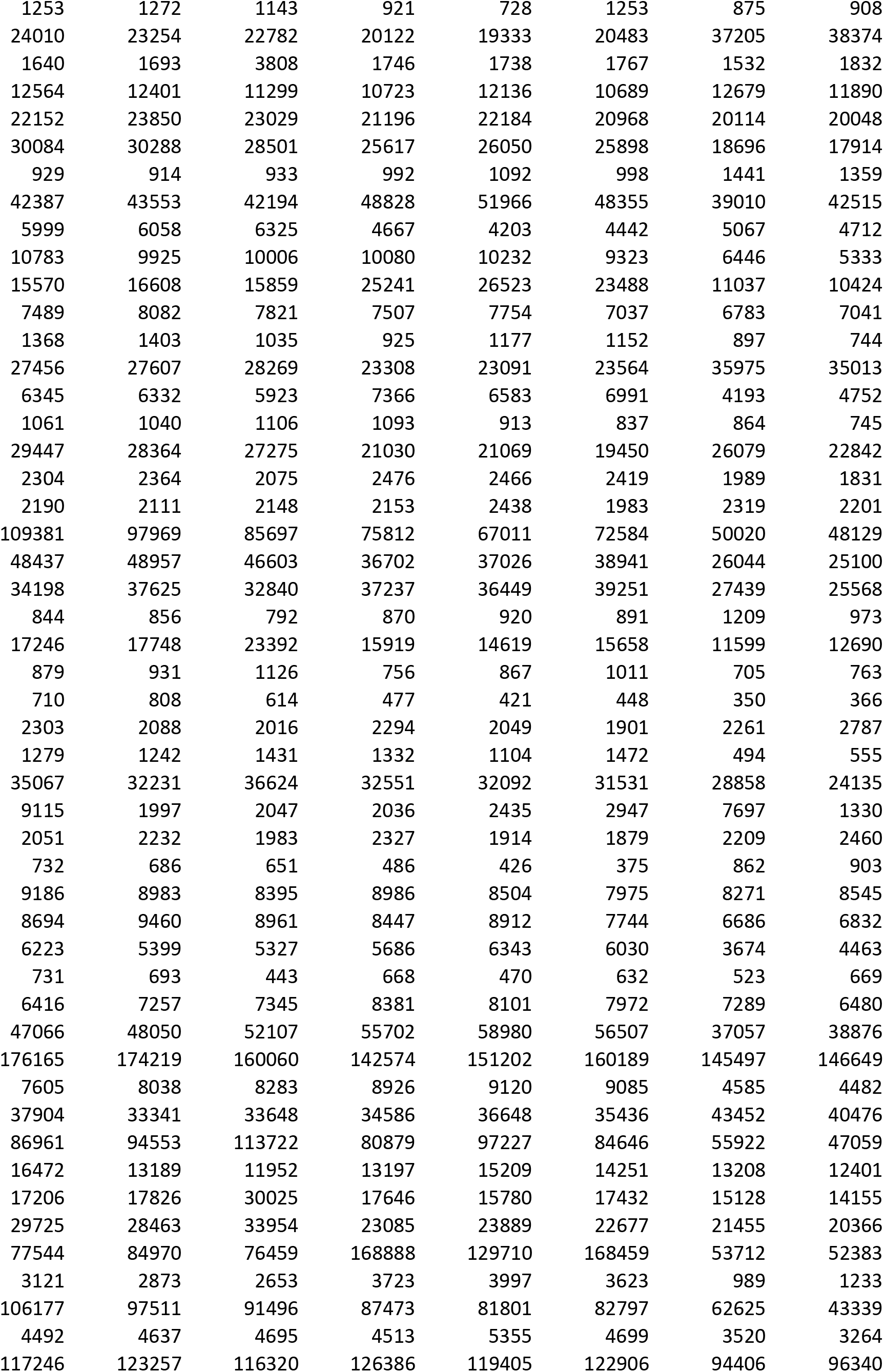
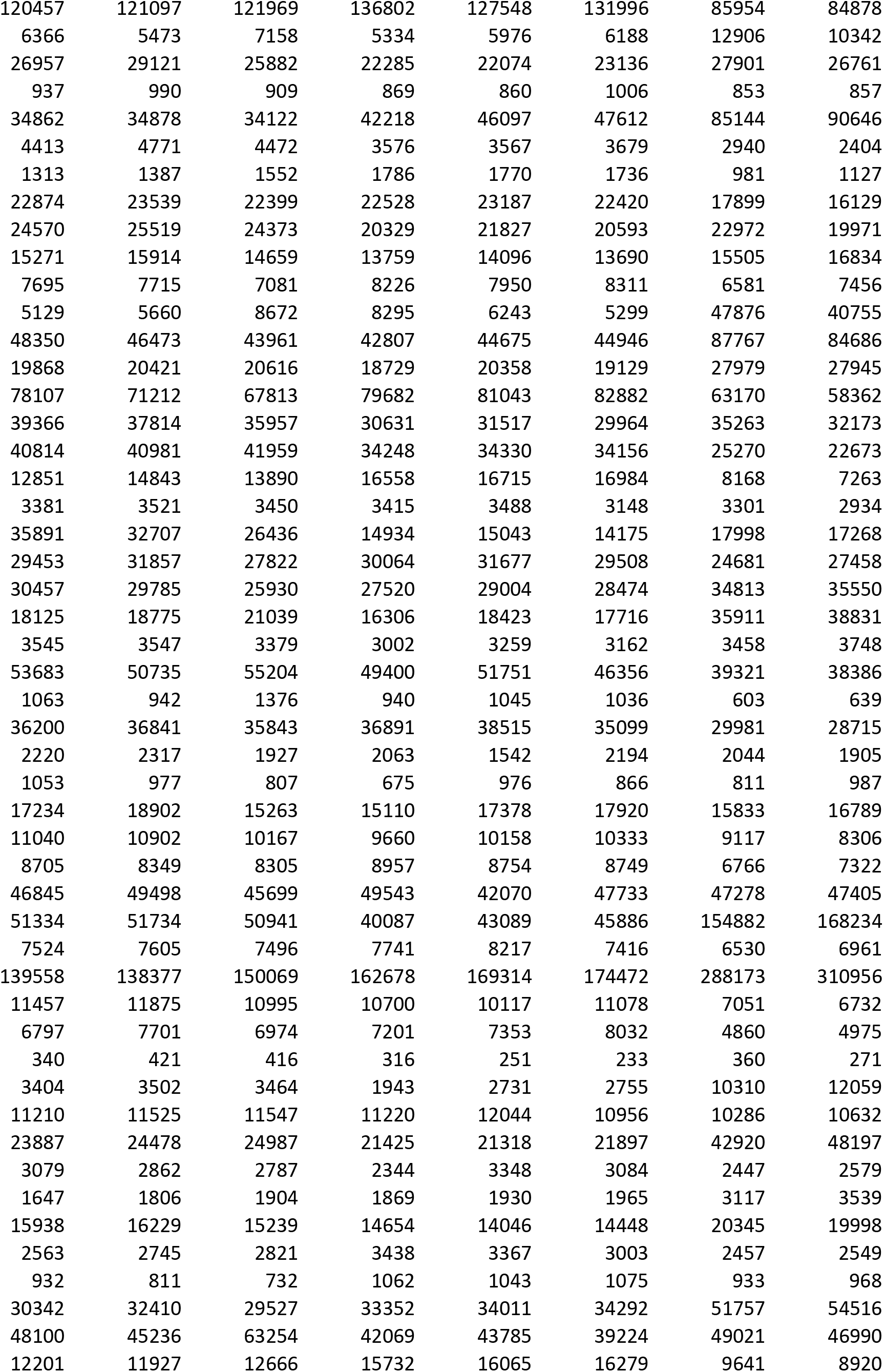
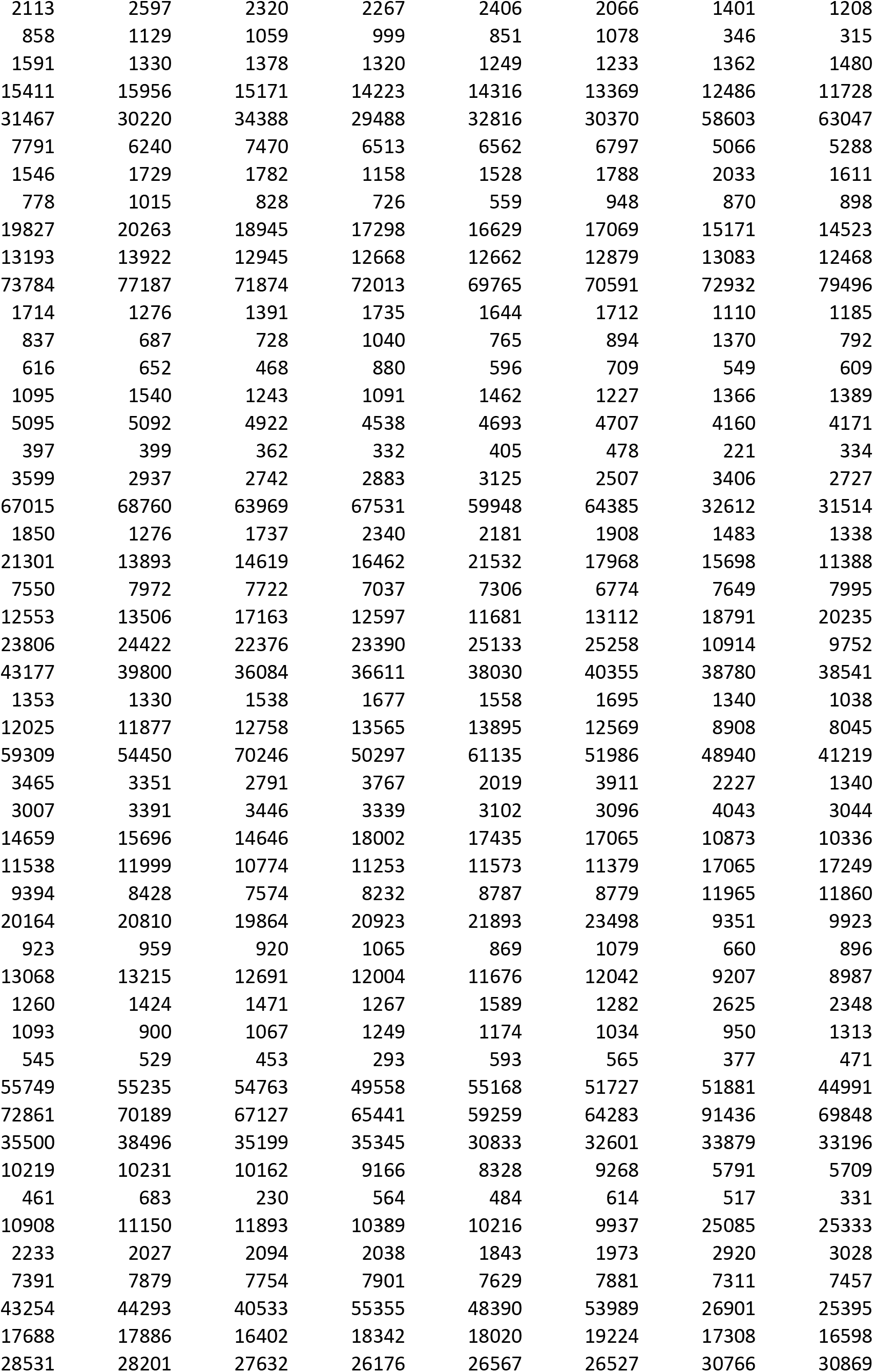
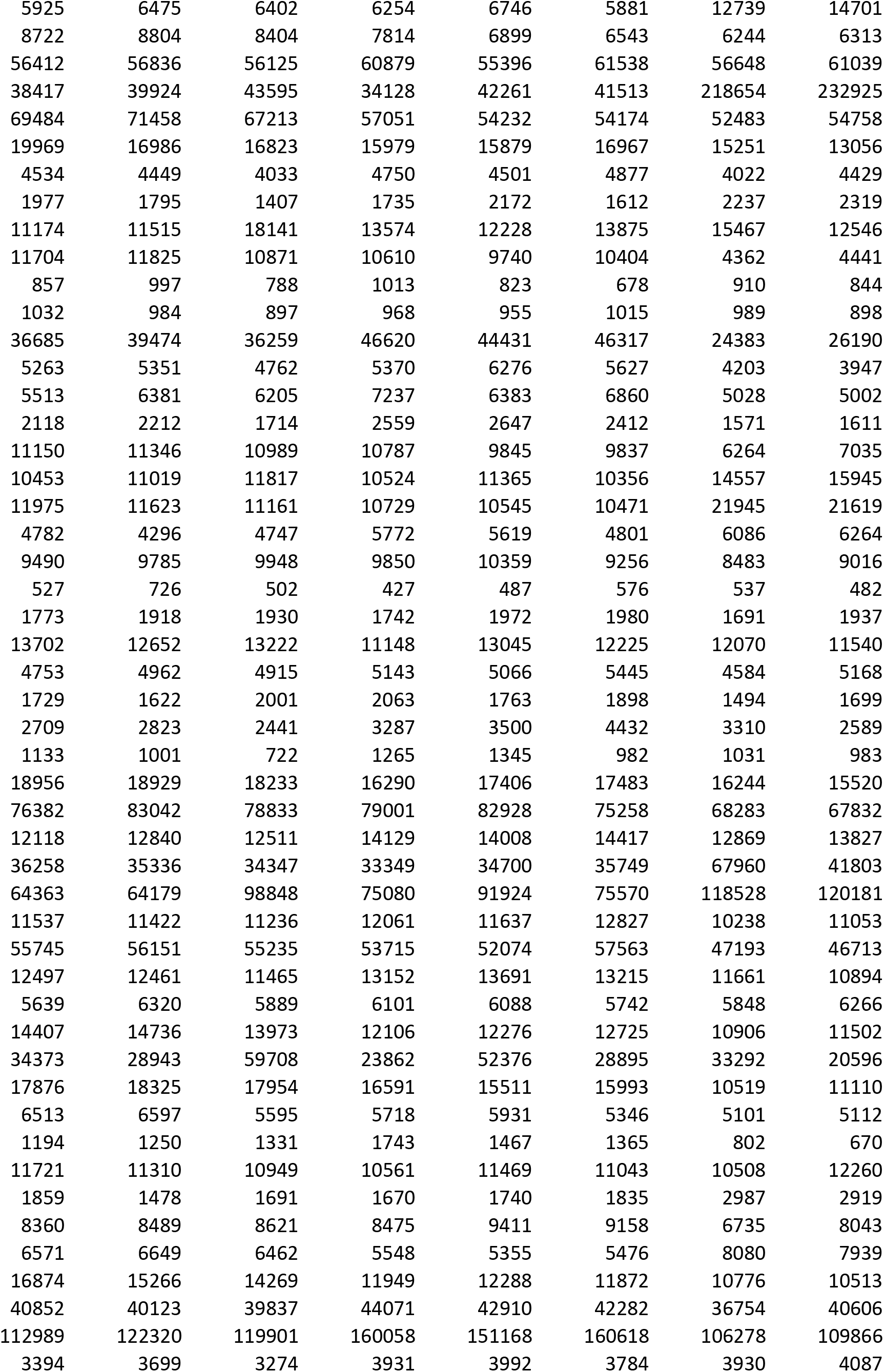
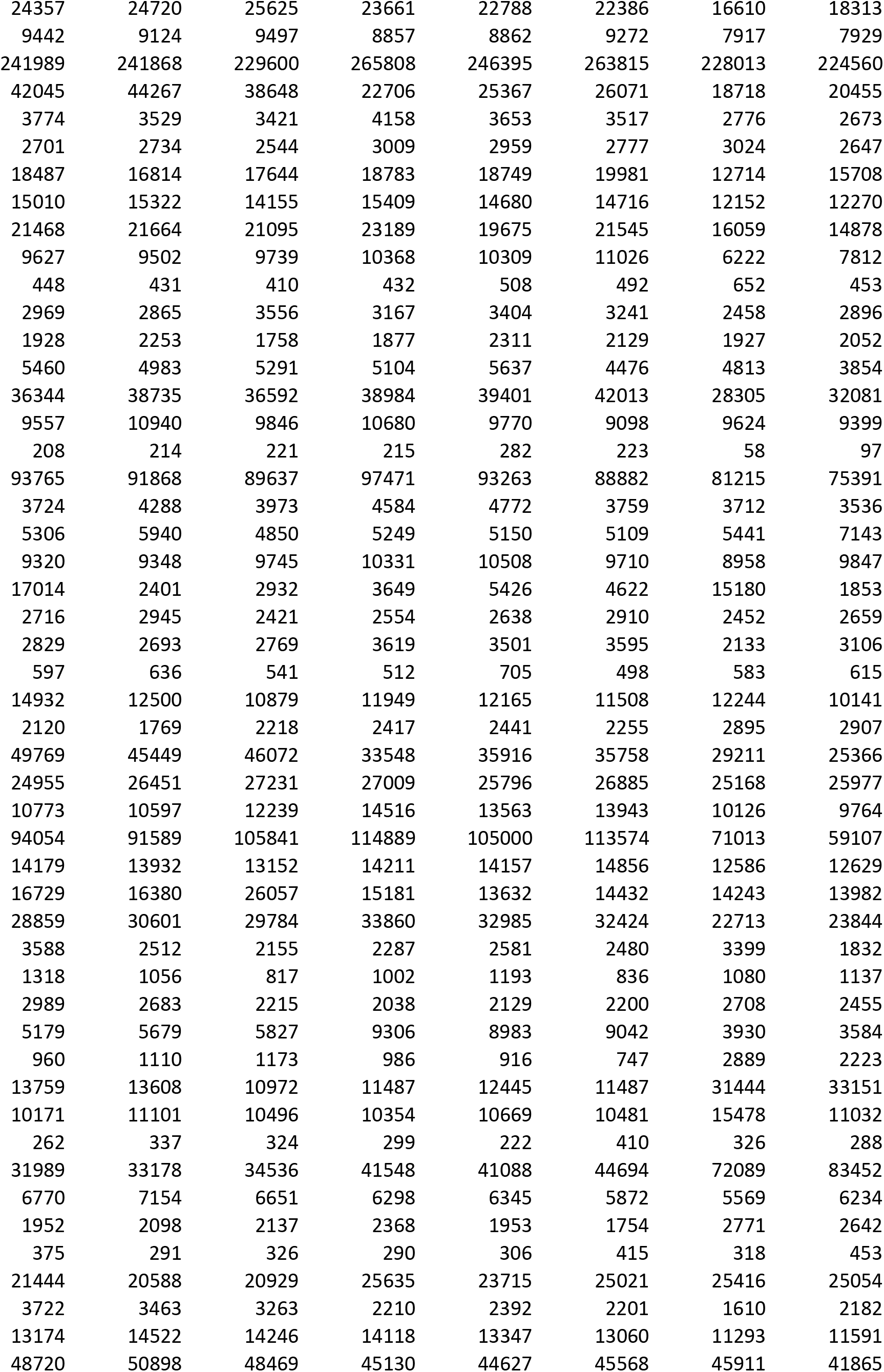
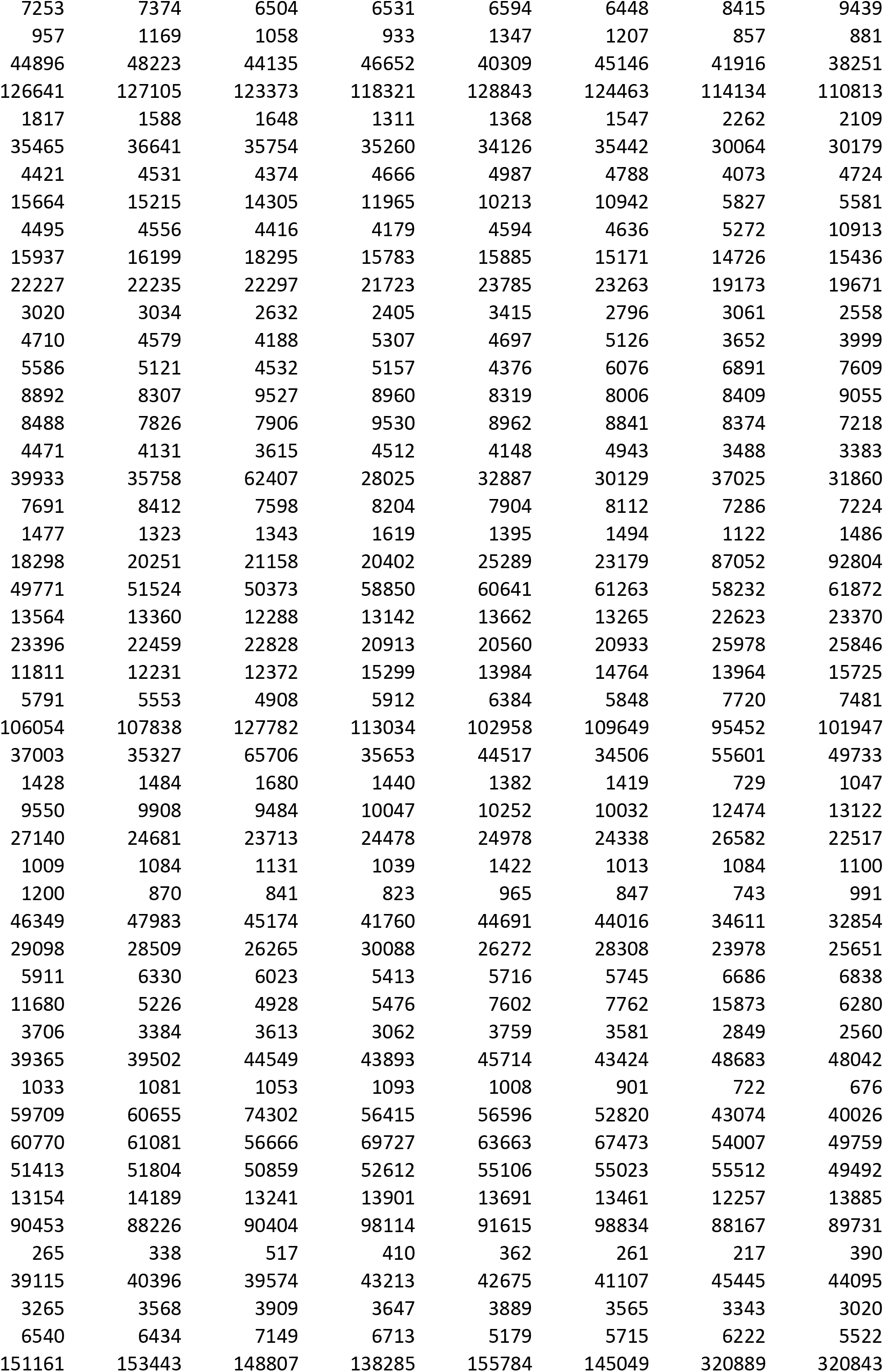
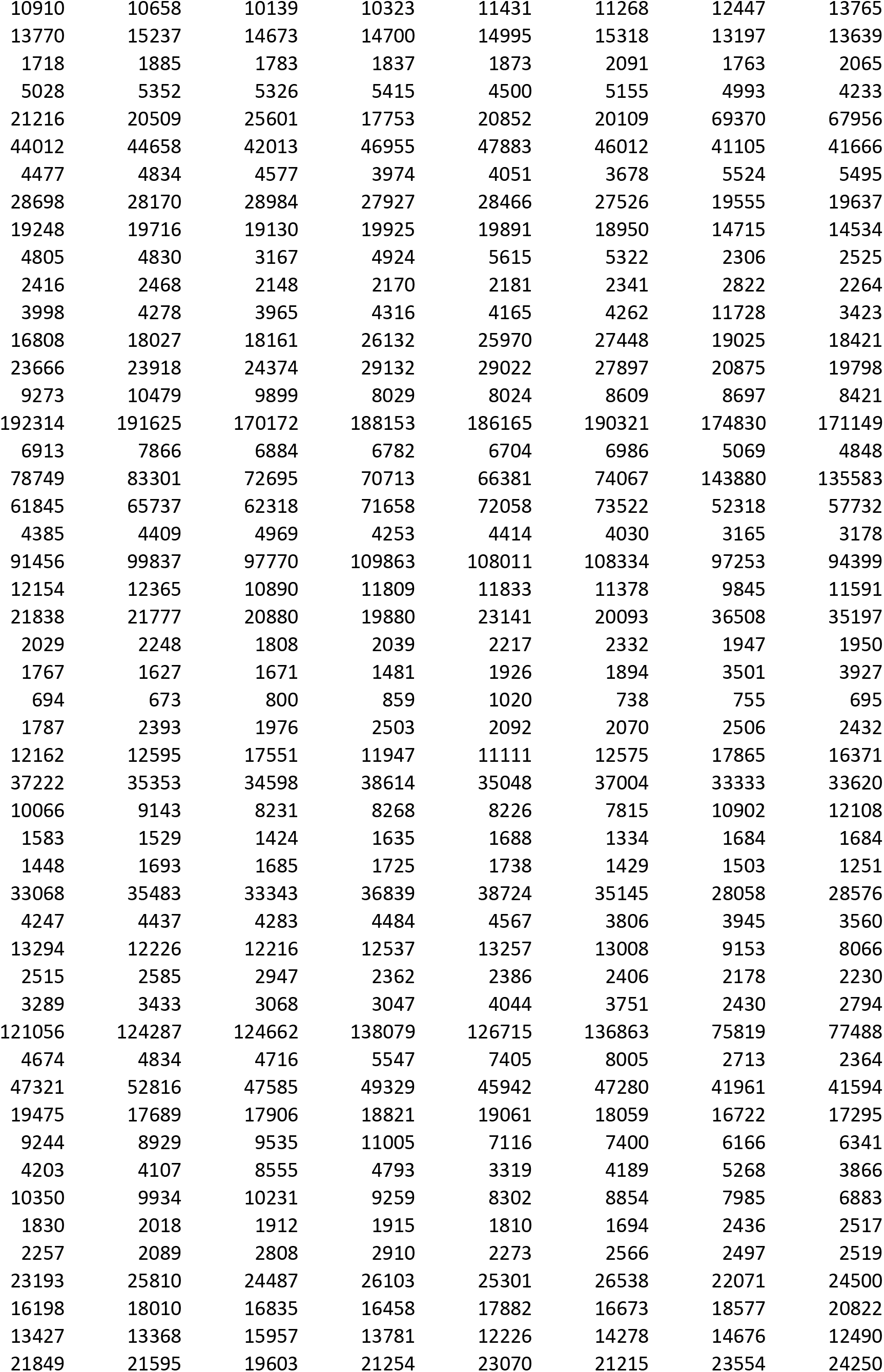
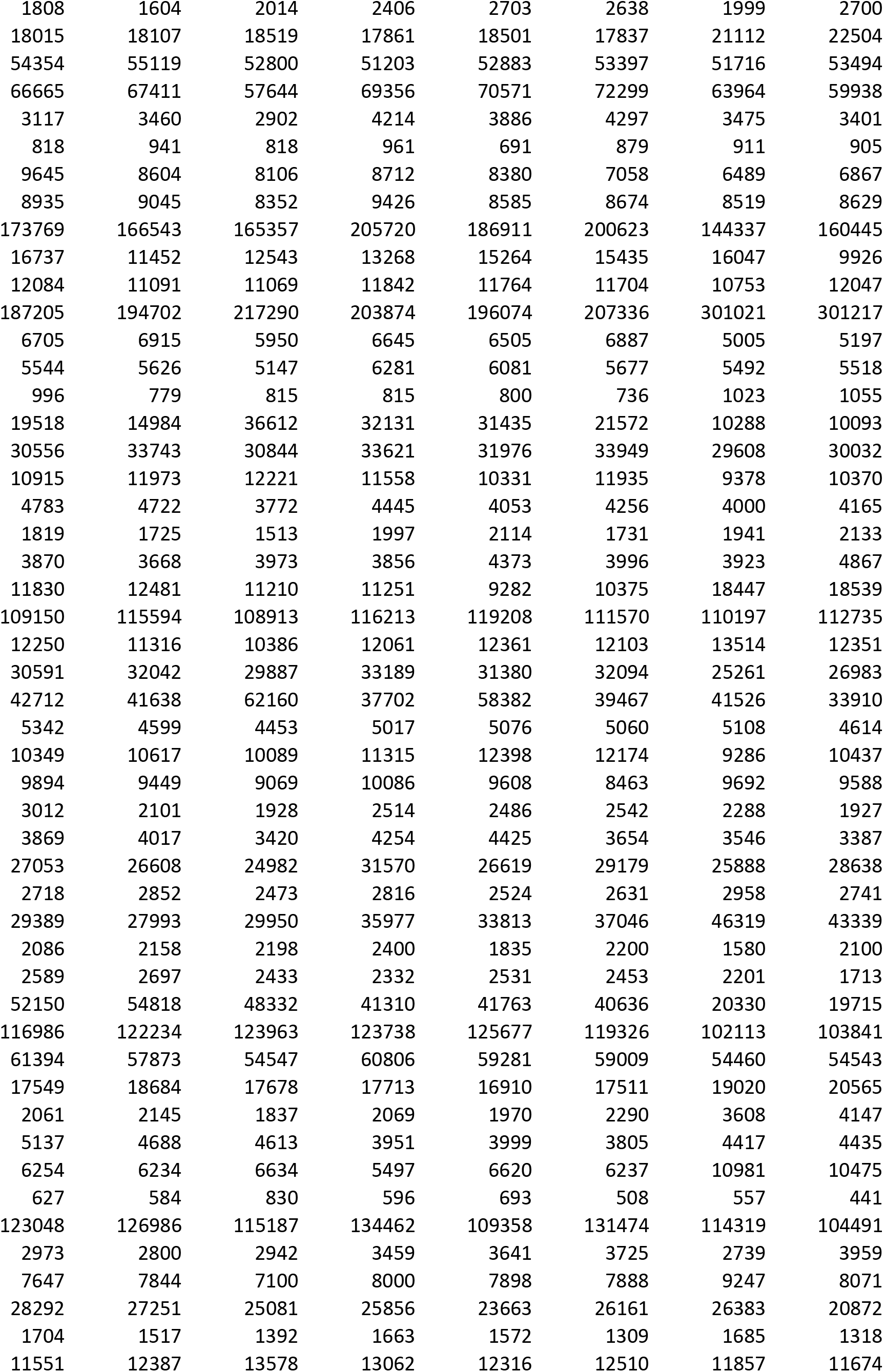
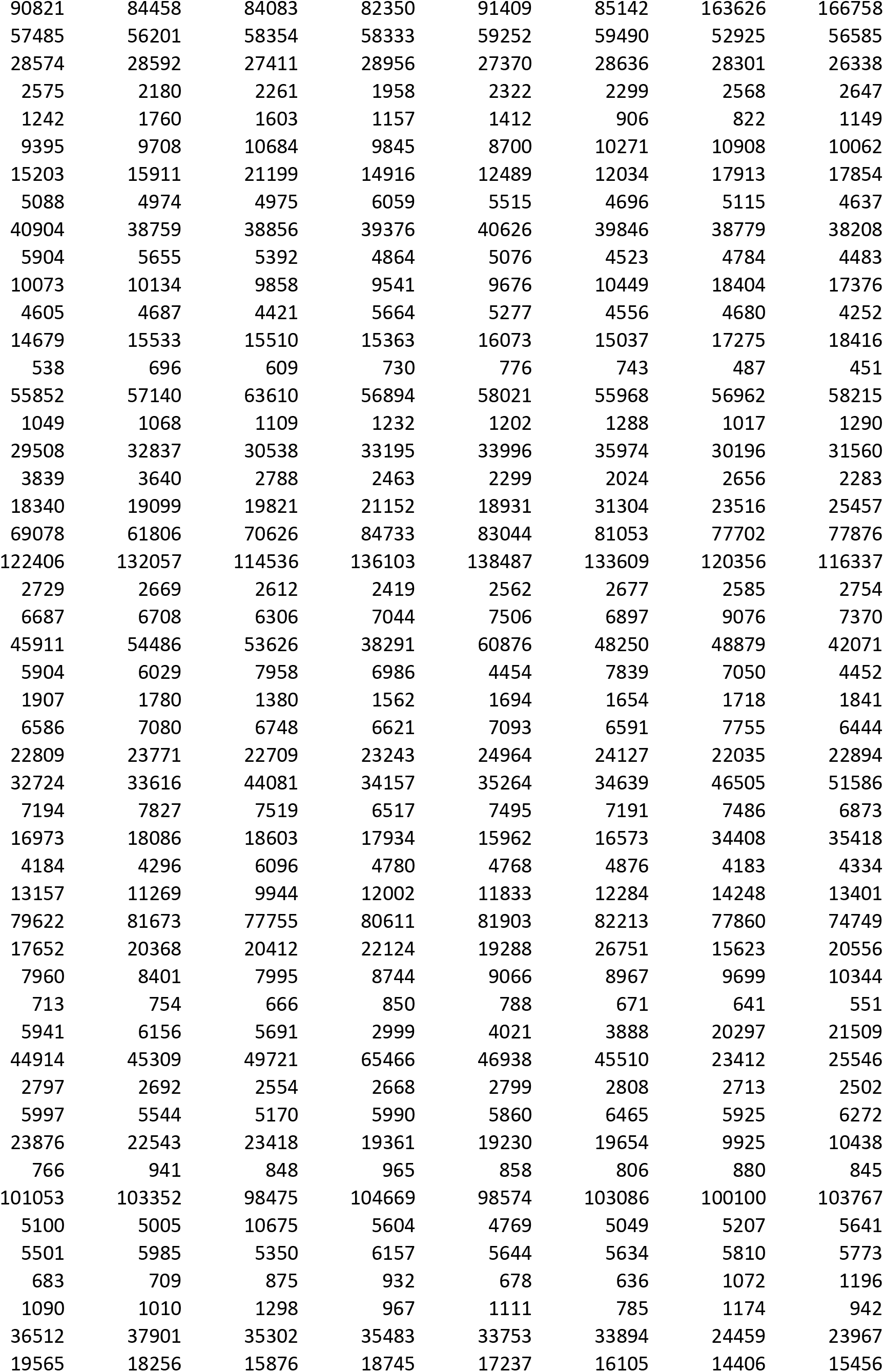
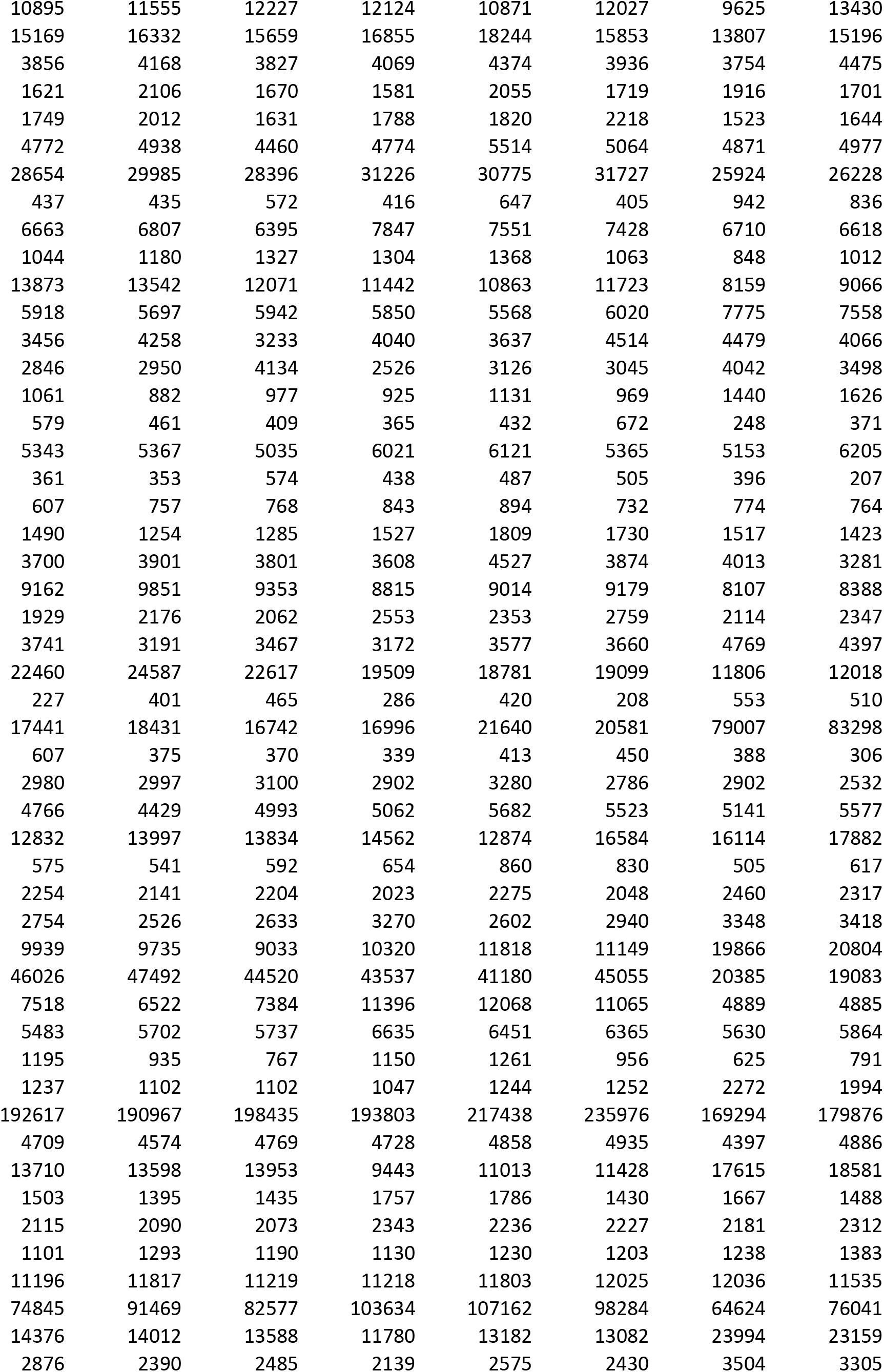
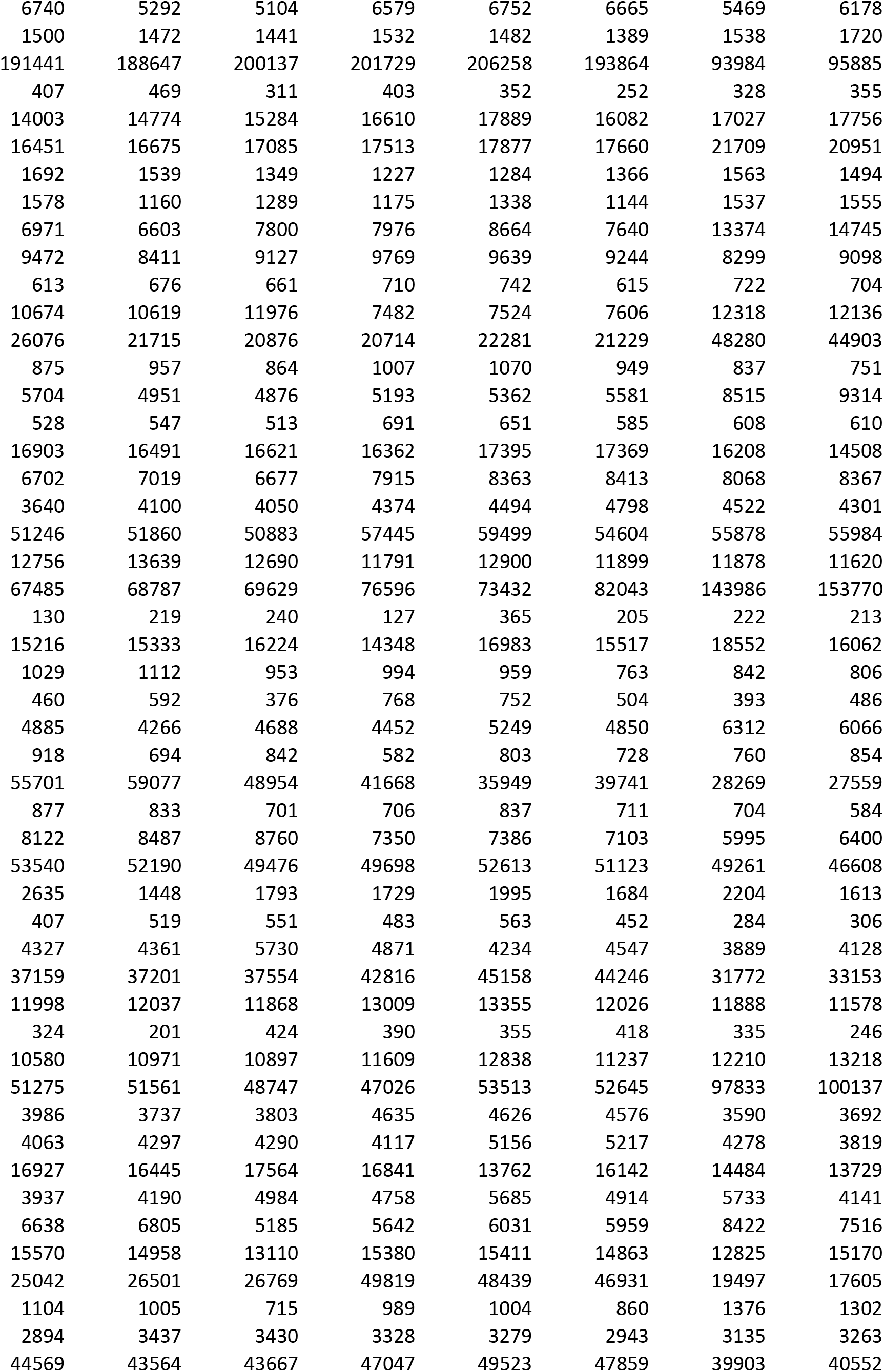
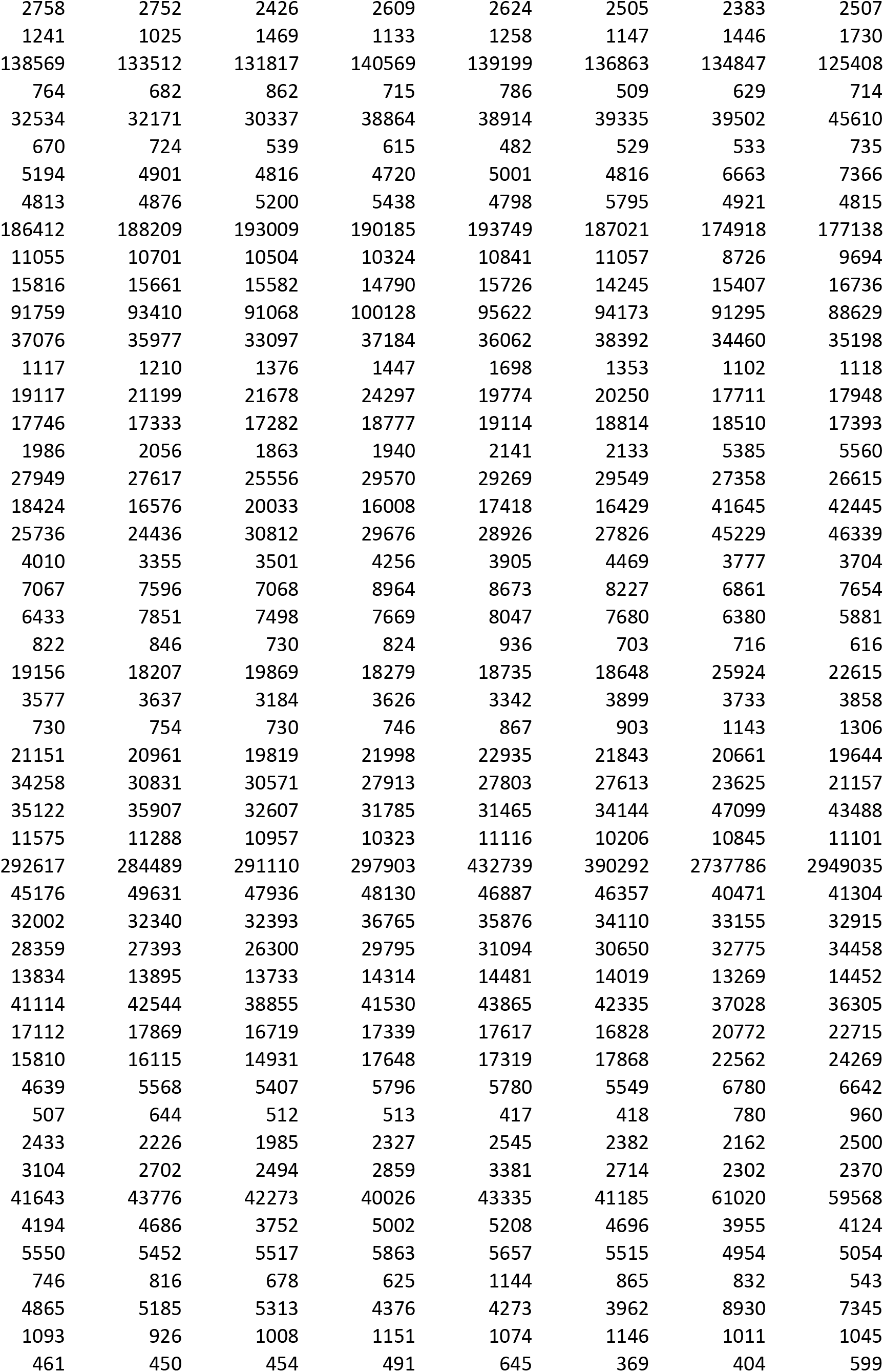
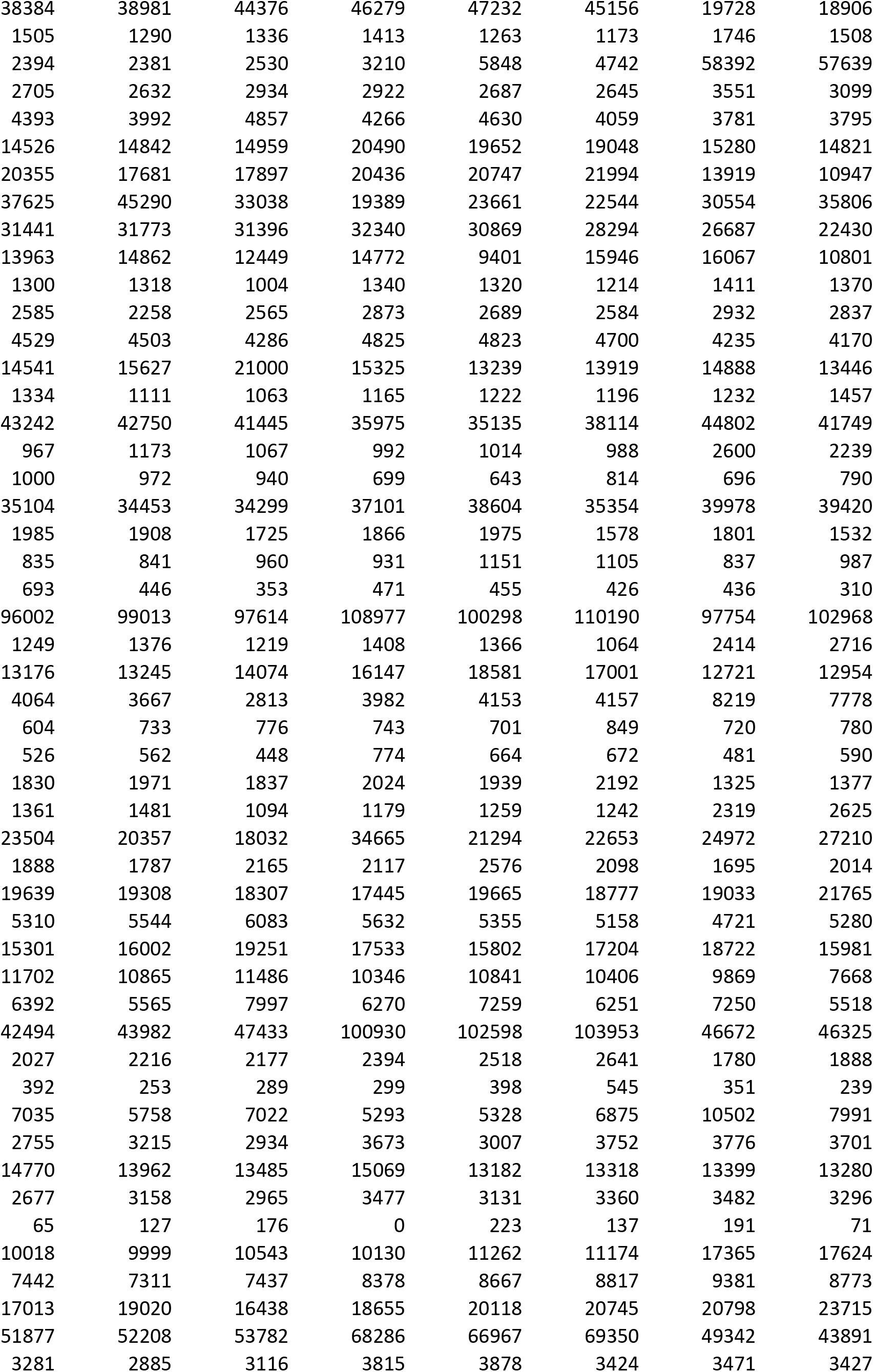
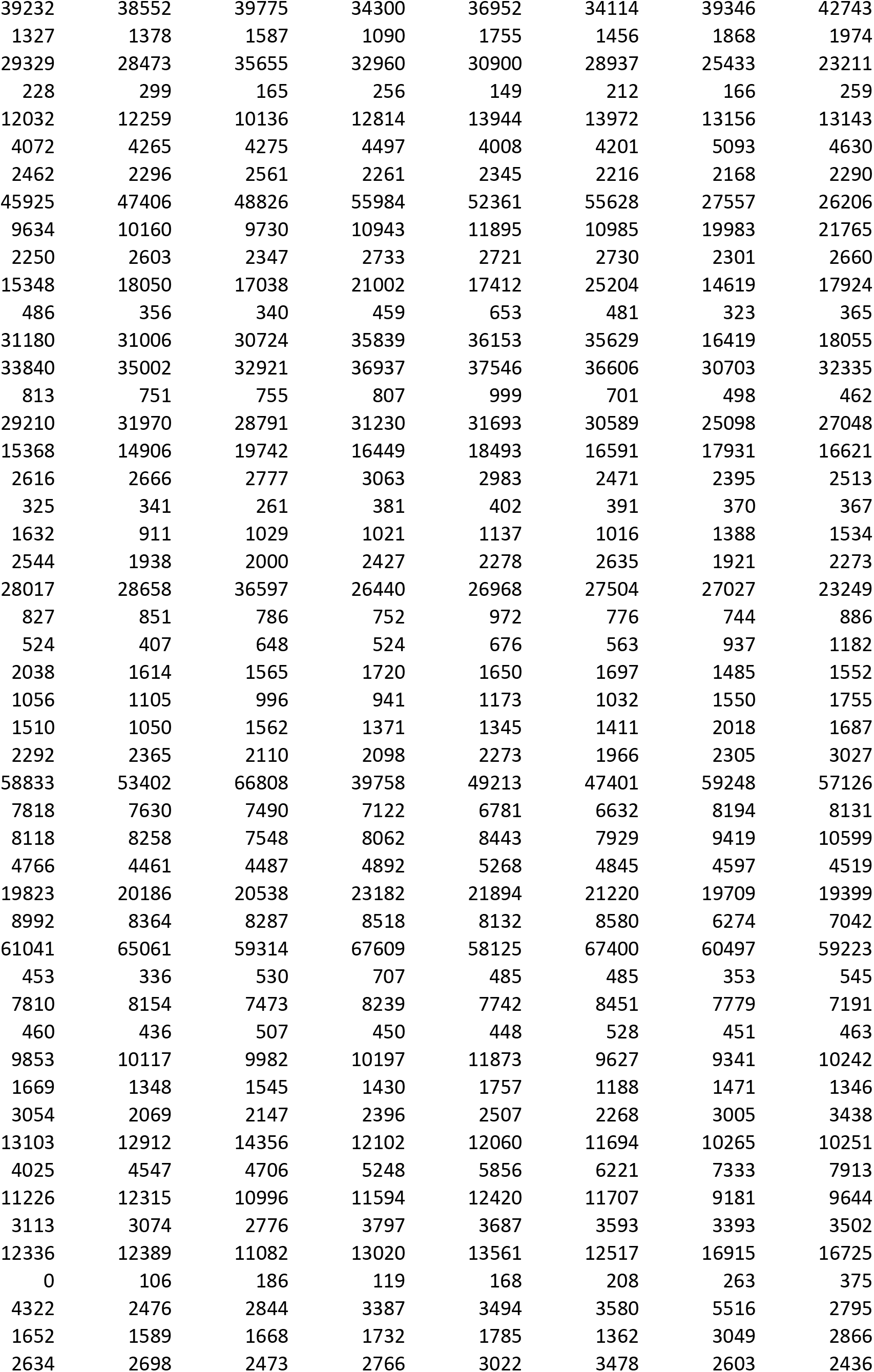
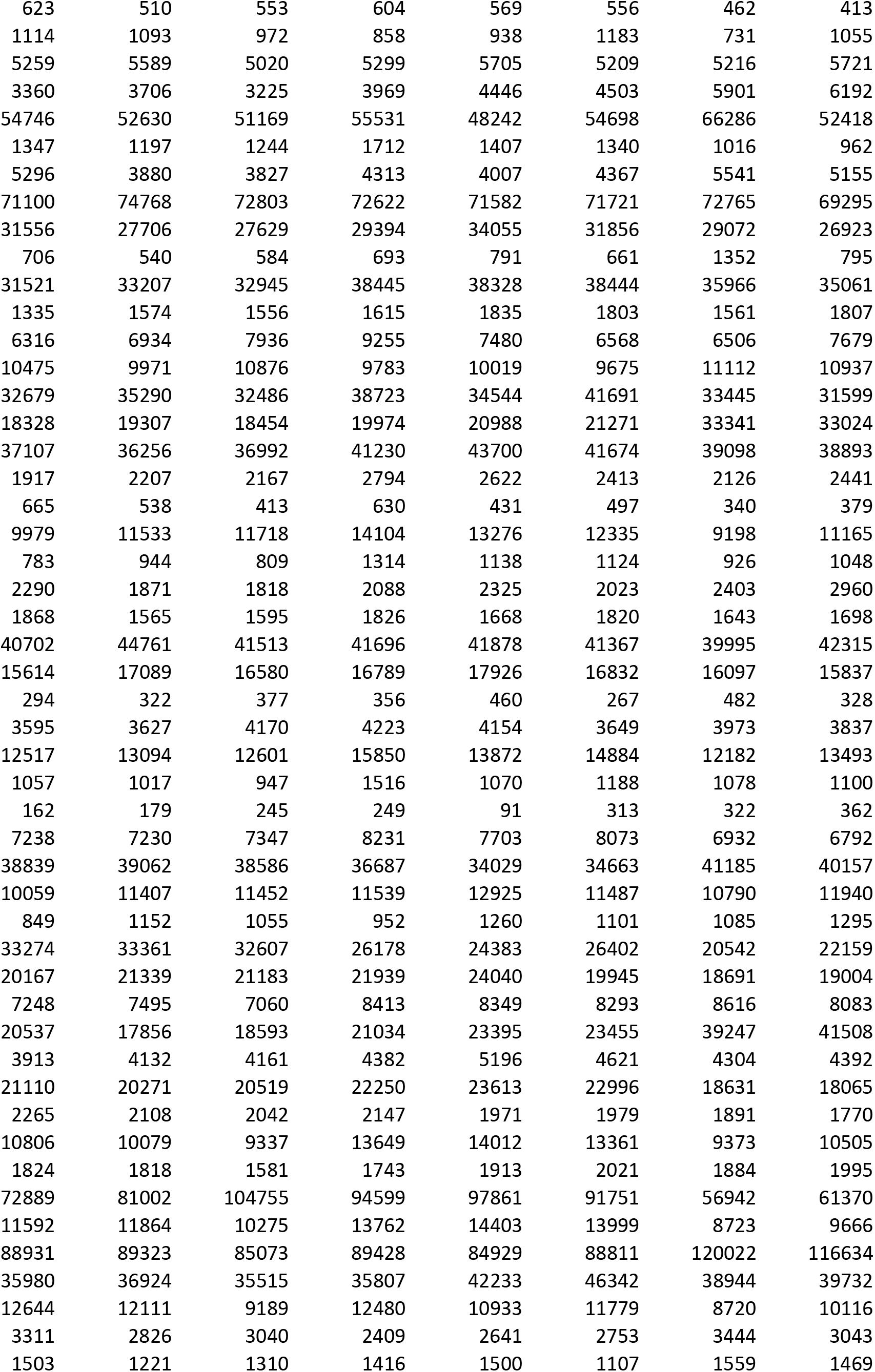
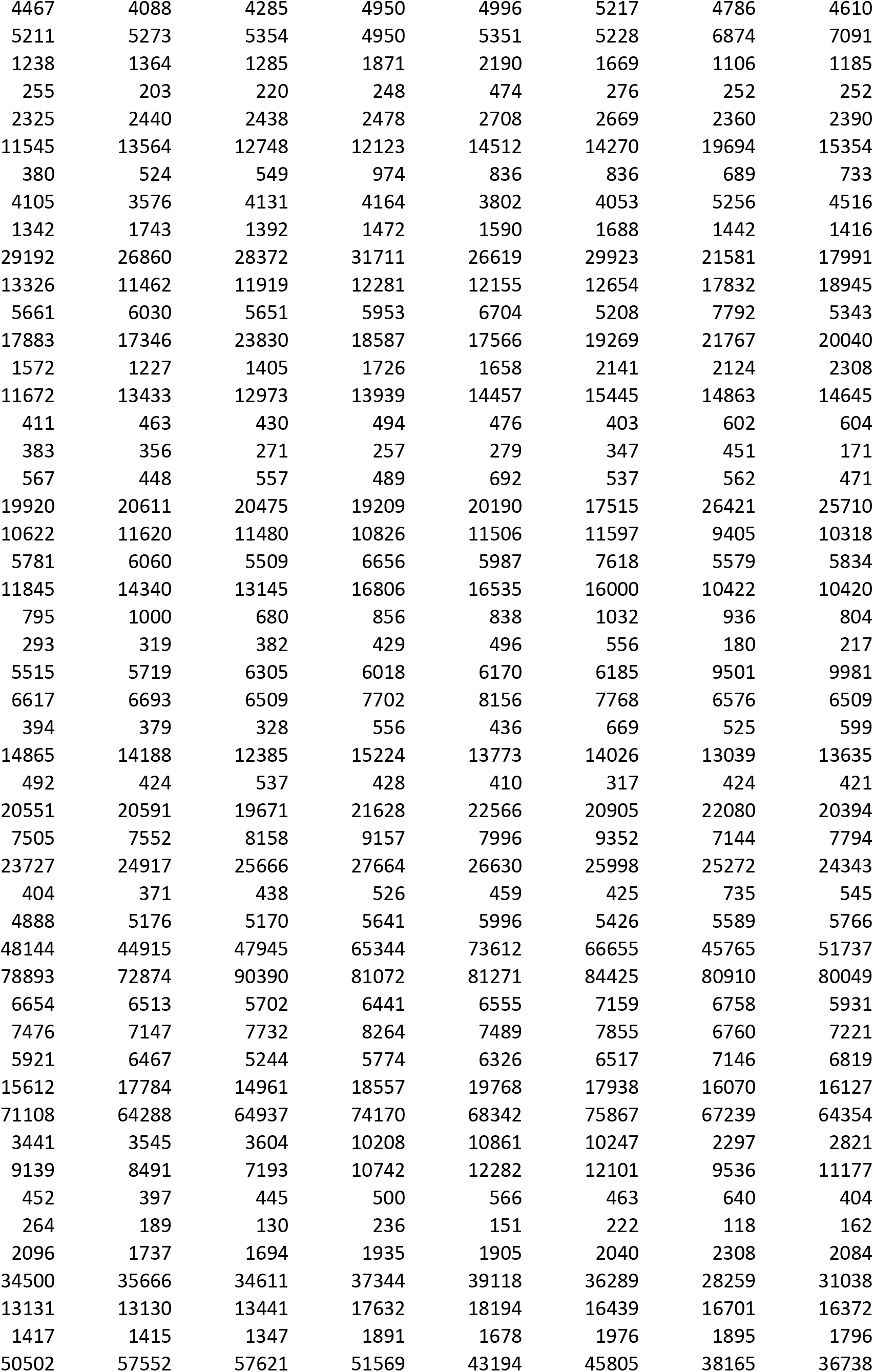
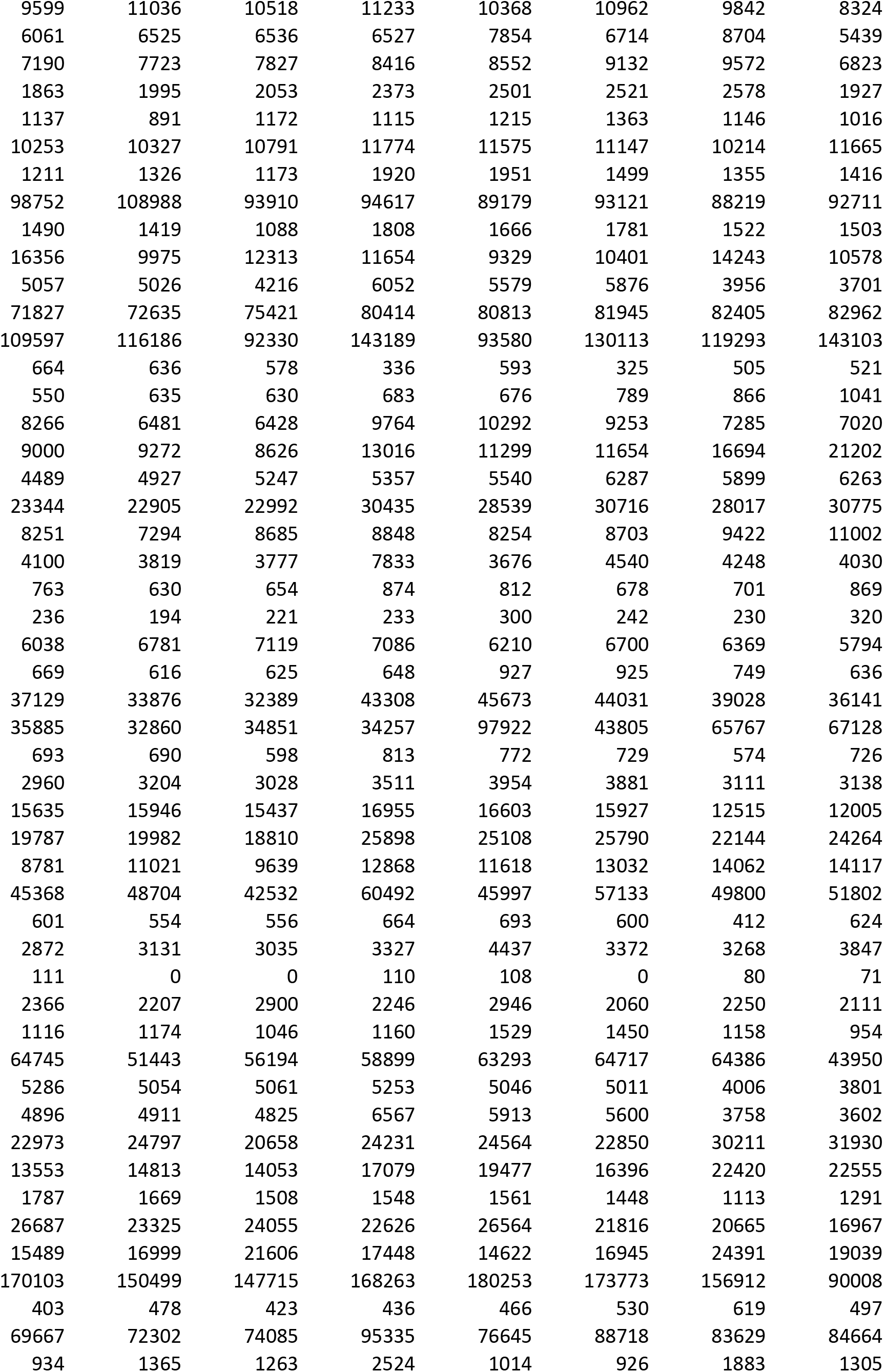
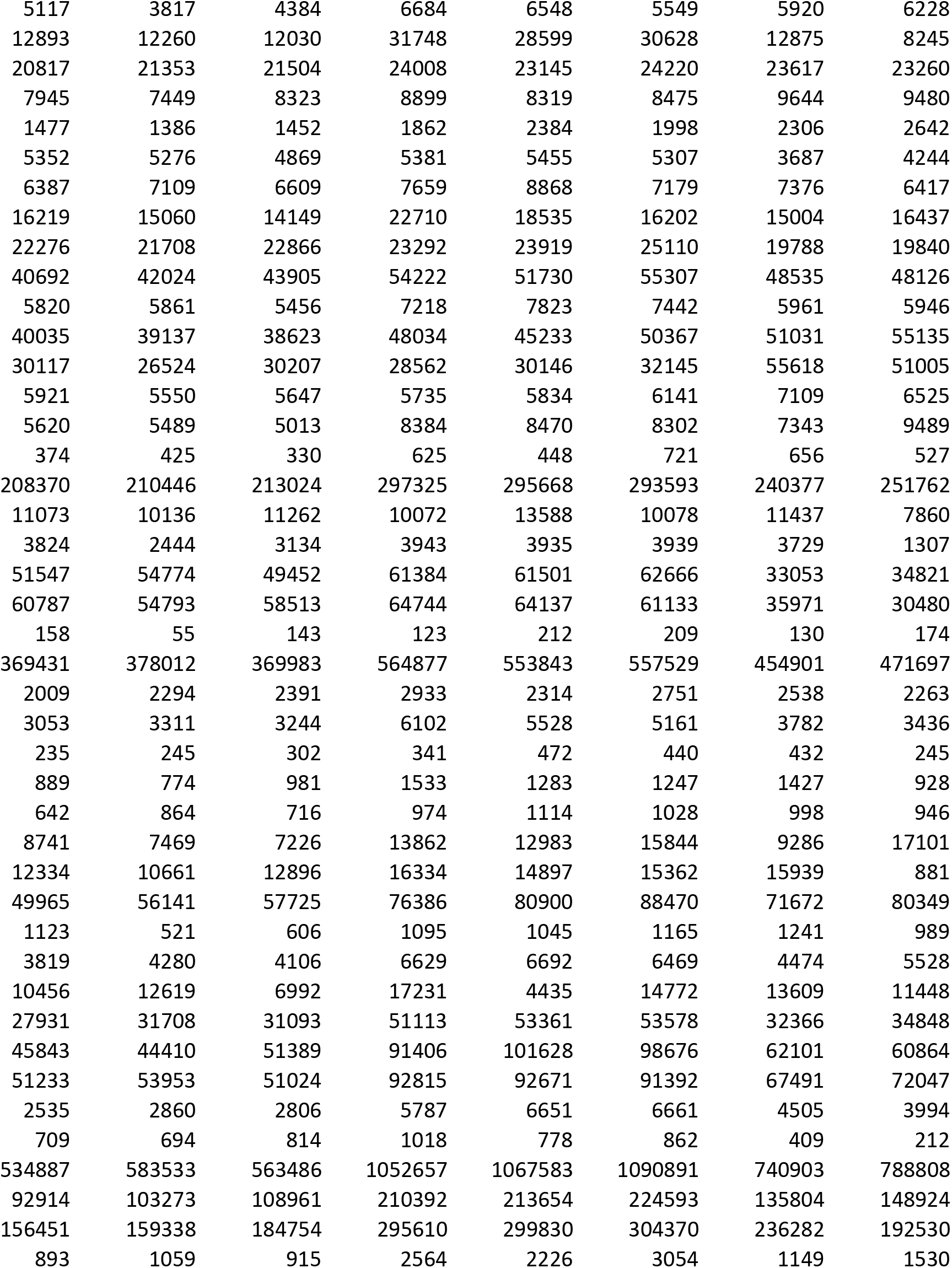
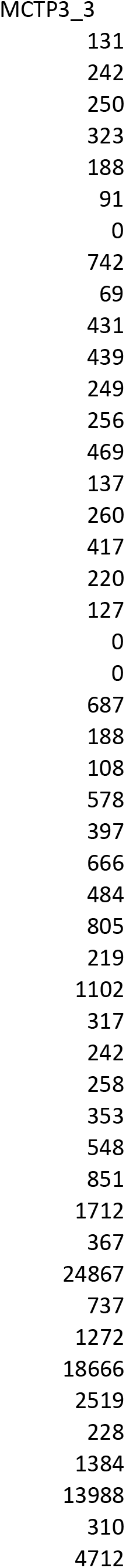
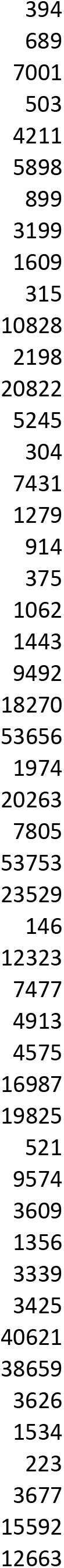
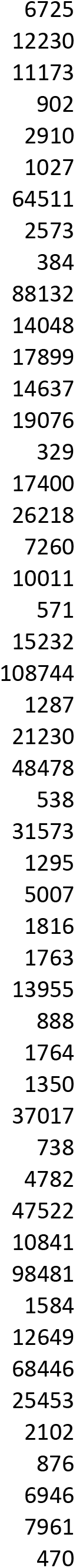
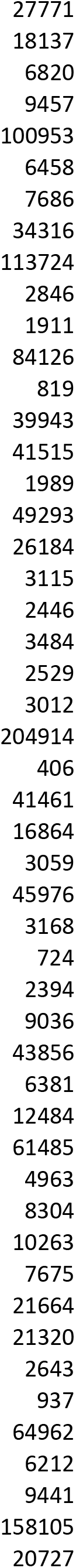
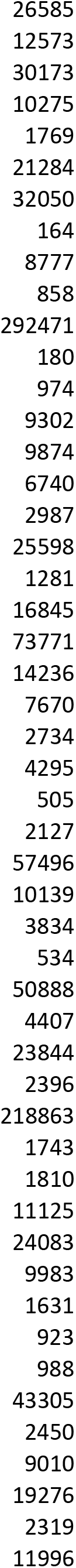
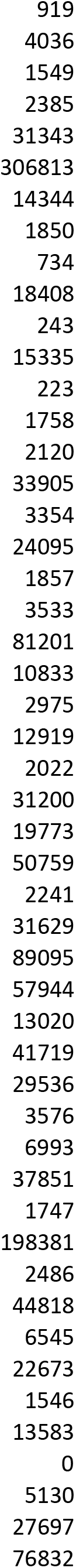
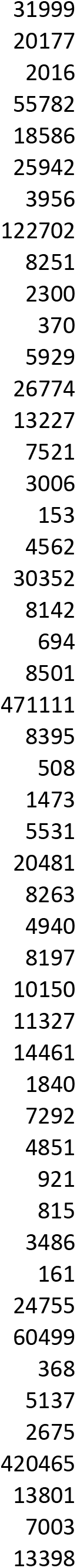
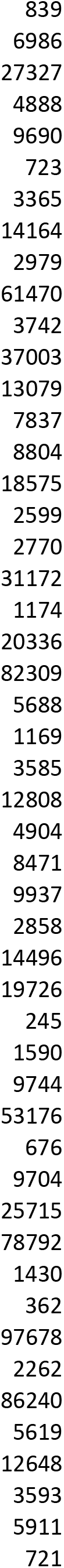
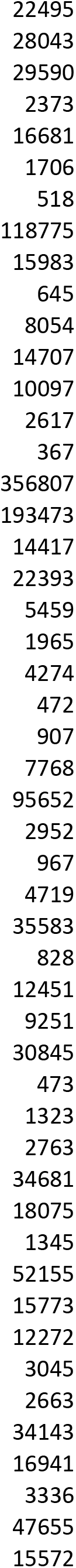
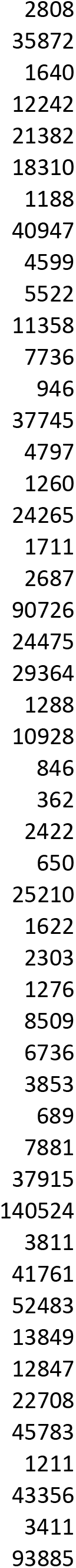
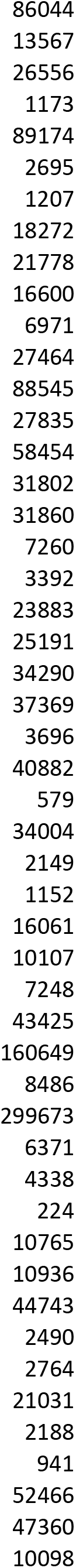
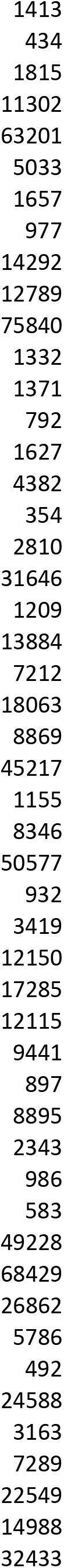
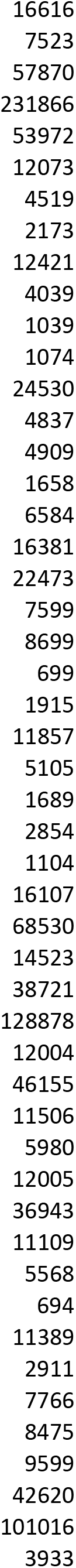
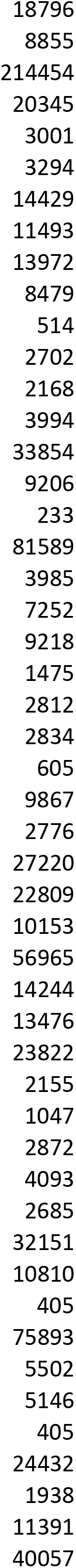
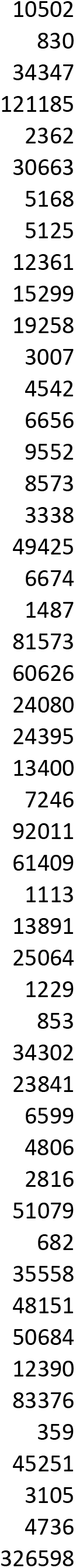
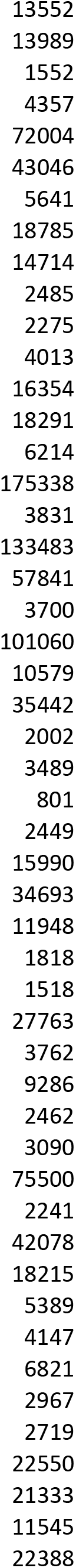
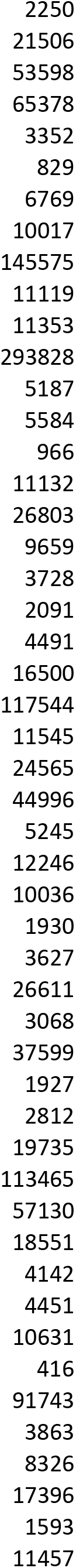
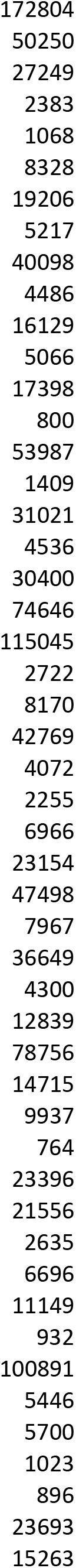
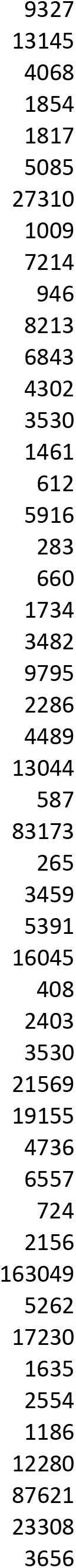
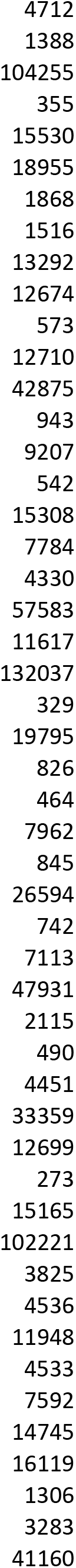
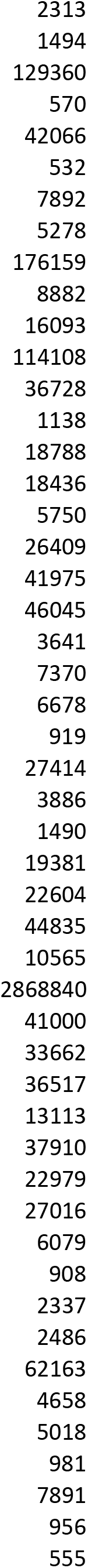
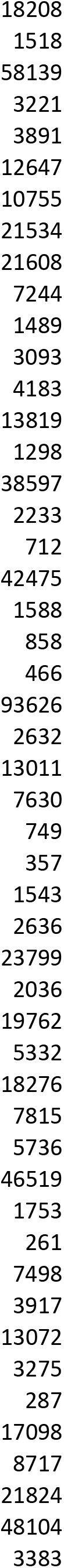
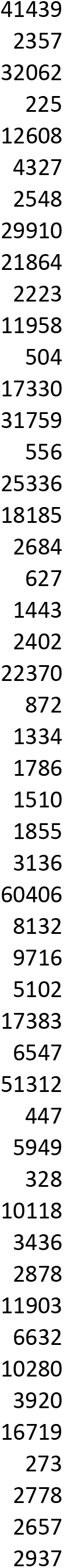
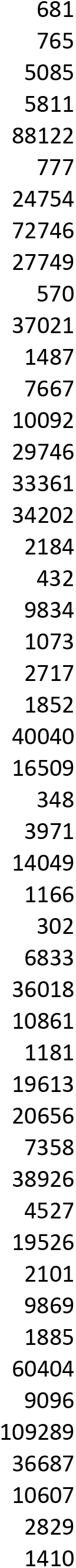
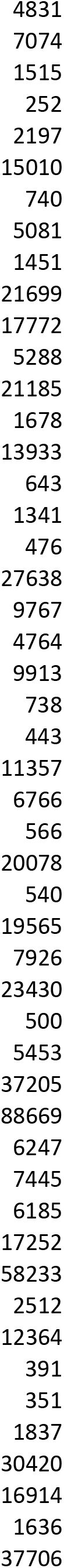
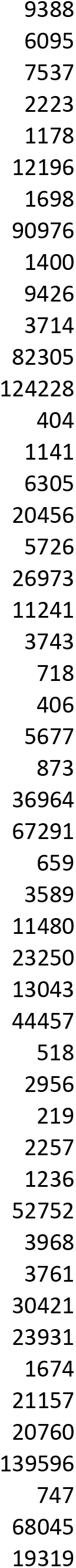
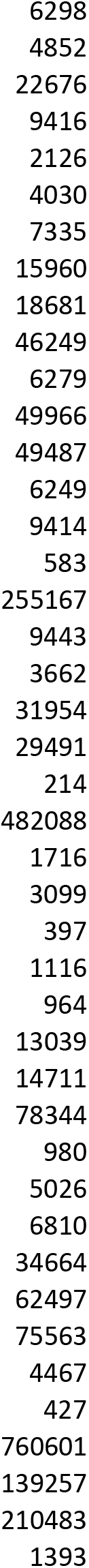

